# The genetic architecture of Parkinson disease in Spain: characterizing population-specific risk, differential haplotype structures, and providing etiologic insight

**DOI:** 10.1101/609016

**Authors:** Sara Bandres-Ciga, Sarah Ahmed, Marya S. Sabir, Cornelis Blauwendraat, Astrid D. Adarmes-Gómez, Inmaculada Bernal-Bernal, Marta Bonilla- Toribio, Dolores Buiza-Rueda, Fátima Carrillo, Mario Carrión-Claro, Pilar Gómez-Garre, Silvia Jesús, Miguel A. Labrador-Espinosa, Daniel Macias, Carlota Méndez-del-Barrio, Teresa Periñán-Tocino, Cristina Tejera-Parrado, Laura Vargas-González, Monica Diez-Fairen, Ignacio Alvarez, Juan Pablo Tartari, María Teresa Buongiorno, Miquel Aguilar, Ana Gorostidi, Jesús Alberto Bergareche, Elisabet Mondragon, Javier Ruiz-Martínez, Oriol Dols-Icardo, Jaime Kulisevsky, Juan Marín-Lahoz, Javier Pagonabarraga, Berta Pascual-Sedano, Mario Ezquerra, Ana Cámara, Yaroslau Compta, Manel Fernández, Rubén Fernández-Santiago, Esteban Muñoz, Eduard Tolosa, Francesc Valldeoriola, Isabel Gonzalez-Aramburu, Antonio Sanchez Rodriguez, María Sierra, Manuel Menéndez-González, Marta Blazquez, Ciara Garcia, Esther Suarez-San Martin, Pedro García-Ruiz, Juan Carlos Martínez-Castrillo, Lydia Vela-Desojo, Clara Ruz, Francisco Javier Barrero, Francisco Escamilla-Sevilla, Adolfo Mínguez-Castellanos, Debora Cerdan, Cesar Tabernero, Maria Jose Gomez Heredia, Francisco Perez Errazquin, Manolo Romero-Acebal, Cici Feliz, Jose Luis Lopez-Sendon, Marina Mata, Irene Martínez Torres, Jonggeol Jeffrey Kim, Janet Brooks, Sara Saez-Atienzar, J Raphael Gibbs, Rafael Jorda, Juan A. Botia, Luis Bonet-Ponce, Karen E Morrison, Carl Clarke, Manuela Tan, Huw Morris, Connor Edsall, Dena Hernandez, Javier Simon-Sanchez, Mike A Nalls, Sonja W. Scholz, Adriano Jimenez-Escrig, Jacinto Duarte, Francisco Vives, Raquel Duran, Janet Hoenicka, Victoria Alvarez, Jon Infante, Maria José Marti, Jordi Clarimón, Adolfo López de Munain, Pau Pastor, Pablo Mir, Andrew Singleton, on behalf of the International Parkinson Disease Genomics Consortium

## Abstract

**Background:** The Iberian Peninsula stands out as having variable levels of population admixture and isolation, making Spain an interesting setting for studying the genetic architecture of neurodegenerative diseases.

**Objectives:** To perform the largest Parkinson disease (PD) genome-wide association study (GWAS) restricted to a single country.

**Methods:** We performed a GWAS for both risk of PD and age-at-onset (AAO) in 7,849 Spanish individuals. Further analyses included population-specific risk haplotype assessments, polygenic risk scoring through machine learning, Mendelian randomization of expression and methylation data to gain insight into disease-associated loci, heritability estimates, genetic correlations and burden analyses.

**Results:** We identified a novel population-specific GWAS signal at *PARK2* associated with AAO. We replicated four genome-wide independent signals associated with PD risk, including *SNCA, LRRK2, KANSL1/MAPT* and *HLA-DQB1*. A significant trend for smaller risk haplotypes at known loci was found compared to similar studies of non-Spanish origin. Seventeen PD-related genes showed functional consequence via two-sample Mendelian randomization in expression and methylation datasets. Long runs of homozygosity at 28 known genes/loci were found to be enriched in cases versus controls.

**Conclusions:** Our data demonstrate the utility of the Spanish risk haplotype substructure for future fine-mapping efforts, showing how leveraging unique and diverse population histories can benefit genetic studies of complex diseases. The present study points to *PARK2* as a major hallmark of PD etiology in Spain.

## INTRODUCTION

Parkinson disease (PD) is a complex disorder arising from the interplay of polygenic risk, environment and stochastic factors occurring in an unpredictable manner ^1^. Over the last 20 years, extensive work in molecular genetics has dissected the underlying genetic cause of several familial and early-onset patients in which disease was inherited in a Mendelian fashion. However, while just a small percentage of PD cases are monogenic, often exhibiting variable penetrance, the vast majority are considered to be sporadic with complex genetic influence.

An important step forward in favor of a genetic contribution to the etiology of idiopathic PD has been taken from genome-wide association studies (GWAS). The implementation of a large-scale, unbiased approach aimed at identifying genetic susceptibility factors has substantially improved our understanding of the pathogenic pathways relevant to disease. To date, 90 loci have been associated with idiopathic PD ^2^. Yet, despite great advances in the field, only a small proportion of the heritable component of PD has been mapped. Total estimates of genetic variation attributed to PD are roughly 21% ^3^.

GWAS meta-analyses have been crucial to identifying and expanding the biological knowledge of novel disease risk factors. However, one of the main challenges is the heterogeneity across cohorts that might mask genetic associations specific to certain populations. Therefore, establishing population□specific catalogues of genetic variation when studying the genetic basis of PD is necessary.

Separated from the rest of Europe by the Pyrenees range of mountains and just 700 miles from the North coast of Africa, the Iberian Peninsula represents a cross-link between two continents and stands out as having remarkably variable levels of admixture, which has been reinforced by linguistic and geopolitical boundaries within the territory ^4^. The Spanish population has a more diverse haplotypic structure in comparison with other European populations and is somehow isolated within itself in terms of global genetic structure ^5^. The long-lasting migratory influences since early centuries and ulterior admixture from a number of civilizations over recent history have left their genetic imprint, thereby creating a particular genome population structure and diversity ^6^. Taken together, these observations make Spain an interesting setting to comprehensively study the genetic architecture of PD and other complex diseases.

Here, we have performed the largest PD genome-wide assessment from a single country to date, by characterizing 7,849 Spanish individuals. We compared our new Spanish dataset with extant data from various European ancestry populations to relate our findings in the context of larger studies. We also analyzed risk profiles, heritability and autozygosity in this population as it relates to PD etiology. Of particular interest are our analyses leveraging the unique population genetic structure of Spain to identify smaller risk haplotype blocks than in less admixed Europeans. We envisage that the data generated from this large study, together with other concomitant efforts underway in other European populations, will be key to shed light on the molecular mechanisms involved in the disease process and might pave the way for future therapeutic interventions.

## METHODS

### Cohort characteristics

A total of 7,849 individuals (4,783 cases and 3,066 neurologically healthy controls) were recruited from 13 centers across Spain. PD patients were diagnosed by expert movement disorders neurologists following the standard criteria of the United Kingdom PD Society Brain Bank ^7^. The respective ethical committees for medical research approved involvement in genetic studies and all participants gave written informed consent. Controls were extensively assessed to rule out any sign of neurological condition. Detailed demographic characteristics of each Spanish sub-cohort are summarized in **Supplementary Table 1,** with an average age at onset (AAO) for cases of 61.23 +/− 11.47, age at recruitment for controls of 61.79 +/− 11.05, 42.27 % of female cases and 56.06 % female controls.

### Genotyping and quality control analyses

Samples were genotyped using the customized NeuroChip Array v.1.0 or v.1.1 (Illumina) ^8^. This array contains a backbone consisting of 306,670 tagging variants (Infinium HumanCore-24 v1.0) that densely cover ancestry informative markers, markers for determination of identity by descent (IBD) and X chromosome single nucleotide polymorphisms (SNPs) for sex determination. Additionally, NeuroChip contains a custom content consisting of 179,467 neurodegenerative disease-related variants.

Genotypes were clustered using Illumina GenomeStudio v.2.0. Quality control (QC) analysis was performed as follows: samples with call rates of less than 95% and whose genetically determined sex from X chromosome heterogeneity did not match that from clinical data were excluded from the analysis. Samples exhibiting excess heterozygosity estimated by an F statistic > +/− 0.25 were also excluded. Once preliminary sample-level QC was completed, SNPs with minor allele frequency (MAF) < 0.01, Hardy-Weinberg Equilibrium (HWE) p-value < 1E-5 and missingness rates > 5 % were excluded. Genetic variants passing QC numbered 433,768 SNPs. Genotyped SNPs thought to be in linkage disequilibrium (LD) in a sliding window of 50 adjacent SNPs which scrolled through the genome at a rate of 5 overlapping SNPs were also removed from the following analyses (as were palindromic SNPs). Next, samples were clustered using principal component analysis (PCA) to evaluate European ancestry as compared to the HapMap3 CEU/TSI populations (<u>International HapMap Consortium, 2003</u>) (**Supplementary Figure 1**). Confirmed European-ancestry samples were extracted and principal components (PCs) 1-20 were used as covariates in all analysis. Samples related at the level of cousins or closer (sharing proportionally more than 18.5 % of alleles) were dropped from the following analysis. After filtering, 4,639 cases and 2,949 controls remained. The data were then imputed using the *Haplotype Reference Consortium r1.1 2016* (http://www.haplotype-reference-consortium.org), under default settings with phasing using the EAGLE option. Imputed variants numbered 9,911,207 after filtering for MAF > 1% and imputation quality (RSQ) > 0.3.

### Genome-wide association study versus Parkinson disease, age at onset and risk haplotype structure

To estimate risk associated with PD, imputed dosages were analyzed using a logistic regression model adjusted for sex, AAO for cases or examination for controls and the first 20 PCs as covariates. Summary statistics were generated using the RVTESTS package ^9^ and filtered for inclusion after meeting a minimum imputation quality of 0.30 and minor allele frequency greater than 1 %. To explore the influence of genetic variation on the AAO of PD cases, a linear regression model, adjusted for the same covariates, was performed (using AAO as the outcome instead of as a covariate).

Given the history of admixture in Spain, we hypothesized that there might be smaller haplotype blocks at risk loci than in less admixed populations ^10^. We compared the size of the 90 independent risk haplotype blocks in Spanish cases with a British ancestry PD cohort composed of 1,478 cases. After standardizing both datasets with the same genotyped SNPs passing identical quality control in both datasets, we determined the size of the haplotype blocks in both populations by using PLINK 1.9 ^11^.

### Whole genome sequencing analysis

To dissect the novel GWAS signal associated with AAO identified in the Spanish population, we performed whole genome sequencing analyses in 5 out of 37 homozygous carriers of the *PARK2* signal. DNA concentration was determined by Qubit fluorescence and normalized to 20 ng/ul. One microgram total genomic DNA was sheared to a target size of 450 bp using the Covaris LE220 ultrasonicator. Library preparation was achieved using TruSeq DNA PCR-Free High Throughput Library Prep Kit and IDT for Illumina TruSeq DNA UD Indexes (96 Indexes, 96 Samples). Sequencing libraries were assessed for size distribution, absence of free adapters and adapter dimers on a Fragment Analyzer. Library quantitation was performed by quantitative PCR using KAPA Library Quantification Kit subsequent normalization to 4 nM. Libraries were clustered on v2.5 flowcell using Illumina cBot 2 System prior to sequencing on Illumina HiSeq X System using paired-end 150 bp reads. BCL files processed with alignment by ISAAC on HAS 2.2 and BAMs were used for quality control assessment of mean coverage, percent duplicates, percent bases > 20x coverage and percent noise sites.

### Genetic risk profiling

Polygenic risk score analysis for PD and AAO was performed as described in detail elsewhere ^12,13^. Briefly, a cumulative genetic risk score was calculated by using the R package PRSice2 ^14^. Permutation testing and p-value after LD pruning was used to identify best P thresholds in external GWAS data (training dataset derived from summary statistics from Nalls et al. 2019 ^2^, excluding Spanish samples) to construct the PRS, allowing us to utilize variants below the common GWAS significance threshold of 5E-08. LD clumping was implemented under default settings (window size = 250kb, r^2^ > 0.1) and using the Spanish dataset (testing dataset), 10,000 permutations were applied to generate empirical P estimates derived P threshold ranging from 5E-08 to 0.5, at a minimum increment of 5E-08. Each permutation test provided a Nagelkerke’s pseudo r^2^ after adjustment for an estimated prevalence of 0.5%, study-specific eigenvectors 1-20, AAO for cases or examination for controls and sex as covariates. GWAS derived P threshold with the highest pseudo r^2^ was selected for further analysis.

### Machine Learning to predict disease status

Summary statistics from the most recent meta-analysis excluding Spanish cohort samples ^2^ were used for initial SNP selection in our Machine Learning (ML) analyses. This analysis utilizes an upcoming software package (GenoML, https://genoml.github.io), an automated Machine Learning tool that optimizes basic machine learning pipelines for genomic data. We used PD GWAS full summary statistics in the ML feature selection process from which we removed samples present in the Spanish PD cohort to avoid any circularity. Prior to analyses, we filtered the Spanish cohort genotype data for MAF > 1% and imputation quality > 0.8. Next, we randomly sampled without replacement 70% of the subjects for training the classifier and the other 30% for its validation. We opted to use PRSice ^14^ to pre-filter variants under default settings yielding 1,521 candidate variants from GWAS at a p-value threshold of 0.0005. In both the training and test sets, the dosage of each of the 1,521 variants per individuals is weighted by the GWAS beta. We then used a Caret ML [https://github.com/topepo/caret] framework with glmnet, xgbTree, xgbDART, xgbLinear and Random forest, with a grid search size of 30- and 10-fold cross validation on the training set. The algorithm maximizing mean area under the curve across iterations of cross-validation in the training set was then fit to the validation set to generate predictions and summary statistics.

### Heritability estimates and genetic correlations

The heritability of PD in the Spanish population was calculated using Linkage Disequilibrium Score Regression (LDSC) ^15^. This method has the ability to detect the contributions to disease risk of variants which do not reach genome significance but does not identify the specific variants contributing to disease risk. Using the same software, genetic correlations between PD and other catalogued GWAS studies were evaluated. The database, *LD Hub*, was used to screen overlapping genetic etiologies across 757 diseases/traits gathered from publicly available resources ^15^. Default settings were used in the analyses and final results were adjusted for multiple testing by using Bonferroni correction.

### Runs of homozygosity

Based on LD-pruned dataset (using previously described parameters), runs of homozygosity (ROHs) were defined using PLINK 1.9 ^11^. We explored ROHs containing at least 10 SNPs and a total length ≥ 1000 Kb, with a rate of scanning windows of at least 0.05 (not containing > 1 heterozygous call or 10 missing calls). In order to explore overall homozygosity between cases and controls, three metrics were assessed, including the number of homozygous segments spread across the genome, total kilobase distance spanned by those segments, and average segment size (autosomes only). Subsequently, ROHs were further investigated for known PD risk gene regions (**Supplementary Table 2**) and PD significant loci from GWAS ^2^ with a window of +/− 1 Mb upstream or downstream. In these analyses, cryptically related PD individuals removed in previous steps were included to identify overrepresented sharing of recessive regions among cases.

### Burden analyses

We examined the contribution of rare variation on disease risk by collapsing the cumulative effect of multiple genetic variants at a per-gene level. We performed the Sequence Kernel Association Test (SKAT) in imputed data after classifying variants into nested categories (based on two maximum minor allele frequency thresholds, (a) < MAF 1% or (b) < MAF 5%, and three functional filters, (1) non-coding variants, (2) only coding and (3) Combined Annotation Dependent Depletion (CADD) likely damaging). Burden analyses were adjusted for the first 20 PCs, AAO (cases) or examination (controls), and sex. These were run using default settings as part of the RVTESTS package ^9^. To adjust for multiple comparisons, we applied Bonferroni correction. Predictions of variant pathogenicity were obtained from ANNOVAR ^16^, based on the CADD algorithm (v1.3, http://cadd.gs.washington.edu) ^17^. In accordance with previous reports ^18^, we selected a stringent CADD C-score threshold ≥ 12.37, representing the top ∼2% most damaging of all possible nucleotide changes in the genome.

### Quantitative trait loci Mendelian randomization

Two-sample Mendelian randomization (MR) was performed to investigate possible functional genomic associations between PD known genes (**Supplementary Table 2**) and nominated loci ^2^ and expression or methylation quantitative trait loci (QTL) using summary statistics from this GWAS of the Spanish population to represent the outcome. Brain and blood QTL association summary statistics from well-curated methylation and expression datasets available via the SMR website (http://cnsgenomics.com/software/smr)^19^ were considered possible exposures in the MR models. These include estimates for methylation and cis-expression across multiple brain regions^20^. We also studied expression patterns in blood from the meta-analyses as described ^21^. Multi-SNP Summary-data-based MR was utilized to generate association estimates via MR between each QTL and local PD risk SNPs that contained 2 or more SNPs under default settings ^19^. The current Spanish dataset was used as a reference for linkage disequilibrium in this analysis. For each reference QTL dataset, p-values were adjusted by false discovery rate.

### Known genetic factors in Parkinson disease and atypical parkinsonism related genes

Considering both related and unrelated samples, we screened for relevant variants in known PD or parkinsonism-related genes. Our criteria included variants tagged as “disease causing mutation”, “possibly disease causing mutation”, “probably disease-associated polymorphism”, “disease-associated polymorphism with additional supporting functional evidence”, “pathogenic” or “possibly pathogenic” in ClinVar ^22^ or Human Gene Mutation Database ^23^. We followed two models: a putative dominant model (either the allele is present only in cases and absent in controls or with a lower frequency in controls) and a putative recessive model (two copies of the allele present only in cases and absent or with a lower frequency in controls). Genes screened are highlighted in **Supplementary Table 2**. We further screened putative structural genomic variation associated with PD in *PARK2* and *SNCA*, since notable genomic arrangements have been detected in these genes ^24,25^. Two metrics were assessed and visualized with R version 3.5.1 ^26^: B allele frequency and log R ratio. These two statistics allow visualization of copy number changes and are described in detail elsewhere ^27^.

## RESULTS

### Genome-wide association for Parkinson disease risk, age at onset and population-specific risk haplotype structure analyses

We identified four genome-wide independent signals associated with PD risk, including *SNCA, LRRK2, KANSL1/MAPT* and *HLA-DQB1* (**Figure 1, Table 1, Supplementary Figure 2**). Additionally, we show trends for association at an uncorrected p-value < 0.05 with 39 of the 90 loci previously identified in the largest PD meta-analysis performed to date ^2^ (**Supplementary Table 3**).

**Table 1.**
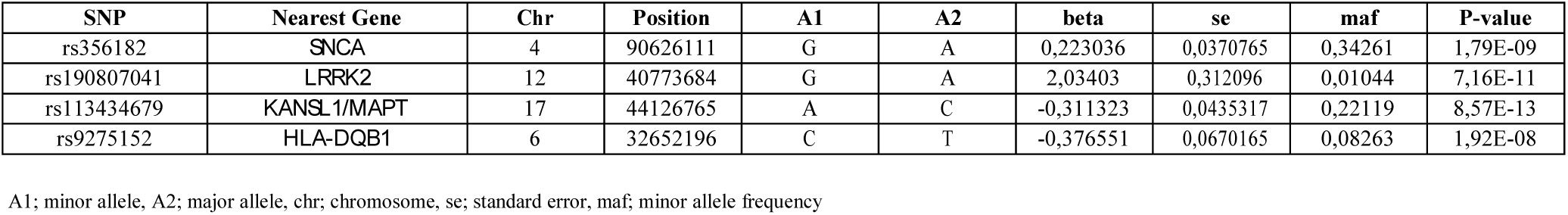
Genome-wide significant loci associated with PD risk in the Spanish population.

**Figure 1.**
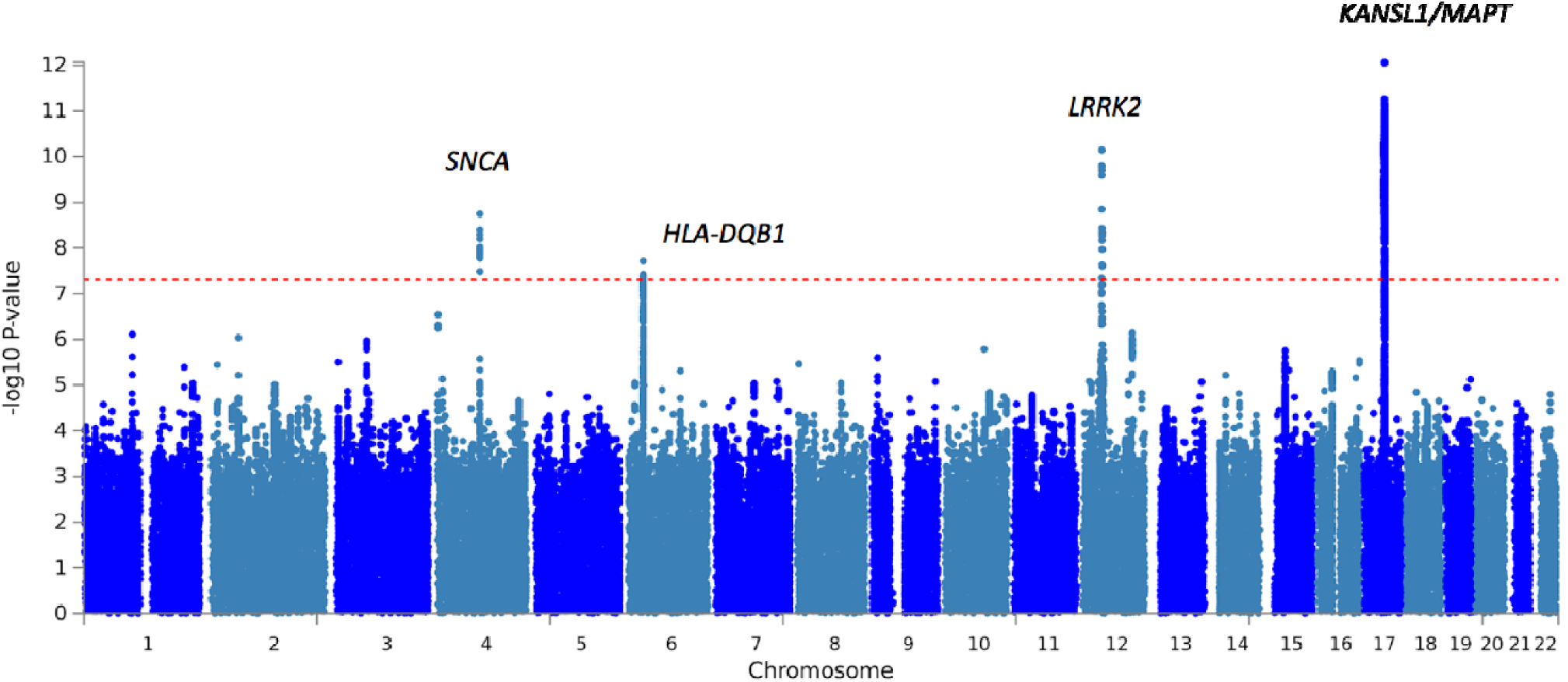
Manhattan plot showing results of PD GWA testing. Based on unrelated individuals (4,639 cases and 2,949 controls) using 9,945,565 SNPs. Four genome-wide significant loci were identified: *SNCA, LRRK2, HLA-DQB1* and *MAPT*.

GWA for AAO of PD revealed a genome-wide significant association signal at *PARK2* (rs9356013, beta= −4.11, se= 0.56, corrected p-value= 4.44 E-13) **(Figure 2, Supplementary Figure 3-4).** The association signal was only observed in the Spanish population and not in other International Parkinson’s Disease Genomics Consortium (IPDGC) datasets ^28^. A total of 37 cases carried the SNP in the homozygous state and 440 cases in the heterozygous state, which represents 9.5% of the Spanish PD cases. After removing rs9356013 homozygous carriers from the linear regression analysis, the signal dropped substantially (beta= −2.64, se = 0.74, p-value=3.41 E-4). The mean AAO for the rs9356013 homozygous carriers was 42.67 +/− 14.58 years while for the heterozygous carriers was 60.07 +/− 12.64 as compared to the AAO of the overall case series which was 61.23 +/− 11.47. Whole-genome sequencing analyses performed in 5 out of the 37 homozygous rs9356013 carriers revealed that all of them carried the deleterious frameshift mutation *PARK2* c.155delA (p.Asn52Metfs), which is located in exon 2 and 39 Kb away from the GWAS signal. Among them, four individuals carried the variant in the homozygous state and one in heterozygous state **(Supplemental Figure 5)**.

**Figure 2.**
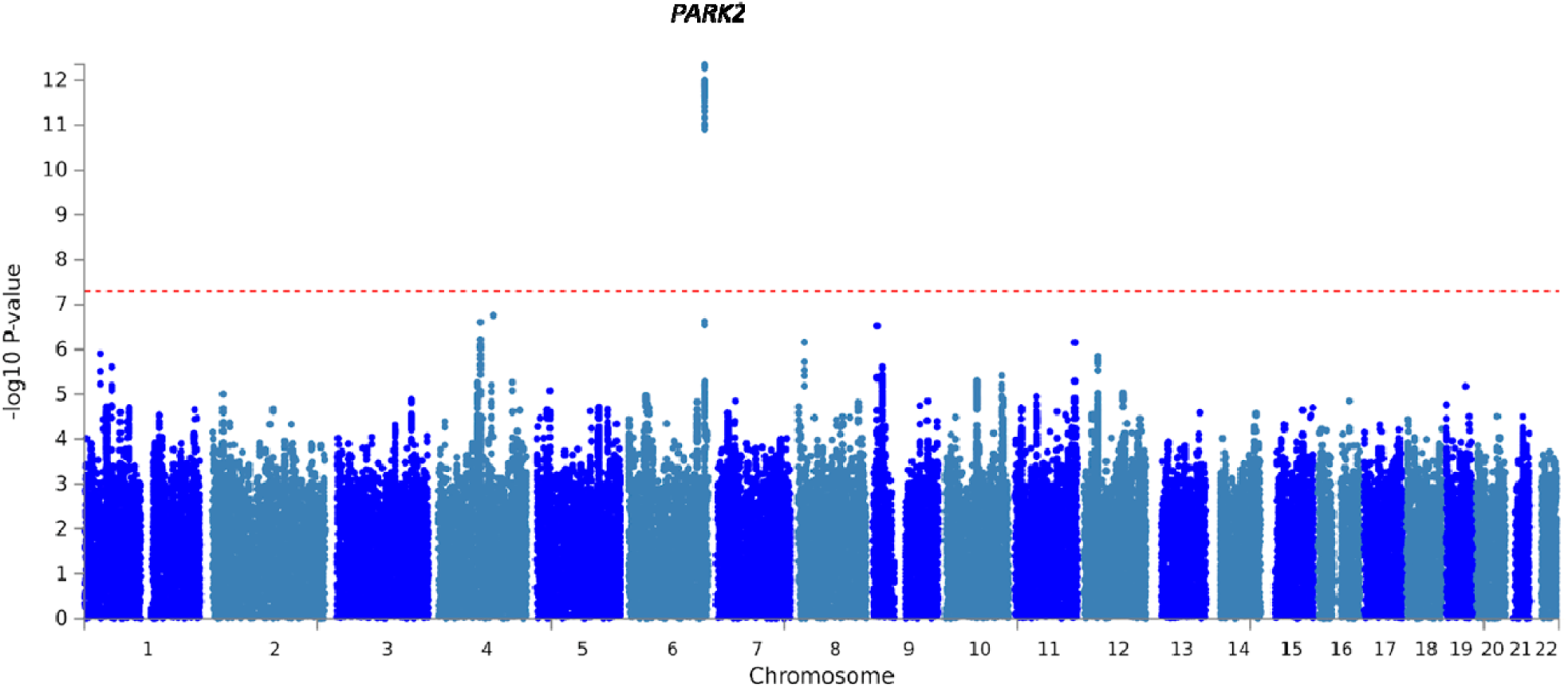
Manhattan plot showing results of PD GWA with age at onset testing. Based on 3,997 unrelated cases with available age at onset information using 9,945,565 SNPs. One genome-wide significant loci was identified: *PARK2.*

Sanger sequencing of the *PARK2* c.155delA variant (p.Asn52Metfs) in 1,275 PD cases followed by conditional analyses was performed to dissect whether the GWAS signal was tagging the indel. Conditional analyses for rs9356013 conditioning on c.155delA suggested that the common variant rs9356013 and the rare indel c.155delA were most likely dependent signals (rs9356013 linear model p-value = 1.13E-05; c.155delA linear model p-value = 3.77E-08; rs9356013 p-value after conditional analysis on c.155delA = 0.02, beta = −3.38, se= 1.46).

Haplotype size at risk loci in a matched cohort of British ancestry PD cases was 13.73 (+/− 32.9) Kb larger than the Spanish at overlapping loci. At consensus genotyped variants, a total of 11 risk haplotype blocks were smaller in the Spanish population than in the British, 23 were the same size, and 2 were larger **(Supplementary Table 4**). Other risk loci did not have multi-SNP genotyped haplotypes spanning top risk variants from external GWAS in both cohorts for comparison.

### Genetic risk profiling versus disease and age-at-onset

After adjusting for appropriate covariates and estimated PD prevalence, an overall pseudo r^2^ between PRS and PD was approximately r^2^=0.026. For each standard deviation from the population mean of the PRS, risk was estimated to be an odds ratio of 1.667 (beta = 0.511, se = 0.027, p-value = 3.63E-79, empirical P after permutation 1.00E-4). This model incorporated a total of 665 SNPs up to p-value < 5.99E-05 in the current GWAS **(Figure 3A-B)**.

**Figure 3.**
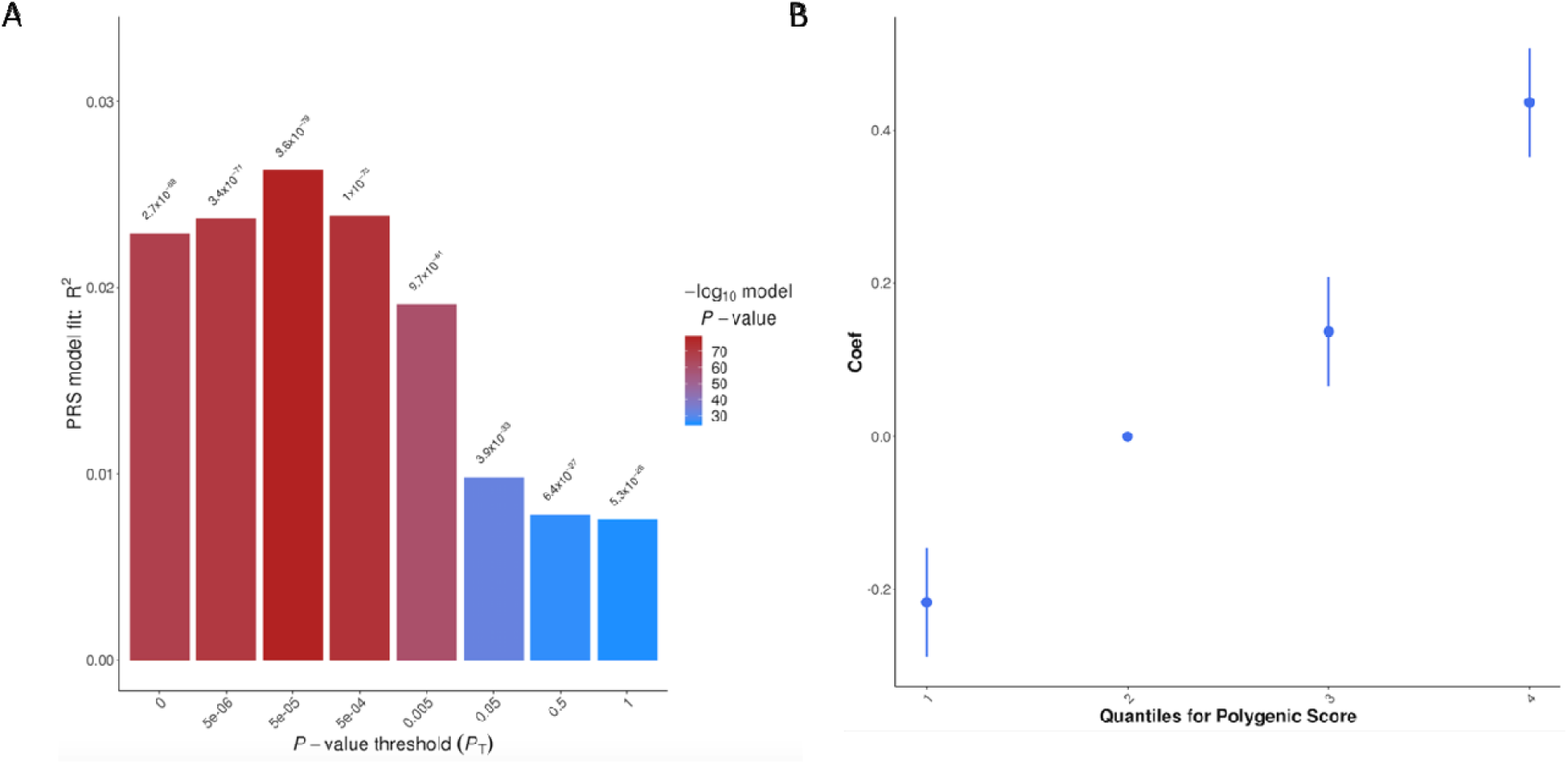
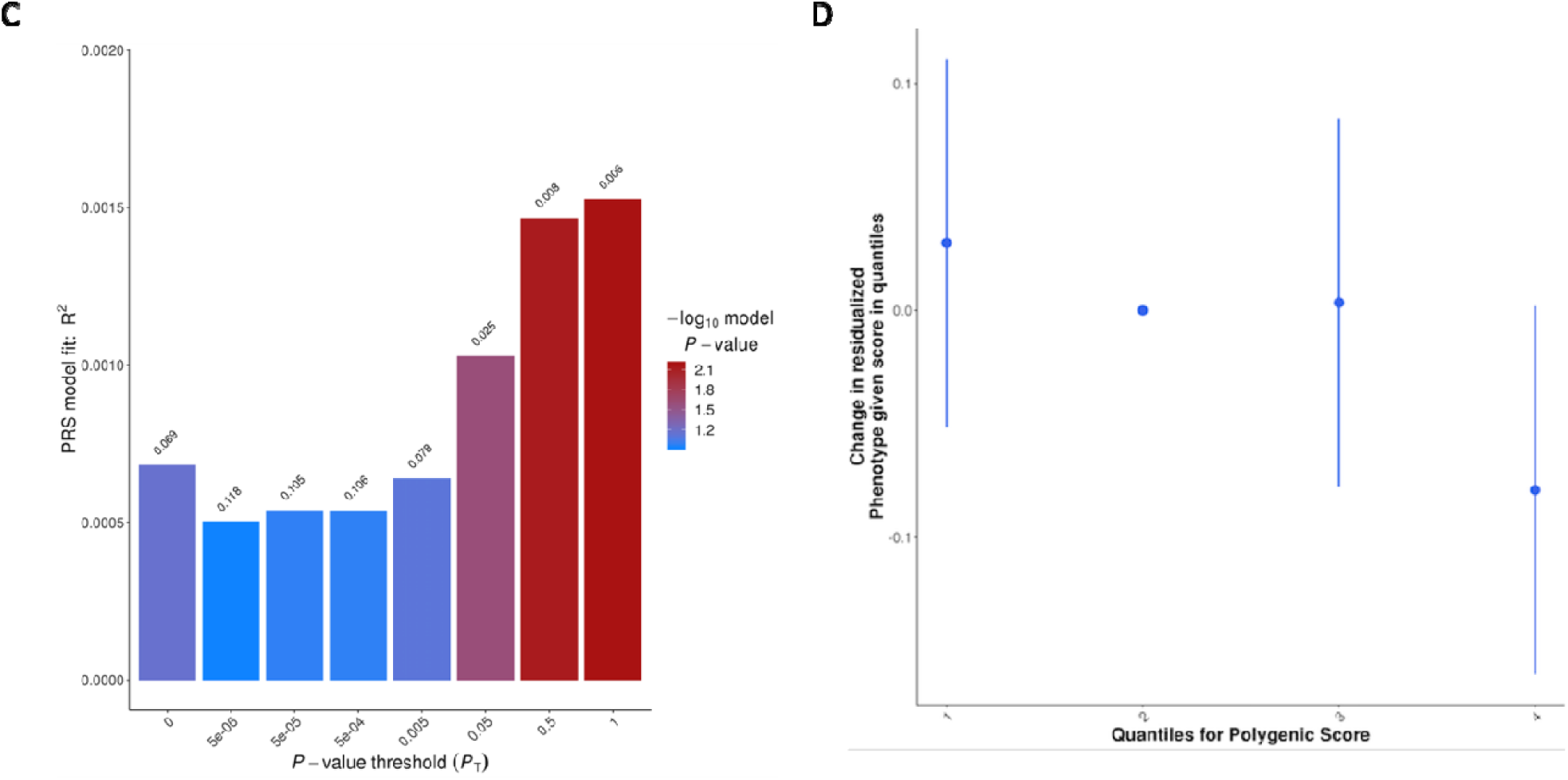
Polygenic risk score versus disease status and age at onset. **A.** Polygenic risk score versus disease status. R^2^ estimates at various p-value thresholds. **B.** Odds ratios by quantile of PD polygenic risk score. **C.** Polygenic risk score versus age at onset. R^2^ estimates at various p-value thresholds**. D.** Odds ratios by quantile of age at onset polygenic risk score.

Similar models were generated for AAO. We observed an association between a one standard deviation increase in the age at onset polygenic score and a near one year earlier onset of disease (beta = −0.944, se = 0.346, p-value = 0.006, empirical P after permutation = 0.031) This model utilized all unlinked variants of interest, pruned down to 271,191 SNPs **(Figure 3C-D)**.

### Machine learning to predict disease status

The xgbDART algorithm ^28^ (extreme gradient boosting approach with additive linear regression trees regularized by drop-out) yields the best AUC as predictor of the probability PD in this Spanish GWAS cohort. The model in the validation set had an AUC of 0.6205 with a sensitivity and specificity of 0.86 and 0.24. This is a ∼1% improvement over using simply polygenic risk scores prediction with PRSice alone, even after employing a two-stage design compared to a previous single phase that could be more overfit than what is presented here.

### Heritability estimates and genetic correlations

SNP heritability estimates by LDSC were estimated to be 28.67 % ± 6.65%. We analyzed cross-trait genetic correlations between PD and 750 other GWAS datasets of interest curated by *LD hub* ^15^. No genetic correlations remained significant after adjusting for multiple testing via false discovery rate. However, when considering an un-adjusted p-value < 0.05, negative correlations were found for body mass index related traits, smoking and alcohol intake, and positive correlations were identified for allergies and physical activity, among others **(Supplementary Table 5).**

### Runs of homozygosity

PD cases in our dataset were shown to have longer ROHs than controls, both with regard to the percentage of the genome within these runs and the average run size. For every 10 Mb increase in ROHs per sample, we noted an odds ratio of 1.02 (beta = 0.02, se = 0.008, p-value = 0.0097), a small but significant increase. Average run size was also associated with PD risk at an odds ratio of 1.244 per 1 Mb increase in average run size (beta = 0.218, se = 0.068, p-value = 0.0013). The total numbers of these ROHs were not significantly different between cases and controls. This suggests that fewer large ROHs might be more closely associated with disease risk than many small ROHs.

We further explored extended runs of homozygosity in known Mendelian PD genes **(Supplementary Table 2)** and the 90 nominated risk loci in the last meta-analysis ^2^. Homozygosity was found enriched in cases versus controls at 28 genes/loci **(Supplementary Table 6)**. However, only a ROH in *PARK2* surpassed Bonferroni correction (p-value threshold = 0.001).

### Burden analyses

We explored the cumulative effect of multiple rare variants at a gene level by grouping them in different categories based on frequency and functionality. We found a significant enrichment of coding variants in the *LRRK2* gene in PD cases compared to controls (*p*-value =□4.51E-16). When we excluded p.G2019S carried by 2.8% of the PD subjects from the analysis, the risk of PD conferred by *LRRK2* is not significant (p-value=0.0611), suggesting that this variant is the main driver of the association. Rare coding variants in *PARK2* were found overrepresented in cases versus controls (*p*-value =□0.008), although the association did not surpassed Bonferroni correction (*p*-value threshold =□0.0002). When focusing only on non-coding variants, *GBA* displayed an association at a p-value = 0.0003 which in this case did not reach multiple testing correction (p-value threshold = 3.15E-6). Finally, when grouping the variants by CADD score, we did not find any prospective novel gene associated with PD in the Spanish population.

### Quantitative trait loci Mendelian randomization

After adjustment for false discovery rate, 17 PD-related genes/loci showed functional consequence via two-sample Mendelian randomization in expression and methylation datasets (**Supplementary Table 7**). Increased expression of *NSF* and *BST1* in blood and *KANSL1, WNT3, KAT8, CD38, HLA-DRB6, TMEM175, HLA-DRB6* and *CTSB* in brain, were found to be inversely associated with PD risk, while a positive risk association was found for *TMEM163, GAK,* and *HLA-DQA1* expression in brain. Disparate results were found across different probes tagging *DGK1*.

Methylation QTL Mendelian randomization analyses revealed 56 CpG sites linked to PD risk in brain after multiple test correction. Increased methylation of *ARHGAP27, TMEM175, CRHR1* and *GAK* was found to be positively associated with disease while *HLA-DRB5, IGSF9B, TMEM163, DGKQ,* showed a negative directionality versus PD risk. Disparate results were found across different probes tagging *KANSL1* and *HLA-DRB5*.

### Known genetic factors in Parkinson disease and atypical parkinsonism related genes

A total of 73 PD or parkinsonism variants annotated as possibly disease associated were identified with higher frequencies in the PD patient sample (see **Supplementary Table 8, Supplementary Figure 6).** Of the identified variants, 28.76% (21/73) were detected in genes responsible for autosomal dominant PD. A total of 19 variants were detected in *LRRK2;* 2.8 % (134/4,783) of the screened PD patients and 0.3 % (12/3,066) of the controls carry the *LRRK2* p.G2019S mutation in the heterozygous state, while one case carried the variant in the homozygous state (*p*□=□2.73×10^-15^; OR□=□8.05, se = 0.26). The *LRRK2* p.Arg1441Gly variant was identified in a case and the *LRRK2* p.Met1869Thr was found in five cases and one control.

Of the variants, 36.9 % (27/73) were identified within autosomal recessive PD genes, including 19 variants in *PARK2* and 8 variants in *PINK1.* 15.78 % of the *PARK2* cases carriers (18/114) and 5.5 % of the *PINK1* carriers were found in the homozygous state. Although NeuroChip was not able to detect any *PARK2* or *PINK1* compound heterozygous carriers, exonic rearrangements were detected in 0.96 % of the screened patients **(Supplementary Figures 7A-B).** *PARK2* deletions were identified among 34 patients, 26 in the heterozygous state and 8 in the homozygous. Duplications were identified in 12 patients.

17.8 % (13/73) of the variants were found in PD risk genes. Eleven *GBA* variants were detected among (175/4,783) patients. The *GBA* p.His490Arg, p.Val437Ile, p.Gly234Glu, p.Val54Leu, p.Lys13Arg, p.Leu29Alafs*18 and p.Leu363Pro variants were found overrepresented in cases versus controls but the association analysis did not reach statistical significance (**Supplementary Table 8).** A 1.1 % (53/4,783) of the patients and 0.3 % (12/3,066) of the controls under study carry the *GBA* p.Asn409Ser mutation in the heterozygous state. A 2 % (97/4,783) of the cases and 1.01 % (31/3,066) of the controls harbored the *GBA* p.Glu365Lys heterozygous polymorphism (*p*□=□0.0006; OR□=□2.005, se = 0.2). The *GBA* p.Asp448His variant was identified in 14 cases and 3 controls respectively (*p*□=□0.07; OR□=□2.99, se =0.6). Furthermore, the previously reported p.Asn613del in *MAPT* was identified in a 87 year old patient with an onset of rest tremor at 62 years and a notable family history of PD and progressive supranuclear palsy ^29^. Finally, fourteen variants were found in four atypical parkinsonism-related genes, including *FBXO7* and *POLG1* (autosomal recessive), as well as *ATP13A2* and *DCTN1* (autosomal dominant), however, their disease significance is uncertain.

## DISCUSSION

As part of a Spanish multicenter massive collaborative effort, we have gathered the largest collection of PD patients and controls from a single country to comprehensively assess the genetics of PD on a genome-wide scale. We have used the same genotyping platform, thus reducing possible batch effects. Here, we dissect population-specific differences in risk and AAO from a genetic perspective and highlight the utility of the Spanish risk haplotype substructure for future fine-mapping efforts.

In concordance with other populations ^2^, our Spanish GWAS on PD risk replicated four loci linked to disease, strengthening once again the role of *SNCA, LRRK2, KANSL1/MAPT* and *HLA-DQB1* in disease etiology. Of note, we identified for the first time in an European population an intronic signal in *PARK2* as a modifier of AAO. Conditional analysis showed a likely dependent effect with c.155delA, highlighting a higher frequency of this deleterious mutation in Spain compared to other populations. Given the shared ancestry, c.155delA has been described at high frequencies in the Iberian Peninsula ^30,31^ and has also been reported to be more common in the Latino population (GnomAD allele count = 39/35,440; frequency = 0.11%) versus Non-Finnish Europeans (GnomAD allele count =31/129,038; frequency = 0.024%).

Genetic testing can help to design an optimized trial with the highest likelihood of providing meaningful and actionable answers. Our study shows that Spain is a valuable resource for identifying and tracking *PARK2* c.155delA carriers to accelerate enrollment for target-specific PD clinical trials.

The fact that risk PD haplotypes are smaller in the Spanish population, in comparison to the less admixed British population, brings to light the importance of exhaustively studying diverse populations. The investigation of admixed populations in GWA studies has significant potential to accelerate the mapping of PD loci.

Importantly, we revealed an overall excess of homozygosity in PD cases versus controls and identified 28 genes/loci exhibiting ROH overrepresented in cases, pointing out the possible existence of disease-causing recessive variants that might be uncovered by future sequencing analysis. Additionally, burden analysis reinforced the contribution of both common and rare variants in *LRRK2*, making Spain an important candidate population for specific *LRRK2* clinical trials. In an effort to explore functional consequences associated to PD risk in the Spanish population, we performed quantitative trait loci Mendelian randomization analyses using expression and methylation data and suggest biological pathways underlying the nominated genes warrant further study.

Recent research has begun to demonstrate the utility of polygenic risk profiling to identify individuals who could benefit from the knowledge of their probabilistic susceptibility to disease, an aspect that is central to clinical decision-making and early disease detection ^32^. Here, we assessed the overall cumulative contribution of common SNPs on disease risk and age at onset. Our PRS derived model for disease risk and age at onset showed expected trends comparable to previous literature ^12,13,33^.

While we have made progress in assessing genetic risk factors for PD in a population-specific manner, there are a number of limitations to our study. First, although all the available PD cases and controls from Spain have been assessed, we are aware of the caveats driven by sample size. Dissection of additional susceptibility genetic risk and phenotypic relationships would have been possible if a larger cohort had been analyzed. In fact, the heritability estimate, of ∼ 28.67 % in this population, indicates that there is a large component of genetic risk yet to be uncovered. We assume there are a considerable number of variants that impact risk for disease outside the limits of what can be accurately detected with a genotyping platform. This could explain the lower observed frequency of certain well-established pathogenic variants and exonic rearrangements when comparing other sequencing studies previously performed in the Spanish population ^33,34.^

We have applied a state-of-the-art machine learning approach in an effort to predict disease status. Our results show that genetic data are not sufficient to accurately predict disease status in a clinical setting by itself when used alone, although this may change in the future when combining genetic with other biomarker data. Entering the era of personalized medicine in which an individual’s genetic makeup will help determine the most suitable therapy, we envisage our collaborative initiative will expand towards identifying, refining, and predicting heritable risk in the Spanish population by combining future large-scale whole-genome sequencing approaches, multi-omics and detailed longitudinal clinical data for translational approaches. We conclude by saying that this is the starting point of a collaborative network of Spanish clinicians and scientists that will continue to pave the road towards future therapeutic interventions.

## Funding agencies

This research was supported in part by the Intramural Research Program of the National Institutes of Health (National Institute on Aging, National Institute of Neurological Disorders and Stroke; project numbers: project numbers 1ZIA-NS003154-03, Z01-AG000949-02 and Z01-ES101986). In addition this work was supported by the Department of Defense (award W81XWH-09-2-0128), and The Michael J Fox Foundation for Parkinson’s Research, the ISCIII Grants PI 15/0878 (Fondos Feder) to VA and PI 15/01013 to JH. This study was supported by grants from the Spanish Ministry of Economy and Competitiveness [PI14/01823, PI16/01575, PI18/01898, [SAF2006-10126 (2006-2009), SAF2010-22329-C02-01 (2010-2012), and

SAF2013-47939-R (2013-2018)].], co-founded by ISCIII (Subdirección General de Evaluación y Fomento de la Investigación) and by Fondo Europeo de Desarrollo Regional (FEDER), the Consejería de Economía, Innovación, Ciencia y Empleo de la Junta de Andalucía [CVI-02526, CTS-7685], the Consejería de Salud y Bienestar Social de la Junta de Andalucía [PI-0437-2012, PI-0471-2013], the Sociedad Andaluza de Neurología, the Jacques and Gloria Gossweiler Foundation, the Fundación Alicia Koplowitz, the Fundación Mutua Madrileña. Pilar Gómez-Garre was supported by the “Miguel Servet” (from ISCIII16 FEDER) and “Nicolás Monardes” (from Andalusian Ministry of Health) programmes. Silvia Jesús Maestre was supported by the “Juan Rodés” programme and Daniel Macías-García was supported by the “Río Hortega” programme (both from ISCIII-FEDER). Cristina Tejera Parrado was supported by VPPI-US from the Universidad de Sevilla. This research has been conducted using samples from the HUVR-IBiS Biobank (Andalusian Public Health System Biobank and ISCIII-Red de Biobancos PT13/0010/0056). This work was also supported by the grant PSI2014-57643 from the Junta de Andalucía to the CTS-438 group and a research award from the Andalusian Society of Neurology.

## AUTHOR CONTRIBUTIONS

Data Acquisition or data contribution

Astrid D. Adarmes-Gómez, Inmaculada Bernal-Bernal, Marta Bonilla-Toribio, Dolores Buiza-Rueda, Fátima Carrillo, Mario Carrión-Claro, Pilar Gómez-Garre, Silvia Jesús, Miguel A. Labrador-Espinosa, Daniel Macias, Carlota Méndez-del-Barrio, Teresa Periñán-Tocino, Cristina Tejera-Parrado, Laura Vargas-González, Monica Diez-Fairen, Ignacio Alvarez, Juan Pablo Tartari, María Teresa Buongiorno, Miquel Aguilar, Ana Gorostidi Pagola, Jesús Alberto Bergareche Yarza, Elisabet Mondragon Rezola, Javier Ruiz-Martínez, Oriol Dols-Icardo, Jaime Kulisevsky, Juan Marín-Lahoz, Javier Pagonabarraga, Berta Pascual-Sedano, Ana Cámara, Yaroslau Compta, Manel Fernández, Rubén Fernández-Santiago, Maria Jose Marti, Esteban Muñoz, Eduard Tolosa, Francesc Valldeoriola, Isabel Gonzalez-Aramburu, Jon Infante, Antonio Sanchez Rodriguez, María Sierra, Manuel Menéndez-González, Marta Blazquez, Ciara Garcia, Esther Suarez-San Martin, Pedro García-Ruiz, Juan Carlos Martínez-Castrillo, Lydia Vela, Francisco Javier Barrero, Francisco Escamilla-Sevilla, Adolfo Mínguez-Castellanos, Debora Cerdan, Jacinto Duarte, Maria Jose Gomez Heredia, Francisco Perez Errazquin, Cici Feliz, Jose Luis Lopez-Sendon, Marina Mata, Irene Martínez Torres, Karen E Morrison, Carl Clarke, Manuela Tan, Huw Morris, Adriano Jimenez-Escrig, Cesar Tabernero, Francisco Vives, Raquel Duran, Janet Hoenicka, Victoria Alvarez, Jon Infante, Mario Ezquerra, Jordi Clarimón, Adolfo López de Munain Arregui, Pau Pastor, Pablo Mir.

Study level analysis and data management

Sara Bandres-Ciga, Sarah Ahmed, Marya Sabir, Cornelis Blauwendraat, Rafael Jorda, Juan A. Botia, Mike Nalls, Sonja Scholz, Connor Edsall, Dena Hernandez, Andrew Singleton.

Design and funding

Sara Bandres-Ciga, Cornelis Blauwendraat, Mike Nalls, Andrew Singleton

Writing – Original Draft

Sara Bandres-Ciga

Critical review and writing the manuscript

All authors

## ACKNOWLEDGMENTS

The authors are grateful to the participants in this study without whom this work would not have been possible.

