## Supplementary materials for "The genetic architecture of Parkinson disease in Spain: characterizing population-specific risk, differential haplotype structures, and providing etiologic insight"

**Supplementary Figure 1.** Spanish cohort with HapMap3 populations; Spanish cohort ancestry; Spanish cohort with African, Asian and European CEU and TSI populations…………………………………………………………………………………………….. 1

**Supplementary Figure 2.** Quantile-Quantile plot of GWA with PD risk……………………………………………………………………………. 3

**Supplementary Figure 3.** Quantile-Quantile plot of GWA with age at onset…………………………………………………………………. 3

**Supplementary Figure 4.** P-value correlation plot between PD GWAS and age of onset……………………………………………….. 4

**Supplementary Figure 5.** *Interactive Genome Viewer* (IGV) screen capture displaying the sequencing reads generated from five PD cases carrying the *PARK2* intronic variant (rs9356013)……………………………………………………………………………………. 5

**Supplementary Figure 6.** Cluster plots of genetic variants in PD and Parkinsonism related genes (overrepresented in cases) generated from Genome Studio……………………………………………………………………………………………………………………………. 6

**Supplementary Figure 7.** Copy number variation plots displaying *PARK2* deletions or duplications……………………………… 20

**Supplementary Table 1.** Demographic characteristics of the Spanish cohort………………………..…………………………………….. 32

**Supplementary Table 2.** Parkinson disease and Parkinsonism related genes under study…………………………………………….  33

**Supplementary Table 3.** Spanish GWAS replication results of known risk loci identified in Nalls et al., 2019………………..…. 34

**Supplementary Table 4**. Risk haplotype estimates at consensus genotyped variants for known GWAS loci in Spanish and UK PD cases……………………………………………………………………………………………………………………………………………………. 37

**Supplementary Table 5.** Cross-trait genetic correlations by linkage disequilibrium score regression…………………………….  40

**Supplementary Table 6.** Runs of homozygosity within PD known genes/loci identified in the Spanish population…………. 51

**Supplementary Table 7.** Significant functional associations for PD known/nominated genes via two-sample Mendelian randomization………………………………………………………………………………………………………………………………………….. 53
**Supplementary Table 8.** Genetic variants in PD and Parkinsonism related genes overrepresented in Spanish cases versus controls…………………………………………………………………………………………………………………………………………………. 55
Consortia members………………………………………………………………………………………………………………………………………… 57

**Supplementary Figure 1.** Spanish cohort with HapMap3 populations; Spanish cohort ancestry; Spanish cohort with African, Asian and European CEU and TSI populations

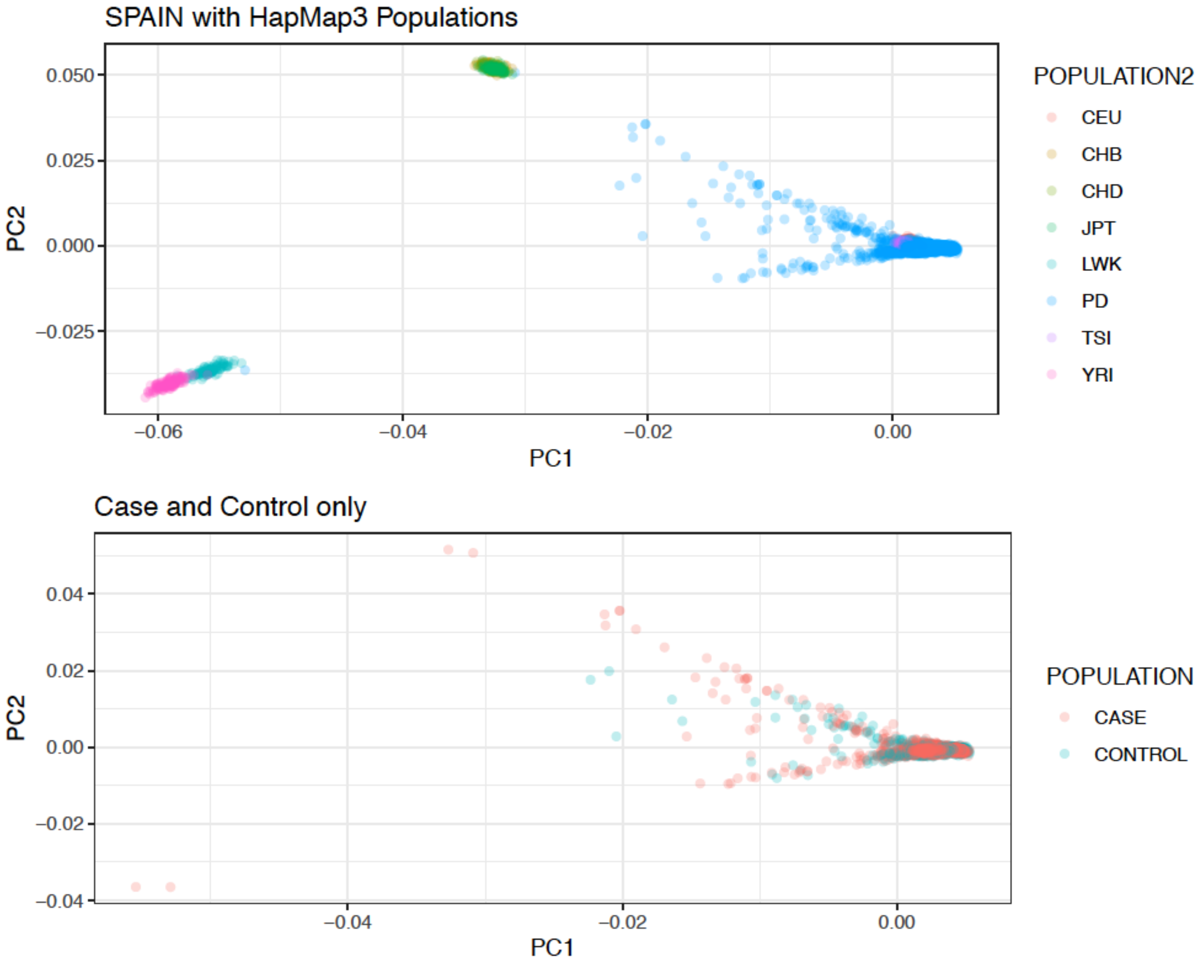

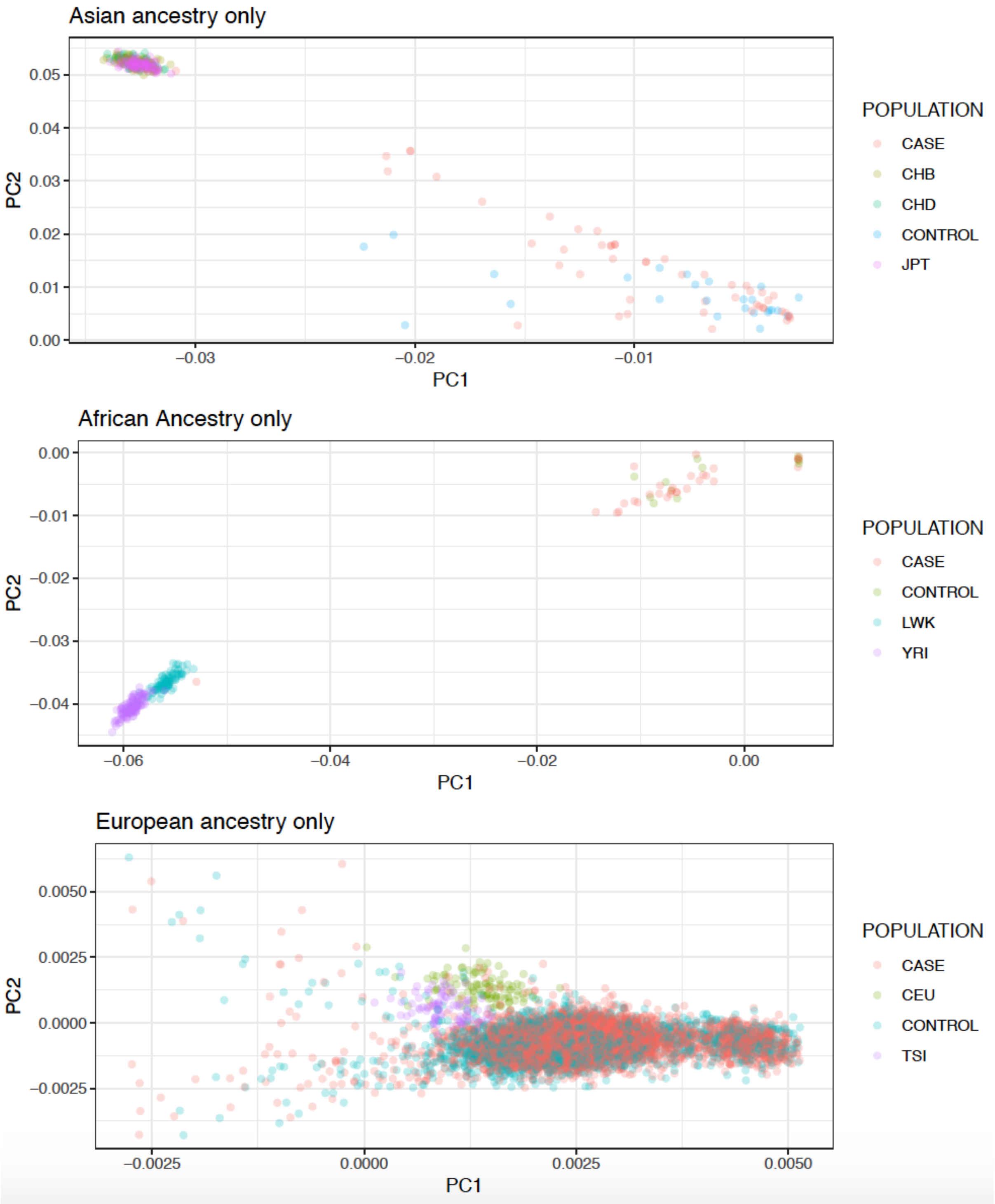
Asian ancestry only
PC2
0.05
0.04
0.03
0.02
0.01
0.00
−0.03
−0.02
−0.01
PC1
POPULATION
CASE
CHB
CHD
CONTROL
JPT
African Ancestry only
PC2
0.00
−0.01
−0.02
−0.03
−0.04
−0.06
−0.04
−0.02
0.00
PC1
POPULATION
CASE
CONTROL
LWK
YRI
European ancestry only
PC2
0.0050
0.0025
0.0000
−0.0025
−0.0025
0.0000
0.0025
0.0050
PC1
POPULATION
CASE
CEU
CONTROL
TSI

**Supplementary Figure 2.** Quantile-Quantile plot of GWA with PD risk

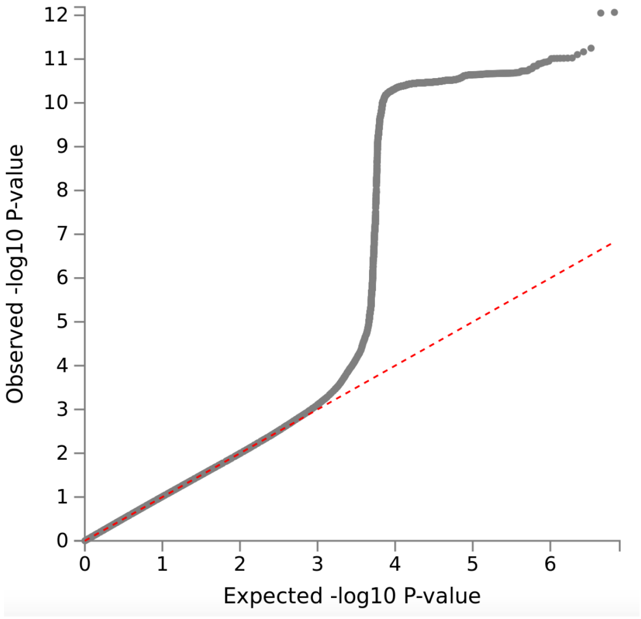

Genomic inflation factor (λ) = 1.038; λ1000=1.010

**Supplementary Figure 3.** Quantile-Quantile plot of GWA with age at onset

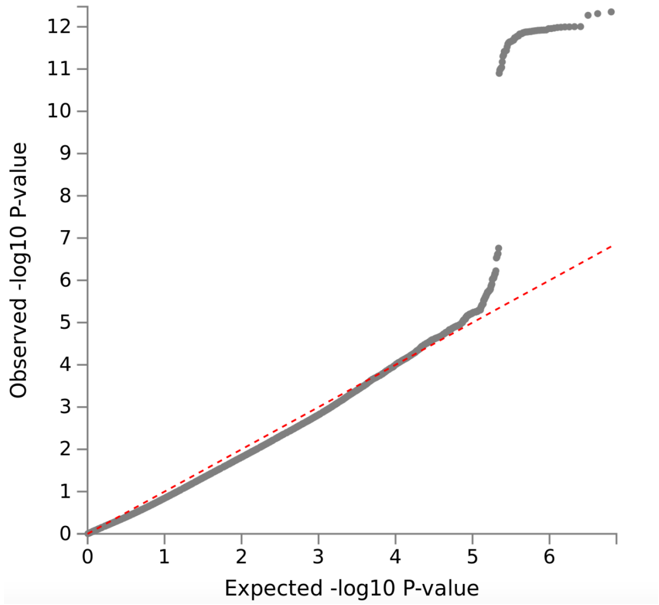

Genomic inflation factor (λ) = 0.825

**Supplementary Figure 4.** P-value correlation plot between PD GWAS and age at onset GWAS.

Log transformed P-values were plotted including the 90 independent PD GWAS signals in the Spanish cohort (x-axis) versus the same signals from the age at onset Spanish GWAS summary statistics (y-axis). SNPs are annotated by their closest gene. Green dots are loci that pass Bonferroni correction and red are loci that did not pass Bonferroni correction.

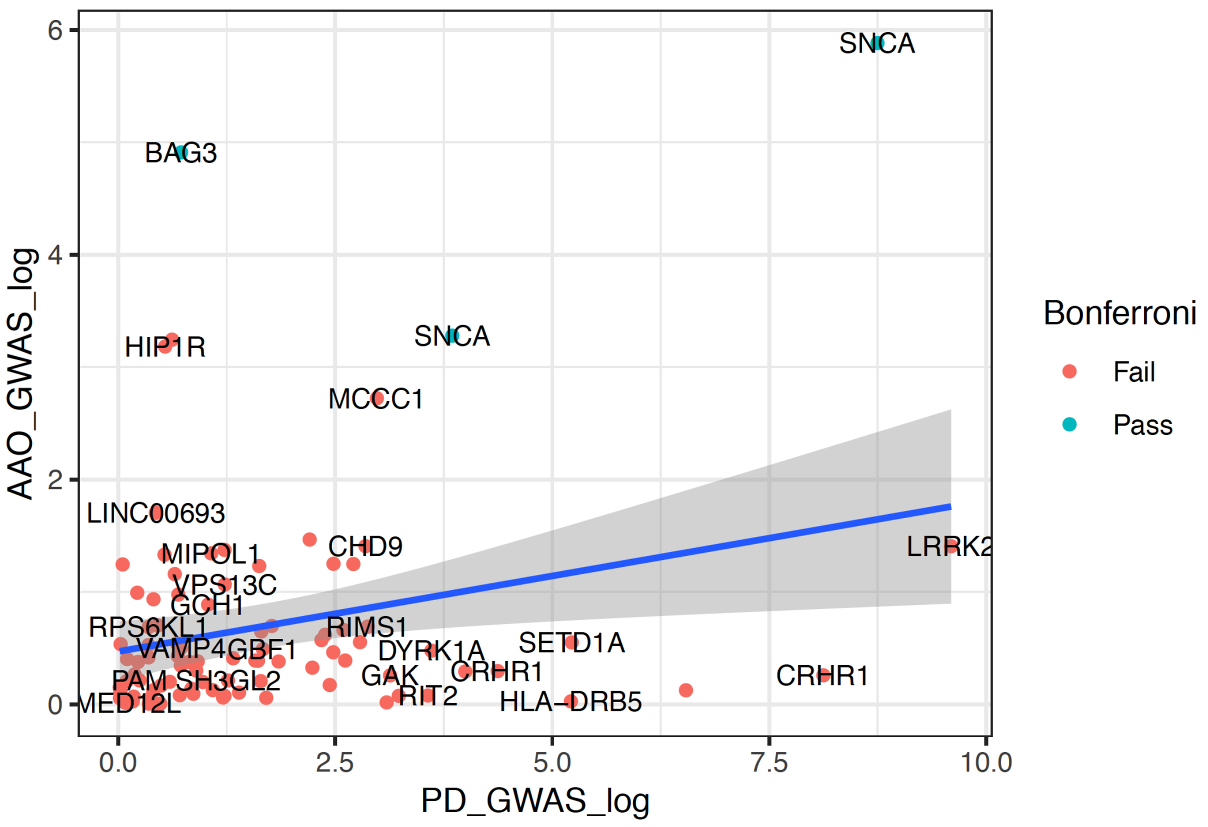

**Supplementary Figure 5.** *Interactive Genome Viewer* (IGV) screen capture displaying the sequencing reads generated from the bam files from five PD cases carrying the *PARK2* intronic variant (rs9356013). The image shows that all these individuals carried a deleterious frameshift mutation in exon 2 which is located 39 Kb away from the GWAS signal (c.155delA). The variant is an indel, so the deletion of a 1-bp leads to the truncation of the protein 28 amino acids later. As highlighted in the image, the variant was found in four cases in homozygous state and one case in heterozygous state.

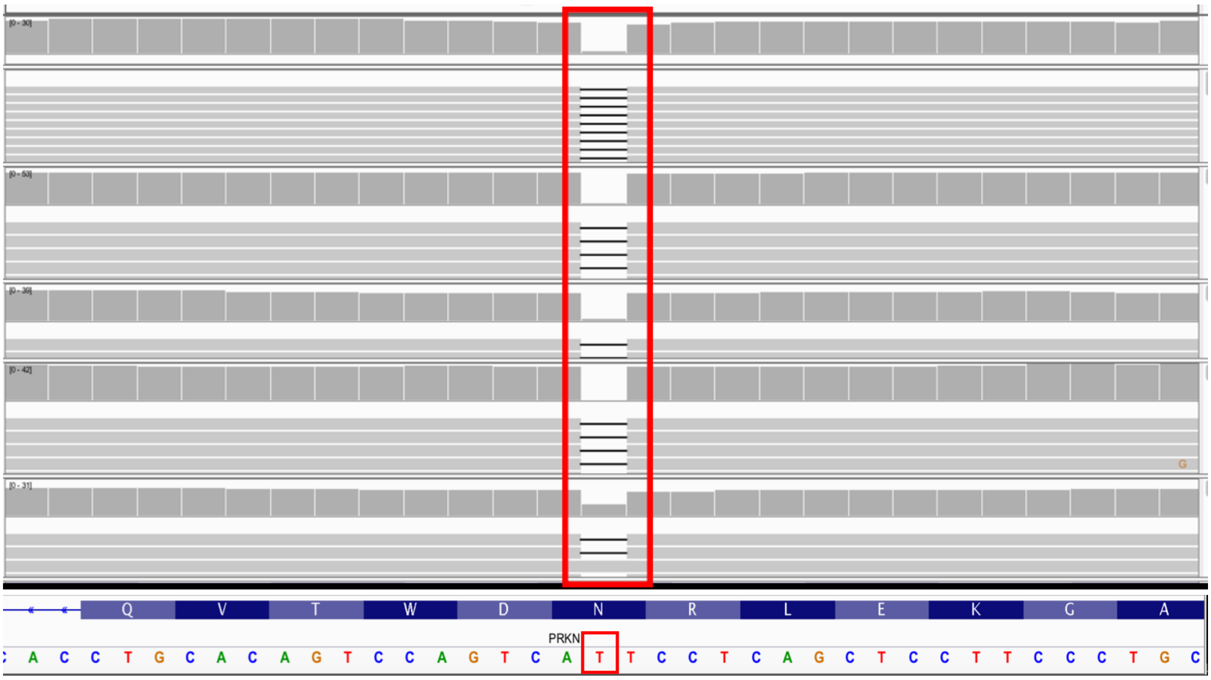

**Supplementary Figure 6.** Cluster plots of genetic variants in PD and Parkinsonism related genes (overrepresented in cases) generated from Genome Studio.

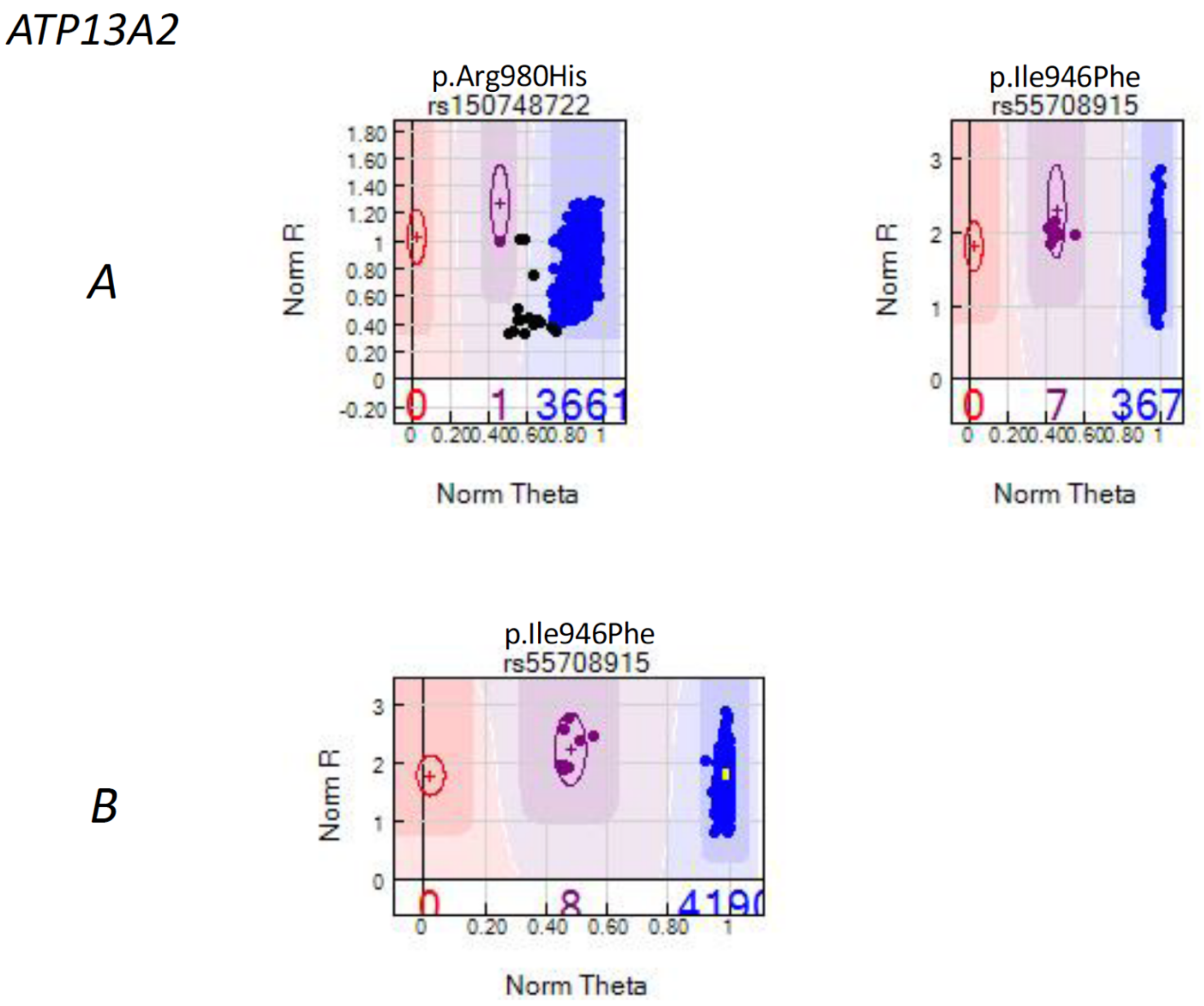

*DCTN1*

*A*

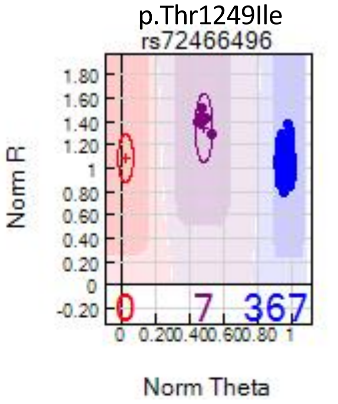
p.Thr1249Ile
rs72466496
Norm R
1.80
1.60
1.40
1.20
1
0.80
0.60
0.40
0.20
0
-0.20
0
7
367
0 0.200.400.600.80 1
Norm Theta

*B*

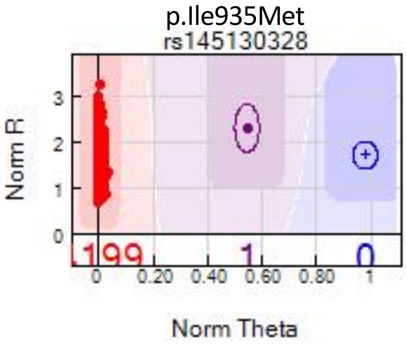
p.Ile935Met
rs145130328
Norm R
3
2
1
0
1
0
0 0.20 0.40 0.60 0.80 1
Norm Theta

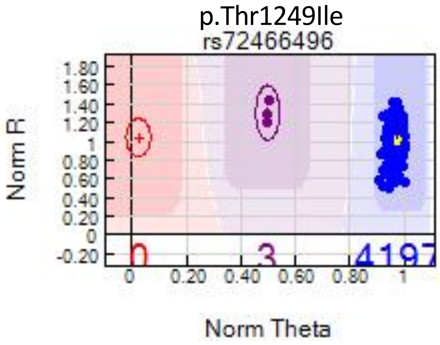
p.Thr1249Ile
rs72466496
Norm R
1.80
1.60
1.40
1.20
1
0.80
0.60
0.40
0.20
0
-0.20
0
3
0 0.20 0.40 0.60 0.80 1
Norm Theta

*FBXO7*

*A*

p.Arg635Gln

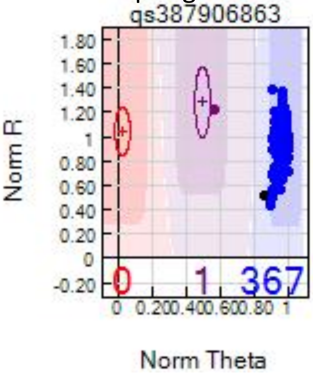

p.Tyr790Term

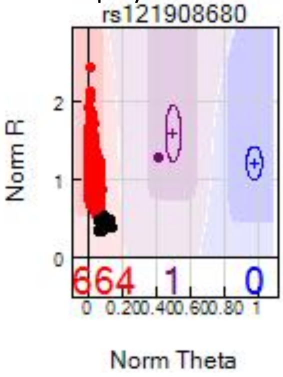

*B*

p.Ala343Thr

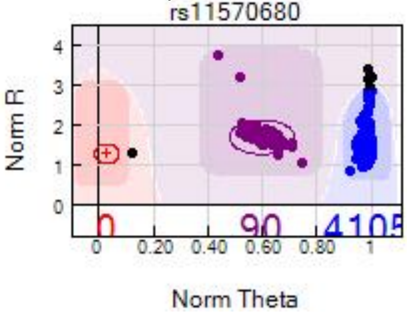

*GBA*

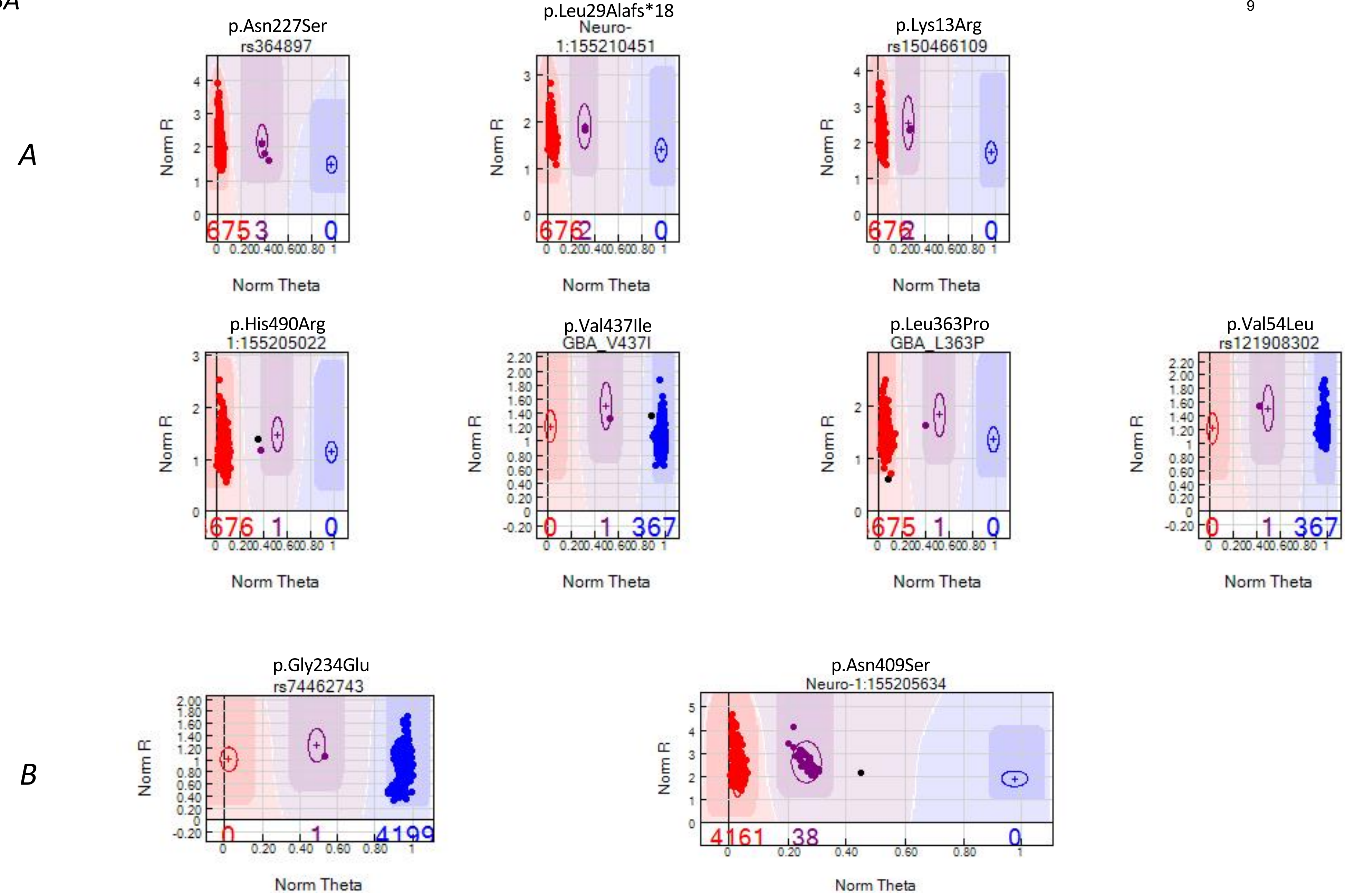
A
p.Asn227Ser
rs364897
675 3 0
p.Leu29Alafs*18
Neuro-
1:155210451
676 2 0
p.Lys13Arg
rs150466109
676 2 0
p.His490Arg
1:155205022
676 1 0
p.Val437Ile
GBA_V437I
0 1 367
p.Leu363Pro
GBA_L363P
675 1 0
p.Val54Leu
rs121908302
0 1 367
B
p.Gly234Glu
rs74462743
0 1 4199
p.Asn409Ser
Neuro-1:155205634
4161 38 0
Norm R
Norm Theta

*LRRK2*

*A*

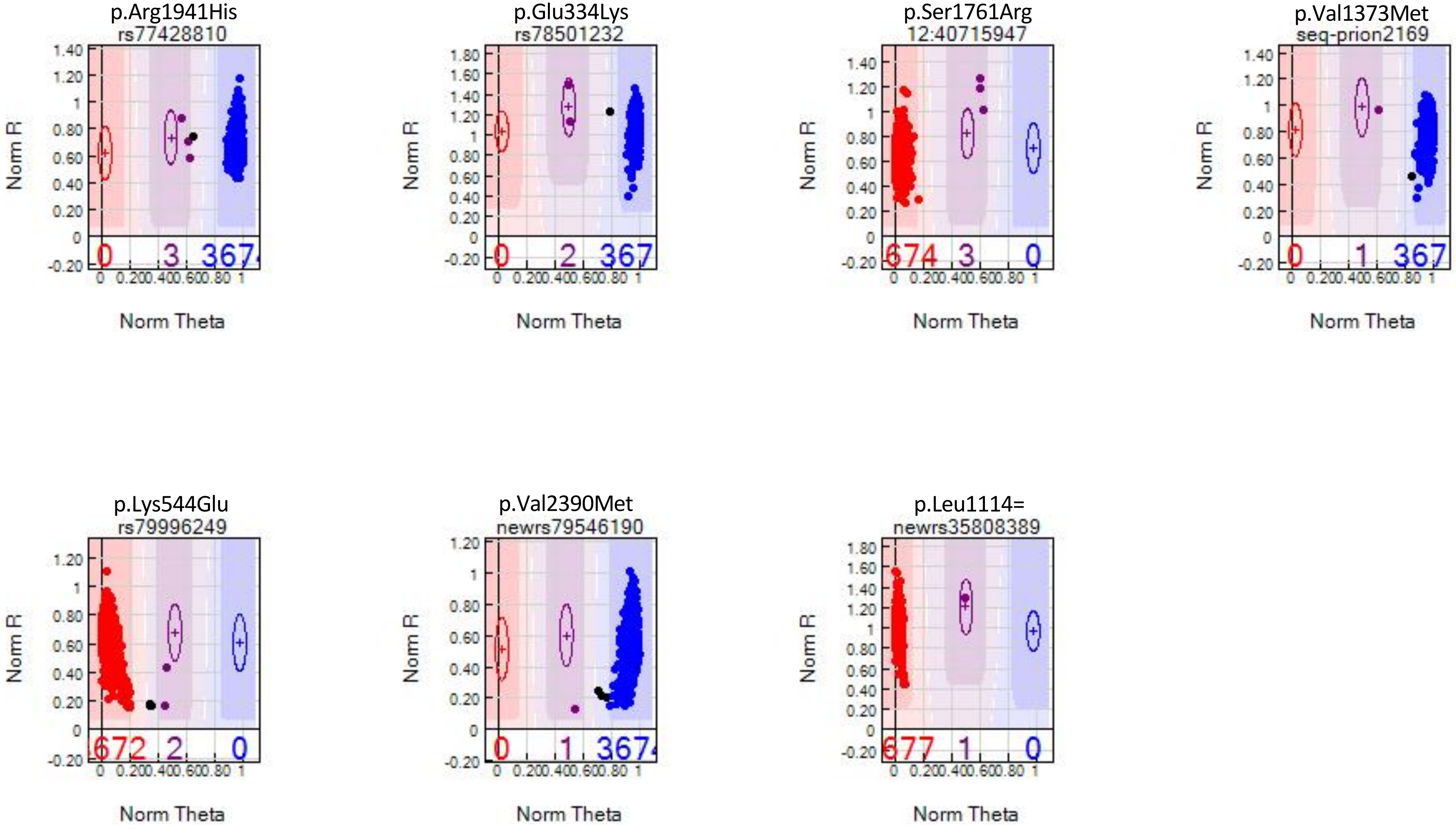
p.Arg1941His
rs77428810
Norm R
Norm Theta
0 3 367
p.Glu334Lys
rs78501232
Norm R
Norm Theta
0 2 367
p.Ser1761Arg
12:40715947
Norm R
Norm Theta
674 3 0
p.Val1373Met
seq-prion2169
Norm R
Norm Theta
0 1 367
p.Lys544Glu
rs79996249
Norm R
Norm Theta
672 2 0
p.Val2390Met
newrs79546190
Norm R
Norm Theta
0 1 367
p.Leu1114=
newrs35808389
Norm R
Norm Theta
677 1 0

*B*

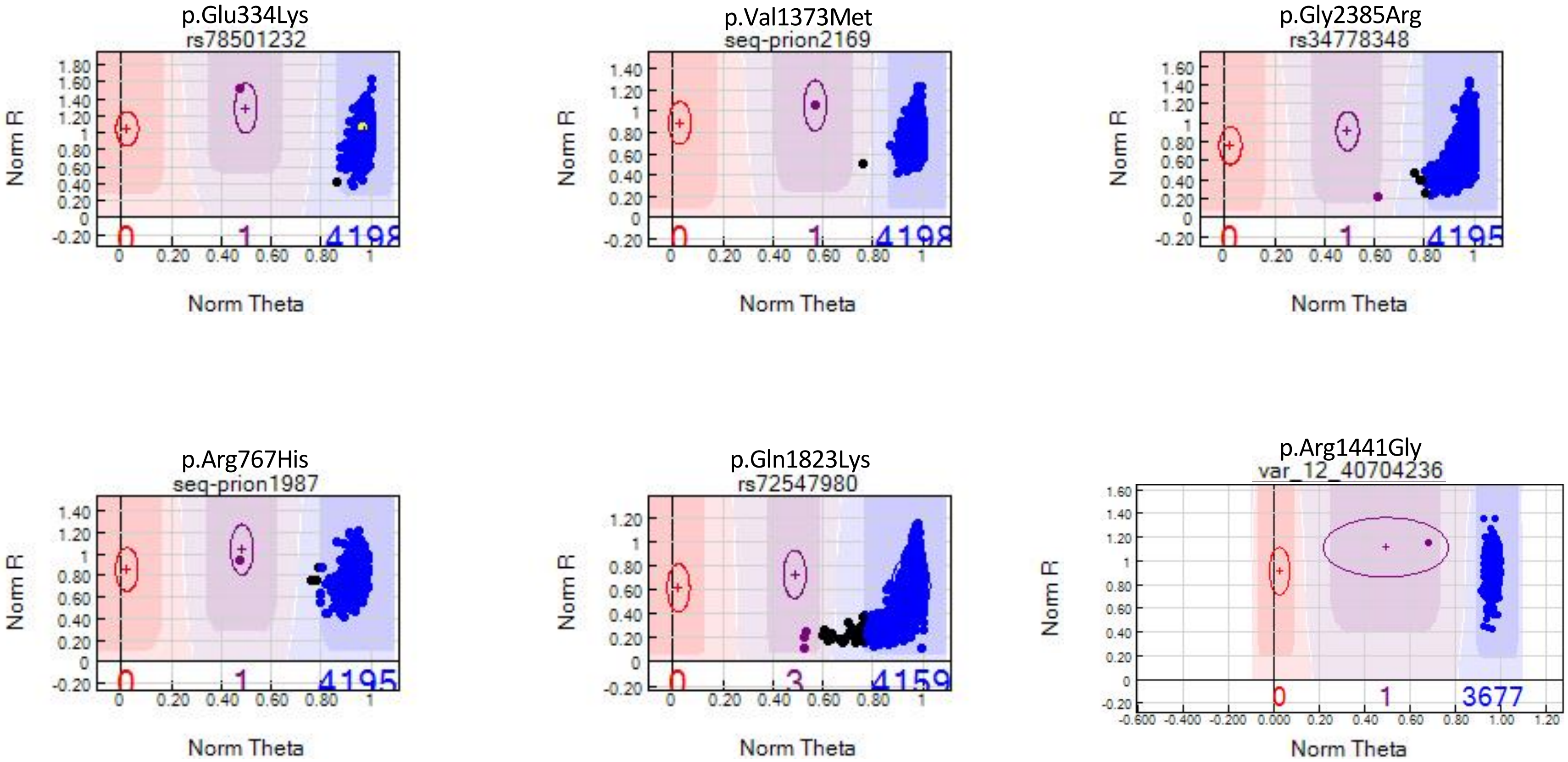
p.Glu334Lys
rs78501232
Norm R
Norm Theta
0
1
4198
p.Val1373Met
seq-prion2169
Norm R
Norm Theta
0
1
4198
p.Gly2385Arg
rs34778348
Norm R
Norm Theta
0
1
4195
p.Arg767His
seq-prion1987
Norm R
Norm Theta
0
1
4195
p.Gln1823Lys
rs72547980
Norm R
Norm Theta
0
3
4159
p.Arg1441Gly
var_12_40704236
Norm R
Norm Theta
0
1
3677

*MAPT*

*A*

p.Asn613del

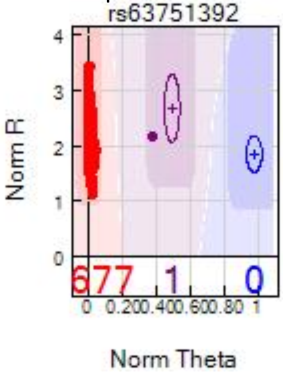

*PARK2*

*A*

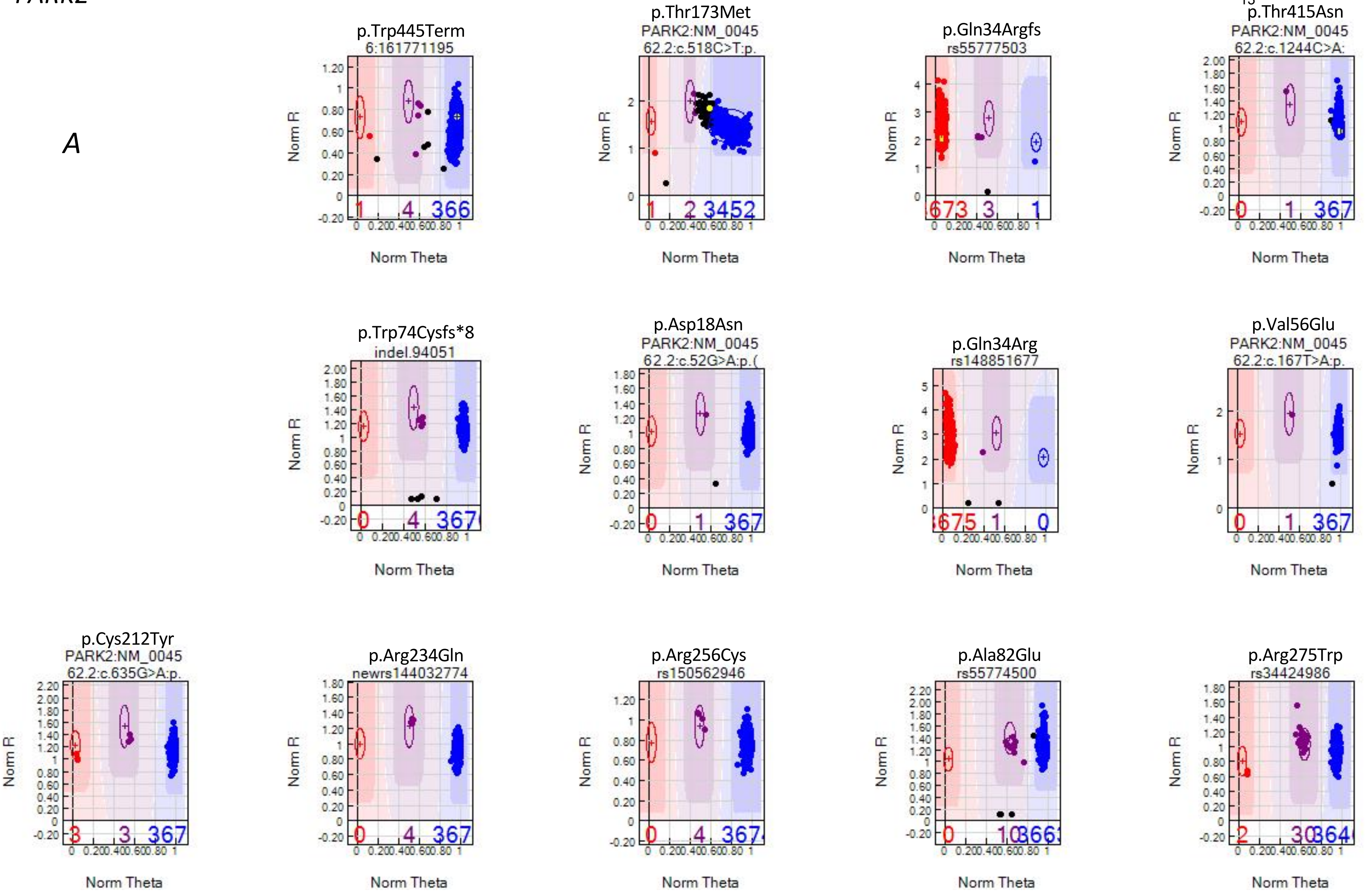
p.Trp445Term
6:161771195
p.Thr173Met
PARK2:NM_0045
62.2:c.518C>T:p.
p.Gln34Argfs
rs55777503
p.Thr415Asn
PARK2:NM_0045
62.2:c.1244C>A:
p.Trp74Cysfs*8
indel.94051
p.Asp18Asn
PARK2:NM_0045
62.2:c.52G>A:p.(
p.Gln34Arg
rs148851677
p.Val56Glu
PARK2:NM_0045
62.2:c.167T>A:p.
p.Cys212Tyr
PARK2:NM_0045
62.2:c.635G>A:p.
p.Arg234Gln
newrs144032774
p.Arg256Cys
rs150562946
p.Ala82Glu
rs55774500
p.Arg275Trp
rs34424986
Norm R
Norm Theta

*B*

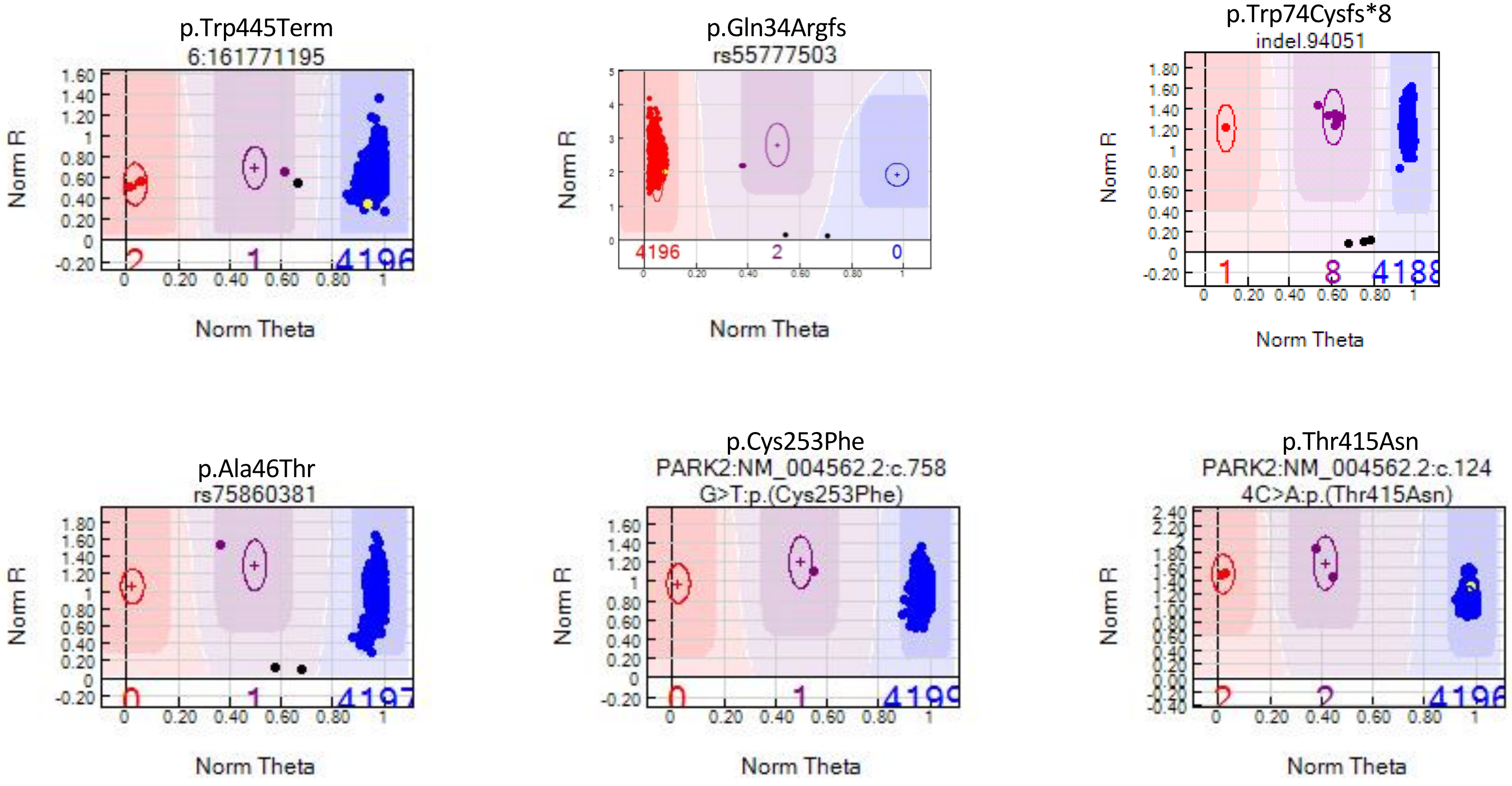
p.Trp445Term
6:161771195
p.Gln34Argfs
rs55777503
p.Trp74Cysfs*8
indel.94051
p.Ala46Thr
rs75860381
p.Cys253Phe
PARK2:NM_004562.2:c.758
G>T:p.(Cys253Phe)
p.Thr415Asn
PARK2:NM_004562.2:c.124
4C>A:p.(Thr415Asn)
Norm R
Norm Theta

*PARK2*

*B (cont.)*

p.Cys212Gly
PARK2:NM_004562.2:c.634
T>G:p.(Cys212Gly)

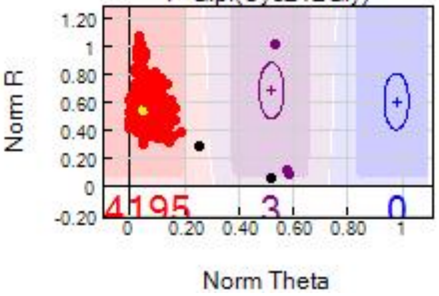

p.Arg42Pro
rs368134308

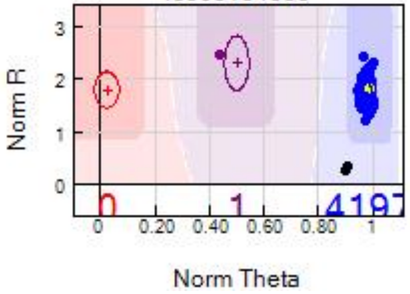

p.Arg234Gln
newrs144032774

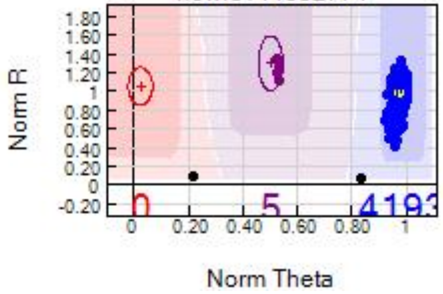

p.Arg275Trp
rs34424986

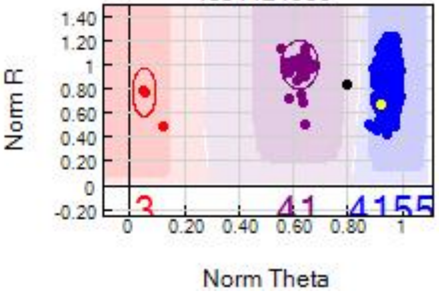

p.Ala82Glu
rs55774500

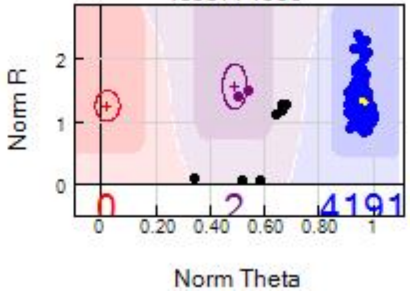

p.Arg256Cys
rs150562946

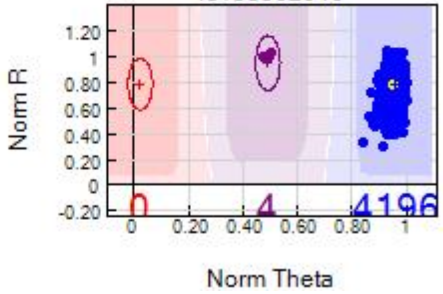

p.Ile298Ser
PARK2:NM_004562.2:c.893
T>G:p.(Ile298Ser)

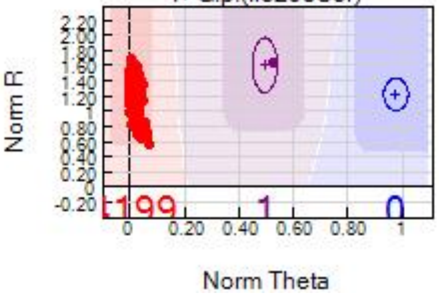

*PINK1*

*A*

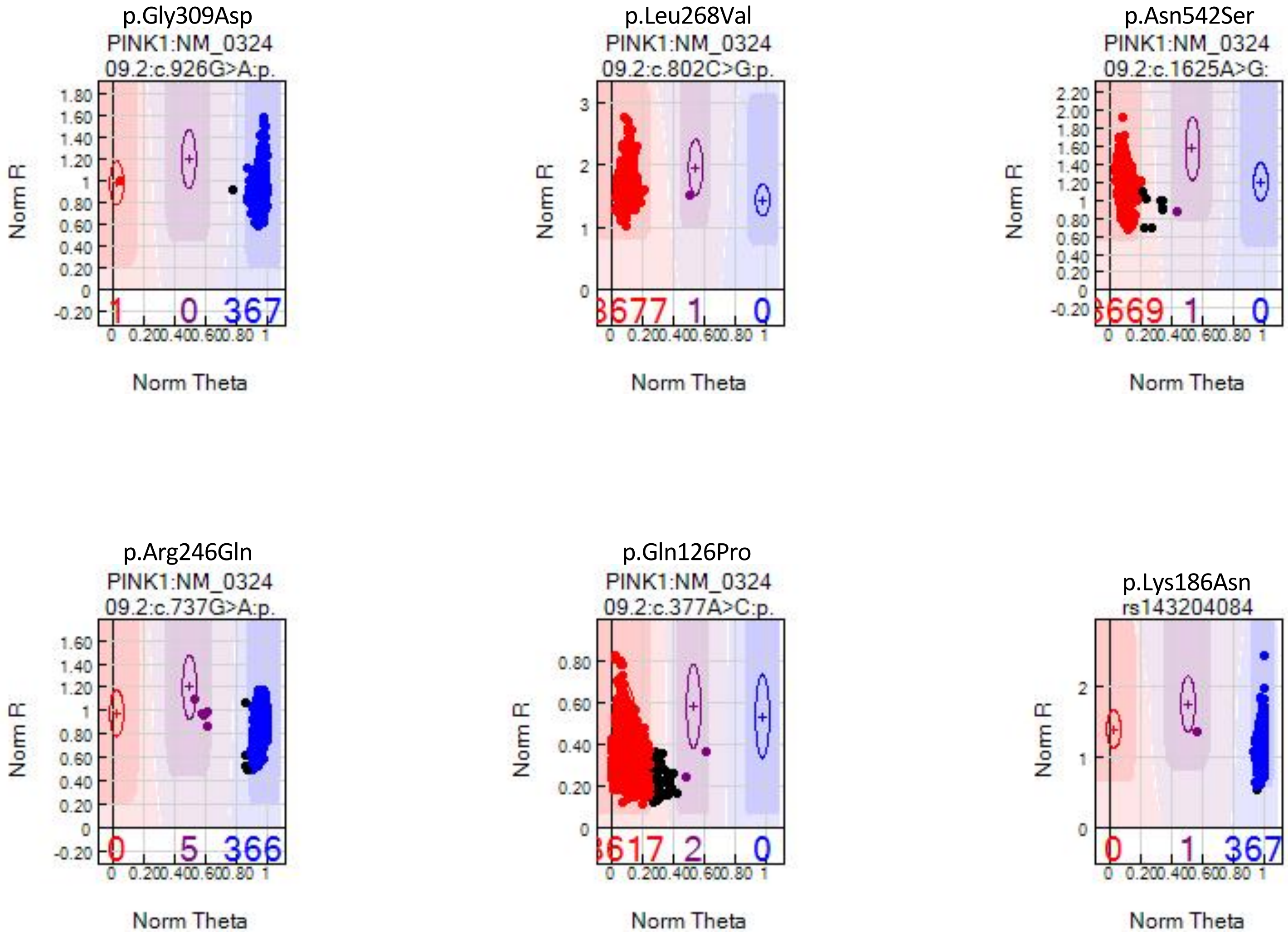
p.Gly309Asp
PINK1:NM_0324
09.2:c.926G>A:p.
Norm R
Norm Theta
1
0
367
p.Leu268Val
PINK1:NM_0324
09.2:c.802C>G:p.
3677
1
0
p.Asn542Ser
PINK1:NM_0324
09.2:c.1625A>G:
3669
1
0
p.Arg246Gln
PINK1:NM_0324
09.2:c.737G>A:p.
0
5
366
p.Gln126Pro
PINK1:NM_0324
09.2:c.377A>C:p.
3617
2
0
p.Lys186Asn
rs143204084
0
1
367

*PINK1*

*B*

p.Leu369Pro

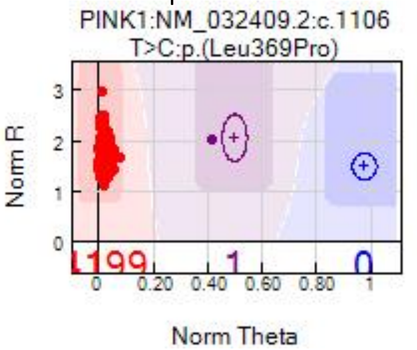

p.Gly32Arg

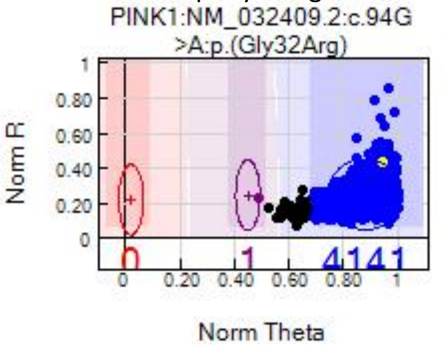

*A*

p.Arg232His

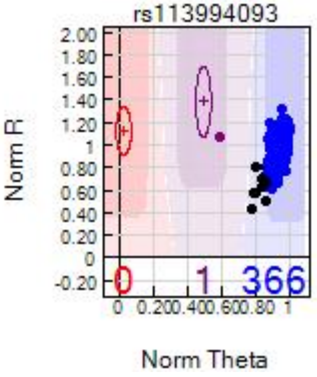

p.Arg579Trp

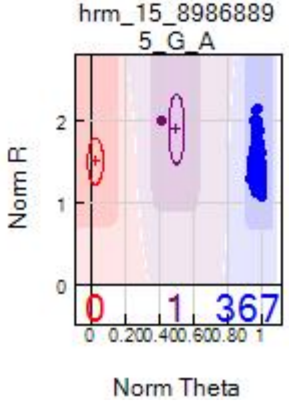

p.Asn468Asp

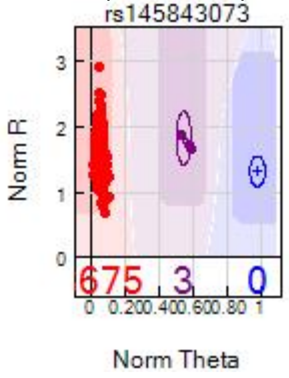

*B*

p.Asn468Asp

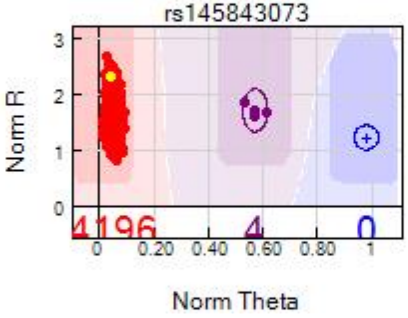

p.Pro587Leu

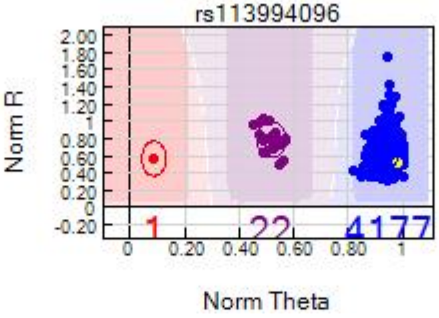

p.Ty831-Cys

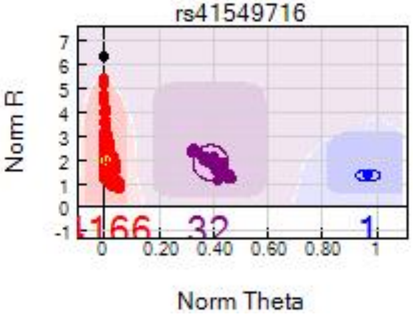

*VPS35*

*A*

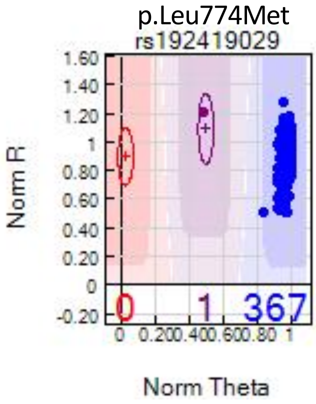
p.Leu774Met
rs192419029
Norm R
1.60
1.40
1.20
1
0.80
0.60
0.40
0.20
0
-0.20
0
1
367
0 0.200.400.600.80 1
Norm Theta

*B*

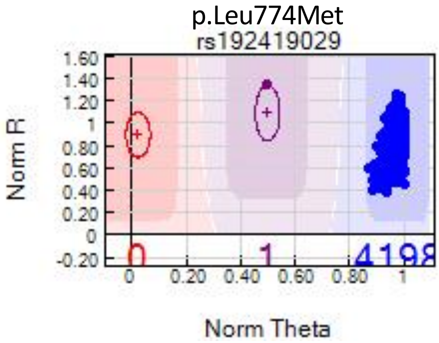
p.Leu774Met
rs192419029
Norm R
1.60
1.40
1.20
1
0.80
0.60
0.40
0.20
0
-0.20
0
1
0 0.20 0.40 0.60 0.80 1
Norm Theta

**Supplementary Figure 7.** Copy number variation plots displaying *PARK2* deletions or duplications

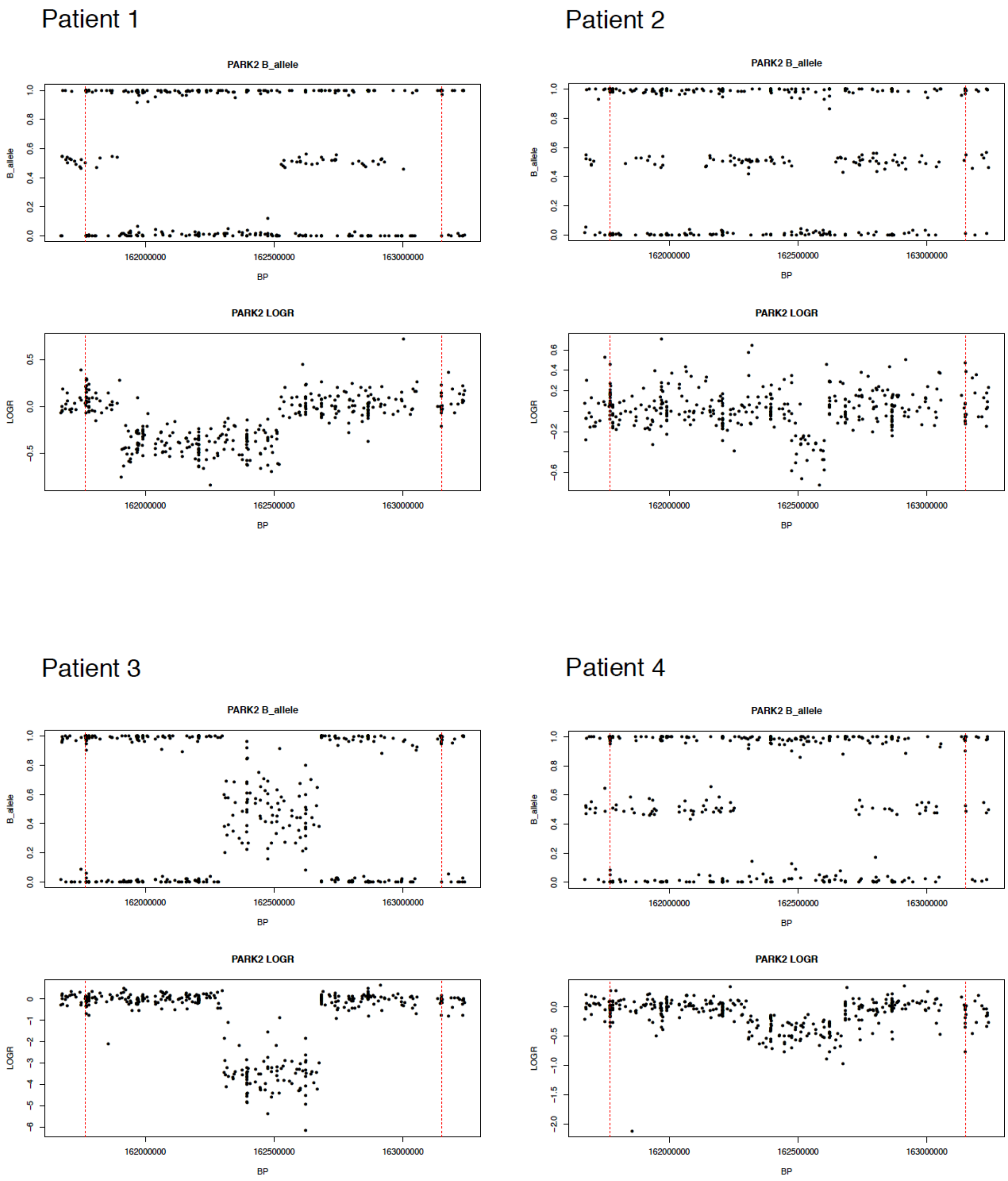

### Patient 35

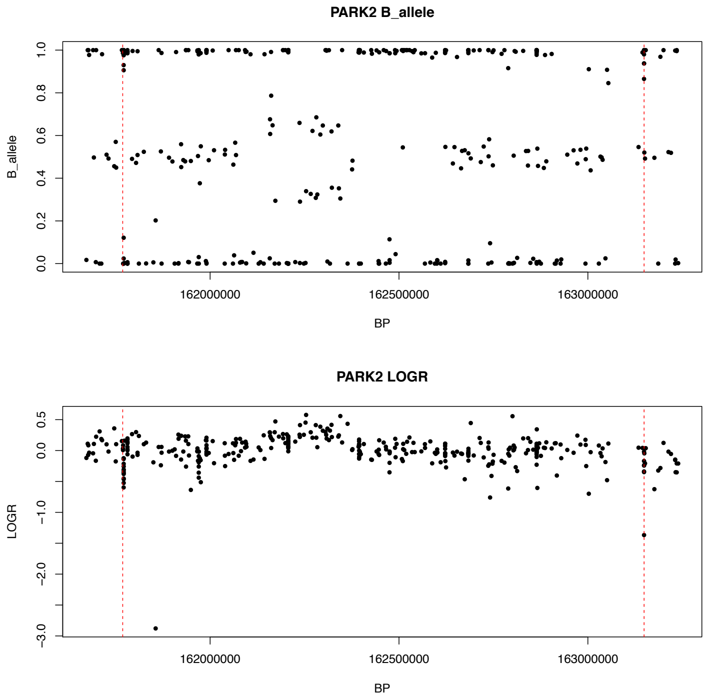

### Patient 36

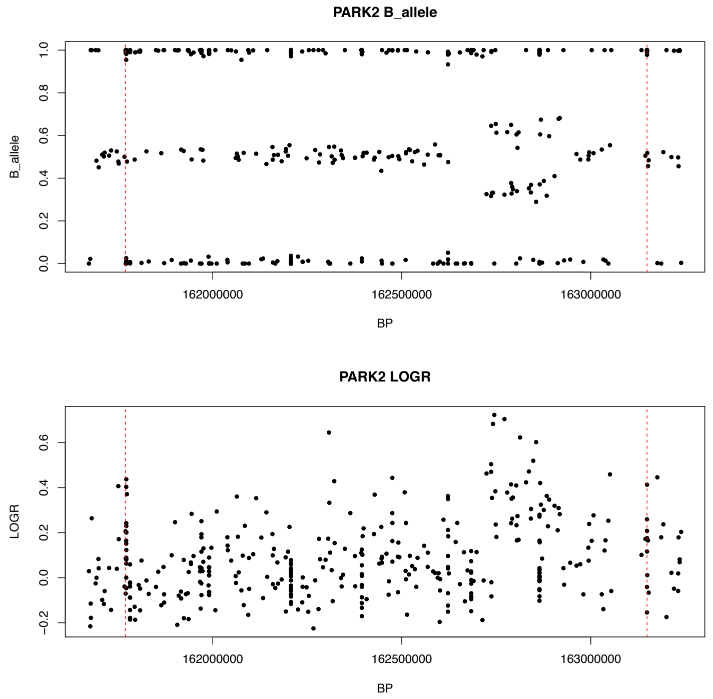

### Patient 37

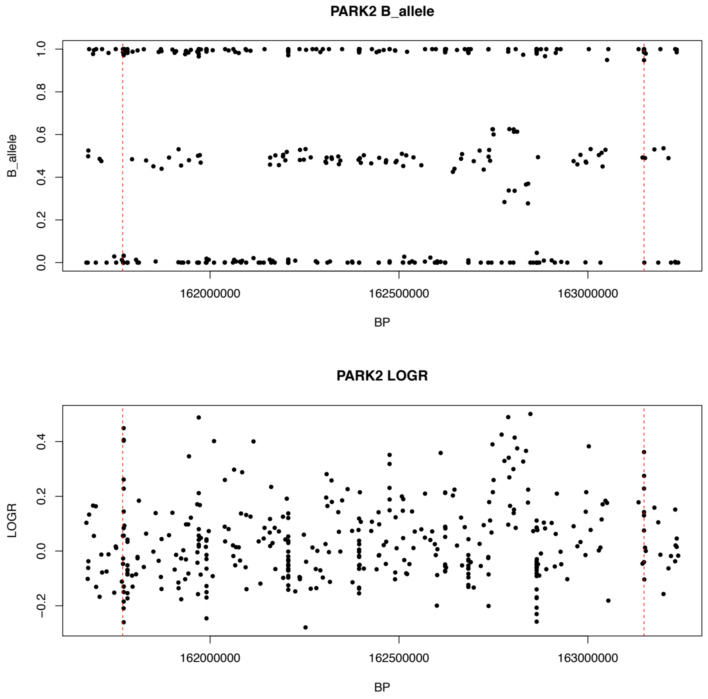

### Patient 38

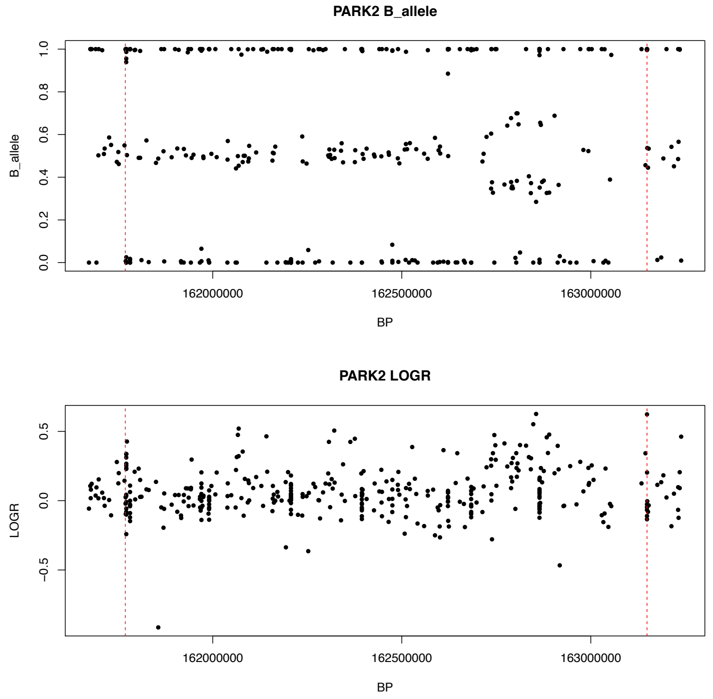

#### Patient 39

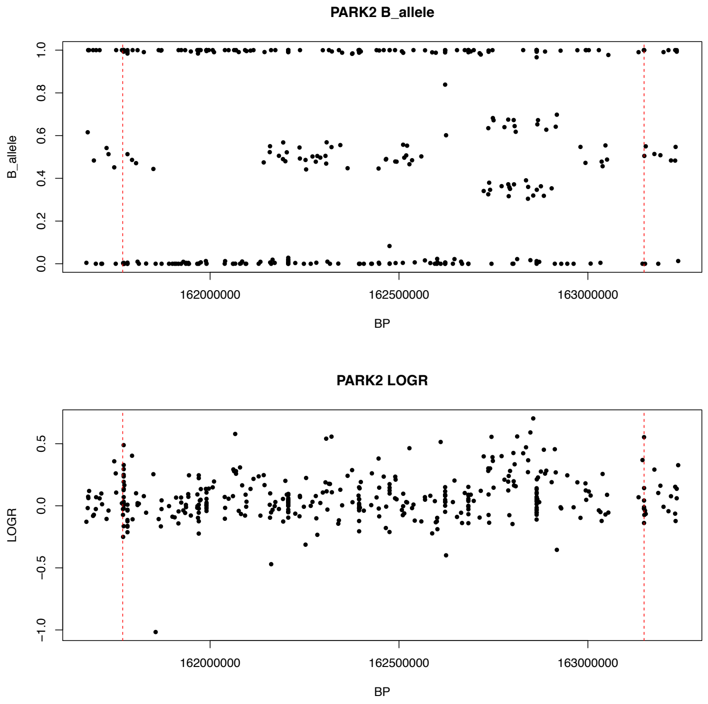

#### Patient 40

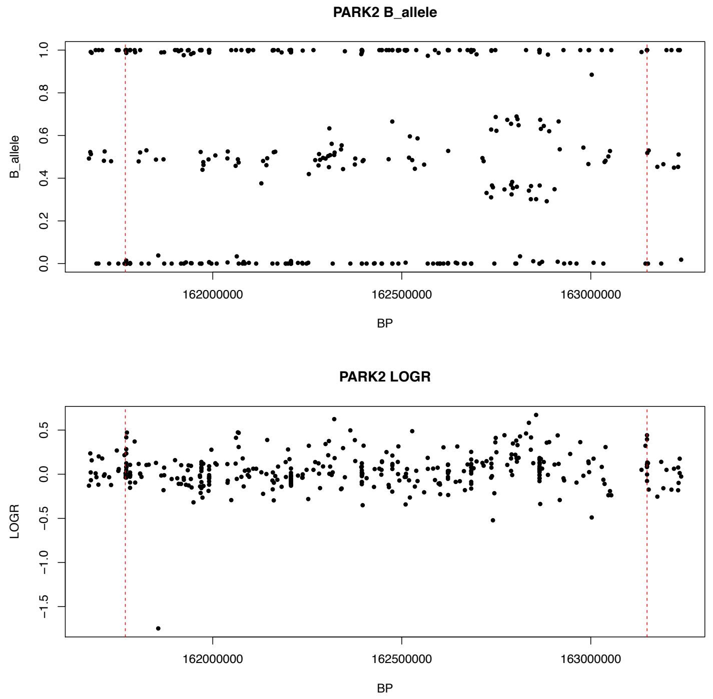

#### Patient 41

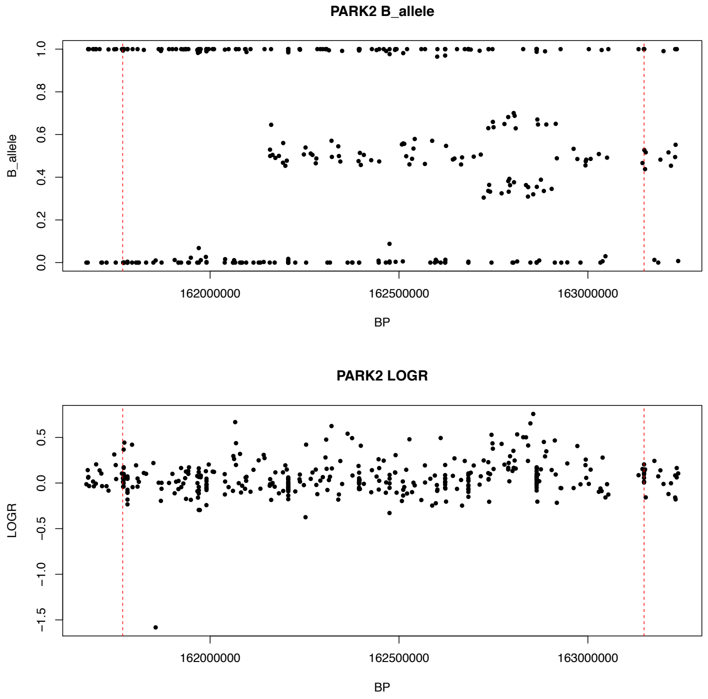

#### Patient 42

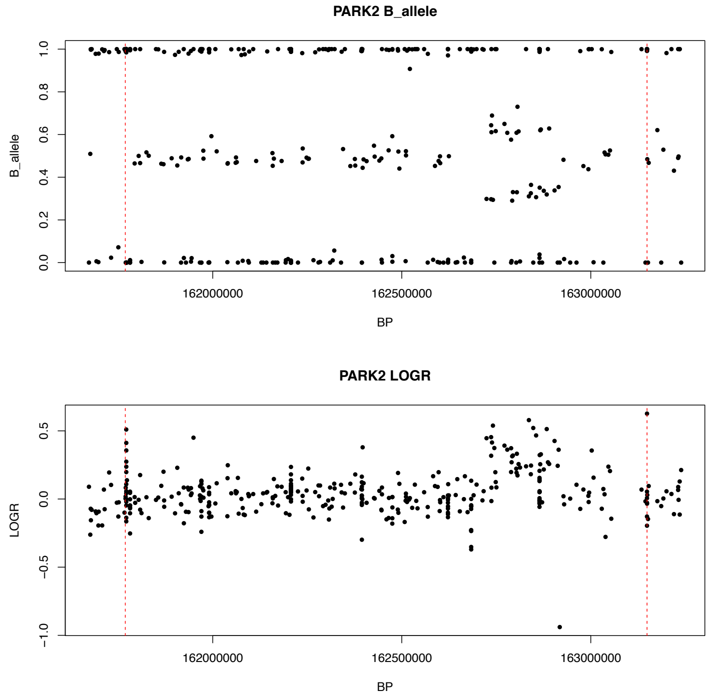

### Patient 43

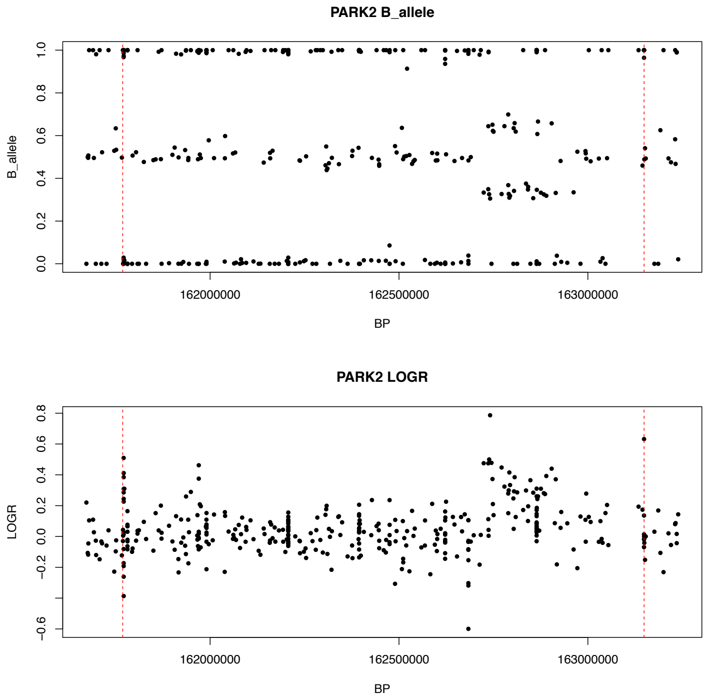

### Patient 44

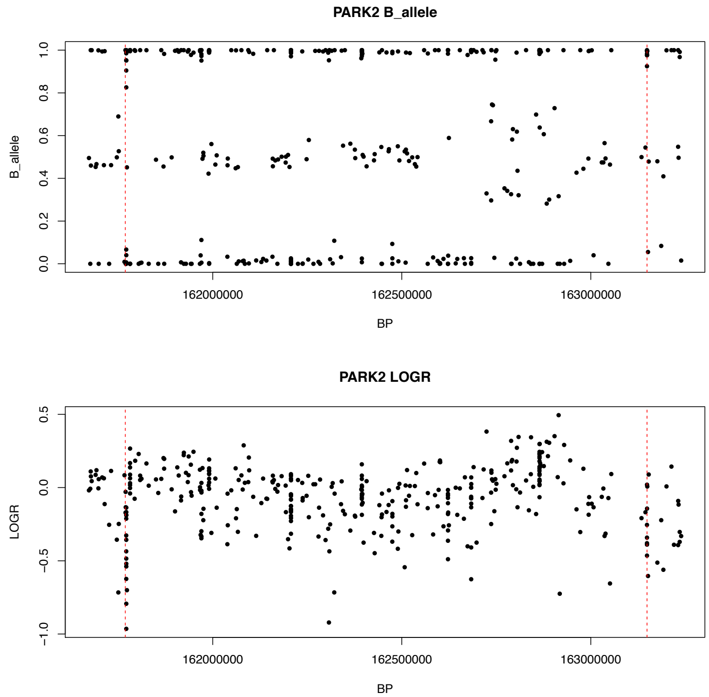

### Patient 45

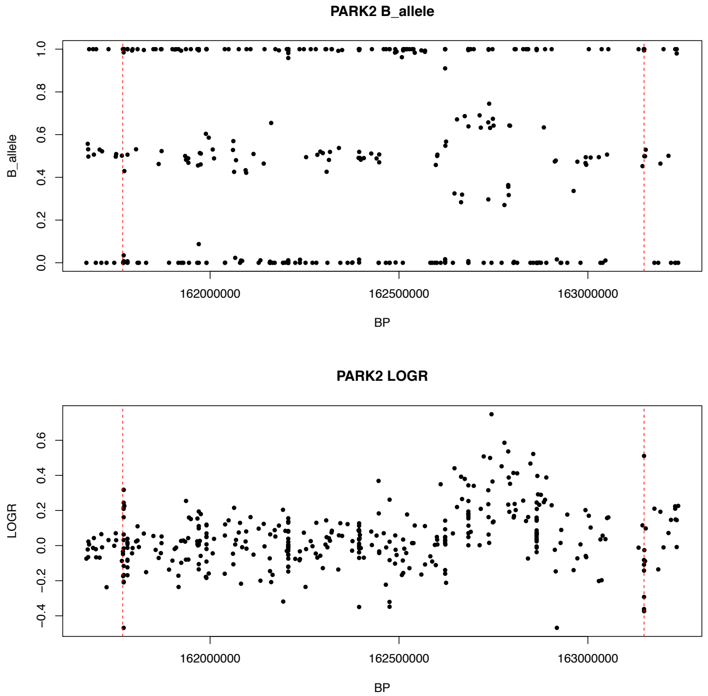

### Patient 46

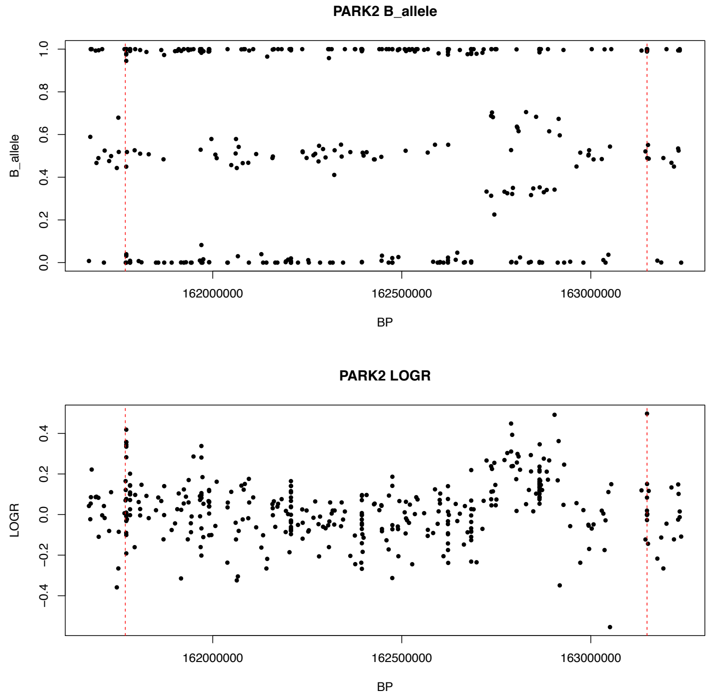

### Patient 5

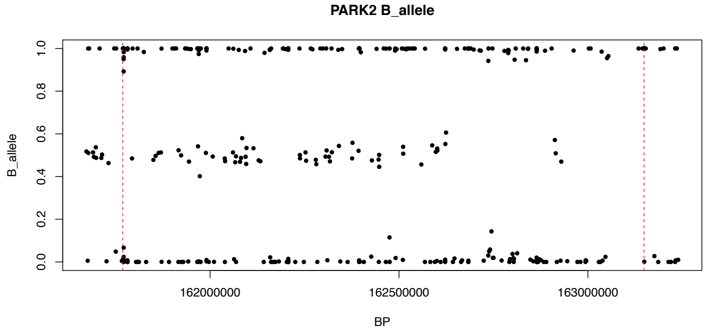

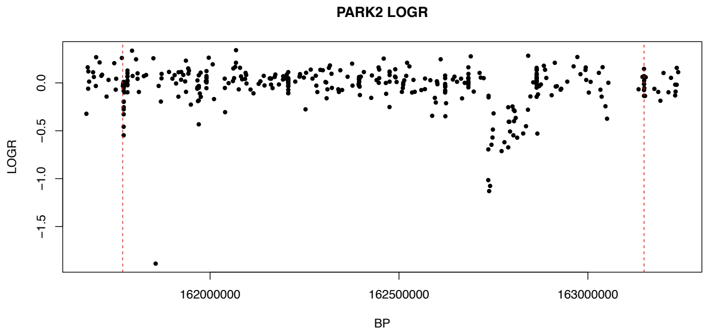

### Patient 6

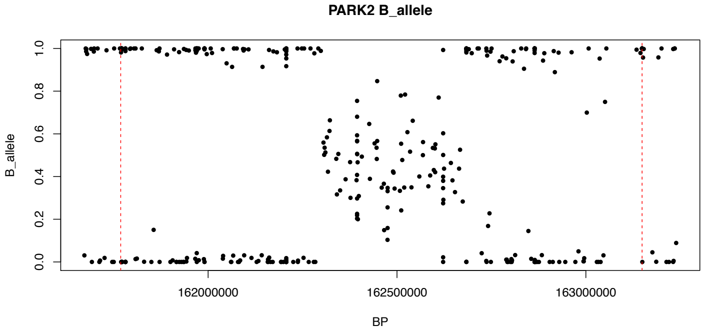

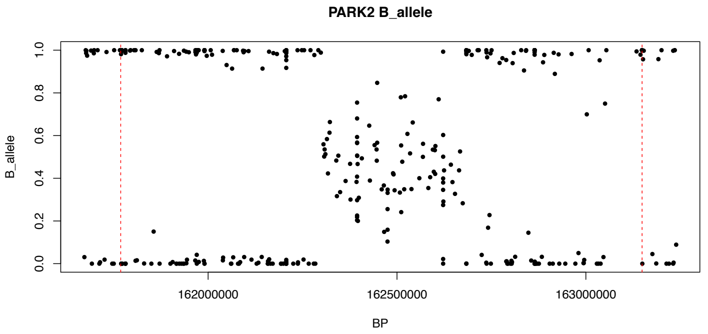

### Patient 7

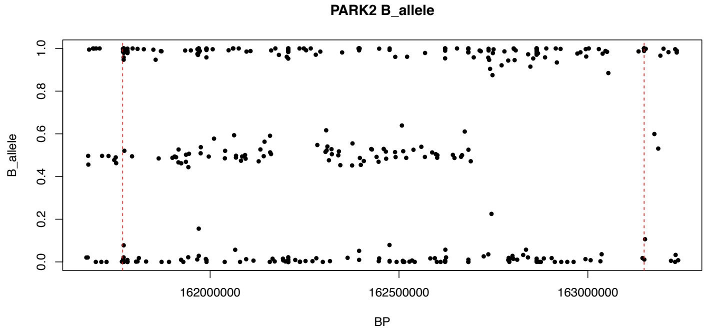

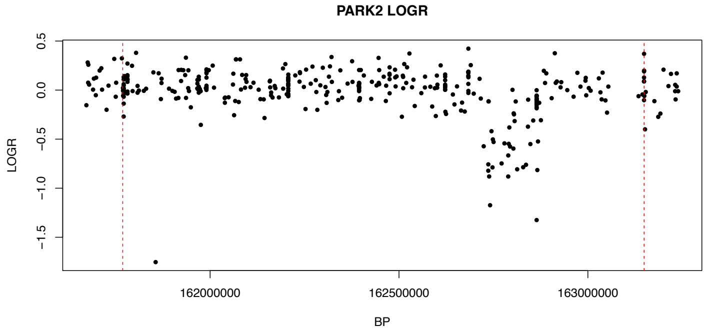

### Patient 8

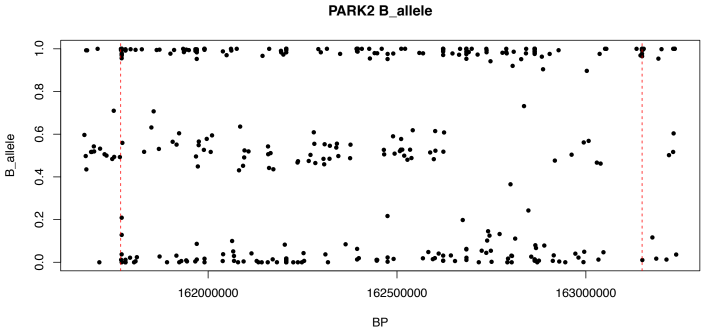

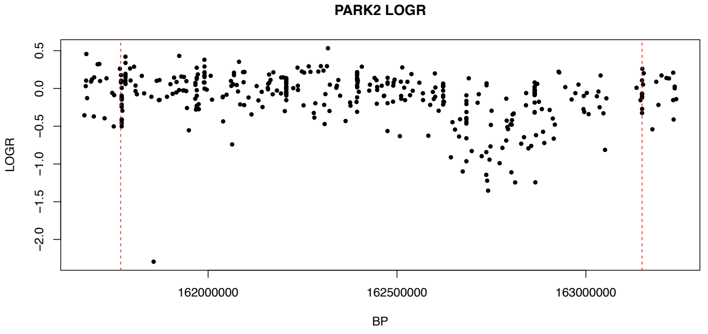

### Patient 9

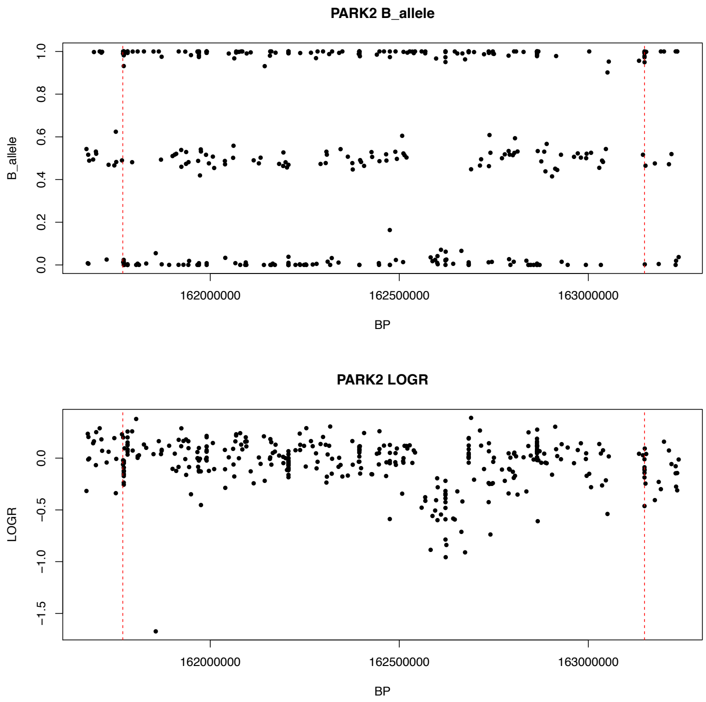

### Patient 10

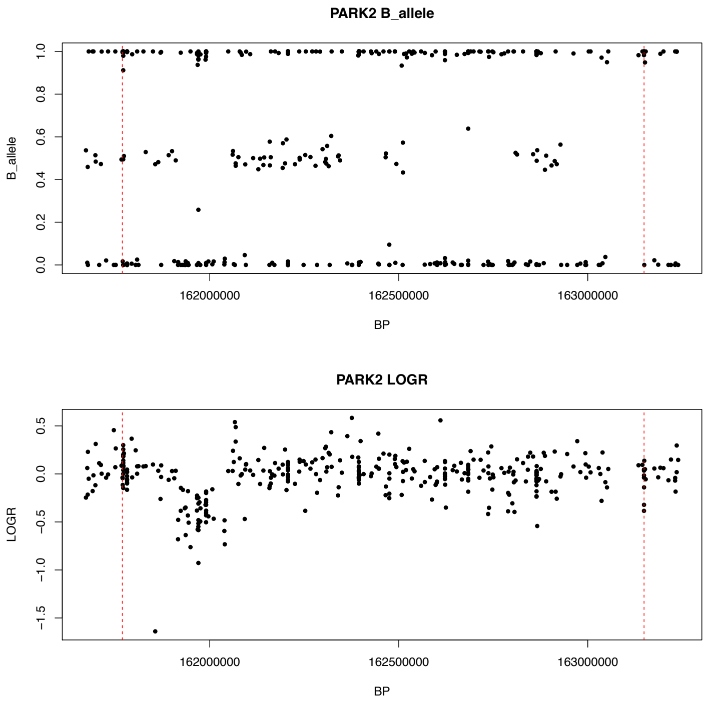

### Patient 11

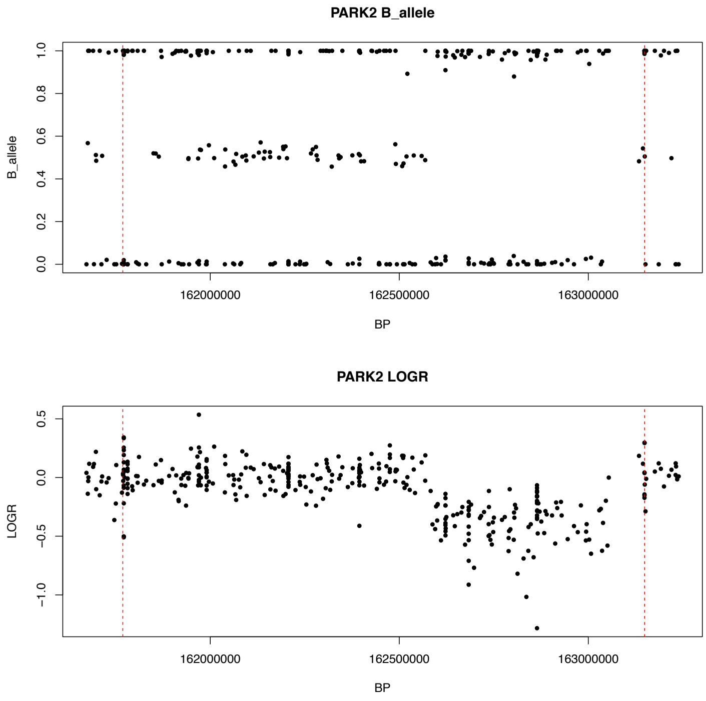

### Patient 12

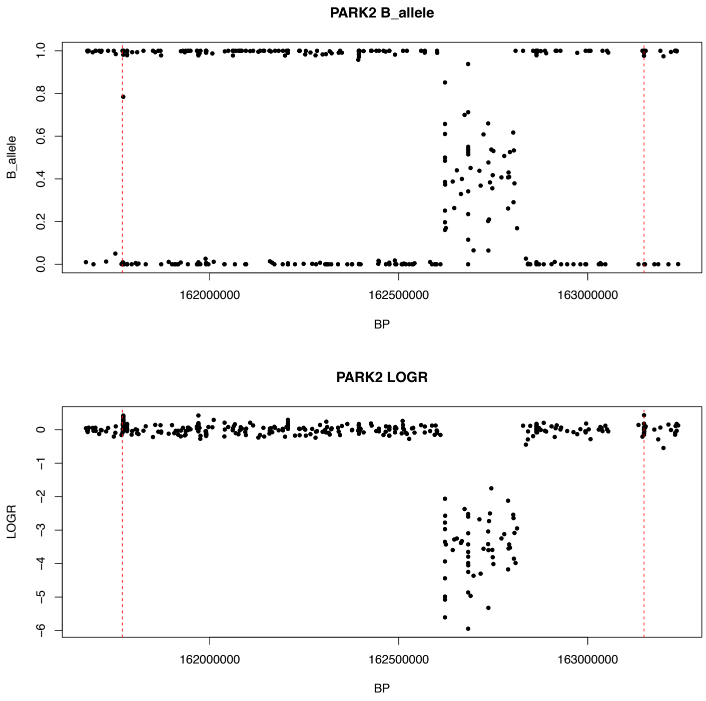

#### Patient 13

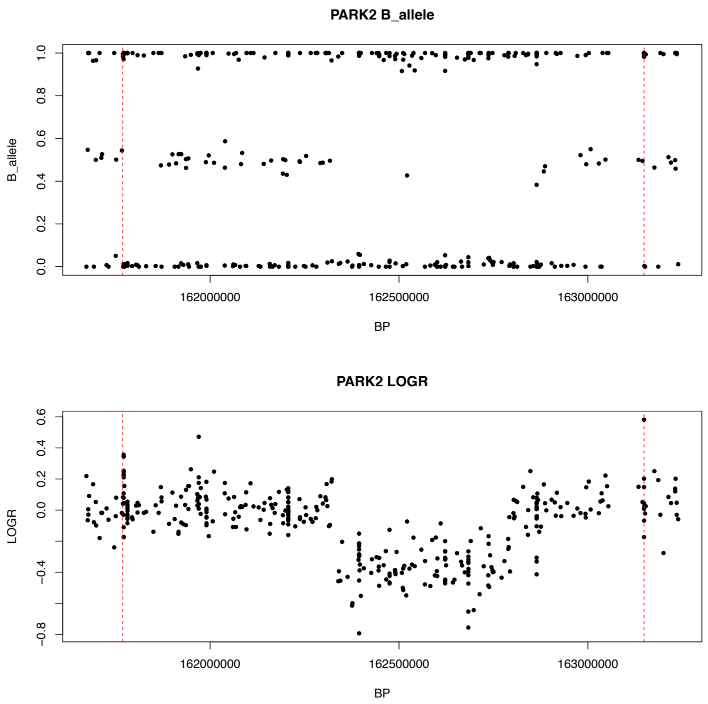

#### Patient 14

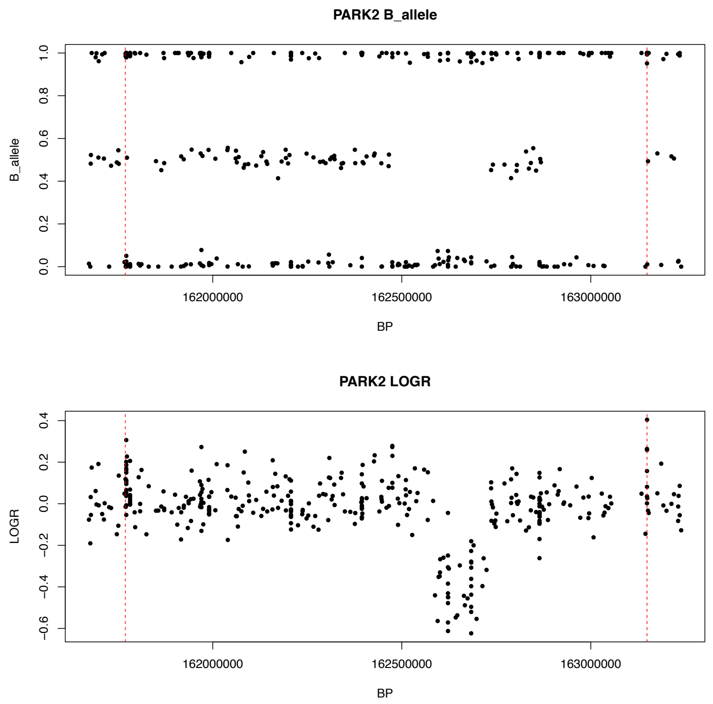

#### Patient 15

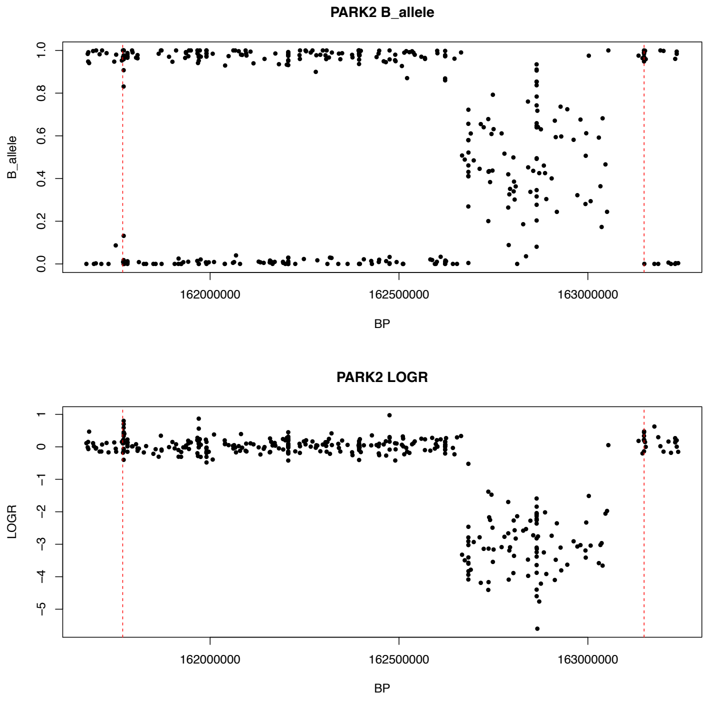

#### Patient 16

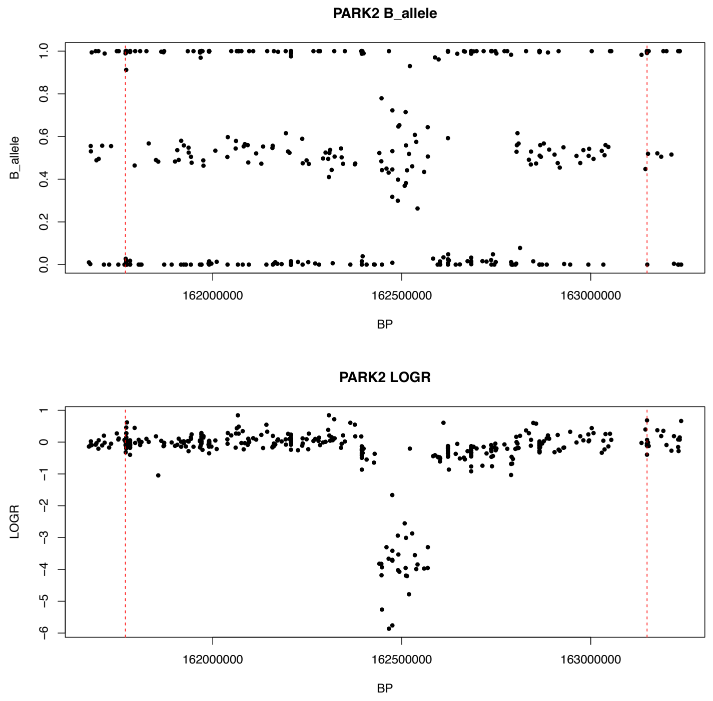

#### Patient 17

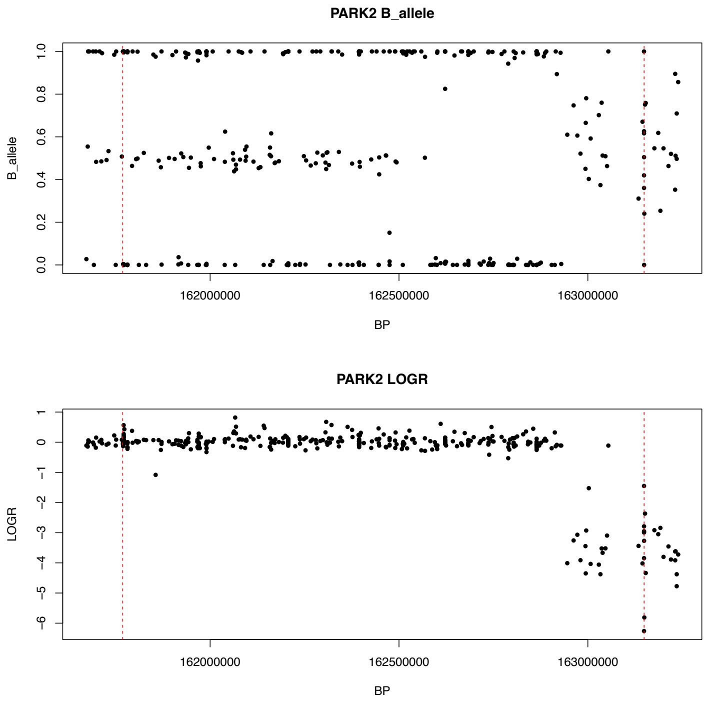

#### Patient 18

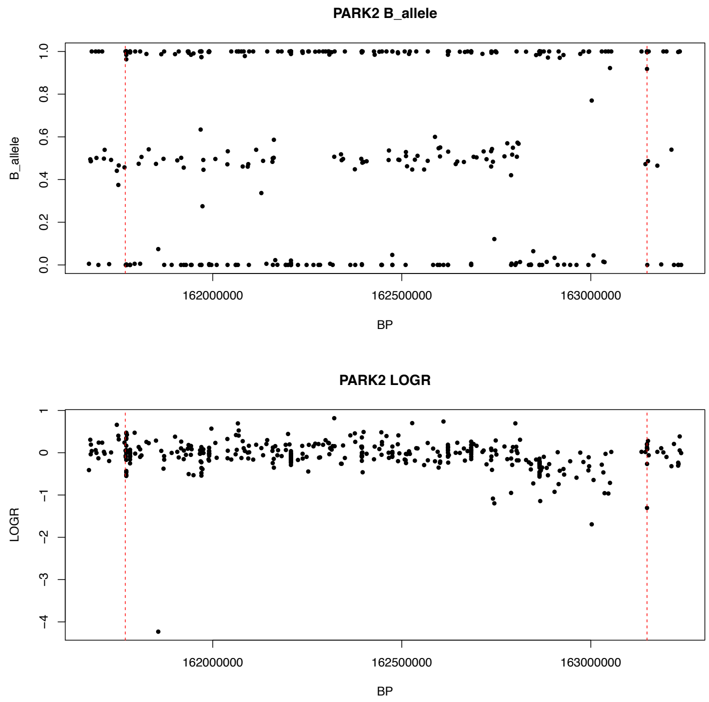

#### Patient 19

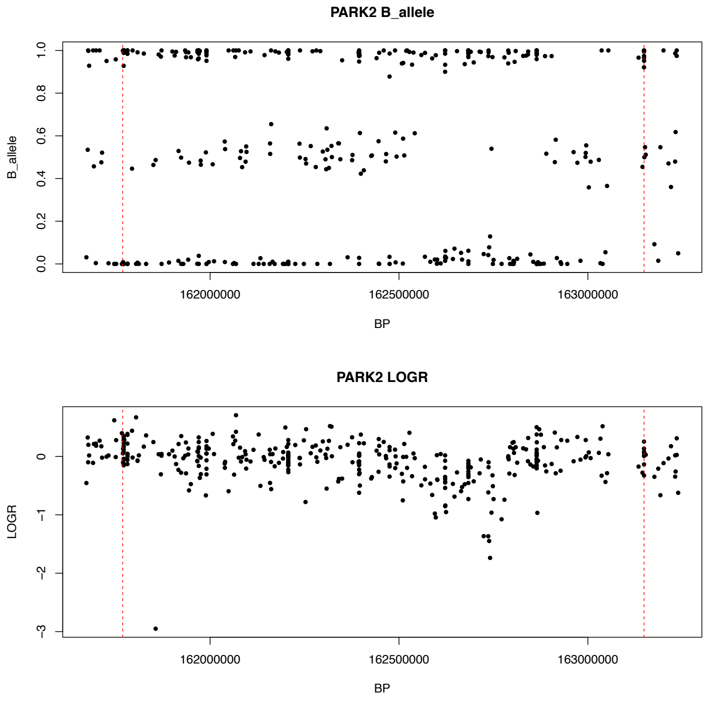

#### Patient 20

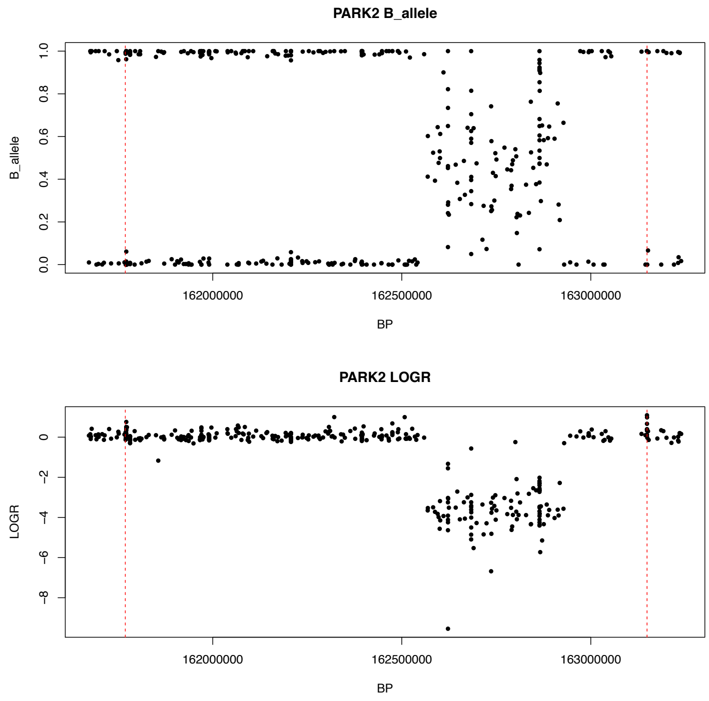

### Patient 21

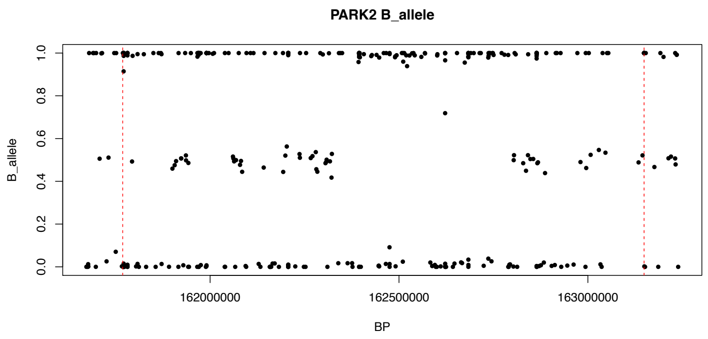

### Patient 22

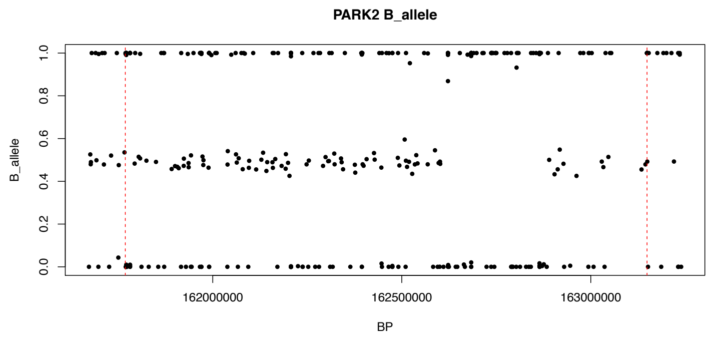

### Patient 23

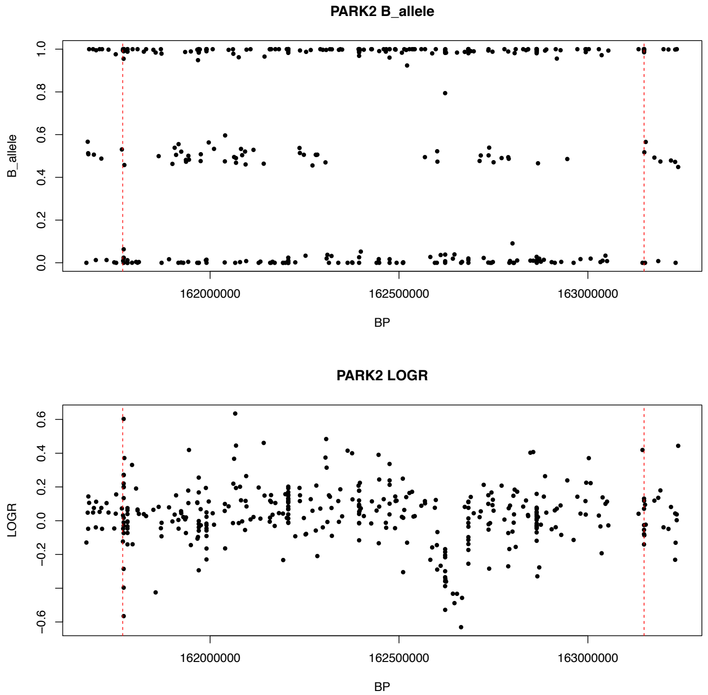

### Patient 24

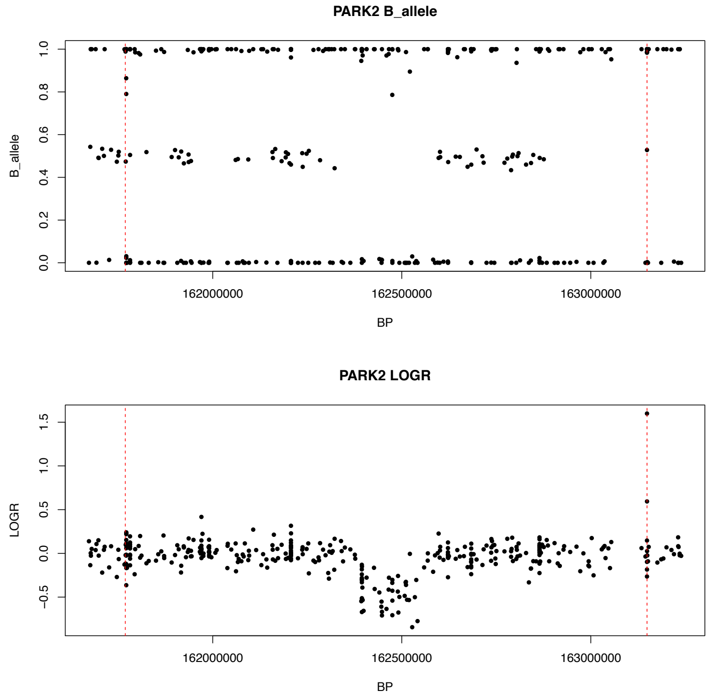

### Patient 25

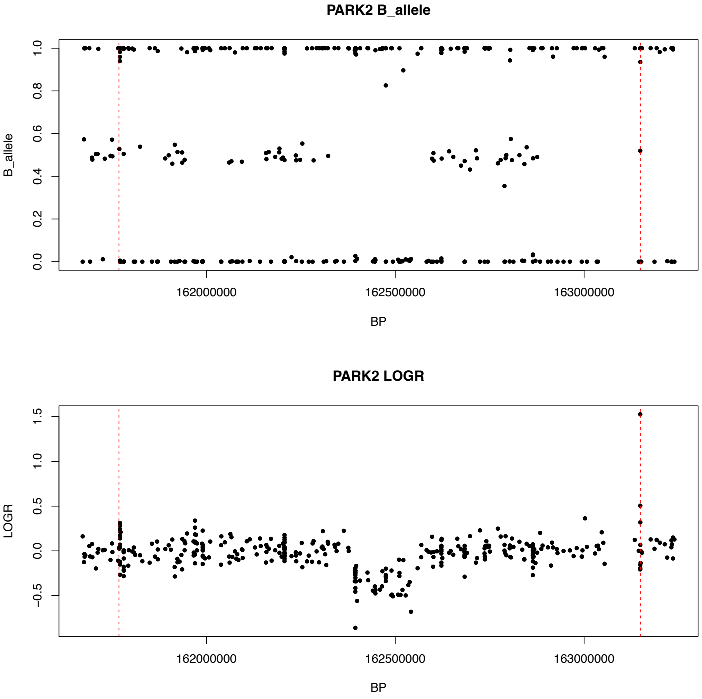

### Patient 26

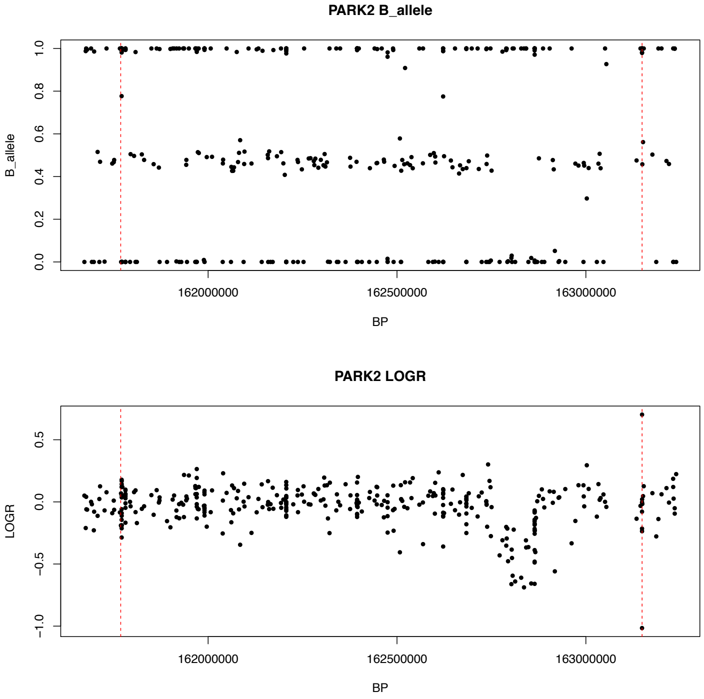

### Patient 27

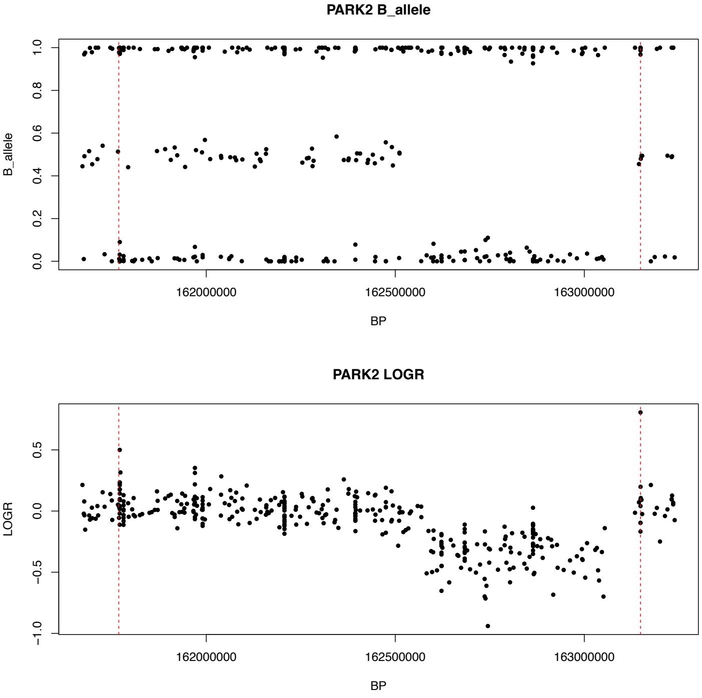

### Patient 28

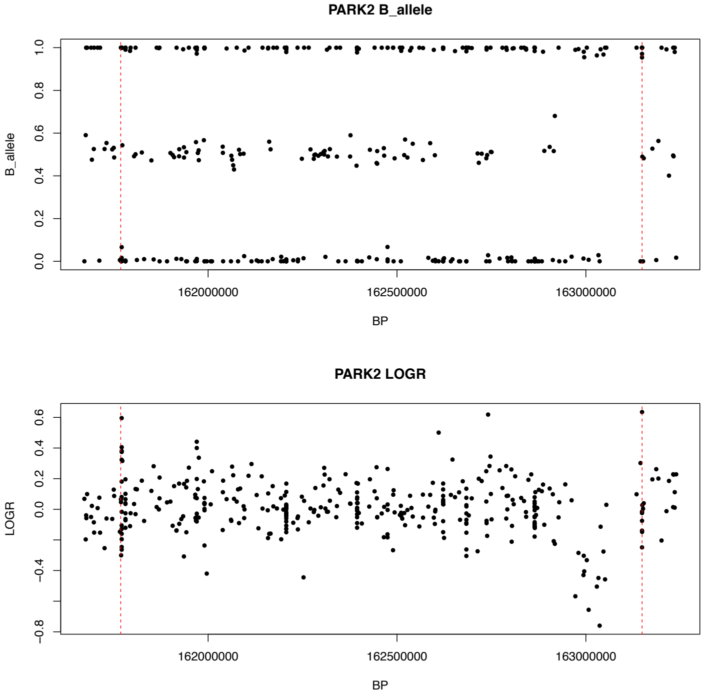

### Patient 29

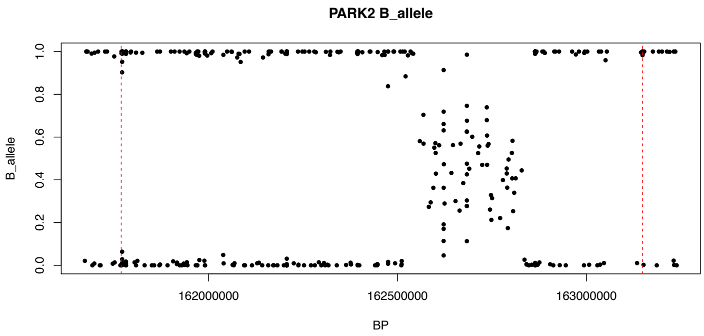

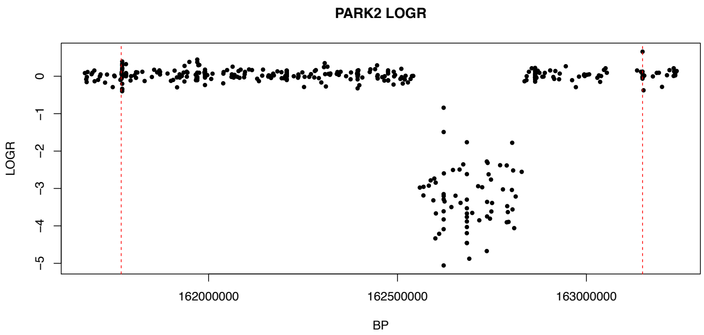

### Patient 30

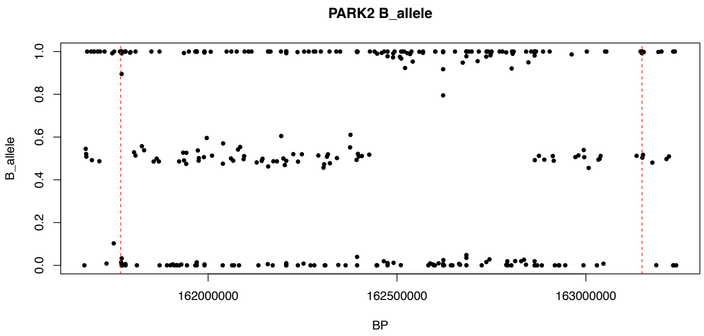

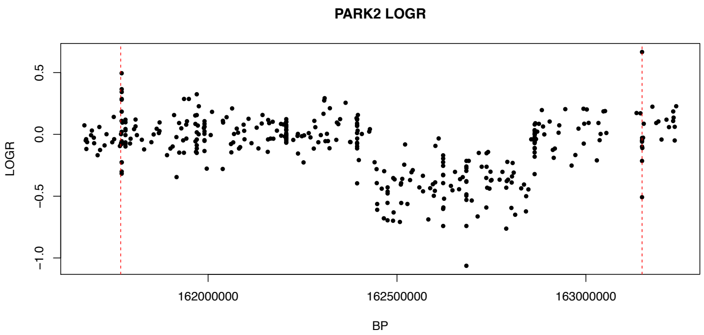

### Patient 31

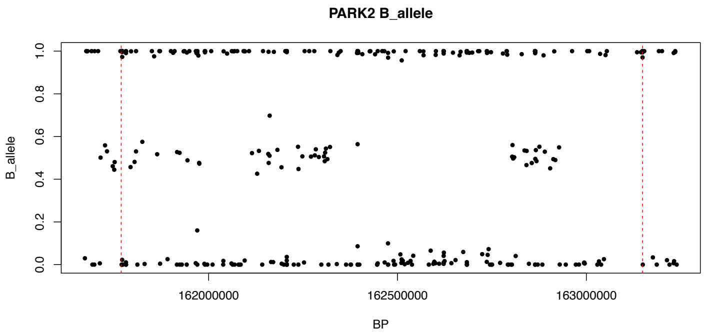

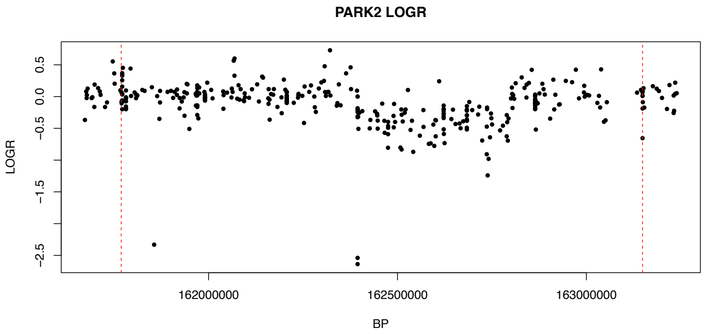

### Patient 32

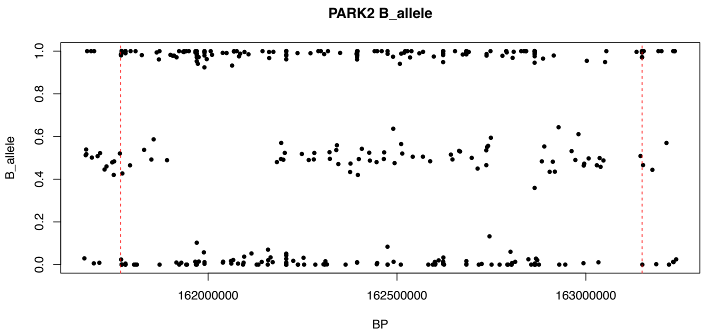

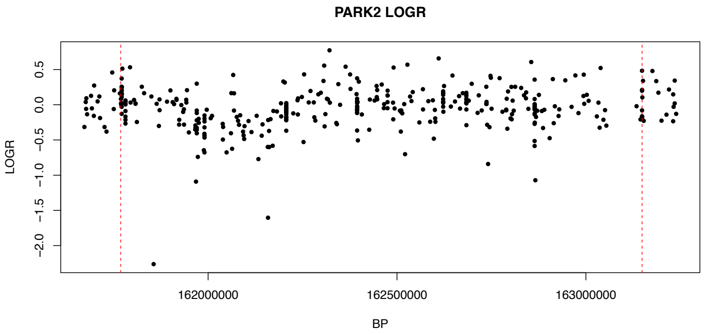

### Patient 33

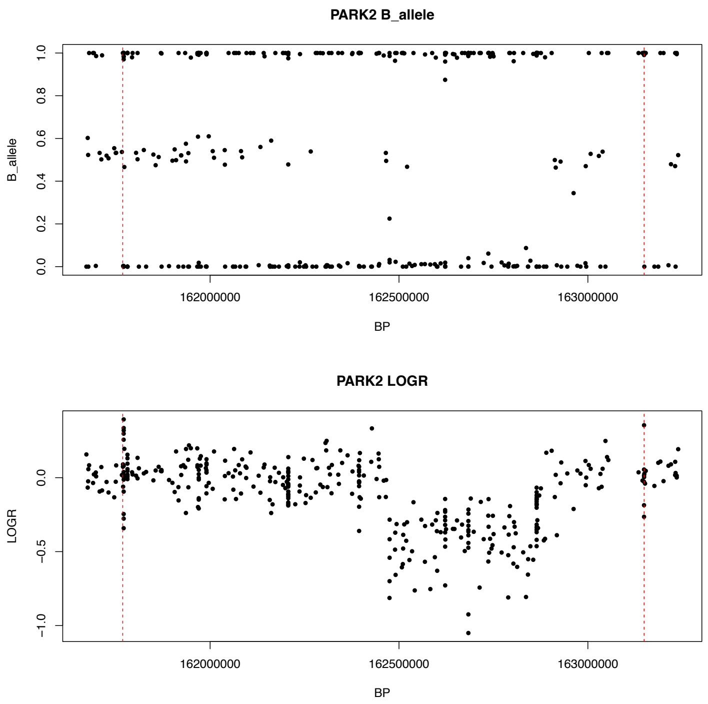

### Patient 34

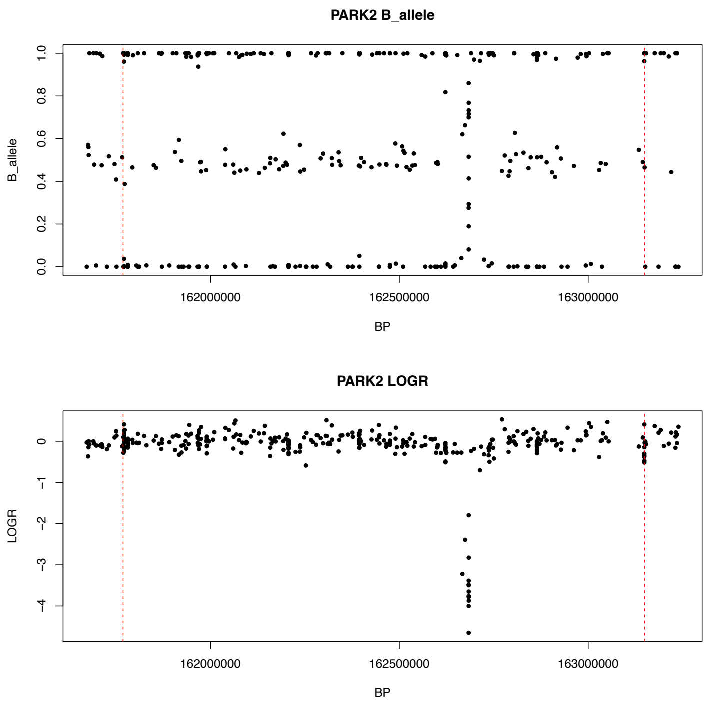

**Supplementary Table 1. Demographic characteristics of the Spanish cohort**

| SITE | Cases (N) | Controls (N) | Total (N) | Female cases (%) | Female control (%) | Case age at onset (mean , SD in years) | Control age at last exam (mean, SD in years) |
|---|---|---|---|---|---|---|---|
| BARCELONA1 | 446 | 455 | 901 | 44,84 | 43,95 | 56.87 (12.36) | 68.53 (9.88) |
| BARCELONA2 | 1026 | 554 | 1580 | 44,54 | 62,27 | 63.19 (12.12) | 65.93 (11.35) |
| BARCELONA3 | 102 | 178 | 280 | 39,21 | 48,31 | 52.53 (12.43) | 46.86 (8.05) |
| BARCELONA4 | 625 | 66 | 691 | 40,48 | 65,15 | 57.02 (12.50) | 63.64 (10.63) |
| SEVILLA | 589 | 1254 | 1843 | 40,23 | 50,31 | 55.07 (12.89) | 53.35 (18.23) |
| MALAGA | 116 | - | 116 | 47,41 | - | 66.4 (9.39) | - |
| MADRID | 9 | 20 | 29 | 66,66 | 60 | 63.22 (7.87) | 52.5 (13.97) |
| OVIEDO | 318 | 67 | 385 | 44,02 | 50,74 | 61.76 (12.95) | 69.89 (5.40) |
| SANTANDER | 411 | 151 | 562 | 43,79 | 65,56 | 64.53 (10.73) | 62.08 (12.28) |
| SAN SEBASTIAN | 804 | 321 | 1125 | 46,01 | 58,25 | 63.46 (11.16) | 73.35 (9.67) |
| GRANADA | 171 | - | 171 | 33,3 | - | 62.24 (11.86) | - |
| SEGOVIA | 166 | - | 166 | 40,82 | - | 68.57 (11.44) | - |
| TOTAL | 4783 | 3066 | 7849 | | | | |

**Supplementary Table 2. Parkinson disease and Parkinsonism related genes under study**

| Gene | Chr | Position | Description |
|---|---|---|---|
| *LRRK2* | 12 | 40590546- 40763087 | Autosomal dominant classical PD |
| *PINK1* | 1 | 20959948 - 20978004 | Autosomal recessive PD |
| *PARK2* | 6 | 161768452 - 163148803 | Autosomal recessive PD |
| *DJ1* | 1 | 8014351 - 8045565 | Autosomal recessive PD |
| *SNCA* | 4 | 90645250 - 90759466 | Autosomal dominant PD with dementia |
| *GBA* | 1 | 155204243 - 155214490 | PD risk with dementia |
| *VPS13C* | 15 | 62144588 - 62352672 | Autosomal recessive PD |
| *CHCHD2* | 7 | 56169262 - 56174269 | Autosomal recessive PD |
| *VPS35* | 16 | 46690054 - 46723430 | Autosomal dominant PD |
| *FBXO7* | 22 | 32870663 - 32894818 | Autosomal recessive PD |
| *ATP13A2* | *1* | 17312453 - 17338423 | Autosomal dominant atypical parkinsonism |
| *PANK2* | *20* | 3869486 - 3907605 | Autosomal recessive, atypical parkinsonism |
| *PLA2G6* | *22* | 38507502 - 38601697 | Autosomal recessive atypical parkinsonism |
| *DCTN1* | *2* | 74588281 - 74619214 | Autosomal dominant atypical parkinsonism |
| *POLG1* | *15* | 89859534 - 89878092 | Autosomal recessive atypical parkinsonism |

Chr; chromosome, Position (referred to hg19)

**Supplementary Table 3. Spanish GWAS replication results of known risk loci identified in Nalls et al., 2019**

| SNP | Chr | Bp | Nearest gene | A1 | A2 | beta | se | maf | P |
|---|---|---|---|---|---|---|---|---|---|
| rs34637584 | 12 | 40734202 | *LRRK2* | A | G | 2,37229 | 0,375093 | 0,00953 | 2,54E-10 |
| rs356182 | 4 | 90626111 | *SNCA* | G | A | 0,223036 | 0,0370765 | 0,34261 | 1,79E-09 |
| rs62053943 | 17 | 43744203 | *CRHR1* | T | C | -0,295669 | 0,0511647 | 0,17489 | 7,52E-09 |
| rs34311866 | 4 | 951947 | *TMEM175* | C | T | 0,242485 | 0,0472576 | 0,16547 | 2,88E-07 |
| rs11150601 | 16 | 30977799 | *SETD1A* | G | A | -0,166897 | 0,0368572 | 0,32723 | 5,95E-06 |
| rs112485576 | 6 | 32578772 | *HLA-DRB5* | A | C | -0,347702 | 0,0768687 | 0,15396 | 6,09E-06 |
| rs117615688 | 17 | 43798308 | *CRHR1* | A | G | -0,297021 | 0,0725574 | 0,09752 | 4,25E-05 |
| rs1474055 | 2 | 169110394 | *STK39* | T | C | 0,187081 | 0,0480997 | 0,15283 | 1,00E-04 |
| rs5019538 | 4 | 90636630 | *SNCA* | G | A | 0,141565 | 0,0372009 | 0,2996 | 0,000142 |
| rs2248244 | 21 | 38852361 | *DYRK1A* | A | G | 0,135448 | 0,0369482 | 0,31448 | 0,000246 |
| rs12456492 | 18 | 40673380 | *RIT2* | G | A | 0,132906 | 0,0364962 | 0,31697 | 0,000271 |
| rs35749011 | 1 | 155135036 | *KRTCAP2* | A | G | 0,750031 | 0,218164 | 0,00823 | 0,000586 |
| rs873786 | 4 | 925376 | *GAK* | T | C | -0,194796 | 0,0577172 | 0,10601 | 0,000738 |
| rs997368 | 6 | 112243291 | *FYN* | G | A | -0,132329 | 0,0395032 | 0,23918 | 0,000809 |
| rs10513789 | 3 | 182760073 | *MCCC1* | G | T | -0,152186 | 0,0464337 | 0,15654 | 0,00105 |
| rs12528068 | 6 | 72487762 | *RIMS1* | T | C | 0,12017 | 0,0374606 | 0,29369 | 0,00134 |
| rs10221156 | 16 | 52969426 | *CHD9* | A | G | -0,183429 | 0,0574851 | 0,09497 | 0,00142 |
| rs620513 | 8 | 16697593 | *FGF20* | T | G | -0,124867 | 0,0396532 | 0,25176 | 0,00164 |
| rs76763715 | 1 | 155205634 | *GBAP1* | C | T | 0,973179 | 0,314228 | 0,00416 | 0,00195 |
| rs6497339 | 16 | 19277493 | *SYT17* | T | A | -0,117843 | 0,0388599 | 0,49192 | 0,00243 |
| rs823118 | 1 | 205723572 | *NUCKS1* | C | T | -0,107755 | 0,0357008 | 0,38694 | 0,00254 |
| rs4698412 | 4 | 15737348 | *BST1* | G | A | -0,0990597 | 0,0337259 | 0,48562 | 0,00331 |
| rs13117519 | 4 | 114369065 | *CAMK2D* | T | C | 0,146826 | 0,0500205 | 0,18663 | 0,00333 |
| rs57891859 | 2 | 135464616 | *TMEM163* | G | A | -0,106623 | 0,0366903 | 0,33598 | 0,00366 |
| rs62333164 | 4 | 170583157 | *CLCN3* | A | G | -0,104158 | 0,0363188 | 0,33008 | 0,00413 |
| rs34025766 | 4 | 17968811 | *LCORL* | A | T | -0,122367 | 0,0431372 | 0,18079 | 0,00456 |
| rs76904798 | 12 | 40614434 | *LRRK2* | T | C | 0,133851 | 0,0485183 | 0,14836 | 0,0058 |
| rs4653767 | 1 | 226916078 | *ITPKB* | C | T | -0,100307 | 0,0366752 | 0,30421 | 0,00624 |
| rs10797576 | 1 | 232664611 | *SIPA1L2* | T | C | 0,121345 | 0,0494854 | 0,14723 | 0,0142 |
| rs12600861 | 17 | 7355621 | *CHRNB1* | C | A | 0,0881833 | 0,0369518 | 0,32999 | 0,017 |
| rs6500328 | 16 | 50736656 | *NOD2* | G | A | -0,0817654 | 0,0350713 | 0,38099 | 0,0197 |
| rs10748818 | 10 | 104015279 | *GBF1* | G | A | 0,118275 | 0,051447 | 0,12725 | 0,0215 |
| rs12951632 | 17 | 40741013 | *RETREG3* | C | T | -0,0853649 | 0,0374104 | 0,29324 | 0,0225 |

| | | | | | | | | | |
|---|---|---|---|---|---|---|---|---|---|
| rs13294100 | 9 | 17579690 | *SH3GL2* | T | G | -0,084288 | 0,0370367 | 0,31128 | 0,0229 |
| rs1867598 | 5 | 60137959 | *ELOVL7* | G | A | 0,119137 | 0,0526915 | 0,123 | 0,0238 |
| rs11658976 | 17 | 44866805 | *WNT3* | G | A | 0,088709 | 0,0394489 | 0,36094 | 0,0245 |
| rs1293298 | 8 | 11712443 | *CTSB* | C | A | -0,101646 | 0,0457512 | 0,26195 | 0,0263 |
| rs199351 | 7 | 23300049 | *GPNMB* | C | A | -0,0695996 | 0,0339813 | 0,45165 | 0,0405 |
| rs2904880 | 16 | 28944396 | *CD19* | C | G | -0,0761626 | 0,0384129 | 0,27357 | 0,0474 |
| rs10756907 | 9 | 17727065 | *SH3GL2* | G | A | 0,0718328 | 0,0373543 | 0,27922 | 0,0545 |
| rs2251086 | 15 | 61997385 | *VPS13C* | T | C | -0,0968268 | 0,0513231 | 0,12557 | 0,0592 |
| rs7938782 | 11 | 10558777 | *RNF141* | G | A | -0,0883287 | 0,046898 | 0,15878 | 0,0596 |
| rs73038319 | 3 | 18361759 | *SATB1* | C | A | 0,262886 | 0,139603 | 0,02077 | 0,0597 |
| rs11950533 | 5 | 134199105 | *C5orf24* | A | C | -0,111364 | 0,059655 | 0,09445 | 0,0619 |
| rs1941685 | 18 | 31304318 | *ASXL3* | T | G | 0,0595049 | 0,034157 | 0,47959 | 0,0815 |
| rs12147950 | 14 | 37989270 | *MIPOL1* | T | C | -0,0621843 | 0,0360926 | 0,45154 | 0,0849 |
| rs11158026 | 14 | 55348869 | *GCH1* | T | C | -0,0658739 | 0,0391027 | 0,25872 | 0,0921 |
| rs55818311 | 19 | 2341047 | *SPPL2B* | C | T | 0,0631666 | 0,0391159 | 0,33663 | 0,106 |
| rs55961674 | 3 | 122196892 | *KPNA1* | T | C | 0,064938 | 0,0419192 | 0,22123 | 0,121 |
| rs76116224 | 2 | 18147848 | *KCNS3* | T | A | -0,111175 | 0,0726913 | 0,06849 | 0,126 |
| rs11557080 | 1 | 205737739 | *RAB29* | A | G | 0,0723827 | 0,0485836 | 0,15112 | 0,136 |
| rs3104783 | 16 | 52636242 | *CASC16* | A | C | 0,0503004 | 0,0343808 | 0,46312 | 0,143 |
| rs114138760 | 1 | 154898185 | *PMVK* | C | G | 0,312074 | 0,217799 | 0,00687 | 0,152 |
| rs72840788 | 10 | 121415685 | *BAG3* | A | G | 0,05769 | 0,0438847 | 0,20042 | 0,189 |
| rs11578699 | 1 | 171719769 | *VAMP4* | T | C | -0,0555585 | 0,0424364 | 0,19475 | 0,19 |
| rs117896735 | 10 | 121536327 | *INPP5F* | A | G | 0,199776 | 0,152455 | 0,01457 | 0,19 |
| rs11610045 | 12 | 133063768 | *FBRSL1* | A | G | 0,0447668 | 0,0345808 | 0,42916 | 0,195 |
| rs186111791 | 12 | 41447669 | *CNTN1* | T | C | 0,370697 | 0,291172 | 0,00499 | 0,203 |
| rs4101061 | 4 | 77147969 | *FAM47E* | G | A | 0,0502194 | 0,0396963 | 0,24956 | 0,206 |
| rs2280104 | 8 | 22525980 | *BIN3* | T | C | 0,0430352 | 0,0343277 | 0,40244 | 0,21 |
| rs8087969 | 18 | 48683589 | *MEX3C* | G | T | 0,0437927 | 0,0359595 | 0,41811 | 0,223 |
| rs61169879 | 17 | 59917366 | *BRIP1* | T | C | 0,0543028 | 0,0462196 | 0,17793 | 0,24 |
| rs6476434 | 9 | 34046391 | *UBAP2* | C | T | 0,045092 | 0,0396047 | 0,25585 | 0,255 |
| rs10847864 | 12 | 123326598 | *HIP1R* | T | G | 0,0427109 | 0,0401887 | 0,37649 | 0,288 |
| rs6854006 | 4 | 77198054 | *FAM47E-STBD1* | T | C | -0,0365313 | 0,0347406 | 0,36931 | 0,293 |
| rs3802920 | 11 | 133787001 | *IGSF9B* | T | G | 0,0426299 | 0,0429361 | 0,21077 | 0,321 |
| rs4140646 | 6 | 27738801 | *LOC100131289* | A | G | 0,0431854 | 0,0441091 | 0,2068 | 0,328 |
| rs75859381 | 6 | 133210361 | *RPS12* | C | T | 0,119614 | 0,128253 | 0,02911 | 0,351 |
| rs1450522 | 3 | 161077630 | *SPTSSB* | G | A | 0,0342789 | 0,0368314 | 0,30585 | 0,352 |
| rs6808178 | 3 | 28705690 | *LINC00693* | T | C | 0,0317566 | 0,0353169 | 0,38927 | 0,369 |

| | | | | | | | | | |
|---|---|---|---|---|---|---|---|---|---|
| rs12497850 | 3 | 48748989 | *IP6K2* | G | T | 0,0307309 | 0,0359268 | 0,33606 | 0,392 |
| rs77351827 | 20 | 6006041 | *CRLS1* | T | C | 0,0475474 | 0,0556763 | 0,11574 | 0,393 |
| rs9568188 | 13 | 49927732 | *CAB39L* | C | T | -0,0335308 | 0,04227 | 0,20747 | 0,428 |
| rs4771268 | 13 | 97865021 | *MBNL2* | T | C | 0,0297526 | 0,0387084 | 0,26355 | 0,442 |
| rs3742785 | 14 | 75373034 | *RPS6KL1* | C | A | -0,0319663 | 0,0418254 | 0,19833 | 0,445 |
| rs2086641 | 8 | 130901909 | *FAM49B* | C | T | 0,0283826 | 0,0374003 | 0,29747 | 0,448 |
| rs850738 | 17 | 42434630 | *FAM171A2* | G | A | 0,0267449 | 0,0352205 | 0,36902 | 0,448 |
| rs26431 | 5 | 102365794 | *PAM* | G | C | -0,0214186 | 0,0375211 | 0,29197 | 0,568 |
| rs896435 | 10 | 15557406 | *ITGA8* | C | T | 0,0203307 | 0,0378775 | 0,28273 | 0,591 |
| rs2269906 | 17 | 42294337 | *UBTF* | C | A | -0,0184981 | 0,0356335 | 0,3484 | 0,604 |
| rs6825004 | 4 | 77110365 | *SCARB2* | G | C | -0,0164801 | 0,0359628 | 0,34469 | 0,647 |
| rs6658353 | 1 | 161469054 | *FCGR2A* | G | C | 0,0150915 | 0,0345883 | 0,48055 | 0,663 |
| rs7134559 | 12 | 46419086 | *SCAF11* | T | C | -0,0146968 | 0,0357578 | 0,41628 | 0,681 |
| rs11707416 | 3 | 151108965 | *MED12L* | A | T | -0,00965181 | 0,0347941 | 0,39043 | 0,781 |
| rs666463 | 17 | 76425480 | *DNAH17* | T | A | 0,0121239 | 0,0453111 | 0,1726 | 0,789 |
| rs979812 | 14 | 88464264 | *GALC* | T | G | 0,00870694 | 0,0339772 | 0,45541 | 0,798 |
| rs9261484 | 6 | 30108683 | *TRIM40* | T | C | 0,00922 | 0,0441818 | 0,19516 | 0,835 |
| rs2042477 | 2 | 96000943 | *KCNIP3* | A | T | 0,00609432 | 0,0437241 | 0,18973 | 0,889 |
| rs11683001 | 2 | 102396963 | *MAP4K4* | A | T | -0,00277821 | 0,0362529 | 0,33456 | 0,939 |
| rs200564078 | 16 | 58587672 | *CNOT1* | T | C | 0,0269392 | 0,391422 | 0,00292 | 0,945 |
| rs76949143 | 7 | 66009851 | *GS1-124K5.11* | A | T | 0,00428635 | 0,076004 | 0,05437 | 0,955 |
| rs12283611 | 11 | 83487277 | *DLG2* | A | C | -0,00169366 | 0,0345875 | 0,45023 | 0,961 |

**Supplementary Table 4. Risk haplotype estimates at consensus genotyped variants for known GWAS loci in Spanish and UK PD cases.**

| | | | | SPAIN | | UK | |
|---|---|---|---|---|---|---|---|
| **SNP** | **Nearest gene** | **Chr** | **Bp** | **Haplotype Length** | **NSNPs** | **Haplotype Length** | **NSNPs** |
| rs112485576 | *HLA-DRB5* | 6 | 32578772 | 16,663 | 3 | 144,918 | 8 |
| rs4140646 | *LOC100131289* | 6 | 27738801 | 12,65 | 2 | 124,193 | 9 |
| rs34025766 | *LCORL* | 4 | 17968811 | 73,6 | 5 | 151,946 | 6 |
| rs12497850 | *IP6K2* | 3 | 48748989 | 20,885 | 3 | 90,116 | 5 |
| rs12951632 | *RETREG3* | 17 | 40741013 | 89,325 | 5 | 149,802 | 6 |
| rs9261484 | *TRIM40* | 6 | 30108683 | 24,795 | 3 | 51,246 | 8 |
| rs2280104 | *BIN3* | 8 | 22525980 | 48,396 | 5 | 62,358 | 7 |
| rs11578699 | *VAMP4* | 1 | 171719769 | 79,35 | 9 | 91,045 | 9 |
| rs2248244 | *DYRK1A* | 21 | 38852361 | 117,828 | 7 | 125,939 | 8 |
| rs1450522 | *SPTSSB* | 3 | 161077630 | 53,4 | 5 | 58,993 | 7 |
| rs12456492 | *RIT2* | 18 | 40673380 | 1,162 | 2 | 4,856 | 3 |
| rs34637584 | *LRRK2* | 12 | 40734202 | 15,802 | 7 | 15,802 | 7 |
| rs62053943 | *CRHR1/MAPT* | 17 | 43744203 | 120,057 | 4 | 120,057 | 4 |
| rs1474055 | *STK39* | 2 | 169110394 | 34,519 | 6 | 34,519 | 6 |
| rs12528068 | *RIMS1* | 6 | 72487762 | 11,622 | 2 | 11,622 | 2 |
| rs10221156 | *CHD9* | 16 | 52969426 | 19,828 | 3 | 19,828 | 3 |
| rs4698412 | *BST1* | 4 | 15737348 | 2,151 | 2 | 2,151 | 2 |
| rs12600861 | *CHRNB1* | 17 | 7355621 | 12,545 | 2 | 12,545 | 2 |
| rs10748818 | *GBF1* | 10 | 104015279 | 123,92 | 7 | 123,92 | 7 |
| rs199351 | *GPNMB* | 7 | 23300049 | 5,906 | 3 | 5,906 | 3 |
| rs11158026 | *GCH1* | 14 | 55348869 | 2,579 | 2 | 2,579 | 2 |
| rs55818311 | *SPPL2B* | 19 | 2341047 | 16,527 | 2 | 16,527 | 2 |
| rs55961674 | *KPNA1* | 3 | 122196892 | 175,314 | 11 | 175,314 | 11 |
| rs117896735 | *INPP5F* | 10 | 121536327 | 100,641 | 5 | 100,641 | 5 |
| rs11610045 | *FBRSL1* | 12 | 133063768 | 8,272 | 2 | 8,272 | 2 |
| rs186111791 | *CNTN1* | 12 | 41447669 | 74,369 | 4 | 74,369 | 4 |
| rs3802920 | *IGSF9B* | 11 | 133787001 | 8,9 | 3 | 8,9 | 3 |
| rs4771268 | *MBNL2* | 13 | 97865021 | 20,404 | 3 | 20,404 | 3 |
| rs26431 | *PAM* | 5 | 102365794 | 168,367 | 4 | 168,367 | 4 |
| rs6825004 | *SCARB2* | 4 | 77110365 | 31,909 | 5 | 31,909 | 5 |
| rs6658353 | *FCGR2A* | 1 | 161469054 | 15,932 | 2 | 15,932 | 2 |
| rs11707416 | *MED12L* | 3 | 151108965 | 41,783 | 5 | 41,783 | 5 |

| | | | | | | | |
|---|---|---|---|---|---|---|---|
| rs979812 | *GALC* | 14 | 88464264 | 66,813 | 8 | 66,813 | 8 |
| rs200564078 | *CNOT1* | 16 | 58587672 | 123,499 | 6 | 123,499 | 6 |
| rs6854006 | *FAM47E-STBD1* | 4 | 77198054 | 25,872 | 8 | 15,073 | 7 |
| rs6500328 | *NOD2* | 16 | 50736656 | 43,248 | 6 | 31,171 | 5 |
| rs356182 | *SNCA* | 4 | 90626111 | NA | NA | NA | NA |
| rs34311866 | *TMEM175* | 4 | 951947 | NA | NA | NA | NA |
| rs11150601 | *SETD1A* | 16 | 30977799 | NA | NA | NA | NA |
| rs117615688 | *CRHR1* | 17 | 43798308 | NA | NA | NA | NA |
| rs5019538 | *SNCA* | 4 | 90636630 | NA | NA | NA | NA |
| rs35749011 | *KRTCAP2* | 1 | 155135036 | NA | NA | NA | NA |
| rs873786 | *GAK* | 4 | 925376 | 50,947 | 4 | NA | NA |
| rs997368 | *FYN* | 6 | 112243291 | NA | NA | NA | NA |
| rs10513789 | *MCCC1* | 3 | 182760073 | NA | NA | NA | NA |
| rs620513 | *FGF20* | 8 | 16697593 | NA | NA | NA | NA |
| rs76763715 | *GBAP1* | 1 | 155205634 | NA | NA | 19,494 | 2 |
| rs6497339 | *SYT17* | 16 | 19277493 | NA | NA | NA | NA |
| rs823118 | *NUCKS1* | 1 | 205723572 | NA | NA | NA | NA |
| rs13117519 | *CAMK2D* | 4 | 114369065 | NA | NA | NA | NA |
| rs57891859 | *TMEM163* | 2 | 135464616 | NA | NA | NA | NA |
| rs62333164 | *CLCN3* | 4 | 170583157 | NA | NA | NA | NA |
| rs76904798 | *LRRK2* | 12 | 40614434 | NA | NA | NA | NA |
| rs4653767 | *ITPKB* | 1 | 226916078 | NA | NA | NA | NA |
| rs10797576 | *SIPA1L2* | 1 | 232664611 | NA | NA | NA | NA |
| rs13294100 | *SH3GL2* | 9 | 17579690 | NA | NA | 8,971 | 2 |
| rs1867598 | *ELOVL7* | 5 | 60137959 | NA | NA | NA | NA |
| rs11658976 | *WNT3* | 17 | 44866805 | NA | NA | NA | NA |
| rs1293298 | *CTSB* | 8 | 11712443 | NA | NA | NA | NA |
| rs2904880 | *CD19* | 16 | 28944396 | NA | NA | NA | NA |
| rs10756907 | *SH3GL2* | 9 | 17727065 | NA | NA | NA | NA |
| rs2251086 | *VPS13C* | 15 | 61997385 | NA | NA | NA | NA |
| rs7938782 | *RNF141* | 11 | 10558777 | NA | NA | NA | NA |
| rs73038319 | *SATB1* | 3 | 18361759 | NA | NA | NA | NA |
| rs11950533 | *C5orf24* | 5 | 134199105 | NA | NA | NA | NA |
| rs1941685 | *ASXL3* | 18 | 31304318 | NA | NA | NA | NA |
| rs12147950 | *MIPOL1* | 14 | 37989270 | NA | NA | NA | NA |
| rs76116224 | *KCNS3* | 2 | 18147848 | NA | NA | NA | NA |

| | | | | | | | |
|---|---|---|---|---|---|---|---|
| rs11557080 | *RAB29* | 1 | 205737739 | NA | NA | NA | NA |
| rs3104783 | *CASC16* | 16 | 52636242 | NA | NA | NA | NA |
| rs114138760 | *PMVK* | 1 | 154898185 | NA | NA | NA | NA |
| rs72840788 | *BAG3* | 10 | 121415685 | NA | NA | NA | NA |
| rs4101061 | *FAM47E* | 4 | 77147969 | NA | NA | NA | NA |
| rs8087969 | *MEX3C* | 18 | 48683589 | NA | NA | NA | NA |
| rs61169879 | *BRIP1* | 17 | 59917366 | NA | NA | NA | NA |
| rs6476434 | *UBAP2* | 9 | 34046391 | NA | NA | NA | NA |
| rs10847864 | *HIP1R* | 12 | 123326598 | NA | NA | NA | NA |
| rs75859381 | *RPS12* | 6 | 133210361 | NA | NA | 52,65 | 5 |
| rs6808178 | *LINC00693* | 3 | 28705690 | NA | NA | NA | NA |
| rs77351827 | *CRLS1* | 20 | 6006041 | NA | NA | NA | NA |
| rs9568188 | *CAB39L* | 13 | 49927732 | NA | NA | NA | NA |
| rs3742785 | *RPS6KL1* | 14 | 75373034 | NA | NA | NA | NA |
| rs2086641 | *FAM49B* | 8 | 130901909 | NA | NA | NA | NA |
| rs850738 | *FAM171A2* | 17 | 42434630 | NA | NA | NA | NA |
| rs896435 | *ITGA8* | 10 | 15557406 | NA | NA | NA | NA |
| rs2269906 | *UBTF* | 17 | 42294337 | NA | NA | NA | NA |
| rs7134559 | *SCAF11* | 12 | 46419086 | NA | NA | NA | NA |
| rs666463 | *DNAH17* | 17 | 76425480 | NA | NA | NA | NA |
| rs2042477 | *KCNIP3* | 2 | 96000943 | NA | NA | NA | NA |
| rs11683001 | *MAP4K4* | 2 | 102396963 | NA | NA | 147,136 | 4 |
| rs76949143 | *GS1-124K5.11* | 7 | 66009851 | NA | NA | NA | NA |
| rs12283611 | *DLG2* | 11 | 83487277 | NA | NA | NA | NA |

Chr; chromosome, NSNPs; number of SNPs, Bp; base pair

NA: Due to poor coverage haplotypes could not be constrcuted for these regions in both UK and SPAIN datasets; Length is specified in Kilobases.

**Supplementary Table 5. Cross-trait genetic correlations by linkage disequilibrium score regression**

| Trait | Source | rg (se) | z | p-value | FDR p-value | h² obs [se] | h² int [se] | gcov int [se] |
|---|---|---|---|---|---|---|---|---|
| Types of physical activity in last 4 weeks: Light DIY (eg: pruning_ watering the lawn) | 0 | 0.2533(0.0769) | 3,2928 | 1,00E-03 | 7,50E-01 | 0.0384(0.0022) | 1.0078(0.0084) | -0.0092(0.0048) |
| Subjective well being | 27089181 | -0.2596(0.0869) | -2,9872 | 2,80E-03 | 7,50E-01 | 0.0246(0.0021) | 1.0015(0.0074) | 0.0118(0.0044) |
| Average weekly spirits intake | 0 | -0.2588(0.0922) | -2,8081 | 5,00E-03 | 7,50E-01 | 0.031(0.0027) | 1.0177(0.0079) | 0.0084(0.0046) |
| Blood clot_ DVT_ bronchitis_ emphysema_ asthma_ rhinitis_ eczema_ allergy diagnosed by doctor: Blood clot in the leg (DVT) | 0 | -0.3371(0.127) | -2,6539 | 8,00E-03 | 7,50E-01 | 0.0104(0.0017) | 1.0108(0.0092) | 0.0078(0.005) |
| Illnesses of mother: Heart disease | 0 | -0.3058(0.1188) | -2,5728 | 1,01E-02 | 7,50E-01 | 0.0154(0.0018) | 1.0109(0.0067) | 0.0088(0.0048) |
| Vascular/heart problems diagnosed by doctor: Heart attack | 0 | -0.2685(0.1044) | -2,5725 | 1,01E-02 | 7,50E-01 | 0.0179(0.0017) | 1.0141(0.0074) | 0.0097(0.005) |
| Cancer diagnosed by doctor | 0 | -0.4077(0.1602) | -2,5441 | 1,10E-02 | 7,50E-01 | 0.0062(0.0016) | 1.0101(0.0064) | 0.0074(0.0047) |
| Cancer code_ self-reported: breast cancer | 0 | -0.3491(0.1379) | -2,5323 | 1,13E-02 | 7,50E-01 | 0.0095(0.0018) | 1.019(0.0079) | 0.013(0.0047) |
| Non-cancer illness code_ self-reported: deep venous thrombosis (dvt) | 0 | -0.3061(0.1236) | -2,4761 | 1,33E-02 | 7,50E-01 | 0.0109(0.0018) | 1.0084(0.0092) | 0.0075(0.005) |
| Tinnitus: Yes_ but not now_ but have in the past | 0 | -0.4175(0.1703) | -2,4518 | 1,42E-02 | 7,50E-01 | 0.0184(0.0044) | 0.9917(0.0064) | 0.0104(0.0044) |
| Non-cancer illness code_ self-reported: heart attack/myocardial infarction | 0 | -0.2539(0.1054) | -2,4093 | 1,60E-02 | 7,50E-01 | 0.0183(0.0017) | 1.0123(0.0073) | 0.0089(0.005) |
| Blood clot_ DVT_ bronchitis_ emphysema_ asthma_ rhinitis_ eczema_ allergy diagnosed by doctor: None of the above | 0 | 0.1425(0.06) | 2,3775 | 1,74E-02 | 7,50E-01 | 0.0734(0.006) | 1.0429(0.0166) | -0.004(0.0052) |
| Vitamin and mineral supplements: Multivitamins +/- minerals | 0 | 0.2157(0.0922) | 2,3405 | 1,93E-02 | 7,50E-01 | 0.0234(0.0019) | 1.0081(0.0076) | -0.0097(0.0048) |
| Arm predicted mass (right) | 0 | -0.1205(0.052) | -2,3181 | 2,04E-02 | 7,50E-01 | 0.2629(0.011) | 1.1068(0.0273) | 0.005(0.0072) |
| ICV | 25607358 | 0.4156(0.1817) | 2,2876 | 2,22E-02 | 7,50E-01 | 0.1679(0.0445) | 1.0025(0.0066) | -0.0087(0.0045) |
| Arm fat-free mass (right) | 0 | -0.1176(0.0522) | -2,2518 | 2,43E-02 | 7,50E-01 | 0.2642(0.011) | 1.1051(0.0272) | 0.0048(0.0072) |
| Non-cancer illness code_ self-reported: pneumothorax | 0 | 0.5735(0.2553) | 2,2465 | 2,47E-02 | 7,50E-01 | 0.0028(0.0015) | 0.988(0.007) | -0.0094(0.0045) |
| Current employment status: Doing unpaid or voluntary work | 0 | -0.2866(0.1298) | -2,208 | 2,72E-02 | 7,50E-01 | 0.0093(0.0014) | 1.0079(0.0062) | 0.0115(0.0046) |
| Chest pain or discomfort walking normally | 0 | -0.276(0.1255) | -2,1986 | 2,79E-02 | 7,50E-01 | 0.0739(0.0095) | 1.0095(0.0071) | 0.0048(0.0047) |
| Pain type(s) experienced in last month: Knee pain | 0 | -0.1537(0.0712) | -2,1582 | 3,09E-02 | 7,50E-01 | 0.0406(0.0024) | 1.015(0.0083) | 0.0057(0.0048) |
| Illnesses of mother: None of the above (group 1) | 0 | 0.2379(0.1105) | 2,1539 | 3,13E-02 | 7,50E-01 | 0.017(0.0018) | 1.0123(0.0072) | -0.0065(0.0054) |
| Arm fat-free mass (left) | 0 | -0.1101(0.0514) | -2,1402 | 3,23E-02 | 7,50E-01 | 0.2632(0.0108) | 1.1115(0.0262) | 0.0047(0.0072) |
| Vitamin and mineral supplements: None of the above | 0 | -0.1818(0.0855) | -2,1265 | 3,35E-02 | 7,50E-01 | 0.0301(0.0018) | 0.9938(0.0066) | 0.0078(0.0048) |
| Number of unsuccessful stop-smoking attempts | 0 | -0.2842(0.1342) | -2,1183 | 3,41E-02 | 7,50E-01 | 0.049(0.0074) | 1.0073(0.0069) | 0.0051(0.0047) |
| Diagnoses - main ICD10: C50 Malignant neoplasm of breast | 0 | -0.3017(0.1428) | -2,1125 | 3,46E-02 | 7,50E-01 | 0.009(0.0017) | 1.0018(0.0075) | 0.0093(0.0046) |
| Number of self-reported cancers | 0 | -0.3337(0.1588) | -2,1014 | 3,56E-02 | 7,50E-01 | 0.0061(0.0015) | 1.0129(0.0065) | 0.0068(0.0045) |
| Arm predicted mass (left) | 0 | -0.1082(0.0517) | -2,0951 | 3,62E-02 | 7,50E-01 | 0.2624(0.0108) | 1.1136(0.0265) | 0.0049(0.0071) |
| Childhood IQ | 23358156 | 0.312(0.1497) | 2,0838 | 3,72E-02 | 7,50E-01 | 0.2751(0.0469) | 1.002(0.0095) | -0.0154(0.0062) |
| Cough on most days | 0 | 0.2605(0.1252) | 2,0807 | 3,75E-02 | 7,50E-01 | 0.0477(0.0063) | 0.9989(0.0071) | -0.0038(0.0048) |
| Whole body water mass | 0 | -0.1075(0.0518) | -2,0742 | 3,81E-02 | 7,50E-01 | 0.2873(0.0123) | 1.1298(0.0294) | 0.0034(0.0073) |
| Basal metabolic rate | 0 | -0.1057(0.0511) | -2,0667 | 3,88E-02 | 7,50E-01 | 0.283(0.0116) | 1.1167(0.0276) | 0.0033(0.0073) |
| Leg predicted mass (right) | 0 | -0.106(0.0514) | -2,0627 | 3,91E-02 | 7,50E-01 | 0.2631(0.0108) | 1.1121(0.0259) | 0.0035(0.0072) |
| Leg fat-free mass (right) | 0 | -0.1048(0.0513) | -2,0423 | 4,11E-02 | 7,50E-01 | 0.2633(0.0108) | 1.1123(0.0259) | 0.0034(0.0072) |
| Whole body fat-free mass | 0 | -0.1061(0.0521) | -2,0379 | 4,16E-02 | 7,50E-01 | 0.2873(0.0123) | 1.1305(0.0297) | 0.0029(0.0074) |
| Types of physical activity in last 4 weeks: None of the above | 0 | -0.1861(0.0916) | -2,0323 | 4,21E-02 | 7,50E-01 | 0.024(0.0017) | 1.0162(0.0071) | 0.0048(0.0048) |
| Nervous feelings | 0 | 0.1224(0.0604) | 2,0277 | 4,26E-02 | 7,50E-01 | 0.0656(0.0039) | 1.0149(0.0125) | -0.003(0.0052) |
| Impedance of arm (right) | 0 | 0.0943(0.0465) | 2,0269 | 4,27E-02 | 7,50E-01 | 0.2346(0.0083) | 1.0829(0.0222) | -0.003(0.0062) |
| Mouth/teeth dental problems: Bleeding gums | 0 | -0.1888(0.0934) | -2,0209 | 4,33E-02 | 7,50E-01 | 0.0221(0.0019) | 1.0114(0.0079) | 0.0079(0.005) |
| Relative age of first facial hair | 0 | 0.1464(0.0731) | 2,0015 | 4,53E-02 | 7,50E-01 | 0.1201(0.0075) | 1.0479(0.0149) | -0.0041(0.0056) |
| Leg predicted mass (left) | 0 | -0.1(0.0507) | -1,9736 | 4,84E-02 | 7,50E-01 | 0.2653(0.0108) | 1.1049(0.0251) | 0.0022(0.0071) |
| Number of operations_ self-reported | 0 | -0.1694(0.0859) | -1,9712 | 4,87E-02 | 7,50E-01 | 0.0379(0.0023) | 1.0372(0.0078) | 0.0041(0.0054) |
| Leg fat-free mass (left) | 0 | -0.0996(0.0507) | -1,9639 | 4,95E-02 | 7,50E-01 | 0.2657(0.0108) | 1.1053(0.0252) | 0.0021(0.0071) |
| Medication for cholesterol_ blood pressure or diabetes: None of the above | 0 | 0.1375(0.0706) | 1,9474 | 5,15E-02 | 7,50E-01 | 0.0878(0.0051) | 1.0323(0.01) | -0.0058(0.0051) |
| Ever smoked | 0 | -0.1273(0.0656) | -1,9406 | 5,23E-02 | 7,50E-01 | 0.0734(0.0032) | 1.0255(0.0094) | 0.0019(0.005) |
| Reason for glasses/contact lenses: For just reading/near work as you are getting older (called presbyopia) | 0 | -0.3957(0.2043) | -1,9367 | 5,28E-02 | 7,50E-01 | 0.0041(0.0017) | 1.1351(0.0082) | 0.0079(0.0051) |
| Non-cancer illness code_ self-reported: diabetes | 0 | -0.1544(0.0798) | -1,9361 | 5,29E-02 | 7,50E-01 | 0.0356(0.0025) | 1.0359(0.0099) | 0.0113(0.005) |
| Childhood obesity | 22484627 | -0.1909(0.0987) | -1,9343 | 5,31E-02 | 7,50E-01 | 0.4104(0.0466) | 0.9261(0.0075) | 0.008(0.0045) |
| Tinnitus: Yes_ now some of the time | 0 | 0.4115(0.2141) | 1,9222 | 5,46E-02 | 7,50E-01 | 0.0099(0.0042) | 0.9978(0.0062) | -0.0022(0.0045) |
| Trunk predicted mass | 0 | -0.1024(0.0535) | -1,9137 | 5,57E-02 | 7,50E-01 | 0.283(0.0128) | 1.1384(0.0312) | 0.0027(0.0074) |
| Weight | 0 | -0.0928(0.0485) | -1,9121 | 5,59E-02 | 7,50E-01 | 0.2647(0.0095) | 1.085(0.0219) | 0.0041(0.007) |
| Rheumatoid Arthritis | 24390342 | -0.1549(0.0811) | -1,9105 | 5,61E-02 | 7,50E-01 | 0.1599(0.0295) | 1.0207(0.0158) | 0.0022(0.0048) |
| Medication for cholesterol_ blood pressure or diabetes: Blood pressure medication | 0 | -0.1263(0.0662) | -1,9077 | 5,64E-02 | 7,50E-01 | 0.1035(0.0056) | 1.0318(0.0101) | 0.0065(0.0055) |
| Length of menstrual cycle | 0 | 0.2743(0.1441) | 1,9044 | 5,69E-02 | 7,50E-01 | 0.1024(0.0207) | 0.9993(0.007) | -0.0068(0.0046) |
| Happiness | 0 | 0.1925(0.1024) | 1,8811 | 6,00E-02 | 7,50E-01 | 0.0613(0.0047) | 1.0053(0.0073) | -0.0022(0.0049) |
| Trunk fat-free mass | 0 | -0.1007(0.0536) | -1,8789 | 6,03E-02 | 7,50E-01 | 0.2845(0.0128) | 1.1392(0.0314) | 0.0023(0.0074) |
| Past tobacco smoking | 0 | 0.1139(0.0608) | 1,8722 | 6,12E-02 | 7,50E-01 | 0.0894(0.0039) | 1.0244(0.0099) | -0.0012(0.0051) |
| Parkinsons disease | 19915575 | -0.2352(0.126) | -1,8674 | 6,18E-02 | 7,50E-01 | 0.3592(0.1126) | 1.1248(0.0087) | 0.0053(0.0048) |
| Impedance of arm (left) | 0 | 0.0881(0.0476) | 1,8517 | 6,41E-02 | 7,50E-01 | 0.2311(0.0084) | 1.0989(0.0221) | -0.0028(0.0063) |
| Leptin_not_adjBMI | 26833098 | -0.2363(0.1278) | -1,8487 | 6,45E-02 | 7,50E-01 | 0.0956(0.0168) | 0.9955(0.0066) | 0.0097(0.0045) |
| Waist circumference | 0 | -0.09(0.0489) | -1,8409 | 6,56E-02 | 7,50E-01 | 0.2(0.0071) | 1.0622(0.0167) | 0.0018(0.0064) |
| Qualifications: O levels/GCSEs or equivalent | 0 | 0.1364(0.075) | 1,8184 | 6,90E-02 | 7,50E-01 | 0.0479(0.0024) | 1.0181(0.0093) | -0.0089(0.0048) |
| Neck/shoulder pain for 3+ months | 0 | 0.3487(0.1938) | 1,7992 | 7,20E-02 | 7,50E-01 | 0.0207(0.006) | 1.0084(0.0071) | -0.0041(0.0046) |
| Duration of vigorous activity | 0 | 0.182(0.1012) | 1,7983 | 7,21E-02 | 7,50E-01 | 0.0361(0.0033) | 0.9916(0.0074) | -0.0051(0.0048) |
| Body mass index (BMI) | 0 | -0.0826(0.046) | -1,7952 | 7,26E-02 | 7,50E-01 | 0.247(0.0082) | 1.0732(0.0185) | 0.0046(0.0067) |
| Asthma | 17611496 | -0.2657(0.1484) | -1,7907 | 7,33E-02 | 7,50E-01 | 0.1121(0.0261) | 1.0137(0.0083) | 0.0017(0.0052) |
| Non-cancer illness code_ self-reported: varicose veins | 0 | -0.3252(0.182) | -1,7871 | 7,39E-02 | 7,50E-01 | 0.0043(0.0014) | 1.0122(0.0063) | -0.0011(0.0044) |
| Falls in the last year | 0 | -0.1316(0.0737) | -1,7857 | 7,41E-02 | 7,50E-01 | 0.0327(0.002) | 1.0037(0.0074) | -0.0014(0.0048) |

| | | | | | | | | |
|---|---|---|---|---|---|---|---|---|
| Forced expiratory volume in 1 second (FEV1) | 28166213 | 0.1422(0.08) | 1,7772 | 7,55E-02 | 7,50E-01 | 0.2579(0.0163) | 0.9722(0.0067) | -0.0044(0.0044) |
| Attention deficit hyperactivity disorder (No GC) | 27663945 | -0.3796(0.2143) | -1,7712 | 7,65E-02 | 7,50E-01 | 0.0729(0.0304) | 1.0076(0.0075) | 0.0076(0.0051) |
| Attention deficit hyperactivity disorder (GC) | 27663945 | -0.3808(0.2152) | -1,7695 | 7,68E-02 | 7,50E-01 | 0.0718(0.03) | 0.993(0.0073) | 0.0076(0.0051) |
| Major depressive disorder | 22472876 | 0.2267(0.1288) | 1,7598 | 7,84E-02 | 7,50E-01 | 0.1682(0.0255) | 1.0044(0.0062) | -0.0072(0.0042) |
| Diagnoses - main ICD10: K76 Other diseases of liver | 0 | 0.383(0.2179) | 1,7577 | 7,88E-02 | 7,50E-01 | 0.0038(0.0012) | 0.9927(0.0063) | -0.0098(0.0047) |
| Transport type for commuting to job workplace: Car/motor vehicle | 0 | -0.2245(0.1283) | -1,7498 | 8,02E-02 | 7,50E-01 | 0.0238(0.003) | 1.0243(0.007) | 0.0031(0.0048) |
| Arm fat mass (left) | 0 | -0.0814(0.0468) | -1,7381 | 8,22E-02 | 7,50E-01 | 0.2316(0.0078) | 1.0667(0.0172) | 0.0049(0.0067) |
| Leg fat mass (right) | 0 | -0.0822(0.0474) | -1,7333 | 8,30E-02 | 7,50E-01 | 0.2305(0.0076) | 1.0695(0.0175) | 0.0049(0.0066) |
| Diagnoses - main ICD10: I83 Varicose veins of lower extremities | 0 | -0.1656(0.0956) | -1,7335 | 8,30E-02 | 7,50E-01 | 0.0185(0.0023) | 1.0362(0.0112) | 0.0017(0.0046) |
| Diabetes diagnosed by doctor | 0 | -0.1255(0.0725) | -1,7303 | 8,36E-02 | 7,50E-01 | 0.0434(0.0027) | 1.038(0.0104) | 0.0076(0.0049) |
| Leg fat mass (left) | 0 | -0.082(0.0474) | -1,7299 | 8,36E-02 | 7,50E-01 | 0.2315(0.0076) | 1.0704(0.0174) | 0.0047(0.0067) |
| Fractured bone site(s): Other bones | 0 | -0.2446(0.1418) | -1,7251 | 8,45E-02 | 7,50E-01 | 0.0092(0.0017) | 0.9995(0.007) | 0.0034(0.0047) |
| Forced Vital capacity(FVC) | 28166213 | 0.1375(0.0802) | 1,7138 | 8,66E-02 | 7,50E-01 | 0.254(0.0151) | 0.9778(0.0068) | -0.0021(0.0045) |
| Types of physical activity in last 4 weeks: Heavy DIY (eg: weeding_ lawn mowing_ carpentry_ digging) | 0 | 0.1264(0.0738) | 1,7135 | 8,66E-02 | 7,50E-01 | 0.0341(0.002) | 1.0063(0.0071) | -0.0126(0.0048) |
| Blood clot_ DVT_ bronchitis_ emphysema_ asthma_ rhinitis_ eczema_ allergy diagnosed by doctor: Hayfever_ allergic rhinitis or eczema | 0 | -0.1055(0.0616) | -1,7125 | 8,68E-02 | 7,50E-01 | 0.0695(0.0051) | 1.048(0.0155) | -0.002(0.005) |
| Celiac disease | 20190752 | -0.2391(0.1408) | -1,698 | 8,95E-02 | 7,50E-01 | 0.301(0.0502) | 1.0649(0.0106) | 0.0063(0.0072) |
| Blood clot_ DVT_ bronchitis_ emphysema_ asthma_ rhinitis_ eczema_ allergy diagnosed by doctor: Asthma | 0 | -0.1092(0.0645) | -1,6936 | 9,03E-02 | 7,50E-01 | 0.0554(0.0054) | 1.0299(0.0143) | 0.0045(0.0056) |
| Forced expiratory volume in 1 second (FEV1) | 21946350 | 0.2209(0.1305) | 1,6928 | 9,05E-02 | 7,50E-01 | 0.1654(0.0212) | 0.8839(0.0112) | -0.0096(0.008) |
| Snoring | 0 | 0.1134(0.0671) | 1,6896 | 9,11E-02 | 7,50E-01 | 0.0595(0.0027) | 1.0243(0.009) | -0.0014(0.0049) |
| Alcohol drinker status: Never | 0 | -0.1812(0.1074) | -1,6873 | 9,15E-02 | 7,50E-01 | 0.0143(0.0016) | 1.0165(0.0067) | 0.0089(0.0045) |
| Impedance of whole body | 0 | 0.0785(0.0465) | 1,6864 | 9,17E-02 | 7,50E-01 | 0.256(0.0097) | 1.0905(0.0227) | -0.001(0.0065) |
| Systolic blood pressure_ automated reading | 0 | -0.0838(0.0498) | -1,6841 | 9,22E-02 | 7,50E-01 | 0.1257(0.0056) | 1.0668(0.0151) | 0.0074(0.0052) |
| Non-cancer illness code_ self-reported: cholelithiasis/gall stones | 0 | 0.2449(0.1455) | 1,6831 | 9,23E-02 | 7,50E-01 | 0.0082(0.0017) | 1.0216(0.0076) | -0.0014(0.0048) |
| Illness_ injury_ bereavement_ stress in last 2 years: Financial difficulties | 0 | -0.1412(0.0841) | -1,6795 | 9,31E-02 | 7,50E-01 | 0.033(0.0021) | 1.0279(0.008) | 0.0036(0.0052) |
| HbA1C | 20858683 | -0.2245(0.1343) | -1,6713 | 9,47E-02 | 7,50E-01 | 0.0633(0.0112) | 0.9998(0.0068) | 0.0024(0.0041) |
| Arm fat mass (right) | 0 | -0.0793(0.0475) | -1,6691 | 9,51E-02 | 7,50E-01 | 0.2308(0.0079) | 1.0701(0.0173) | 0.0039(0.0068) |
| Ever used hormone-replacement therapy (HRT) | 0 | 0.1539(0.0924) | 1,6662 | 9,57E-02 | 7,50E-01 | 0.0402(0.0032) | 0.9992(0.0072) | -0.0039(0.0047) |
| Current employment status: Unable to work because of sickness or disability | 0 | -0.1637(0.0985) | -1,6624 | 9,64E-02 | 7,50E-01 | 0.0245(0.0019) | 1.0174(0.0075) | 0.0031(0.0049) |
| Smoking status: Previous | 0 | -0.1195(0.0719) | -1,6611 | 9,67E-02 | 7,50E-01 | 0.051(0.0028) | 1.022(0.0092) | 0.0024(0.0048) |
| Body fat | 26833246 | -0.135(0.0814) | -1,6593 | 9,71E-02 | 7,50E-01 | 0.1046(0.0083) | 0.9054(0.0064) | 0.0024(0.0049) |
| Coronary artery disease | 26343387 | -0.122(0.0738) | -1,6543 | 9,81E-02 | 7,50E-01 | 0.0776(0.0054) | 1.0445(0.0092) | 0.0074(0.0045) |
| Non-cancer illness code_ self-reported: emphysema/chronic bronchitis | 0 | -0.2143(0.1301) | -1,6472 | 9,95E-02 | 7,50E-01 | 0.0097(0.0016) | 0.9873(0.0068) | 0.0024(0.0046) |
| Underlying (primary) cause of death: ICD10: J84.1 Other interstitial pulmonary diseases with fibrosis | 0 | 0.5398(0.3286) | 1,643 | 1,00E-01 | 7,50E-01 | 0.0944(0.0644) | 0.9945(0.0069) | -0.0036(0.0048) |
| Non-cancer illness code_ self-reported: back problem | 0 | -0.3007(0.184) | -1,6336 | 1,02E-01 | 7,57E-01 | 0.0044(0.0013) | 1.0079(0.0062) | 0.004(0.0042) |
| Worrier / anxious feelings | 0 | 0.0993(0.0609) | 1,6311 | 1,03E-01 | 7,57E-01 | 0.0786(0.0049) | 1.0021(0.0122) | -0.0028(0.0054) |
| Eczema | 26482879 | -0.2105(0.1302) | -1,6167 | 1,06E-01 | 7,64E-01 | 0.0666(0.0148) | 1.0236(0.0072) | -0.0065(0.0044) |
| Whole body fat mass | 0 | -0.0773(0.0479) | -1,6146 | 1,06E-01 | 7,64E-01 | 0.2345(0.0077) | 1.073(0.0178) | 0.0048(0.0068) |
| Forced expiratory volume in 1-second (FEV1)_ predicted percentage | 0 | 0.1175(0.0731) | 1,6064 | 1,08E-01 | 7,64E-01 | 0.1975(0.0097) | 1.0369(0.0097) | -0.0034(0.0058) |
| Maternal smoking around birth | 0 | -0.1296(0.0807) | -1,6056 | 1,08E-01 | 7,64E-01 | 0.0498(0.0028) | 1.0353(0.0092) | 0.0039(0.0052) |
| Frequency of travelling from home to job workplace | 0 | 0.2058(0.1291) | 1,5943 | 1,11E-01 | 7,70E-01 | 0.0218(0.0028) | 1.0169(0.007) | -0.0064(0.005) |
| Impedance of leg (right) | 0 | 0.0765(0.0482) | 1,5862 | 1,13E-01 | 7,70E-01 | 0.2386(0.0093) | 1.0919(0.0222) | -0.002(0.0066) |
| Townsend deprivation index at recruitment | 0 | -0.1397(0.0881) | -1,5845 | 1,13E-01 | 7,70E-01 | 0.0334(0.002) | 1.0355(0.0072) | 0.0052(0.0054) |
| Primary sclerosing cholangitis | 27992413 | 0.1969(0.1246) | 1,5805 | 1,14E-01 | 7,70E-01 | 0.3891(0.1618) | 0.9948(0.0161) | -0.0012(0.0052) |
| Relative age voice broke | 0 | 0.1337(0.0847) | 1,5781 | 1,15E-01 | 7,70E-01 | 0.0719(0.0052) | 1.0113(0.0084) | -0.0031(0.0046) |
| Illnesses of mother: Chronic bronchitis/emphysema | 0 | -0.1856(0.1179) | -1,5743 | 1,15E-01 | 7,70E-01 | 0.0126(0.0018) | 1.0035(0.0067) | 0.0053(0.0046) |
| Valine | 27005778 | 0.3044(0.1944) | 1,5657 | 1,17E-01 | 7,77E-01 | 0.0529(0.0196) | 1.0177(0.0073) | -0.0039(0.0044) |
| Chronic Kidney Disease | 26831199 | 0.2615(0.1681) | 1,5563 | 1,20E-01 | 7,80E-01 | 0.0175(0.0056) | 1.0186(0.0087) | -0.004(0.0052) |
| Non-cancer illness code_ self-reported: asthma | 0 | -0.1006(0.0648) | -1,552 | 1,21E-01 | 7,80E-01 | 0.0561(0.0055) | 1.0305(0.0143) | 0.0052(0.0057) |
| Pain type(s) experienced in last month: Pain all over the body | 0 | -0.2077(0.1348) | -1,5405 | 1,23E-01 | 7,80E-01 | 0.0099(0.0016) | 1.0049(0.0072) | 0.0043(0.0049) |
| Non-cancer illness code_ self-reported: migraine | 0 | -0.1534(0.0998) | -1,5361 | 1,25E-01 | 7,80E-01 | 0.0166(0.0018) | 1.0119(0.0073) | 0.0051(0.0046) |
| Diagnoses - main ICD10: N92 Excessive_ frequent and irregular menstruation | 0 | -0.2309(0.1504) | -1,5352 | 1,25E-01 | 7,80E-01 | 0.0073(0.0017) | 1.0002(0.0072) | 0.0022(0.0047) |
| Pain type(s) experienced in last month: Hip pain | 0 | -0.1484(0.0967) | -1,5348 | 1,25E-01 | 7,80E-01 | 0.0235(0.0019) | 1.0178(0.0074) | 0.0085(0.0049) |
| Had menopause | 0 | 0.2266(0.1482) | 1,5295 | 1,26E-01 | 7,80E-01 | 0.0198(0.0038) | 1.0034(0.0066) | -0.0044(0.0047) |
| Forced expiratory volume in 1 second (FEV1)/Forced Vital capacity(FVC) | 26635082 | 0.1769(0.1158) | 1,5276 | 1,27E-01 | 7,80E-01 | 0.1117(0.0156) | 0.9817(0.0071) | -0.0044(0.0049) |
| Bipolar disorder | 21926972 | 0.1306(0.0857) | 1,5242 | 1,28E-01 | 7,80E-01 | 0.4338(0.0374) | 1.0221(0.0073) | 0.0012(0.0047) |
| Hip circumference | 0 | -0.077(0.0506) | -1,5237 | 1,28E-01 | 7,80E-01 | 0.2171(0.0082) | 1.0792(0.0191) | 0.0026(0.0069) |
| Number of pregnancy terminations | 0 | 0.2061(0.1367) | 1,5073 | 1,32E-01 | 7,84E-01 | 0.061(0.0102) | 1.007(0.0076) | -0.0038(0.005) |
| Work/job satisfaction | 0 | 0.1731(0.1151) | 1,5037 | 1,33E-01 | 7,84E-01 | 0.0496(0.0068) | 0.9934(0.0068) | -0.0077(0.0045) |
| Non-cancer illness code_ self-reported: high cholesterol | 0 | -0.1283(0.0853) | -1,5027 | 1,33E-01 | 7,84E-01 | 0.04(0.0055) | 1.0566(0.0275) | 0.0065(0.0057) |
| Time spent driving | 0 | -0.1341(0.0893) | -1,5012 | 1,33E-01 | 7,84E-01 | 0.0422(0.0028) | 1.0014(0.007) | 0.0072(0.005) |
| Diagnoses - main ICD10: R10 Abdominal and pelvic pain | 0 | -0.1775(0.1183) | -1,5002 | 1,34E-01 | 7,84E-01 | 0.012(0.0017) | 0.9965(0.0072) | 0.0026(0.0047) |
| Tyrosine | 27005778 | 0.2578(0.1733) | 1,4876 | 1,37E-01 | 7,84E-01 | 0.0768(0.0283) | 0.9991(0.0065) | -0.0008(0.0043) |
| Leptin_adjBMI | 26833098 | -0.1986(0.1336) | -1,486 | 1,37E-01 | 7,84E-01 | 0.0912(0.0179) | 1.0048(0.007) | 0.0111(0.0044) |
| Mouth/teeth dental problems: Mouth ulcers | 0 | 0.1214(0.0817) | 1,4856 | 1,37E-01 | 7,84E-01 | 0.0285(0.0037) | 1.0324(0.0109) | -0.0079(0.0048) |
| Diagnoses - main ICD10: M16 Coxarthrosis [arthrosis of hip] | 0 | -0.173(0.1182) | -1,4642 | 1,43E-01 | 7,92E-01 | 0.0103(0.0018) | 1.0106(0.008) | 0.003(0.0045) |
| Diagnoses - main ICD10: S09 Other and unspecified injuries of head | 0 | -0.2974(0.2031) | -1,4644 | 1,43E-01 | 7,92E-01 | 0.0039(0.0016) | 0.9943(0.0066) | 0.0062(0.0044) |
| Obesity class 2 | 23563607 | -0.1204(0.0825) | -1,4583 | 1,45E-01 | 7,92E-01 | 0.1817(0.0123) | 1.004(0.0099) | 0.0019(0.006) |
| Extreme height | 23563607 | -0.1252(0.0864) | -1,4497 | 1,47E-01 | 7,92E-01 | 1.2326(0.1063) | 1.0353(0.0177) | 0.0192(0.0079) |
| Diagnoses - main ICD10: J33 Nasal polyp | 0 | -0.392(0.2709) | -1,4472 | 1,48E-01 | 7,92E-01 | 0.0029(0.0015) | 1.0078(0.0073) | 0.0074(0.0049) |
| Medication for cholesterol_ blood pressure or diabetes: Cholesterol lowering medication | 0 | -0.1209(0.0837) | -1,4453 | 1,48E-01 | 7,92E-01 | 0.0675(0.0065) | 1.0329(0.0142) | 0.0037(0.005) |

| | | | | | | | | |
|---|---|---|---|---|---|---|---|---|
| Ferritin | 25352340 | 0.2376(0.165) | 1,4404 | 1,50E-01 | 7,92E-01 | 0.0926(0.0269) | 1.0276(0.0095) | -0.0084(0.0059) |
| Transport type for commuting to job workplace: Walk | 0 | 0.1977(0.1373) | 1,4399 | 1,50E-01 | 7,92E-01 | 0.017(0.0027) | 1.0023(0.0063) | -0.0042(0.0047) |
| Trunk fat mass | 0 | -0.0695(0.0483) | -1,4389 | 1,50E-01 | 7,92E-01 | 0.2313(0.0076) | 1.0774(0.0183) | 0.0034(0.0068) |
| Light smokers_ at least 100 smokes in lifetime | 0 | -0.1293(0.09) | -1,4373 | 1,51E-01 | 7,92E-01 | 0.0762(0.0074) | 1.0057(0.0072) | 0.0031(0.0045) |
| Wheeze or whistling in the chest in last year | 0 | -0.1093(0.0761) | -1,4362 | 1,51E-01 | 7,92E-01 | 0.0594(0.003) | 1.0326(0.0096) | 0.0018(0.0061) |
| Target heart rate achieved | 0 | 0.2535(0.1768) | 1,4339 | 1,52E-01 | 7,92E-01 | 0.0341(0.0111) | 1.0062(0.0071) | -0.0032(0.0044) |
| Risk taking | 0 | -0.0997(0.0696) | -1,4329 | 1,52E-01 | 7,92E-01 | 0.0555(0.0029) | 1.0056(0.0093) | 0.0048(0.0054) |
| Diagnoses - main ICD10: H25 Senile cataract | 0 | -0.3544(0.249) | -1,4236 | 1,55E-01 | 7,96E-01 | 0.0031(0.0014) | 1.014(0.007) | 0.005(0.0048) |
| Diagnoses - main ICD10: I25 Chronic ischaemic heart disease | 0 | -0.14(0.0985) | -1,4208 | 1,55E-01 | 7,96E-01 | 0.0192(0.0021) | 1.0218(0.0075) | 0.0014(0.0052) |
| Fractured/broken bones in last 5 years | 0 | -0.1449(0.1023) | -1,4161 | 1,57E-01 | 7,96E-01 | 0.0185(0.0021) | 1.0083(0.0074) | 0.0000048619(0.0049) |
| Illnesses of father: Chronic bronchitis/emphysema | 0 | -0.1823(0.1299) | -1,4029 | 1,61E-01 | 7,96E-01 | 0.0143(0.0019) | 1.0249(0.0068) | 0.0081(0.0047) |
| Illnesses of mother: Breast cancer | 0 | -0.2256(0.161) | -1,4013 | 1,61E-01 | 7,96E-01 | 0.0094(0.0019) | 1.0062(0.0081) | 0.0005(0.0052) |
| Multiple sclerosis | 21833088 | 0.3303(0.2361) | 1,3991 | 1,62E-01 | 7,96E-01 | 0.0532(0.0278) | 1.0586(0.0084) | -0.009(0.0053) |
| Frequency of other exercises in last 4 weeks | 0 | 0.1982(0.1417) | 1,3991 | 1,62E-01 | 7,96E-01 | 0.0216(0.0033) | 0.9978(0.0063) | -0.0034(0.005) |
| Diagnoses - main ICD10: I80 Phlebitis and thrombophlebitis | 0 | -0.1937(0.1389) | -1,3941 | 1,63E-01 | 7,96E-01 | 0.0082(0.0017) | 0.9932(0.0083) | 0.0018(0.0044) |
| Frequency of walking for pleasure in last 4 weeks | 0 | -0.1219(0.0876) | -1,3915 | 1,64E-01 | 7,96E-01 | 0.0349(0.0026) | 1.006(0.0076) | 0.005(0.0044) |
| Mean Pallidum | 25607358 | 0.2323(0.1678) | 1,3842 | 1,66E-01 | 7,96E-01 | 0.1645(0.0412) | 0.9774(0.0069) | -0.0095(0.0048) |
| Non-cancer illness code_ self-reported: allergy or anaphylactic reaction to drug | 0 | 1.0492(0.7608) | 1,3791 | 1,68E-01 | 7,96E-01 | 0.0011(0.0013) | 1.0074(0.0062) | -0.0121(0.0044) |
| Citrate | 27005778 | 0.2326(0.1699) | 1,3691 | 1,71E-01 | 7,96E-01 | 0.0693(0.0222) | 1.0126(0.0076) | -0.0014(0.0044) |
| Impedance of leg (left) | 0 | 0.0654(0.0478) | 1,3662 | 1,72E-01 | 7,96E-01 | 0.2394(0.0095) | 1.0822(0.0223) | 0.0013(0.0067) |
| Non-cancer illness code_ self-reported: hypertrophic cardiomyopathy (hcm / hocm) | 0 | -0.4742(0.3474) | -1,3649 | 1,72E-01 | 7,96E-01 | 0.0016(0.0013) | 1.0017(0.0066) | 0.0066(0.0044) |
| Number of self-reported non-cancer illnesses | 0 | -0.0843(0.0619) | -1,3614 | 1,73E-01 | 7,96E-01 | 0.0608(0.0025) | 1.0275(0.01) | 0.0013(0.0047) |
| Heel bone mineral density (BMD) T-score_ automated (left) | 0 | 0.0736(0.0541) | 1,3611 | 1,74E-01 | 7,96E-01 | 0.2964(0.035) | 1.0345(0.0208) | -0.0018(0.0053) |
| Diagnoses - main ICD10: Z47 Other orthopaedic follow-up care | 0 | -0.2955(0.2174) | -1,3594 | 1,74E-01 | 7,96E-01 | 0.0036(0.0013) | 0.9968(0.0064) | 0.0053(0.0049) |
| Medication for pain relief_ constipation_ heartburn: Aspirin | 0 | -0.1282(0.0944) | -1,3578 | 1,75E-01 | 7,96E-01 | 0.0209(0.0018) | 1.0138(0.007) | 0.0024(0.0051) |
| Qualifications: A levels/AS levels or equivalent | 0 | 0.0835(0.0616) | 1,3556 | 1,75E-01 | 7,96E-01 | 0.0964(0.0036) | 1.0528(0.0119) | -0.0089(0.0057) |
| Non-cancer illness code_ self-reported: mania/bipolar disorder/manic depression | 0 | -0.2385(0.1779) | -1,3406 | 1,80E-01 | 7,96E-01 | 0.0057(0.0014) | 1.0005(0.0061) | 0.0073(0.005) |
| Exposure to tobacco smoke at home | 0 | 0.1798(0.1343) | 1,3392 | 1,81E-01 | 7,96E-01 | 0.0114(0.0016) | 0.9995(0.0062) | -0.0004(0.0051) |
| Exposure to tobacco smoke outside home | 0 | -0.1305(0.0977) | -1,3359 | 1,82E-01 | 7,96E-01 | 0.0317(0.0025) | 1.0205(0.0084) | 0.0005(0.0055) |
| Age at last live birth | 0 | 0.1039(0.0778) | 1,3356 | 1,82E-01 | 7,96E-01 | 0.0876(0.0059) | 1.0178(0.0081) | -0.002(0.005) |
| Non-cancer illness code_ self-reported: nasal polyps | 0 | -0.2476(0.1856) | -1,3339 | 1,82E-01 | 7,96E-01 | 0.0048(0.0015) | 1.0073(0.0062) | 0.0015(0.0047) |
| Average weekly champagne plus white wine intake | 0 | 0.1235(0.0927) | 1,3327 | 1,83E-01 | 7,96E-01 | 0.0299(0.0026) | 1.0421(0.0077) | 0.0032(0.0045) |
| Friendships satisfaction | 0 | 0.1368(0.1029) | 1,3303 | 1,83E-01 | 7,96E-01 | 0.062(0.0051) | 0.9987(0.007) | 0.001(0.0051) |
| Serum creatinine | 26831199 | -0.099(0.0744) | -1,3301 | 1,84E-01 | 7,96E-01 | 0.1077(0.0097) | 0.9749(0.0133) | 0.0059(0.0053) |
| Infant head circumference | 22504419 | 0.1951(0.147) | 1,3271 | 1,85E-01 | 7,96E-01 | 0.2267(0.0444) | 0.9908(0.0068) | -0.0045(0.0046) |
| Illnesses of mother: None of the above (group 2) | 0 | 0.1966(0.1483) | 1,3257 | 1,85E-01 | 7,96E-01 | 0.0092(0.0017) | 1.0002(0.0065) | -0.0079(0.005) |
| Alcohol intake frequency. | 0 | -0.085(0.0641) | -1,3254 | 1,85E-01 | 7,96E-01 | 0.0822(0.004) | 1.0474(0.0107) | 0.007(0.0058) |
| Non-cancer illness code_ self-reported: sleep apnoea | 0 | -0.2948(0.2229) | -1,3228 | 1,86E-01 | 7,96E-01 | 0.0034(0.0014) | 1.0029(0.0068) | 0.0034(0.0048) |
| PGC cross-disorder analysis | 23453885 | 0.1127(0.0854) | 1,3192 | 1,87E-01 | 7,96E-01 | 0.17(0.0131) | 1.0141(0.0114) | -0.0033(0.0061) |
| Qualifications: None of the above | 0 | -0.0724(0.055) | -1,3172 | 1,88E-01 | 7,96E-01 | 0.0965(0.0039) | 1.0682(0.0114) | 0.0077(0.0053) |
| Used an inhaler for chest within last hour | 0 | -0.289(0.2219) | -1,3024 | 1,93E-01 | 7,96E-01 | 0.0044(0.0015) | 1.0017(0.0062) | 0.0028(0.005) |
| Worry too long after embarrassment | 0 | 0.0839(0.0648) | 1,2963 | 1,95E-01 | 7,96E-01 | 0.0643(0.0035) | 1.0187(0.0102) | -0.0012(0.0052) |
| Waist-to-hip ratio | 25673412 | 0.086(0.0664) | 1,2941 | 1,96E-01 | 7,96E-01 | 0.1116(0.0071) | 0.9217(0.009) | -0.009(0.0048) |
| Parents age at death | 27015805 | 0.2241(0.1731) | 1,2942 | 1,96E-01 | 7,96E-01 | 0.029(0.0068) | 1.0159(0.0062) | -0.0026(0.0046) |
| Illnesses of siblings: Severe depression | 0 | -0.1702(0.1318) | -1,2911 | 1,97E-01 | 7,96E-01 | 0.0139(0.0018) | 1.0049(0.0067) | 0.0068(0.0047) |
| Bilateral oophorectomy (both ovaries removed) | 0 | -0.1651(0.1281) | -1,288 | 1,98E-01 | 7,96E-01 | 0.0206(0.0033) | 1.0121(0.0073) | -0.0004(0.0048) |
| Average number of double bonds in a fatty acid chain | 27005778 | 0.1925(0.1495) | 1,2876 | 1,98E-01 | 7,96E-01 | 0.1662(0.0544) | 1.0016(0.0074) | -0.0043(0.0043) |
| Lung adenocarcinoma | 27488534 | -0.2249(0.1747) | -1,2871 | 1,98E-01 | 7,96E-01 | 0.0313(0.0113) | 1.0189(0.0064) | 0.0042(0.0046) |
| Pain type(s) experienced in last month: Stomach or abdominal pain | 0 | 0.1224(0.0952) | 1,2861 | 1,98E-01 | 7,96E-01 | 0.0216(0.0017) | 1.0059(0.007) | -0.0052(0.0049) |
| Frequency of tiredness / lethargy in last 2 weeks | 0 | -0.0882(0.069) | -1,2783 | 2,01E-01 | 7,96E-01 | 0.0596(0.0029) | 1.0259(0.0093) | 0.0028(0.0055) |
| Breastfed as a baby | 0 | -0.1199(0.0939) | -1,2764 | 2,02E-01 | 7,96E-01 | 0.0262(0.0023) | 1.0206(0.0074) | 0.0037(0.0044) |
| Blood clot_ DVT_ bronchitis_ emphysema_ asthma_ rhinitis_ eczema_ allergy diagnosed by doctor: Emphysema/chronic bronchitis | 0 | -0.1529(0.1198) | -1,2761 | 2,02E-01 | 7,96E-01 | 0.0122(0.0016) | 0.9861(0.0071) | 0.0012(0.0047) |
| Non-cancer illness code_ self-reported: angina | 0 | -0.1234(0.0968) | -1,2748 | 2,02E-01 | 7,96E-01 | 0.0219(0.0019) | 1.0236(0.0073) | -0.0005(0.0054) |
| Knee pain for 3+ months | 0 | 0.2114(0.1661) | 1,2732 | 2,03E-01 | 7,96E-01 | 0.0259(0.0068) | 1.0009(0.0066) | -0.0045(0.0047) |
| Vascular/heart problems diagnosed by doctor: None of the above | 0 | 0.0688(0.0542) | 1,2702 | 2,04E-01 | 7,96E-01 | 0.1121(0.0046) | 1.0596(0.0152) | 0.0036(0.0057) |
| Arm fat percentage (left) | 0 | -0.06(0.0473) | -1,268 | 2,05E-01 | 7,96E-01 | 0.2189(0.0069) | 1.0733(0.0175) | 0.005(0.0065) |
| Difference in height between adolescence and adulthood; age 14 | 23449627 | 0.1874(0.1481) | 1,2657 | 2,06E-01 | 7,96E-01 | 0.4431(0.107) | 0.9814(0.0065) | -0.007(0.0043) |
| Diagnoses - main ICD10: R14 Flatulence and related conditions | 0 | -0.2969(0.236) | -1,258 | 2,08E-01 | 8,00E-01 | 0.0031(0.0013) | 0.993(0.0067) | 0.002(0.0049) |
| Non-cancer illness code_ self-reported: hyperthyroidism/thyrotoxicosis | 0 | 0.1817(0.1452) | 1,2511 | 2,11E-01 | 8,06E-01 | 0.0087(0.0026) | 1.0047(0.008) | -0.0082(0.0049) |
| Diagnoses - main ICD10: M24 Other specific joint derangements | 0 | -0.2841(0.2277) | -1,2477 | 2,12E-01 | 8,06E-01 | 0.0035(0.0013) | 0.9879(0.0061) | 0.0064(0.0045) |
| Age of smoking initiation | 20418890 | 0.2688(0.2166) | 1,2408 | 2,15E-01 | 8,06E-01 | 0.0614(0.0192) | 0.9984(0.0064) | -0.0003(0.005) |
| Forced expiratory volume in 1 second (FEV1) | 26635082 | 0.135(0.1089) | 1,2396 | 2,15E-01 | 8,06E-01 | 0.1389(0.0155) | 0.9926(0.0066) | -0.0007(0.0049) |
| Creatinine (enzymatic) in urine | 0 | -0.0741(0.0598) | -1,2394 | 2,15E-01 | 8,06E-01 | 0.0655(0.0029) | 1.0198(0.0087) | 0.0039(0.0051) |
| Diagnoses - main ICD10: K50 Crohns disease [regional enteritis] | 0 | -0.2323(0.1886) | -1,2319 | 2,18E-01 | 8,13E-01 | 0.0043(0.0013) | 1.0029(0.0064) | 0.000023522(0.0042) |
| Frequency of heavy DIY in last 4 weeks | 0 | -0.1672(0.1361) | -1,2292 | 2,19E-01 | 8,13E-01 | 0.0206(0.0034) | 1.0093(0.006) | 0.0025(0.004) |
| Number of days/week of moderate physical activity 10+ minutes | 0 | 0.0928(0.0761) | 1,2197 | 2,23E-01 | 8,23E-01 | 0.0417(0.0026) | 1.0128(0.0081) | 0.0043(0.005) |
| Sodium in urine | 0 | -0.0734(0.0605) | -1,2141 | 2,25E-01 | 8,23E-01 | 0.0708(0.0031) | 1.0306(0.0101) | 0.0033(0.0052) |
| Diagnoses - main ICD10: K22 Other diseases of oesophagus | 0 | -0.2931(0.2416) | -1,213 | 2,25E-01 | 8,23E-01 | 0.0037(0.0015) | 1.0055(0.0065) | 0.0061(0.0047) |
| Fasting glucose main effect | 22581228 | -0.1338(0.1109) | -1,2066 | 2,28E-01 | 8,26E-01 | 0.0828(0.0122) | 1.0051(0.009) | 0.0022(0.0042) |
| Vascular/heart problems diagnosed by doctor: Angina | 0 | -0.1196(0.0992) | -1,2055 | 2,28E-01 | 8,26E-01 | 0.0214(0.0019) | 1.0265(0.0073) | -0.0015(0.0056) |

| | | | | | | | | |
|---|---|---|---|---|---|---|---|---|
| Number of full sisters | 0 | -0.1163(0.0971) | -1,1976 | 2,31E-01 | 8,33E-01 | 0.0184(0.0018) | 1.0406(0.0066) | -0.0002(0.0046) |
| Diagnoses - main ICD10: R07 Pain in throat and chest | 0 | -0.1193(0.1003) | -1,19 | 2,34E-01 | 8,33E-01 | 0.0163(0.0018) | 1.0044(0.0075) | 0.0033(0.0051) |
| Obesity class 3 | 23563607 | -0.1371(0.1158) | -1,1833 | 2,37E-01 | 8,33E-01 | 0.1189(0.0146) | 0.9805(0.0092) | 0.0001(0.0054) |
| Cancer code_ self-reported: basal cell carcinoma | 0 | -0.1506(0.1275) | -1,1816 | 2,37E-01 | 8,33E-01 | 0.01(0.0018) | 0.9982(0.0076) | 0.0033(0.0048) |
| Medication for cholesterol_ blood pressure_ diabetes_ or take exogenous hormones: Hormone replacement therapy | 0 | 0.1609(0.1364) | 1,1798 | 2,38E-01 | 8,33E-01 | 0.0154(0.0027) | 1.0013(0.0061) | -0.0039(0.0047) |
| Mineral and other dietary supplements: Selenium | 0 | -0.1988(0.1685) | -1,1796 | 2,38E-01 | 8,33E-01 | 0.0059(0.0015) | 1.0016(0.007) | 0.0108(0.0046) |
| Height; Females at age 10 and males at age 12 | 23449627 | -0.1408(0.1196) | -1,1771 | 2,39E-01 | 8,33E-01 | 0.4353(0.0474) | 0.9521(0.0079) | 0.0093(0.0053) |
| Duration of strenuous sports | 0 | 0.2365(0.2011) | 1,1762 | 2,40E-01 | 8,33E-01 | 0.0303(0.0132) | 0.9941(0.0067) | -0.007(0.0045) |
| Reproduciblity of spirometry measurement using ERS/ATS criteria | 0 | -0.1513(0.1288) | -1,1741 | 2,40E-01 | 8,33E-01 | 0.014(0.0023) | 1.0081(0.0072) | 0.0035(0.0047) |
| Types of transport used (excluding work): Public transport | 0 | 0.1093(0.0944) | 1,1577 | 2,47E-01 | 8,45E-01 | 0.0224(0.0018) | 1.0255(0.0065) | -0.0041(0.005) |
| Home area population density - urban or rural: Scotland - Large Urban Area | 0 | -0.1798(0.1553) | -1,1574 | 2,47E-01 | 8,45E-01 | 0.0108(0.0031) | 2.1365(0.0147) | 0.002(0.0068) |
| Difference in height between childhood and adulthood; age 8 | 23449627 | 0.1375(0.119) | 1,1556 | 2,48E-01 | 8,45E-01 | 0.3291(0.0524) | 0.9721(0.0078) | -0.0081(0.0045) |
| Current tobacco smoking | 0 | -0.0833(0.0721) | -1,1557 | 2,48E-01 | 8,45E-01 | 0.0545(0.0029) | 1.0143(0.0093) | -0.0052(0.0054) |
| Blood clot_ DVT_ bronchitis_ emphysema_ asthma_ rhinitis_ eczema_ allergy diagnosed by doctor: Blood clot in the lung | 0 | -0.1904(0.1655) | -1,1504 | 2,50E-01 | 8,48E-01 | 0.0056(0.0016) | 1.0084(0.0076) | 0.005(0.0043) |
| Neuroticism score | 0 | 0.0671(0.0585) | 1,1479 | 2,51E-01 | 8,48E-01 | 0.1167(0.0062) | 1.0198(0.0134) | -0.0038(0.0055) |
| Non-cancer illness code_ self-reported: pulmonary embolism +/- dvt | 0 | -0.1928(0.1689) | -1,1413 | 2,54E-01 | 8,54E-01 | 0.0053(0.0016) | 1.0076(0.0076) | 0.0054(0.0044) |
| Hearing difficulty/problems: Yes | 0 | 0.0954(0.0839) | 1,137 | 2,56E-01 | 8,57E-01 | 0.0411(0.0023) | 1.0158(0.0089) | -0.0106(0.0057) |
| Diagnoses - main ICD10: K80 Cholelithiasis | 0 | 0.1502(0.1327) | 1,1318 | 2,58E-01 | 8,60E-01 | 0.0093(0.0019) | 1.0243(0.0086) | -0.0009(0.0048) |
| Types of transport used (excluding work): Cycle | 0 | 0.0933(0.0829) | 1,1254 | 2,60E-01 | 8,63E-01 | 0.0242(0.0018) | 1.027(0.0069) | -0.0027(0.0043) |
| Non-cancer illness code_ self-reported: depression | 0 | 0.1193(0.1063) | 1,1222 | 2,62E-01 | 8,66E-01 | 0.0183(0.0017) | 0.9999(0.0071) | -0.0061(0.0046) |
| Ever had bowel cancer screening | 0 | -0.1228(0.1102) | -1,1144 | 2,65E-01 | 8,72E-01 | 0.0128(0.0017) | 1.0206(0.0071) | 0.0087(0.0046) |
| Had major operations | 0 | -0.1305(0.1178) | -1,1078 | 2,68E-01 | 8,77E-01 | 0.0246(0.0035) | 1.006(0.0072) | 0.0044(0.0046) |
| Glycoprotein acetyls; mainly a1-acid glycoprotein | 27005778 | 0.1849(0.1672) | 1,1056 | 2,69E-01 | 8,77E-01 | 0.1(0.0281) | 0.9856(0.0063) | -0.0047(0.0043) |
| Urinary albumin-to-creatinine ratio | 26631737 | 0.1616(0.1471) | 1,0982 | 2,72E-01 | 8,78E-01 | 0.044(0.0086) | 0.9971(0.0067) | -0.0071(0.0049) |
| Diagnoses - main ICD10: K43 Ventral hernia | 0 | 0.1764(0.1606) | 1,0979 | 2,72E-01 | 8,78E-01 | 0.0056(0.0014) | 0.9921(0.0065) | -0.0022(0.0047) |
| Non-cancer illness code_ self-reported: gastro-oesophageal reflux (gord) / gastric reflux | 0 | 0.1369(0.125) | 1,0951 | 2,74E-01 | 8,78E-01 | 0.0113(0.0016) | 1.015(0.0071) | 0.0022(0.0048) |
| Diagnoses - main ICD10: G56 Mononeuropathies of upper limb | 0 | -0.1268(0.1161) | -1,0924 | 2,75E-01 | 8,78E-01 | 0.0135(0.0018) | 1.0098(0.0072) | -0.0028(0.0045) |
| Forced expiratory volume in 1-second (FEV1)_ Best measure | 0 | 0.0715(0.066) | 1,083 | 2,79E-01 | 8,78E-01 | 0.1608(0.0075) | 1.0596(0.0136) | -0.005(0.0063) |
| Diagnoses - main ICD10: J44 Other chronic obstructive pulmonary disease | 0 | 0.1941(0.1795) | 1,081 | 2,80E-01 | 8,78E-01 | 0.0045(0.0014) | 0.9962(0.0066) | -0.0044(0.0042) |
| Had other major operations | 0 | -0.1205(0.1118) | -1,0785 | 2,81E-01 | 8,78E-01 | 0.0302(0.0032) | 1.0103(0.0069) | 0.0047(0.005) |
| Ever depressed for a whole week | 0 | -0.1045(0.0971) | -1,0762 | 2,82E-01 | 8,78E-01 | 0.0619(0.0053) | 0.9942(0.0069) | 0.0097(0.005) |
| Ulcerative colitis | 26192919 | 0.106(0.0986) | 1,075 | 2,82E-01 | 8,78E-01 | 0.2627(0.0318) | 1.0438(0.0096) | -0.0043(0.0051) |
| Ever taken oral contraceptive pill | 0 | -0.1312(0.1224) | -1,0723 | 2,84E-01 | 8,78E-01 | 0.0195(0.0029) | 1.0107(0.007) | 0.0052(0.0048) |
| Diagnoses - main ICD10: K44 Diaphragmatic hernia | 0 | -0.1767(0.1651) | -1,07 | 2,85E-01 | 8,78E-01 | 0.0052(0.0014) | 1.0057(0.0066) | -0.0005(0.0046) |
| Forced expiratory volume in 1-second (FEV1) | 0 | 0.0694(0.0649) | 1,0694 | 2,85E-01 | 8,78E-01 | 0.1456(0.0064) | 1.0682(0.0137) | -0.0072(0.0065) |
| Arm fat percentage (right) | 0 | -0.0518(0.0486) | -1,0665 | 2,86E-01 | 8,78E-01 | 0.2154(0.0068) | 1.0781(0.0173) | 0.0043(0.0066) |
| Attention deficit hyperactivity disorder | 20732625 | -0.2204(0.2069) | -1,0651 | 2,87E-01 | 8,78E-01 | 0.2326(0.091) | 1.0056(0.0061) | 0.0045(0.0047) |
| Duration of walks | 0 | -0.0848(0.0797) | -1,0651 | 2,87E-01 | 8,78E-01 | 0.047(0.0028) | 1.0056(0.0083) | -0.0015(0.0046) |
| Omega-3 fatty acids | 27005778 | -0.1534(0.1443) | -1,0628 | 2,88E-01 | 8,78E-01 | 0.1586(0.0423) | 0.9897(0.0076) | 0.0005(0.0044) |
| Diagnoses - main ICD10: S66 Injury of muscle and tendon at wrist and hand level | 0 | -0.2902(0.2748) | -1,056 | 2,91E-01 | 8,80E-01 | 0.0023(0.0012) | 0.9997(0.0065) | 0.0072(0.005) |
| Former alcohol drinker | 0 | 0.1483(0.1413) | 1,0503 | 2,94E-01 | 8,80E-01 | 0.1225(0.0209) | 0.9969(0.0059) | -0.0081(0.0045) |
| Fasting insulin main effect | 22581228 | 0.1345(0.1284) | 1,0475 | 2,95E-01 | 8,80E-01 | 0.0654(0.0098) | 1.0174(0.007) | -0.009(0.0046) |
| Long-standing illness_ disability or infirmity | 0 | -0.0731(0.0702) | -1,0401 | 2,98E-01 | 8,80E-01 | 0.0496(0.0024) | 1.0276(0.0086) | -0.0021(0.0052) |
| Cancer code_ self-reported: prostate cancer | 0 | -0.1546(0.1488) | -1,0389 | 2,99E-01 | 8,80E-01 | 0.0065(0.0015) | 1.0126(0.0072) | -0.0003(0.0044) |
| Prospective memory result | 0 | 0.1026(0.0989) | 1,0377 | 2,99E-01 | 8,80E-01 | 0.0564(0.0051) | 1.0122(0.0077) | 0.0004(0.0047) |
| Fasting proinsulin | 20081858 | -0.2012(0.1948) | -1,0331 | 3,02E-01 | 8,80E-01 | 0.1497(0.0745) | 0.9858(0.0083) | 0.0075(0.0046) |
| Taking other prescription medications | 0 | -0.0805(0.078) | -1,0319 | 3,02E-01 | 8,80E-01 | 0.0431(0.0022) | 1.0158(0.0084) | 0.0044(0.0052) |
| Leg fat percentage (left) | 0 | -0.0512(0.0497) | -1,0294 | 3,03E-01 | 8,80E-01 | 0.2075(0.006) | 1.0848(0.0165) | 0.004(0.0065) |
| Weight change compared with 1 year ago | 0 | -0.1439(0.1399) | -1,0289 | 3,04E-01 | 8,80E-01 | 0.0087(0.0015) | 1.0214(0.0066) | 0.0011(0.0051) |
| Leucine | 27005778 | 0.2203(0.2152) | 1,0239 | 3,06E-01 | 8,80E-01 | 0.0436(0.0207) | 1.01(0.0066) | -0.003(0.0046) |
| Heel bone mineral density (BMD) T-score_ automated | 0 | 0.0512(0.05) | 1,0233 | 3,06E-01 | 8,80E-01 | 0.2911(0.0331) | 1.0889(0.0348) | -0.0057(0.007) |
| Diagnoses - main ICD10: O75 Other complications of labour and delivery_ not elsewhere classified | 0 | 0.246(0.2404) | 1,023 | 3,06E-01 | 8,80E-01 | 0.003(0.0012) | 0.9872(0.0057) | -0.0021(0.005) |
| Illness_ injury_ bereavement_ stress in last 2 years: Serious illness_ injury or assault of a close relative | 0 | 0.1269(0.1244) | 1,0197 | 3,08E-01 | 8,80E-01 | 0.0112(0.0017) | 1.0108(0.0071) | -0.0091(0.005) |
| Isoleucine | 27005778 | 0.1742(0.1711) | 1,018 | 3,09E-01 | 8,80E-01 | 0.0699(0.0234) | 0.9952(0.0065) | -0.002(0.0046) |
| Ever highly irritable/argumentative for 2 days | 0 | -0.1104(0.1085) | -1,0176 | 3,09E-01 | 8,80E-01 | 0.0508(0.0059) | 0.9992(0.0076) | -0.002(0.0047) |
| Serum creatinine (non-diabetes) | 26831199 | -0.0796(0.0786) | -1,0124 | 3,11E-01 | 8,80E-01 | 0.1149(0.0109) | 0.9791(0.0134) | 0.0035(0.0053) |
| Number of children ever born | 27798627 | -0.0904(0.0896) | -1,009 | 3,13E-01 | 8,80E-01 | 0.0252(0.0019) | 0.9709(0.0072) | 0.002(0.005) |
| Neo-openness to experience | 21173776 | -0.1706(0.1695) | -1,0069 | 3,14E-01 | 8,80E-01 | 0.1107(0.0274) | 0.9908(0.0066) | 0.004(0.0049) |
| Body fat percentage | 0 | -0.0496(0.0493) | -1,0061 | 3,14E-01 | 8,80E-01 | 0.2162(0.0067) | 1.0829(0.0174) | 0.0039(0.0066) |
| Non-cancer illness code_ self-reported: hayfever/allergic rhinitis | 0 | -0.0855(0.0852) | -1,0031 | 3,16E-01 | 8,80E-01 | 0.0223(0.0023) | 1.0189(0.0084) | 0.0014(0.0046) |
| Free cholesterol in very large HDL | 27005778 | 0.2291(0.2284) | 1,0027 | 3,16E-01 | 8,80E-01 | 0.0405(0.0245) | 1.0161(0.0133) | -0.0088(0.0046) |
| Handedness (chirality/laterality): Use both right and left hands equally | 0 | -0.1904(0.1901) | -1,0014 | 3,17E-01 | 8,80E-01 | 0.0045(0.0014) | 1.0045(0.0068) | 0.0026(0.0045) |
| Femoral neck bone mineral density | 22504420 | -0.0754(0.0755) | -0,9995 | 3,18E-01 | 8,80E-01 | 0.2964(0.0257) | 0.9809(0.0084) | 0.0079(0.0044) |
| Medication for cholesterol_ blood pressure_ diabetes_ or take exogenous hormones: Cholesterol lowering medication | 0 | -0.089(0.0892) | -0,9978 | 3,18E-01 | 8,80E-01 | 0.0507(0.0056) | 1.0244(0.0138) | 0.0013(0.0049) |
| Illnesses of father: Lung cancer | 0 | -0.1199(0.1211) | -0,9905 | 3,22E-01 | 8,88E-01 | 0.0147(0.0019) | 0.9939(0.0071) | -0.0038(0.0048) |
| Trunk fat percentage | 0 | -0.0488(0.0498) | -0,9795 | 3,27E-01 | 8,98E-01 | 0.2062(0.0068) | 1.0841(0.0175) | 0.0035(0.0066) |
| Non-cancer illness code_ self-reported: gout | 0 | -0.0843(0.0861) | -0,9788 | 3,28E-01 | 8,98E-01 | 0.0243(0.0062) | 0.9939(0.0171) | 0.0023(0.0051) |
| Years of schooling 2016 | 27225129 | -0.0556(0.0571) | -0,9735 | 3,30E-01 | 9,00E-01 | 0.1264(0.0046) | 0.927(0.0099) | 0.0104(0.0054) |
| Heel bone mineral density (BMD) T-score_ automated (right) | 0 | 0.0538(0.0556) | 0,9685 | 3,33E-01 | 9,05E-01 | 0.2971(0.0355) | 1.0346(0.0218) | -0.0001(0.0054) |
| Mouth/teeth dental problems: None of the above | 0 | 0.0784(0.0814) | 0,9626 | 3,36E-01 | 9,06E-01 | 0.0366(0.002) | 1.0206(0.0077) | 0.0014(0.0056) |

| | | | | | | | | |
|---|---|---|---|---|---|---|---|---|
| Diagnoses - main ICD10: R31 Unspecified haematuria | 0 | -0.2042(0.2124) | -0,9614 | 3,36E-01 | 9,06E-01 | 0.0033(0.0015) | 1.0065(0.0062) | 0.0015(0.0043) |
| Smoking status: Current | 0 | -0.0706(0.0736) | -0,9585 | 3,38E-01 | 9,09E-01 | 0.0513(0.0028) | 1.0146(0.0092) | -0.0056(0.0053) |
| Diagnoses - main ICD10: C44 Other malignant neoplasms of skin | 0 | -0.1171(0.1233) | -0,9497 | 3,42E-01 | 9,15E-01 | 0.0097(0.0023) | 1.0092(0.0088) | 0.0013(0.0047) |
| Illnesses of siblings: Heart disease | 0 | -0.1092(0.1152) | -0,9475 | 3,43E-01 | 9,15E-01 | 0.0175(0.0022) | 1.0107(0.0069) | 0.0042(0.0048) |
| Maximum heart rate during fitness test | 0 | 0.1337(0.1423) | 0,9392 | 3,48E-01 | 9,26E-01 | 0.0648(0.0132) | 1.0179(0.0085) | -0.0047(0.0049) |
| Non-cancer illness code_ self-reported: crohns disease | 0 | 0.146(0.1575) | 0,9265 | 3,54E-01 | 9,37E-01 | 0.0054(0.0014) | 1.0049(0.0067) | -0.0054(0.0042) |
| Diagnoses - main ICD10: I10 Essential (primary) hypertension | 0 | -0.2388(0.2582) | -0,9249 | 3,55E-01 | 9,37E-01 | 0.0025(0.0013) | 0.9975(0.0061) | 0.0054(0.0046) |
| Non-cancer illness code_ self-reported: pernicious anaemia | 0 | 0.1586(0.1716) | 0,924 | 3,56E-01 | 9,37E-01 | 0.0037(0.0013) | 0.9905(0.0065) | -0.0081(0.0042) |
| Leg pain on walking | 0 | -0.0907(0.0998) | -0,9091 | 3,63E-01 | 9,44E-01 | 0.0549(0.0052) | 1.0126(0.0068) | 0.004(0.0045) |
| Medication for pain relief_ constipation_ heartburn: Ibuprofen (e.g. Nurofen) | 0 | 0.0854(0.0942) | 0,9072 | 3,64E-01 | 9,44E-01 | 0.0221(0.0018) | 0.9996(0.0068) | -0.0047(0.0047) |
| Vitamin and mineral supplements: Vitamin C | 0 | 0.1092(0.1204) | 0,9067 | 3,65E-01 | 9,44E-01 | 0.0129(0.0016) | 0.9997(0.0072) | 0.0011(0.0048) |
| Alcohol intake versus 10 years previously | 0 | -0.0704(0.0777) | -0,9055 | 3,65E-01 | 9,44E-01 | 0.0345(0.0024) | 1.0296(0.0082) | 0.0027(0.0052) |
| Ever unenthusiastic/disinterested for a whole week | 0 | -0.0937(0.1037) | -0,9036 | 3,66E-01 | 9,44E-01 | 0.0567(0.0064) | 1.0001(0.0075) | 0.0069(0.005) |
| Pulse wave Arterial Stiffness index | 0 | -0.1036(0.1149) | -0,9012 | 3,68E-01 | 9,44E-01 | 0.0386(0.005) | 1.008(0.0071) | 0.0016(0.0044) |
| Neuroticism | 27089181 | 0.0705(0.0785) | 0,8985 | 3,69E-01 | 9,44E-01 | 0.0885(0.0076) | 0.9896(0.0119) | 0.0002(0.0056) |
| Years of schooling (proxy cognitive performance) | 25201988 | 0.0749(0.0837) | 0,8953 | 3,71E-01 | 9,44E-01 | 0.1062(0.0078) | 1.027(0.0096) | -0.001(0.0058) |
| Non-cancer illness code_ self-reported: uterine fibroids | 0 | 0.1435(0.1602) | 0,8954 | 3,71E-01 | 9,44E-01 | 0.0069(0.0014) | 1.0077(0.0069) | 0.0003(0.0048) |
| Vitamin and mineral supplements: Vitamin B | 0 | -0.1235(0.1393) | -0,8865 | 3,75E-01 | 9,44E-01 | 0.0074(0.0016) | 0.999(0.0068) | 0.0019(0.0041) |
| Mean Thalamus | 25607358 | -0.1603(0.1809) | -0,8861 | 3,76E-01 | 9,44E-01 | 0.1151(0.0355) | 0.9814(0.0062) | 0.0079(0.0044) |
| Why stopped smoking: Health precaution | 0 | 0.099(0.1121) | 0,8827 | 3,77E-01 | 9,44E-01 | 0.0538(0.0062) | 0.9995(0.0064) | 0.0006(0.0048) |
| Leg fat percentage (right) | 0 | -0.0442(0.0501) | -0,8821 | 3,78E-01 | 9,44E-01 | 0.2077(0.0061) | 1.0813(0.0167) | 0.0025(0.0065) |
| Duration walking for pleasure | 0 | 0.0979(0.111) | 0,8817 | 3,78E-01 | 9,44E-01 | 0.0254(0.0024) | 1.0108(0.0069) | -0.0039(0.0047) |
| Diagnoses - main ICD10: M25 Other joint disorders_ not elsewhere classified | 0 | -0.3757(0.4264) | -0,8811 | 3,78E-01 | 9,44E-01 | 0.0013(0.0013) | 1.0145(0.0065) | 0.0003(0.005) |
| Medication for cholesterol_ blood pressure_ diabetes_ or take exogenous hormones: None of the above | 0 | 0.069(0.0787) | 0,8764 | 3,81E-01 | 9,44E-01 | 0.0657(0.0045) | 1.0172(0.0085) | 0.0023(0.0053) |
| Diagnoses - main ICD10: F31 Bipolar affective disorder | 0 | -0.1657(0.1891) | -0,8762 | 3,81E-01 | 9,44E-01 | 0.0049(0.0015) | 0.9869(0.0073) | 0.0043(0.0047) |
| Mouth/teeth dental problems: Toothache | 0 | -0.1193(0.138) | -0,8647 | 3,87E-01 | 9,44E-01 | 0.0084(0.0016) | 1.0043(0.0071) | -0.0008(0.0051) |
| Diagnoses - main ICD10: K51 Ulcerative colitis | 0 | -0.1567(0.1823) | -0,8599 | 3,90E-01 | 9,44E-01 | 0.0048(0.0016) | 1.0127(0.0065) | -0.0006(0.0049) |
| Medication for cholesterol_ blood pressure or diabetes: Insulin | 0 | -0.1602(0.1867) | -0,8585 | 3,91E-01 | 9,44E-01 | 0.01(0.0033) | 1.0102(0.0072) | 0.0029(0.0049) |
| Underlying (primary) cause of death: ICD10: E85.4 Organ-limited amyloidosis | 0 | 0.2547(0.2969) | 0,8581 | 3,91E-01 | 9,44E-01 | 0.0792(0.0639) | 0.9986(0.0076) | -0.008(0.0049) |
| Usual walking pace | 0 | 0.0561(0.0658) | 0,8539 | 3,93E-01 | 9,44E-01 | 0.0779(0.0028) | 1.0275(0.0097) | 0.0011(0.0055) |
| Diagnoses - main ICD10: L03 Cellulitis | 0 | 0.1584(0.1859) | 0,8518 | 3,94E-01 | 9,44E-01 | 0.004(0.0015) | 1.0034(0.0067) | -0.0134(0.0041) |
| Reason for reducing amount of alcohol drunk: Health precaution | 0 | 0.0874(0.1026) | 0,8516 | 3,94E-01 | 9,44E-01 | 0.0455(0.0046) | 0.9975(0.007) | 0.0003(0.0044) |
| 22:6 docosahexaenoic acid | 27005778 | -0.1469(0.1735) | -0,8465 | 3,97E-01 | 9,44E-01 | 0.1339(0.0367) | 0.9942(0.0074) | -0.0005(0.0044) |
| Headaches for 3+ months | 0 | -0.1061(0.1259) | -0,8432 | 3,99E-01 | 9,44E-01 | 0.0543(0.0075) | 1.0078(0.0066) | 0.0024(0.005) |
| Medication for pain relief_ constipation_ heartburn: Laxatives (e.g. Dulcolax_ Senokot) | 0 | -0.1037(0.123) | -0,8431 | 3,99E-01 | 9,44E-01 | 0.0115(0.0017) | 1.0085(0.0069) | 0.0025(0.0048) |
| Hearing aid user | 0 | 0.1084(0.1293) | 0,8384 | 4,02E-01 | 9,44E-01 | 0.017(0.0027) | 1.0048(0.0068) | -0.0043(0.005) |
| Description of average fatty acid chain length; not actual carbon number | 27005778 | -0.1379(0.1647) | -0,8374 | 4,02E-01 | 9,44E-01 | 0.0985(0.0364) | 0.9838(0.0068) | -0.0005(0.0044) |
| Vascular/heart problems diagnosed by doctor: High blood pressure | 0 | -0.0445(0.0534) | -0,832 | 4,05E-01 | 9,44E-01 | 0.1152(0.005) | 1.0603(0.0164) | -0.0052(0.0056) |
| Fathers age at death | 0 | 0.0782(0.0944) | 0,8286 | 4,07E-01 | 9,44E-01 | 0.0333(0.003) | 1.0239(0.0075) | 0.0066(0.0054) |
| Diagnoses - main ICD10: M67 Other disorders of synovium and tendon | 0 | -0.2118(0.2558) | -0,8282 | 4,08E-01 | 9,44E-01 | 0.0021(0.0013) | 0.9996(0.0062) | 0.0042(0.0045) |
| Eye problems/disorders: Cataract | 0 | -0.1255(0.1516) | -0,828 | 4,08E-01 | 9,44E-01 | 0.0226(0.0044) | 1.0143(0.0072) | 0.0046(0.0045) |
| Average weekly intake of other alcoholic drinks | 0 | -0.2157(0.2613) | -0,8255 | 4,09E-01 | 9,44E-01 | 0.0101(0.0062) | 1.0014(0.0066) | -0.004(0.0047) |
| Noisy workplace | 0 | -0.0789(0.0959) | -0,8224 | 4,11E-01 | 9,44E-01 | 0.0614(0.0057) | 1.0274(0.0077) | 0.0037(0.0052) |
| Adopted as a child | 0 | 0.1475(0.1798) | 0,8204 | 4,12E-01 | 9,44E-01 | 0.0058(0.0014) | 1.0021(0.0068) | -0.0051(0.0046) |
| Forced vital capacity (FVC) | 0 | 0.0503(0.0613) | 0,8196 | 4,13E-01 | 9,44E-01 | 0.1686(0.0075) | 1.0958(0.0171) | -0.0046(0.0065) |
| Number of days/week walked 10+ minutes | 0 | 0.0602(0.074) | 0,8135 | 4,16E-01 | 9,44E-01 | 0.0422(0.0022) | 1.0096(0.0075) | 0.0017(0.0051) |
| Other eye problems | 0 | -0.123(0.1519) | -0,8099 | 4,18E-01 | 9,44E-01 | 0.0083(0.0014) | 1.0271(0.007) | 0.0004(0.005) |
| Non-cancer illness code_ self-reported: enlarged prostate | 0 | -0.1406(0.1739) | -0,8082 | 4,19E-01 | 9,44E-01 | 0.0056(0.0015) | 1.006(0.0066) | 0.0025(0.0046) |
| Forced expiratory volume in 1 second (FEV1)/Forced Vital capacity(FVC) | 28166213 | 0.0623(0.0775) | 0,8039 | 4,22E-01 | 9,44E-01 | 0.2564(0.0206) | 0.972(0.0084) | -0.008(0.0046) |
| Guilty feelings | 0 | -0.0553(0.0689) | -0,8022 | 4,22E-01 | 9,44E-01 | 0.0506(0.0028) | 1.0089(0.0089) | 0.0018(0.0053) |
| Forced vital capacity (FVC)_ Best measure | 0 | 0.0497(0.062) | 0,802 | 4,23E-01 | 9,44E-01 | 0.1871(0.0087) | 1.0823(0.0166) | -0.0035(0.0063) |
| Duration of heavy DIY | 0 | -0.0822(0.1026) | -0,8009 | 4,23E-01 | 9,44E-01 | 0.0331(0.0038) | 1.0082(0.0068) | -0.0032(0.0046) |
| Morning/evening person (chronotype) | 0 | -0.0439(0.0549) | -0,7999 | 4,24E-01 | 9,44E-01 | 0.116(0.0044) | 1.0415(0.0118) | 0.0088(0.0058) |
| Extreme bmi | 23563607 | -0.0753(0.0947) | -0,7957 | 4,26E-01 | 9,44E-01 | 0.689(0.0553) | 1.0305(0.0113) | 0.0002(0.0066) |
| Comparative height size at age 10 | 0 | -0.0481(0.0606) | -0,7948 | 4,27E-01 | 9,44E-01 | 0.2404(0.0127) | 1.1119(0.0353) | 0.0058(0.007) |
| Diagnoses - main ICD10: F43 Reaction to severe stress_ and adjustment disorders | 0 | -0.3347(0.422) | -0,7931 | 4,28E-01 | 9,44E-01 | 0.001(0.0013) | 1.0016(0.0058) | 0.0054(0.0047) |
| Diagnoses - main ICD10: M23 Internal derangement of knee | 0 | -0.1066(0.1346) | -0,792 | 4,28E-01 | 9,44E-01 | 0.008(0.0019) | 1.0051(0.0074) | 0.0008(0.0045) |
| Reason for glasses/contact lenses: For long-sightedness_ i.e. for distance and near_ but particularly for near tasks like reading (called hypermetropia) | 0 | -0.1152(0.1457) | -0,7907 | 4,29E-01 | 9,44E-01 | 0.0099(0.0016) | 1.0637(0.007) | 0.0028(0.0048) |
| Intelligence | 28530673 | 0.0526(0.0667) | 0,7892 | 4,30E-01 | 9,44E-01 | 0.1881(0.0104) | 1.0093(0.0082) | -0.0094(0.0043) |
| Pulse wave peak to peak time | 0 | 0.0883(0.1122) | 0,7878 | 4,31E-01 | 9,44E-01 | 0.0393(0.0048) | 1.0095(0.007) | -0.0019(0.0044) |
| Frequency of light DIY in last 4 weeks | 0 | -0.0846(0.1075) | -0,7872 | 4,31E-01 | 9,44E-01 | 0.023(0.003) | 1.0035(0.0064) | 0.0057(0.0046) |
| Illnesses of father: Prostate cancer | 0 | 0.1187(0.1508) | 0,7874 | 4,31E-01 | 9,44E-01 | 0.0077(0.0018) | 1.0133(0.0079) | 0.0025(0.005) |
| Non-cancer illness code_ self-reported: muscle or soft tissue injuries | 0 | 0.2182(0.2779) | 0,7853 | 4,32E-01 | 9,44E-01 | 0.0021(0.0015) | 1.0016(0.0073) | -0.0038(0.0046) |
| Types of physical activity in last 4 weeks: Other exercises (eg: swimming_ cycling_ keep fit_ bowling) | 0 | 0.058(0.0739) | 0,7843 | 4,33E-01 | 9,44E-01 | 0.0458(0.0023) | 1.0152(0.0084) | -0.0009(0.0052) |
| Qualifications: NVQ or HND or HNC or equivalent | 0 | -0.0931(0.1188) | -0,7836 | 4,33E-01 | 9,44E-01 | 0.015(0.0017) | 1.011(0.007) | -0.0009(0.0053) |
| Glutamine | 27005778 | 0.1343(0.1717) | 0,782 | 4,34E-01 | 9,44E-01 | 0.0613(0.0201) | 1.0174(0.0072) | 0.0006(0.0043) |
| Qualifications: CSEs or equivalent | 0 | 0.0786(0.1007) | 0,7811 | 4,35E-01 | 9,44E-01 | 0.0209(0.0019) | 1.0596(0.0069) | -0.0105(0.0051) |
| Mood swings | 0 | 0.0497(0.0638) | 0,778 | 4,37E-01 | 9,44E-01 | 0.0697(0.0032) | 1.0191(0.0099) | -0.0035(0.0051) |
| Average number of methylene groups per a double bond | 27005778 | -0.1179(0.1516) | -0,7775 | 4,37E-01 | 9,44E-01 | 0.1951(0.0792) | 0.9954(0.0078) | 0.0012(0.0047) |
| Diagnoses - main ICD10: I30 Acute pericarditis | 0 | 0.2156(0.2775) | 0,7767 | 4,37E-01 | 9,44E-01 | 0.0021(0.0014) | 0.9957(0.0068) | -0.0014(0.0047) |

| | | | | | | | | |
|---|---|---|---|---|---|---|---|---|
| Urate | 23263486 | 0.0487(0.063) | 0,7731 | 4,40E-01 | 9,44E-01 | 0.1757(0.0589) | 0.9542(0.0488) | -0.0074(0.0052) |
| Diagnoses - main ICD10: N81 Female genital prolapse | 0 | 0.1142(0.1479) | 0,7722 | 4,40E-01 | 9,44E-01 | 0.0072(0.0015) | 1.0234(0.0065) | -0.0036(0.0046) |
| Fracture resulting from simple fall | 0 | -0.1337(0.1734) | -0,7709 | 4,41E-01 | 9,44E-01 | 0.049(0.0138) | 0.9929(0.0063) | 0.0042(0.0044) |
| Diagnoses - main ICD10: S76 Injury of muscle and tendon at hip and thigh level | 0 | 0.2107(0.2743) | 0,7681 | 4,42E-01 | 9,44E-01 | 0.0018(0.0012) | 0.9973(0.0064) | 0.0049(0.0047) |
| Diagnoses - main ICD10: M10 Gout | 0 | -0.1861(0.2445) | -0,7609 | 4,47E-01 | 9,49E-01 | 0.0023(0.0015) | 0.9994(0.0068) | 0.0011(0.0045) |
| Time spent using computer | 0 | -0.0493(0.0649) | -0,7593 | 4,48E-01 | 9,49E-01 | 0.0961(0.0043) | 1.0434(0.0095) | -0.0003(0.0056) |
| Triglycerides | 20686565 | 0.0537(0.0709) | 0,7575 | 4,49E-01 | 9,49E-01 | 0.1686(0.0298) | 0.9686(0.0163) | -0.0088(0.0051) |
| Alanine | 27005778 | 0.1311(0.1743) | 0,7523 | 4,52E-01 | 9,49E-01 | 0.0842(0.0266) | 1.012(0.0074) | -0.0026(0.0046) |
| Time from waking to first cigarette | 0 | -0.1325(0.1764) | -0,7512 | 4,53E-01 | 9,49E-01 | 0.0827(0.0258) | 1.0162(0.0075) | 0.0048(0.0048) |
| Diagnoses - main ICD10: T84 Complications of internal orthopaedic prosthetic devices_ implants and grafts | 0 | -0.1299(0.1732) | -0,7503 | 4,53E-01 | 9,49E-01 | 0.0053(0.0016) | 0.991(0.0072) | 0.0037(0.0047) |
| Mineral and other dietary supplements: Zinc | 0 | 0.09(0.1205) | 0,747 | 4,55E-01 | 9,49E-01 | 0.0112(0.0015) | 0.988(0.006) | 0.0043(0.0047) |
| Number of live births | 0 | -0.0615(0.0826) | -0,7449 | 4,56E-01 | 9,49E-01 | 0.0613(0.0043) | 1.0083(0.0084) | 0.0052(0.0051) |
| Non-cancer illness code_ self-reported: anxiety/panic attacks | 0 | 0.1276(0.172) | 0,7419 | 4,58E-01 | 9,49E-01 | 0.0058(0.0012) | 1.0008(0.0061) | -0.0003(0.0048) |
| Types of physical activity in last 4 weeks: Walking for pleasure (not as a means of transport) | 0 | 0.058(0.0783) | 0,7407 | 4,59E-01 | 9,49E-01 | 0.0376(0.0021) | 1.0389(0.0071) | -0.0004(0.0054) |
| Back pain for 3+ months | 0 | -0.092(0.125) | -0,7361 | 4,62E-01 | 9,49E-01 | 0.0408(0.0058) | 0.9922(0.0067) | -0.0002(0.0048) |
| Diagnoses - main ICD10: J34 Other disorders of nose and nasal sinuses | 0 | -0.1785(0.2424) | -0,7362 | 4,62E-01 | 9,49E-01 | 0.0026(0.0013) | 1.0074(0.0065) | -0.0006(0.005) |
| Cancer code_ self-reported: small intestine/small bowel cancer | 0 | -0.2747(0.3735) | -0,7356 | 4,62E-01 | 9,49E-01 | 0.0013(0.0015) | 0.9981(0.0071) | 0.0009(0.0043) |
| Total cholesterol in very large HDL | 27005778 | 0.2057(0.2798) | 0,735 | 4,62E-01 | 9,49E-01 | 0.0262(0.0225) | 1.015(0.0114) | -0.0069(0.0045) |
| Diagnoses - main ICD10: M20 Acquired deformities of fingers and toes | 0 | -0.0963(0.1313) | -0,7338 | 4,63E-01 | 9,49E-01 | 0.0093(0.0016) | 1.0008(0.0073) | 0.0028(0.005) |
| Amyotrophic lateral sclerosis | 27455348 | 0.1322(0.1822) | 0,7252 | 4,68E-01 | 9,56E-01 | 0.0482(0.0132) | 0.9943(0.0064) | -0.0026(0.0049) |
| Serum cystatin c | 26831199 | -0.0795(0.11) | -0,7228 | 4,70E-01 | 9,58E-01 | 0.1681(0.0688) | 0.9621(0.0145) | 0.0063(0.0046) |
| Creatinine | 27005778 | 0.1119(0.1557) | 0,7187 | 4,72E-01 | 9,59E-01 | 0.1119(0.0256) | 1.0135(0.0072) | -0.0032(0.0045) |
| Non-cancer illness code_ self-reported: hypertension | 0 | -0.0387(0.0545) | -0,711 | 4,77E-01 | 9,62E-01 | 0.1134(0.0049) | 1.0554(0.0164) | -0.005(0.0058) |
| Diagnoses - main ICD10: R04 Haemorrhage from respiratory passages | 0 | -0.1519(0.2137) | -0,7108 | 4,77E-01 | 9,62E-01 | 0.0029(0.0015) | 0.996(0.0064) | 0.0065(0.0044) |
| Number of treatments/medications taken | 0 | -0.0496(0.07) | -0,7086 | 4,79E-01 | 9,62E-01 | 0.0603(0.0025) | 1.0322(0.0089) | -0.0016(0.0053) |
| Non-cancer illness code_ self-reported: vitiligo | 0 | 0.2183(0.309) | 0,7067 | 4,80E-01 | 9,62E-01 | 0.0015(0.0014) | 0.994(0.007) | -0.0042(0.0043) |
| Hair/balding pattern: Pattern 3 | 0 | 0.0628(0.0889) | 0,7057 | 4,80E-01 | 9,62E-01 | 0.0524(0.0118) | 1.0241(0.0162) | -0.0042(0.0048) |
| Non-cancer illness code_ self-reported: hypopituitarism | 0 | 0.235(0.3331) | 0,7055 | 4,81E-01 | 9,62E-01 | 0.0014(0.0012) | 0.9912(0.0062) | -0.0053(0.0045) |
| Tobacco smoking: Ex-smoker | 0 | -0.0756(0.1077) | -0,7018 | 4,83E-01 | 9,63E-01 | 0.0741(0.008) | 1.01(0.0078) | -0.0015(0.0048) |
| Smoking/smokers in household | 0 | 0.0974(0.1416) | 0,688 | 4,91E-01 | 9,74E-01 | 0.0098(0.0019) | 1.0113(0.0076) | -0.0015(0.005) |
| Non-cancer illness code_ self-reported: eczema/dermatitis | 0 | 0.1052(0.153) | 0,6877 | 4,92E-01 | 9,74E-01 | 0.0079(0.0022) | 1.0168(0.0067) | -0.0043(0.0049) |
| Total cholesterol in large VLDL | 27005778 | -0.1095(0.1599) | -0,6849 | 4,93E-01 | 9,74E-01 | 0.1181(0.0297) | 0.9924(0.0066) | -0.000088008(0.0048) |
| Birth weight | 0 | 0.049(0.0716) | 0,6847 | 4,94E-01 | 9,74E-01 | 0.1086(0.0061) | 1.0493(0.0136) | -0.0026(0.0056) |
| Medication for cholesterol_ blood pressure_ diabetes_ or take exogenous hormones: Blood pressure medication | 0 | -0.0446(0.0653) | -0,683 | 4,95E-01 | 9,74E-01 | 0.1031(0.0066) | 1.0336(0.0116) | -0.0034(0.0054) |
| Alzheimers disease | 24162737 | -0.1156(0.1706) | -0,6775 | 4,98E-01 | 9,77E-01 | 0.0421(0.021) | 1.0672(0.0294) | 0.0049(0.0051) |
| Mineral and other dietary supplements: Fish oil (including cod liver oil) | 0 | 0.0623(0.0921) | 0,6767 | 4,99E-01 | 9,77E-01 | 0.0268(0.0018) | 1.0058(0.007) | 0.0022(0.0052) |
| Fractured bone site(s): Ankle | 0 | -0.1094(0.1629) | -0,6718 | 5,02E-01 | 9,79E-01 | 0.0063(0.0016) | 0.9948(0.0069) | 0.001(0.0049) |
| Anorexia Nervosa | 24514567 | -0.0503(0.0753) | -0,6678 | 5,04E-01 | 9,79E-01 | 0.5602(0.0317) | 0.8958(0.007) | -0.0024(0.0047) |
| Non-cancer illness code_ self-reported: ulcerative colitis | 0 | -0.1133(0.1697) | -0,6676 | 5,04E-01 | 9,79E-01 | 0.0062(0.0017) | 1.009(0.0078) | -0.0033(0.0047) |
| Phospholipids in medium LDL | 27005778 | 0.1502(0.2263) | 0,664 | 5,07E-01 | 9,83E-01 | 0.0695(0.0435) | 1.0088(0.0193) | -0.003(0.0047) |
| Types of transport used (excluding work): Walk | 0 | -0.0568(0.0863) | -0,658 | 5,11E-01 | 9,87E-01 | 0.0333(0.0023) | 1.0342(0.0087) | 0.0041(0.0051) |
| LDL cholesterol | 20686565 | -0.0659(0.1009) | -0,6526 | 5,14E-01 | 9,87E-01 | 0.1021(0.0293) | 1.0524(0.0487) | 0.0008(0.0057) |
| Acetoacetate | 27005778 | 0.1282(0.1969) | 0,651 | 5,15E-01 | 9,87E-01 | 0.076(0.0285) | 0.9796(0.0074) | -0.0005(0.0048) |
| Birth weight | 27680694 | -0.0542(0.0837) | -0,647 | 5,18E-01 | 9,87E-01 | 0.103(0.0066) | 1.0392(0.0095) | -0.0024(0.0047) |
| Birth weight of first child | 0 | 0.05(0.0774) | 0,6467 | 5,18E-01 | 9,87E-01 | 0.1067(0.0063) | 1.0237(0.0099) | -0.0001(0.0057) |
| Job involves shift work | 0 | -0.0661(0.1029) | -0,6423 | 5,21E-01 | 9,87E-01 | 0.0301(0.0028) | 1.0023(0.0064) | 0.0008(0.0047) |
| Mineral and other dietary supplements: None of the above | 0 | -0.0535(0.0841) | -0,6357 | 5,25E-01 | 9,87E-01 | 0.0302(0.002) | 1.0091(0.0067) | -0.0033(0.0047) |
| Diagnoses - main ICD10: N40 Hyperplasia of prostate | 0 | -0.1029(0.1624) | -0,6333 | 5,27E-01 | 9,87E-01 | 0.0061(0.0015) | 1.0091(0.0068) | 0.0038(0.0044) |
| Illnesses of siblings: None of the above (group 1) | 0 | 0.0537(0.0851) | 0,6317 | 5,28E-01 | 9,87E-01 | 0.0364(0.0026) | 1.0134(0.0078) | -0.0012(0.0051) |
| Why stopped smoking: Doctors advice | 0 | 0.1204(0.1925) | 0,6254 | 5,32E-01 | 9,87E-01 | 0.0192(0.0051) | 0.9963(0.0058) | -0.004(0.0048) |
| Illnesses of father: High blood pressure | 0 | 0.063(0.1015) | 0,6201 | 5,35E-01 | 9,87E-01 | 0.0189(0.0018) | 1.0138(0.007) | -0.008(0.0051) |
| Mean Hippocampus | 25607358 | -0.1145(0.1867) | -0,6132 | 5,40E-01 | 9,87E-01 | 0.138(0.0406) | 0.9872(0.0067) | -0.0024(0.0052) |
| Duration of moderate activity | 0 | -0.0605(0.0989) | -0,6117 | 5,41E-01 | 9,87E-01 | 0.0317(0.0023) | 1.0094(0.0075) | -0.0001(0.0052) |
| Years of schooling 2013 | 23722424 | 0.0527(0.0865) | 0,6096 | 5,42E-01 | 9,87E-01 | 0.0838(0.0064) | 1.0189(0.0091) | -0.0004(0.0058) |
| Forearm Bone mineral density | 26367794 | 0.1444(0.2376) | 0,608 | 5,43E-01 | 9,87E-01 | 0.0887(0.0444) | 1.0109(0.0068) | -0.0029(0.0043) |
| Pulse wave reflection index | 0 | -0.0637(0.1048) | -0,6077 | 5,43E-01 | 9,87E-01 | 0.0548(0.0052) | 1.0076(0.0073) | 0.0011(0.0049) |
| Age when periods started (menarche) | 0 | 0.0317(0.0524) | 0,605 | 5,45E-01 | 9,87E-01 | 0.2095(0.0109) | 1.0029(0.013) | -0.0103(0.0053) |
| Illness_ injury_ bereavement_ stress in last 2 years: Serious illness_ injury or assault to yourself | 0 | -0.0681(0.1127) | -0,6047 | 5,45E-01 | 9,87E-01 | 0.0141(0.0016) | 1.0117(0.0067) | -0.0074(0.0048) |
| Suffer from nerves | 0 | 0.0463(0.0779) | 0,5943 | 5,52E-01 | 9,87E-01 | 0.045(0.0032) | 1.0002(0.0097) | -0.0035(0.0052) |
| Duration of fitness test | 0 | -0.1259(0.2119) | -0,594 | 5,53E-01 | 9,87E-01 | 0.0232(0.0089) | 1.0032(0.0064) | 0.0016(0.0046) |
| Mean diameter for LDL particles | 27005778 | -0.147(0.2476) | -0,5939 | 5,53E-01 | 9,87E-01 | 0.0471(0.0266) | 1.0008(0.012) | 0.0026(0.0047) |
| Diagnoses - main ICD10: M54 Dorsalgia | 0 | -0.0771(0.1304) | -0,5912 | 5,54E-01 | 9,87E-01 | 0.01(0.0015) | 1.0049(0.0071) | -0.0003(0.005) |
| Depressive symptoms | 27089181 | -0.0553(0.0936) | -0,5903 | 5,55E-01 | 9,87E-01 | 0.047(0.0036) | 1.0013(0.0069) | 0.0015(0.0052) |
| Standing height | 0 | -0.0322(0.0546) | -0,5899 | 5,55E-01 | 9,87E-01 | 0.4494(0.0235) | 1.2494(0.0595) | -0.0009(0.0082) |
| Waist circumference | 25673412 | 0.0397(0.0677) | 0,5858 | 5,58E-01 | 9,87E-01 | 0.1199(0.0052) | 0.8462(0.0076) | -0.0095(0.0049) |
| Former vs Current smoker | 20418890 | -0.0851(0.1461) | -0,5829 | 5,60E-01 | 9,87E-01 | 0.063(0.0104) | 1(0.0067) | 0.0005(0.0044) |
| Concentration of very large HDL particles | 27005778 | 0.1128(0.194) | 0,5816 | 5,61E-01 | 9,87E-01 | 0.0588(0.0258) | 1.0038(0.0143) | -0.0083(0.0046) |
| Illness_ injury_ bereavement_ stress in last 2 years: Marital separation/divorce | 0 | -0.1173(0.2018) | -0,5812 | 5,61E-01 | 9,87E-01 | 0.0042(0.0014) | 1.009(0.0067) | 0.0037(0.0046) |
| Non-cancer illness code_ self-reported: psoriasis | 0 | -0.0918(0.1583) | -0,5799 | 5,62E-01 | 9,87E-01 | 0.0069(0.0016) | 1.0179(0.008) | 0.0019(0.0047) |
| Diagnoses - main ICD10: R69 Unknown and unspecified causes of morbidity | 0 | -0.109(0.1895) | -0,5752 | 5,65E-01 | 9,87E-01 | 0.005(0.0014) | 0.9991(0.0067) | 0.0057(0.005) |

| | | | | | | | | |
|---|---|---|---|---|---|---|---|---|
| Diagnoses - main ICD10: B37 Candidiasis | 0 | 0.9102(1.6102) | 0,5653 | 5,72E-01 | 9,87E-01 | 0.0006(0.0014) | 1.001(0.0062) | -0.0114(0.0043) |
| Home area population density - urban or rural: England/Wales - Village - less sparse | 0 | 0.1756(0.3109) | 0,5648 | 5,72E-01 | 9,87E-01 | 0.0019(0.0015) | 1.0246(0.0065) | 0.0013(0.0052) |
| Diagnoses - main ICD10: I48 Atrial fibrillation and flutter | 0 | -0.0654(0.1171) | -0,559 | 5,76E-01 | 9,87E-01 | 0.0132(0.0022) | 1.0097(0.0078) | -0.0013(0.0044) |
| Diagnoses - main ICD10: K52 Other non-infective gastro-enteritis and colitis | 0 | -0.0985(0.1762) | -0,5587 | 5,76E-01 | 9,87E-01 | 0.0048(0.0013) | 1.0039(0.0062) | -0.002(0.0043) |
| Diagnoses - main ICD10: N19 Unspecified renal failure | 0 | 0.1337(0.2396) | 0,5581 | 5,77E-01 | 9,87E-01 | 0.003(0.0014) | 0.9992(0.0065) | -0.0057(0.0045) |
| Free cholesterol to esterified cholesterol ratio | 27005778 | 0.1553(0.2795) | 0,5556 | 5,79E-01 | 9,87E-01 | 0.0577(0.0515) | 1.0127(0.0129) | -0.0039(0.0046) |
| Getting up in morning | 0 | -0.035(0.0638) | -0,5483 | 5,84E-01 | 9,87E-01 | 0.0689(0.0032) | 1.0376(0.0106) | -0.0015(0.0055) |
| Ever stopped smoking for 6+ months | 0 | -0.0998(0.1827) | -0,5462 | 5,85E-01 | 9,87E-01 | 0.02(0.0058) | 1.0013(0.007) | 0.0058(0.0043) |
| Diagnoses - main ICD10: K40 Inguinal hernia | 0 | -0.0505(0.0925) | -0,5458 | 5,85E-01 | 9,87E-01 | 0.0184(0.0022) | 0.9916(0.0069) | 0.001(0.0049) |
| Sleep duration | 27494321 | 0.0504(0.0923) | 0,5457 | 5,85E-01 | 9,87E-01 | 0.056(0.0051) | 1.0177(0.0072) | -0.0016(0.0045) |
| Job involves mainly walking or standing | 0 | 0.0415(0.0762) | 0,5449 | 5,86E-01 | 9,87E-01 | 0.08(0.0046) | 1.0307(0.009) | -0.0045(0.0051) |
| Total cholesterol in LDL | 27005778 | 0.1281(0.2378) | 0,5386 | 5,90E-01 | 9,87E-01 | 0.0654(0.0515) | 1.0141(0.024) | -0.0019(0.0048) |
| Potassium in urine | 0 | -0.0375(0.0696) | -0,5383 | 5,90E-01 | 9,87E-01 | 0.0423(0.0022) | 1.0183(0.0083) | 0.0031(0.0047) |
| Total cholesterol in medium VLDL | 27005778 | -0.0831(0.1543) | -0,5381 | 5,91E-01 | 9,87E-01 | 0.1265(0.0355) | 0.9926(0.0076) | 0.0022(0.0046) |
| Fractured bone site(s): Wrist | 0 | -0.0924(0.1717) | -0,5381 | 5,91E-01 | 9,87E-01 | 0.0061(0.0016) | 1.0099(0.0068) | 0.0006(0.0051) |
| Average weekly fortified wine intake | 0 | 0.0685(0.1278) | 0,5361 | 5,92E-01 | 9,87E-01 | 0.0127(0.0022) | 1.0168(0.0066) | -0.0013(0.004) |
| Diagnoses - main ICD10: N20 Calculus of kidney and ureter | 0 | 0.0835(0.1562) | 0,5347 | 5,93E-01 | 9,87E-01 | 0.008(0.0015) | 1.0081(0.0065) | -0.0029(0.0047) |
| Illness_ injury_ bereavement_ stress in last 2 years: Death of a close relative | 0 | 0.0889(0.1682) | 0,5285 | 5,97E-01 | 9,87E-01 | 0.006(0.0015) | 1.005(0.0063) | -0.0054(0.0048) |
| Ever manic/hyper for 2 days | 0 | 0.0838(0.1594) | 0,5256 | 5,99E-01 | 9,87E-01 | 0.0192(0.0047) | 0.9943(0.0065) | 0.0005(0.0042) |
| Average weekly red wine intake | 0 | 0.0412(0.0784) | 0,5251 | 6,00E-01 | 9,87E-01 | 0.0556(0.0034) | 1.015(0.008) | -0.0023(0.0052) |
| Eye problems/disorders: Diabetes related eye disease | 0 | 0.0733(0.14) | 0,5234 | 6,01E-01 | 9,87E-01 | 0.0198(0.0044) | 1.0067(0.0062) | -0.0009(0.0042) |
| Forced expiratory volume in 1 second (FEV1)/Forced Vital capacity(FVC) | 21946350 | 0.0591(0.113) | 0,5229 | 6,01E-01 | 9,87E-01 | 0.1219(0.0135) | 0.9296(0.008) | -0.0053(0.0055) |
| Illnesses of father: None of the above (group 1) | 0 | 0.07(0.1343) | 0,5213 | 6,02E-01 | 9,87E-01 | 0.0143(0.0018) | 1.0235(0.0067) | -0.0007(0.0051) |
| Diagnoses - main ICD10: K21 Gastro-oesophageal reflux disease | 0 | -0.0811(0.1562) | -0,5194 | 6,04E-01 | 9,87E-01 | 0.0061(0.0013) | 1.0024(0.0058) | 0.0094(0.0049) |
| Duration of light DIY | 0 | 0.0489(0.0949) | 0,516 | 6,06E-01 | 9,87E-01 | 0.0349(0.0034) | 1.0041(0.0076) | -0.0045(0.0045) |
| Diagnoses - main ICD10: M72 Fibroblastic disorders | 0 | -0.0677(0.1315) | -0,5148 | 6,07E-01 | 9,87E-01 | 0.0106(0.003) | 1.0168(0.0208) | -0.000070892(0.0049) |
| Ever had hysterectomy (womb removed) | 0 | -0.063(0.1242) | -0,5069 | 6,12E-01 | 9,87E-01 | 0.0221(0.0034) | 1.0055(0.0076) | 0.0035(0.0046) |
| Triglycerides in small HDL | 27005778 | -0.0991(0.196) | -0,5054 | 6,13E-01 | 9,87E-01 | 0.0632(0.0271) | 0.9998(0.008) | 0.0025(0.0043) |
| Time spent watching television (TV) | 0 | -0.0264(0.0522) | -0,5053 | 6,13E-01 | 9,87E-01 | 0.099(0.004) | 1.0577(0.0108) | -0.000089822(0.0057) |
| Forced Vital capacity(FVC) | 26635082 | 0.0543(0.1076) | 0,5047 | 6,14E-01 | 9,87E-01 | 0.1437(0.0143) | 0.9868(0.0063) | 0.0022(0.0047) |
| Lung cancer (all) | 24880342 | -0.0587(0.1164) | -0,5037 | 6,14E-01 | 9,87E-01 | 0.1272(0.0294) | 1.0072(0.0077) | -0.0007(0.0047) |
| Difficulty not smoking for 1 day | 0 | -0.0677(0.1356) | -0,4997 | 6,17E-01 | 9,87E-01 | 0.1133(0.0217) | 0.9885(0.0064) | -0.003(0.0047) |
| Neuroticism | 24828478 | 0.0758(0.1527) | 0,4965 | 6,20E-01 | 9,87E-01 | 0.0152(0.0034) | 1.0074(0.0063) | -0.0017(0.0048) |
| Non-cancer illness code_ self-reported: rheumatoid arthritis | 0 | -0.0978(0.1978) | -0,4945 | 6,21E-01 | 9,87E-01 | 0.0048(0.0014) | 1.0041(0.0066) | -0.0016(0.0051) |
| Age started oral contraceptive pill | 0 | 0.0505(0.1027) | 0,4919 | 6,23E-01 | 9,87E-01 | 0.0485(0.0042) | 1.0149(0.0075) | 0.0031(0.0049) |
| Other serious medical condition/disability diagnosed by doctor | 0 | -0.0435(0.089) | -0,4886 | 6,25E-01 | 9,87E-01 | 0.0249(0.0018) | 1.0118(0.007) | 0.0011(0.0044) |
| Diagnoses - main ICD10: I20 Angina pectoris | 0 | -0.0749(0.1536) | -0,4877 | 6,26E-01 | 9,87E-01 | 0.0079(0.0015) | 1.0101(0.007) | -0.0012(0.005) |
| Diagnoses - main ICD10: D25 Leiomyoma of uterus | 0 | -0.0793(0.1627) | -0,4872 | 6,26E-01 | 9,87E-01 | 0.0059(0.0014) | 0.9929(0.0062) | -0.007(0.0048) |
| Transport type for commuting to job workplace: Cycle | 0 | 0.0482(0.0994) | 0,485 | 6,28E-01 | 9,87E-01 | 0.031(0.0033) | 1.0162(0.0075) | -0.001(0.0047) |
| Pulse rate | 0 | -0.0394(0.0816) | -0,4835 | 6,29E-01 | 9,87E-01 | 0.1336(0.0122) | 1.026(0.0144) | 0.0009(0.0059) |
| Non-cancer illness code_ self-reported: osteoporosis | 0 | 0.0507(0.1053) | 0,4814 | 6,30E-01 | 9,87E-01 | 0.0146(0.0018) | 1.0009(0.0074) | -0.0027(0.0048) |
| Cancer code_ self-reported: malignant melanoma | 0 | -0.0892(0.1856) | -0,4809 | 6,31E-01 | 9,87E-01 | 0.0046(0.0015) | 0.9942(0.0061) | -0.0021(0.0043) |
| Comparative body size at age 10 | 0 | -0.0242(0.0505) | -0,4798 | 6,31E-01 | 9,87E-01 | 0.1445(0.0089) | 1.0179(0.0159) | 0.001(0.0054) |
| Diagnoses - main ICD10: K29 Gastritis and duodenitis | 0 | -0.0684(0.1427) | -0,4793 | 6,32E-01 | 9,87E-01 | 0.0072(0.0015) | 1.0051(0.0064) | -0.0028(0.0045) |
| Mouth/teeth dental problems: Dentures | 0 | -0.0332(0.0698) | -0,476 | 6,34E-01 | 9,87E-01 | 0.0471(0.0026) | 1.0387(0.0093) | -0.0049(0.005) |
| Concentration of small LDL particles | 27005778 | 0.0982(0.2075) | 0,4733 | 6,36E-01 | 9,87E-01 | 0.0907(0.0447) | 1.0076(0.0161) | -0.0028(0.0047) |
| Father still alive | 0 | 0.0743(0.1576) | 0,4717 | 6,37E-01 | 9,87E-01 | 0.0066(0.0015) | 1.0161(0.0065) | -0.0036(0.005) |
| Number of older siblings | 0 | 0.1608(0.3419) | 0,4703 | 6,38E-01 | 9,87E-01 | 0.0047(0.0051) | 1.0082(0.006) | 0.0018(0.0045) |
| Number of days/week of vigorous physical activity 10+ minutes | 0 | -0.0366(0.078) | -0,4696 | 6,39E-01 | 9,87E-01 | 0.0377(0.0023) | 1.0159(0.0081) | 0.0133(0.0048) |
| Doctor diagnosed hayfever or allergic rhinitis | 0 | 0.046(0.0982) | 0,4682 | 6,40E-01 | 9,87E-01 | 0.0772(0.0088) | 1.0097(0.0084) | -0.0055(0.0045) |
| Seen a psychiatrist for nerves_ anxiety_ tension or depression | 0 | 0.0382(0.0819) | 0,4672 | 6,40E-01 | 9,87E-01 | 0.0309(0.0021) | 1.0048(0.0079) | -0.0022(0.0051) |
| Illnesses of father: Heart disease | 0 | -0.0441(0.0944) | -0,467 | 6,41E-01 | 9,87E-01 | 0.0294(0.0024) | 1.0026(0.0085) | -0.0017(0.0049) |
| Diagnoses - main ICD10: D12 Benign neoplasm of colon_ rectum_ anus and anal canal | 0 | -0.0647(0.1388) | -0,4661 | 6,41E-01 | 9,87E-01 | 0.0086(0.0014) | 1.0036(0.0063) | 0.003(0.0048) |
| Non-cancer illness code_ self-reported: ankylosing spondylitis | 0 | -0.1574(0.3404) | -0,4622 | 6,44E-01 | 9,87E-01 | 0.0015(0.0014) | 1.0028(0.0064) | 0.0026(0.0047) |
| Seen doctor (GP) for nerves_ anxiety_ tension or depression | 0 | -0.0306(0.0663) | -0,4613 | 6,45E-01 | 9,87E-01 | 0.0606(0.003) | 1.0158(0.0093) | 0.0000079098(0.0054) |
| Diagnoses - main ICD10: E04 Other non-toxic goitre | 0 | 0.0749(0.1631) | 0,4592 | 6,46E-01 | 9,87E-01 | 0.0056(0.0014) | 0.9907(0.0063) | -0.0006(0.0044) |
| Total cholesterol in medium LDL | 27005778 | 0.1103(0.2403) | 0,4589 | 6,46E-01 | 9,87E-01 | 0.0652(0.0477) | 1.0132(0.0222) | -0.002(0.0049) |
| Illnesses of mother: Diabetes | 0 | -0.0468(0.1022) | -0,4577 | 6,47E-01 | 9,87E-01 | 0.0203(0.002) | 1.0159(0.0091) | 0.0035(0.0046) |
| Type of tobacco previously smoked: Cigars or pipes | 0 | 0.1026(0.2244) | 0,4575 | 6,47E-01 | 9,87E-01 | 0.0138(0.0053) | 0.9981(0.0061) | -0.0063(0.0046) |
| Total lipids in small LDL | 27005778 | 0.0979(0.2154) | 0,4543 | 6,50E-01 | 9,87E-01 | 0.0839(0.0468) | 1.0083(0.0179) | -0.0025(0.0048) |
| Medication for pain relief_ constipation_ heartburn: None of the above | 0 | 0.0313(0.069) | 0,4534 | 6,50E-01 | 9,87E-01 | 0.0539(0.0026) | 1.0311(0.009) | -0.0011(0.0052) |
| Sitting height ratio | 25865494 | 0.0553(0.1226) | 0,4515 | 6,52E-01 | 9,87E-01 | 0.217(0.0283) | 0.98(0.0075) | -0.0043(0.0051) |
| Reason for glasses/contact lenses: For short-sightedness_ i.e. only or mainly for distance viewing such as driving_ cinema etc (called myopia) | 0 | 0.0414(0.0918) | 0,4506 | 6,52E-01 | 9,87E-01 | 0.0246(0.0021) | 1.1183(0.0083) | -0.0057(0.0051) |
| Non-cancer illness code_ self-reported: diverticular disease/diverticulitis | 0 | 0.0486(0.1084) | 0,4485 | 6,54E-01 | 9,87E-01 | 0.0126(0.0016) | 0.9945(0.007) | -0.0004(0.0049) |
| Age of first birth | 27798627 | 0.0327(0.0732) | 0,4459 | 6,56E-01 | 9,87E-01 | 0.0621(0.0036) | 0.9527(0.0078) | 0.0009(0.0047) |
| Current employment status: Looking after home and/or family | 0 | 0.067(0.1504) | 0,4455 | 6,56E-01 | 9,87E-01 | 0.0062(0.0014) | 0.9998(0.0067) | 0.0013(0.0046) |
| Tinnitus: Yes_ now most or all of the time | 0 | 0.0554(0.1246) | 0,4443 | 6,57E-01 | 9,87E-01 | 0.0327(0.0046) | 1.0018(0.0067) | -0.0074(0.0047) |
| Irritability | 0 | -0.0315(0.0709) | -0,4441 | 6,57E-01 | 9,87E-01 | 0.0677(0.0042) | 0.9927(0.011) | 0.0007(0.0056) |
| Phospholipids in chylomicrons and largest VLDL particles | 27005778 | 0.0741(0.1673) | 0,4428 | 6,58E-01 | 9,87E-01 | 0.0973(0.0264) | 0.9857(0.0065) | -0.0047(0.0048) |
| Schizophrenia | 25056061 | 0.0274(0.0623) | 0,4388 | 6,61E-01 | 9,87E-01 | 0.4569(0.0195) | 1.0514(0.0133) | 0.0045(0.0065) |

| | | | | | | | | |
|---|---|---|---|---|---|---|---|---|
| Current employment status: Retired | 0 | 0.0617(0.1407) | 0,4388 | 6,61E-01 | 9,87E-01 | 0.0082(0.0015) | 1.017(0.0071) | 0.0046(0.0045) |
| Child birth length | 25281659 | 0.055(0.1259) | 0,4371 | 6,62E-01 | 9,87E-01 | 0.1708(0.0237) | 0.9928(0.0072) | -0.0025(0.005) |
| Daytime dozing / sleeping (narcolepsy) | 0 | 0.0311(0.0715) | 0,4352 | 6,63E-01 | 9,87E-01 | 0.0484(0.0027) | 1.0102(0.0085) | -0.0049(0.0048) |
| Pain type(s) experienced in last month: Facial pain | 0 | 0.0621(0.1426) | 0,4352 | 6,63E-01 | 9,87E-01 | 0.0071(0.0014) | 0.9964(0.0062) | -0.003(0.0048) |
| Frequency of depressed mood in last 2 weeks | 0 | -0.0305(0.0702) | -0,4345 | 6,64E-01 | 9,87E-01 | 0.0418(0.0027) | 1.0129(0.0081) | -0.0017(0.0049) |
| Pain type(s) experienced in last month: None of the above | 0 | 0.0303(0.07) | 0,4329 | 6,65E-01 | 9,87E-01 | 0.0543(0.0026) | 1.038(0.0083) | -0.0008(0.0052) |
| Diagnoses - main ICD10: S52 Fracture of forearm | 0 | 0.0628(0.1472) | 0,4267 | 6,70E-01 | 9,87E-01 | 0.0062(0.0018) | 0.9994(0.0069) | 0.0017(0.0046) |
| Concentration of medium LDL particles | 27005778 | 0.096(0.2262) | 0,4243 | 6,71E-01 | 9,87E-01 | 0.0773(0.05) | 1.0118(0.0203) | -0.0023(0.0047) |
| Total cholesterol in small LDL | 27005778 | 0.0952(0.2265) | 0,4203 | 6,74E-01 | 9,87E-01 | 0.07(0.0448) | 1.0163(0.0193) | -0.0011(0.0048) |
| Diagnoses - main ICD10: Z09 Follow-up examination after treatment for conditions other than malignant neoplasms | 0 | -0.0907(0.2162) | -0,4194 | 6,75E-01 | 9,87E-01 | 0.0029(0.0013) | 0.9994(0.0061) | 0.0107(0.0043) |
| Mean diameter for HDL particles | 27005778 | -0.0636(0.1523) | -0,4174 | 6,76E-01 | 9,87E-01 | 0.1213(0.0299) | 1.0084(0.0182) | -0.0036(0.0048) |
| Concentration of medium HDL particles | 27005778 | -0.0866(0.208) | -0,4165 | 6,77E-01 | 9,87E-01 | 0.062(0.0266) | 0.9953(0.0078) | 0.0059(0.0043) |
| Illnesses of mother: Alzheimers disease/dementia | 0 | -0.0911(0.2192) | -0,4157 | 6,78E-01 | 9,87E-01 | 0.0036(0.0045) | 1.0348(0.034) | -0.0002(0.0049) |
| Illnesses of siblings: None of the above (group 2) | 0 | 0.0697(0.1679) | 0,4149 | 6,78E-01 | 9,87E-01 | 0.0087(0.0018) | 1.0034(0.0071) | -0.0054(0.0044) |
| Diastolic blood pressure_ automated reading | 0 | -0.0212(0.0512) | -0,4145 | 6,79E-01 | 9,87E-01 | 0.1344(0.0058) | 1.0667(0.0155) | 0.0028(0.005) |
| Number of incorrect matches in round | 0 | 0.027(0.0653) | 0,414 | 6,79E-01 | 9,87E-01 | 0.0544(0.0027) | 1.0042(0.009) | 0.0011(0.0049) |
| Financial situation satisfaction | 0 | 0.0415(0.1006) | 0,4123 | 6,80E-01 | 9,87E-01 | 0.0569(0.0058) | 1.0058(0.0077) | -0.0079(0.0047) |
| Total cholesterol in HDL | 27005778 | 0.0651(0.158) | 0,412 | 6,80E-01 | 9,87E-01 | 0.0894(0.0295) | 1.0078(0.0128) | -0.0025(0.0046) |
| Non-cancer illness code_ self-reported: iron deficiency anaemia | 0 | 0.0781(0.1911) | 0,4088 | 6,83E-01 | 9,87E-01 | 0.0045(0.0013) | 0.9901(0.006) | -0.0035(0.0045) |
| Tense / highly strung | 0 | 0.0281(0.069) | 0,4067 | 6,84E-01 | 9,87E-01 | 0.0565(0.0032) | 0.9963(0.0096) | 0.002(0.0051) |
| Free cholesterol in large VLDL | 27005778 | -0.0636(0.1579) | -0,4024 | 6,87E-01 | 9,87E-01 | 0.125(0.0304) | 0.9896(0.0066) | -0.001(0.005) |
| Diagnoses - main ICD10: R55 Syncope and collapse | 0 | -0.074(0.1844) | -0,4016 | 6,88E-01 | 9,87E-01 | 0.0051(0.0013) | 1.0092(0.0069) | 0.0019(0.0047) |
| Frequency of stair climbing in last 4 weeks | 0 | 0.0319(0.0804) | 0,3966 | 6,92E-01 | 9,87E-01 | 0.0351(0.0021) | 1.0156(0.0078) | 0.009(0.0052) |
| Diagnoses - main ICD10: H26 Other cataract | 0 | -0.0524(0.1322) | -0,3966 | 6,92E-01 | 9,87E-01 | 0.009(0.0016) | 0.9942(0.0073) | -0.0022(0.0045) |
| Mouth/teeth dental problems: Painful gums | 0 | 0.0632(0.1599) | 0,3953 | 6,93E-01 | 9,87E-01 | 0.0065(0.0015) | 1.0053(0.0062) | 0.0006(0.0045) |
| Mean platelet volume | 22139419 | -0.0426(0.1083) | -0,3931 | 6,94E-01 | 9,87E-01 | 0.319(0.0509) | 0.9811(0.0104) | 0.001(0.0049) |
| Apolipoprotein A-I | 27005778 | -0.0728(0.187) | -0,3896 | 6,97E-01 | 9,87E-01 | 0.0658(0.0317) | 1.0064(0.0118) | 0.00006984(0.0043) |
| Cholesterol esters in large VLDL | 27005778 | -0.0562(0.1445) | -0,3893 | 6,97E-01 | 9,87E-01 | 0.1666(0.0347) | 0.9783(0.0069) | 0.0005(0.0047) |
| Total lipids in medium HDL | 27005778 | -0.082(0.2107) | -0,3892 | 6,97E-01 | 9,87E-01 | 0.0589(0.0268) | 0.9968(0.0081) | 0.0058(0.0043) |
| Phospholipids in large LDL | 27005778 | 0.0946(0.2431) | 0,3889 | 6,97E-01 | 9,87E-01 | 0.0591(0.0453) | 1.0173(0.0214) | -0.0021(0.0046) |
| Number of children fathered | 0 | -0.0468(0.1221) | -0,3831 | 7,02E-01 | 9,87E-01 | 0.0317(0.0037) | 1.0115(0.007) | -0.0019(0.005) |
| Triglycerides in very large HDL | 27005778 | -0.0647(0.1691) | -0,3825 | 7,02E-01 | 9,87E-01 | 0.0819(0.026) | 1.0147(0.0261) | -0.0034(0.0041) |
| Phospholipids in small VLDL | 27005778 | -0.0586(0.1543) | -0,38 | 7,04E-01 | 9,87E-01 | 0.1178(0.0355) | 0.9989(0.0085) | 0.0022(0.0044) |
| Concentration of large LDL particles | 27005778 | 0.0874(0.2303) | 0,3796 | 7,04E-01 | 9,87E-01 | 0.0739(0.0515) | 1.0156(0.0217) | -0.0021(0.0046) |
| Fathers age at death | 27015805 | 0.0529(0.1397) | 0,3787 | 7,05E-01 | 9,87E-01 | 0.0388(0.0069) | 1.018(0.0068) | 0.0038(0.0044) |
| Diagnoses - main ICD10: I21 Acute myocardial infarction | 0 | -0.0486(0.1288) | -0,377 | 7,06E-01 | 9,87E-01 | 0.0099(0.0016) | 1.0065(0.0072) | -0.0006(0.0048) |
| Diagnoses - main ICD10: I84 Haemorrhoids | 0 | 0.0536(0.1435) | 0,3734 | 7,09E-01 | 9,87E-01 | 0.0089(0.0016) | 1.0032(0.0074) | -0.0009(0.0046) |
| Diagnoses - main ICD10: E03 Other hypothyroidism | 0 | 0.4431(1.1927) | 0,3715 | 7,10E-01 | 9,87E-01 | 0.0004(0.0012) | 0.9985(0.006) | -0.0092(0.0046) |
| Diagnoses - main ICD10: R35 Polyuria | 0 | 0.0756(0.2053) | 0,3682 | 7,13E-01 | 9,87E-01 | 0.0033(0.0015) | 0.9966(0.0068) | -0.0051(0.0045) |
| Average weekly beer plus cider intake | 0 | -0.0269(0.0732) | -0,3673 | 7,13E-01 | 9,87E-01 | 0.051(0.0034) | 1.0635(0.0094) | -0.0072(0.0052) |
| Number of trend entries | 0 | -0.0719(0.1966) | -0,3655 | 7,15E-01 | 9,87E-01 | 0.0323(0.0098) | 1.0059(0.0068) | 0.0008(0.0049) |
| Triglycerides in large VLDL | 27005778 | 0.0585(0.1608) | 0,3638 | 7,16E-01 | 9,87E-01 | 0.1125(0.0288) | 0.99(0.0062) | -0.0017(0.0048) |
| HOMA-B | 20081858 | -0.0437(0.1207) | -0,3618 | 7,18E-01 | 9,87E-01 | 0.0851(0.0124) | 0.9897(0.0064) | -0.0024(0.0046) |
| Sleep duration | 0 | 0.0198(0.0554) | 0,3578 | 7,21E-01 | 9,87E-01 | 0.0707(0.0036) | 1.0241(0.01) | 0.0003(0.005) |
| Total lipids in small HDL | 27005778 | 0.085(0.2379) | 0,3571 | 7,21E-01 | 9,87E-01 | 0.0523(0.0272) | 1.0031(0.0074) | 0.0038(0.0048) |
| Total lipids in very large VLDL | 27005778 | 0.0562(0.1586) | 0,3544 | 7,23E-01 | 9,87E-01 | 0.1367(0.0304) | 0.9795(0.0069) | -0.004(0.0049) |
| Cholesterol esters in medium VLDL | 27005778 | -0.0519(0.1469) | -0,3534 | 7,24E-01 | 9,87E-01 | 0.1515(0.0378) | 0.987(0.008) | 0.0006(0.0045) |
| Phospholipids in large HDL | 27005778 | -0.0524(0.1489) | -0,3516 | 7,25E-01 | 9,87E-01 | 0.1275(0.0311) | 0.9988(0.0138) | -0.0006(0.0046) |
| Why stopped smoking: None of the above | 0 | -0.0435(0.1239) | -0,3509 | 7,26E-01 | 9,87E-01 | 0.0402(0.006) | 0.9976(0.0069) | -0.0029(0.0048) |
| Alcohol drinker status: Previous | 0 | -0.0399(0.1141) | -0,3501 | 7,26E-01 | 9,87E-01 | 0.0127(0.0015) | 1.0081(0.0067) | -0.0014(0.0046) |
| Age at first live birth | 0 | 0.0226(0.0649) | 0,3486 | 7,27E-01 | 9,87E-01 | 0.1648(0.0073) | 1.0477(0.0088) | 0.0018(0.0051) |
| Types of transport used (excluding work): Car/motor vehicle | 0 | 0.0311(0.0896) | 0,3466 | 7,29E-01 | 9,87E-01 | 0.0245(0.0018) | 1.0089(0.0073) | -0.0009(0.005) |
| Inflammatory Bowel Disease (Euro) | 26192919 | 0.0283(0.0818) | 0,3461 | 7,29E-01 | 9,87E-01 | 0.3331(0.0332) | 1.0498(0.0096) | -0.0014(0.0052) |
| Total lipids in large LDL | 27005778 | 0.0796(0.2313) | 0,344 | 7,31E-01 | 9,87E-01 | 0.0735(0.0526) | 1.0142(0.0223) | -0.002(0.0046) |
| Hair/balding pattern: Pattern 4 | 0 | -0.0214(0.0623) | -0,3438 | 7,31E-01 | 9,87E-01 | 0.161(0.0179) | 1.1048(0.0228) | -0.001(0.0057) |
| Free cholesterol in large LDL | 27005778 | 0.0845(0.2462) | 0,3435 | 7,31E-01 | 9,87E-01 | 0.0599(0.049) | 1.0172(0.0237) | -0.0026(0.0048) |
| Total lipids in medium LDL | 27005778 | 0.0763(0.2228) | 0,3425 | 7,32E-01 | 9,87E-01 | 0.079(0.051) | 1.0101(0.021) | -0.0018(0.0047) |
| Non-cancer illness code_ self-reported: bone disorder | 0 | 0.0953(0.2801) | 0,3404 | 7,34E-01 | 9,87E-01 | 0.0018(0.0012) | 1.0037(0.0059) | 0.0002(0.0042) |
| Number of full brothers | 0 | 0.0353(0.1046) | 0,3379 | 7,35E-01 | 9,87E-01 | 0.0189(0.0018) | 1.0273(0.0072) | -0.0017(0.005) |
| Diagnoses - main ICD10: K20 Oesophagitis | 0 | -0.1091(0.3253) | -0,3355 | 7,37E-01 | 9,87E-01 | 0.0018(0.0013) | 1.008(0.0064) | 0.0059(0.0049) |
| Total lipids in large VLDL | 27005778 | 0.0506(0.1516) | 0,3338 | 7,39E-01 | 9,87E-01 | 0.1376(0.0304) | 0.9871(0.0063) | -0.002(0.0047) |
| Urinary albumin-to-creatinine ratio (non-diabetes) | 26631737 | 0.0496(0.1492) | 0,3324 | 7,40E-01 | 9,87E-01 | 0.0516(0.0101) | 0.9959(0.0071) | -0.0021(0.0049) |
| Concentration of large HDL particles | 27005778 | -0.0481(0.145) | -0,332 | 7,40E-01 | 9,87E-01 | 0.1332(0.0311) | 0.9998(0.0148) | -0.0013(0.0046) |
| Systemic lupus erythematosus | 26502338 | 0.0362(0.1094) | 0,3312 | 7,41E-01 | 9,87E-01 | 0.3718(0.0658) | 1.1068(0.0096) | -0.0044(0.0049) |
| Hand grip strength (left) | 0 | -0.0202(0.061) | -0,3307 | 7,41E-01 | 9,87E-01 | 0.1023(0.0042) | 1.0581(0.0123) | 0.0026(0.0056) |
| Overweight | 23563607 | 0.0248(0.0752) | 0,3293 | 7,42E-01 | 9,87E-01 | 0.1092(0.0066) | 1.0243(0.0096) | -0.0068(0.0053) |
| Mineral and other dietary supplements: Calcium | 0 | 0.0344(0.1048) | 0,3288 | 7,42E-01 | 9,87E-01 | 0.0121(0.0016) | 0.9986(0.0065) | -0.0023(0.0043) |
| Number of cigarettes currently smoked daily (current cigarette smokers) | 0 | 0.0475(0.1452) | 0,3271 | 7,44E-01 | 9,87E-01 | 0.1004(0.0262) | 1.001(0.0068) | -0.0036(0.0044) |
| Phospholipids in medium HDL | 27005778 | -0.0629(0.1946) | -0,3233 | 7,46E-01 | 9,87E-01 | 0.0643(0.0236) | 0.9954(0.0079) | 0.0054(0.0044) |
| Diagnoses - main ICD10: K35 Acute appendicitis | 0 | -0.0924(0.2871) | -0,3219 | 7,48E-01 | 9,87E-01 | 0.0023(0.0013) | 1.0041(0.0069) | -0.0006(0.0044) |

| | | | | | | | | |
|---|---|---|---|---|---|---|---|---|
| Free cholesterol in medium VLDL | 27005778 | -0.0522(0.1623) | -0,3214 | 7,48E-01 | 9,87E-01 | 0.1144(0.0338) | 0.9961(0.0071) | 0.0014(0.0047) |
| Diagnoses - main ICD10: K62 Other diseases of anus and rectum | 0 | 0.0654(0.2037) | 0,3211 | 7,48E-01 | 9,87E-01 | 0.0051(0.0015) | 1.0007(0.0069) | -0.0001(0.0047) |
| Triglycerides in very large VLDL | 27005778 | 0.0508(0.1586) | 0,3204 | 7,49E-01 | 9,87E-01 | 0.1213(0.0296) | 0.9815(0.007) | -0.004(0.005) |
| Glucose | 27005778 | 0.0515(0.1616) | 0,3188 | 7,50E-01 | 9,87E-01 | 0.081(0.0186) | 0.9952(0.0067) | 0.002(0.0042) |
| Sleeplessness / insomnia | 0 | -0.0218(0.0687) | -0,3166 | 7,52E-01 | 9,87E-01 | 0.0616(0.0027) | 1.0195(0.0086) | 0.0042(0.0057) |
| Cholesterol esters in medium HDL | 27005778 | -0.0659(0.2083) | -0,3162 | 7,52E-01 | 9,87E-01 | 0.0548(0.0288) | 1.0008(0.009) | 0.0051(0.0041) |
| Heart rate | 23583979 | -0.0289(0.0915) | -0,3159 | 7,52E-01 | 9,87E-01 | 0.0852(0.0092) | 1.0116(0.0086) | 0.004(0.005) |
| Age at Menarche | 25231870 | -0.0185(0.0587) | -0,3146 | 7,53E-01 | 9,87E-01 | 0.202(0.0106) | 0.9538(0.0114) | 0.0065(0.0053) |
| Platelet count | 22139419 | 0.024(0.0765) | 0,3134 | 7,54E-01 | 9,87E-01 | 0.1191(0.0114) | 0.9839(0.0079) | 0.0034(0.0041) |
| Diagnoses - main ICD10: R11 Nausea and vomiting | 0 | 0.0999(0.3224) | 0,3099 | 7,57E-01 | 9,87E-01 | 0.0012(0.0012) | 0.9947(0.0063) | 0.0013(0.0042) |
| Hand grip strength (right) | 0 | 0.0185(0.0599) | 0,3085 | 7,58E-01 | 9,87E-01 | 0.1028(0.0043) | 1.0625(0.0136) | 0.000065549(0.0055) |
| Types of physical activity in last 4 weeks: Strenuous sports | 0 | -0.03(0.0976) | -0,3076 | 7,58E-01 | 9,87E-01 | 0.0219(0.0018) | 1.0113(0.0072) | 0.0052(0.0051) |
| Non-accidental death in close genetic family | 0 | 0.0511(0.1662) | 0,3074 | 7,59E-01 | 9,87E-01 | 0.0178(0.0045) | 0.9956(0.0064) | 0.0003(0.0049) |
| Ever vs never smoked | 20418890 | 0.0287(0.0939) | 0,3055 | 7,60E-01 | 9,87E-01 | 0.0742(0.0066) | 0.9972(0.006) | -0.0065(0.0044) |
| Albumin | 27005778 | 0.0638(0.2088) | 0,3053 | 7,60E-01 | 9,87E-01 | 0.0568(0.0252) | 0.9902(0.0066) | -0.0041(0.004) |
| Ever had prostate specific antigen (PSA) test | 0 | 0.0309(0.1018) | 0,304 | 7,61E-01 | 9,87E-01 | 0.0339(0.0035) | 1.0036(0.0073) | 0.0005(0.0045) |
| Loneliness_ isolation | 0 | 0.0256(0.0843) | 0,3036 | 7,61E-01 | 9,87E-01 | 0.0375(0.0023) | 1.0232(0.0078) | -0.0079(0.0053) |
| Total cholesterol in large LDL | 27005778 | 0.0726(0.2408) | 0,3016 | 7,63E-01 | 9,87E-01 | 0.0618(0.0474) | 1.015(0.0227) | -0.0018(0.0047) |
| Current employment status: In paid employment or self-employed | 0 | 0.0417(0.1386) | 0,3011 | 7,63E-01 | 9,87E-01 | 0.008(0.0015) | 1.0198(0.0067) | -0.0067(0.0047) |
| Phospholipids in large VLDL | 27005778 | -0.0467(0.156) | -0,2994 | 7,65E-01 | 9,87E-01 | 0.1169(0.0309) | 0.9907(0.0065) | -0.000091861(0.0047) |
| Alcohol usually taken with meals | 0 | -0.0193(0.0649) | -0,2977 | 7,66E-01 | 9,87E-01 | 0.1094(0.0051) | 1.06(0.0093) | 0.0051(0.0047) |
| Vitamin and mineral supplements: Vitamin D | 0 | 0.0509(0.1718) | 0,2961 | 7,67E-01 | 9,87E-01 | 0.006(0.0016) | 0.9966(0.0071) | -0.0016(0.0044) |
| Non-cancer illness code_ self-reported: bladder problem (not cancer) | 0 | 0.0793(0.2698) | 0,294 | 7,69E-01 | 9,87E-01 | 0.0021(0.0013) | 1.0014(0.0069) | -0.0016(0.0044) |
| Diagnoses - main ICD10: M17 Gonarthrosis [arthrosis of knee] | 0 | -0.0308(0.1052) | -0,2927 | 7,70E-01 | 9,87E-01 | 0.0139(0.0017) | 1.0067(0.0072) | -0.0009(0.0041) |
| Distance between home and job workplace | 0 | 0.0461(0.1582) | 0,2914 | 7,71E-01 | 9,87E-01 | 0.0116(0.0033) | 1.0072(0.0069) | 0.0032(0.0045) |
| Total lipids in medium VLDL | 27005778 | 0.0448(0.1545) | 0,2897 | 7,72E-01 | 9,87E-01 | 0.1394(0.0343) | 0.9872(0.0067) | -0.0008(0.0047) |
| Ratio of bisallylic groups to total fatty acids | 27005778 | 0.036(0.1261) | 0,2859 | 7,75E-01 | 9,87E-01 | 0.2689(0.0982) | 0.9856(0.0072) | -0.0005(0.0047) |
| Non-cancer illness code_ self-reported: malabsorption/coeliac disease | 0 | 0.0265(0.0939) | 0,2828 | 7,77E-01 | 9,87E-01 | 0.0171(0.0123) | 0.9941(0.0216) | -0.0027(0.0046) |
| Frequency of tenseness / restlessness in last 2 weeks | 0 | 0.0212(0.0753) | 0,281 | 7,79E-01 | 9,87E-01 | 0.043(0.0026) | 1.0162(0.009) | -0.0003(0.0056) |
| Non-cancer illness code_ self-reported: type 2 diabetes | 0 | -0.0434(0.1548) | -0,2803 | 7,79E-01 | 9,87E-01 | 0.0063(0.0014) | 1.0109(0.0065) | -0.0033(0.0045) |
| HDL cholesterol | 20686565 | 0.0232(0.0839) | 0,2769 | 7,82E-01 | 9,87E-01 | 0.1275(0.0234) | 1.0557(0.0493) | 0.0048(0.005) |
| Hip circumference | 25673412 | 0.0184(0.0665) | 0,2767 | 7,82E-01 | 9,87E-01 | 0.1256(0.0057) | 0.8622(0.0081) | -0.007(0.005) |
| Serum total triglycerides | 27005778 | -0.0409(0.1515) | -0,2702 | 7,87E-01 | 9,87E-01 | 0.127(0.0361) | 0.9927(0.0075) | 0.0009(0.0046) |
| Total lipids in large HDL | 27005778 | -0.0397(0.1469) | -0,2699 | 7,87E-01 | 9,87E-01 | 0.1309(0.0314) | 1.001(0.0146) | -0.0019(0.0046) |
| Chest pain or discomfort | 0 | -0.0221(0.0824) | -0,2688 | 7,88E-01 | 9,87E-01 | 0.0328(0.0021) | 1.0178(0.0078) | 0.0026(0.0051) |
| Cholesterol esters in medium LDL | 27005778 | 0.0605(0.2254) | 0,2684 | 7,88E-01 | 9,87E-01 | 0.0779(0.0507) | 1.0102(0.021) | -0.0013(0.0048) |
| Cholesterol esters in large LDL | 27005778 | 0.0604(0.2284) | 0,2643 | 7,92E-01 | 9,87E-01 | 0.0748(0.052) | 1.0121(0.022) | -0.0016(0.0047) |
| Loud music exposure frequency | 0 | -0.0388(0.1471) | -0,2635 | 7,92E-01 | 9,87E-01 | 0.0289(0.0046) | 1.0127(0.0069) | 0.0009(0.0051) |
| Height_2010 | 20881960 | -0.0144(0.0558) | -0,2579 | 7,97E-01 | 9,87E-01 | 0.2842(0.0161) | 1.0186(0.0181) | 0.0067(0.0058) |
| Total lipids in very large HDL | 27005778 | 0.0551(0.2144) | 0,2573 | 7,97E-01 | 9,87E-01 | 0.0532(0.0272) | 1.0177(0.0145) | -0.0069(0.0046) |
| Non-cancer illness code_ self-reported: retinal detachment | 0 | -0.0477(0.1854) | -0,2572 | 7,97E-01 | 9,87E-01 | 0.0037(0.0013) | 1.0023(0.0065) | 0.0012(0.004) |
| Non-cancer illness code_ self-reported: chronic obstructive airways disease/copd | 0 | -0.0421(0.1699) | -0,2481 | 8,04E-01 | 9,87E-01 | 0.0057(0.0015) | 0.9933(0.007) | -0.0017(0.0047) |
| Concentration of very large VLDL particles | 27005778 | 0.0375(0.1531) | 0,2449 | 8,07E-01 | 9,87E-01 | 0.1241(0.0296) | 0.9841(0.0068) | -0.0002(0.0043) |
| Ever had stillbirth_ spontaneous miscarriage or termination | 0 | 0.0336(0.1373) | 0,2444 | 8,07E-01 | 9,87E-01 | 0.0193(0.0032) | 0.996(0.0077) | -0.004(0.0047) |
| Medication for pain relief_ constipation_ heartburn: Omeprazole (e.g. Zanprol) | 0 | 0.0269(0.1105) | 0,2431 | 8,08E-01 | 9,87E-01 | 0.018(0.0017) | 1.02(0.0071) | 0.0017(0.005) |
| Illnesses of siblings: Stroke | 0 | 0.0569(0.2347) | 0,2425 | 8,08E-01 | 9,87E-01 | 0.0044(0.0018) | 1.0078(0.0066) | -0.0026(0.0049) |
| Diagnoses - main ICD10: C61 Malignant neoplasm of prostate | 0 | -0.0442(0.1824) | -0,2424 | 8,09E-01 | 9,87E-01 | 0.0043(0.0015) | 1.0177(0.0068) | -0.0024(0.0046) |
| Triglycerides in medium VLDL | 27005778 | 0.0412(0.1722) | 0,2391 | 8,11E-01 | 9,87E-01 | 0.0973(0.0304) | 1.0002(0.0064) | 0.0002(0.0047) |
| Illnesses of father: None of the above (group 2) | 0 | -0.0375(0.1569) | -0,2388 | 8,11E-01 | 9,87E-01 | 0.0092(0.0017) | 1.0015(0.0072) | -0.0016(0.005) |
| Serum total cholesterol | 27005778 | -0.0518(0.2176) | -0,2379 | 8,12E-01 | 9,87E-01 | 0.0626(0.0385) | 1.0225(0.0154) | -0.0003(0.0046) |
| Pain type(s) experienced in last month: Back pain | 0 | 0.0187(0.0791) | 0,2363 | 8,13E-01 | 9,87E-01 | 0.0391(0.0023) | 1.0249(0.009) | 0.0026(0.0057) |
| Hair/balding pattern: Pattern 2 | 0 | -0.0211(0.0897) | -0,2357 | 8,14E-01 | 9,87E-01 | 0.051(0.0062) | 1.0351(0.013) | -0.00007992(0.0051) |
| Overall health rating | 0 | -0.0149(0.0637) | -0,2333 | 8,16E-01 | 9,87E-01 | 0.0992(0.0032) | 1.0624(0.0105) | -0.0015(0.0064) |
| Total lipids in chylomicrons and largest VLDL particles | 27005778 | 0.038(0.163) | 0,2332 | 8,16E-01 | 9,87E-01 | 0.1198(0.0275) | 0.9909(0.0066) | -0.0049(0.0048) |
| Lung cancer | 27488534 | -0.0245(0.1055) | -0,232 | 8,17E-01 | 9,87E-01 | 0.3125(0.0664) | 1.0143(0.0071) | -0.0009(0.0044) |
| Non-cancer illness code_ self-reported: arthritis (nos) | 0 | -0.066(0.2883) | -0,2288 | 8,19E-01 | 9,87E-01 | 0.002(0.0014) | 1.014(0.0066) | -0.0034(0.0045) |
| Concentration of IDL particles | 27005778 | 0.0506(0.2212) | 0,2286 | 8,19E-01 | 9,87E-01 | 0.0746(0.044) | 1.0175(0.0182) | -0.0016(0.0044) |
| Total Cholesterol | 20686565 | -0.0192(0.0847) | -0,2268 | 8,21E-01 | 9,87E-01 | 0.1328(0.0249) | 1.0211(0.035) | -0.0027(0.0053) |
| Triglycerides in IDL | 27005778 | 0.04(0.1773) | 0,2259 | 8,21E-01 | 9,87E-01 | 0.1084(0.0353) | 1.01(0.0148) | -0.0004(0.0042) |
| Crohns disease | 26192919 | -0.0186(0.0829) | -0,2241 | 8,23E-01 | 9,87E-01 | 0.4918(0.0582) | 1.021(0.0105) | 0.0013(0.0048) |
| Qualifications: College or University degree | 0 | 0.0118(0.0528) | 0,224 | 8,23E-01 | 9,87E-01 | 0.167(0.0053) | 1.1195(0.0136) | 0.0044(0.0065) |
| Wears glasses or contact lenses | 0 | 0.0212(0.0959) | 0,2213 | 8,25E-01 | 9,87E-01 | 0.016(0.0017) | 1.0008(0.0075) | -0.0047(0.005) |
| Age completed full time education | 0 | 0.0146(0.0662) | 0,2204 | 8,26E-01 | 9,87E-01 | 0.0835(0.0044) | 1.0638(0.0093) | -0.005(0.0048) |
| Phospholipids in very large HDL | 27005778 | 0.0388(0.1766) | 0,2199 | 8,26E-01 | 9,87E-01 | 0.0895(0.0293) | 1.0086(0.0162) | -0.0072(0.0048) |
| Mineral and other dietary supplements: Glucosamine | 0 | -0.0197(0.0898) | -0,219 | 8,27E-01 | 9,87E-01 | 0.0241(0.002) | 1.0087(0.0075) | 0.0051(0.0047) |
| Adiponectin | 22479202 | -0.0311(0.1423) | -0,2187 | 8,27E-01 | 9,87E-01 | 0.1179(0.0264) | 1.0206(0.0112) | 0.0023(0.0052) |
| Mean time to correctly identify matches | 0 | -0.014(0.0656) | -0,2131 | 8,31E-01 | 9,87E-01 | 0.0733(0.003) | 1.0294(0.0093) | 0.001(0.0053) |
| Doctor diagnosed asthma | 0 | 0.0223(0.105) | 0,2124 | 8,32E-01 | 9,87E-01 | 0.0684(0.0098) | 0.9985(0.008) | 0.0038(0.005) |
| Job involves heavy manual or physical work | 0 | -0.0154(0.0735) | -0,2098 | 8,34E-01 | 9,87E-01 | 0.0863(0.0045) | 1.0366(0.0089) | 0.004(0.0049) |
| Total lipids in IDL | 27005778 | 0.0466(0.2239) | 0,208 | 8,35E-01 | 9,87E-01 | 0.0732(0.0455) | 1.0183(0.019) | -0.0015(0.0045) |

| | | | | | | | | |
|---|---|---|---|---|---|---|---|---|
| Non-cancer illness code_ self-reported: glaucoma | 0 | -0.0265(0.1273) | -0,2078 | 8,35E-01 | 9,87E-01 | 0.0109(0.0017) | 1.0144(0.0079) | 0.004(0.005) |
| Pain type(s) experienced in last month: Headache | 0 | 0.0149(0.0721) | 0,2072 | 8,36E-01 | 9,87E-01 | 0.0441(0.0027) | 1.0061(0.0093) | -0.0058(0.005) |
| Phospholipids in medium VLDL | 27005778 | -0.0337(0.1629) | -0,2069 | 8,36E-01 | 9,87E-01 | 0.1134(0.0337) | 0.9974(0.007) | 0.0011(0.0046) |
| Illnesses of mother: High blood pressure | 0 | -0.0173(0.0837) | -0,2067 | 8,36E-01 | 9,87E-01 | 0.0294(0.0022) | 1.0166(0.0086) | -0.0011(0.0052) |
| Lumbar spine bone mineral density | 22504420 | 0.0158(0.0789) | 0,1997 | 8,42E-01 | 9,87E-01 | 0.2615(0.0254) | 1.0145(0.0088) | -0.0004(0.0044) |
| Miserableness | 0 | -0.0127(0.0638) | -0,1993 | 8,42E-01 | 9,87E-01 | 0.0633(0.0035) | 1.0193(0.0107) | -0.0003(0.005) |
| Serumurate overweight | 25811787 | 0.0227(0.114) | 0,1991 | 8,42E-01 | 9,87E-01 | 0.4824(0.2704) | 0.9747(0.0235) | 0.0001(0.0043) |
| Transferrin | 25352340 | 0.0316(0.1585) | 0,1991 | 8,42E-01 | 9,87E-01 | 0.1303(0.065) | 1.0692(0.0205) | 0.001(0.0064) |
| Family relationship satisfaction | 0 | -0.0204(0.1075) | -0,1895 | 8,50E-01 | 9,87E-01 | 0.0585(0.0051) | 1.0031(0.0075) | 0.0071(0.0048) |
| Phenylalanine | 27005778 | 0.0368(0.1956) | 0,1883 | 8,51E-01 | 9,87E-01 | 0.0485(0.0219) | 1.0057(0.0068) | 0.0037(0.0041) |
| Transport type for commuting to job workplace: Public transport | 0 | 0.02(0.1072) | 0,1865 | 8,52E-01 | 9,87E-01 | 0.0309(0.0034) | 1.0248(0.0074) | 0.0002(0.0044) |
| Squamous cell lung cancer | 27488534 | 0.0281(0.1527) | 0,184 | 8,54E-01 | 9,87E-01 | 0.0405(0.0118) | 1.0084(0.0065) | -0.0019(0.0043) |
| Peak expiratory flow (PEF) | 0 | 0.0132(0.0719) | 0,1837 | 8,54E-01 | 9,87E-01 | 0.0889(0.0047) | 1.0572(0.0121) | -0.003(0.0061) |
| Non-cancer illness code_ self-reported: kidney stone/ureter stone/bladder stone | 0 | -0.0282(0.1539) | -0,183 | 8,55E-01 | 9,87E-01 | 0.0089(0.0016) | 1.002(0.0074) | -0.0052(0.0045) |
| Concentration of large VLDL particles | 27005778 | -0.0284(0.1579) | -0,1797 | 8,57E-01 | 9,87E-01 | 0.1311(0.0351) | 0.9758(0.0068) | -0.0002(0.0046) |
| Why reduced smoking: Illness or ill health | 0 | -0.0385(0.2158) | -0,1785 | 8,58E-01 | 9,87E-01 | 0.112(0.0409) | 0.9978(0.0061) | 0.0052(0.0048) |
| Health satisfaction | 0 | 0.0168(0.0952) | 0,1766 | 8,60E-01 | 9,87E-01 | 0.0829(0.0056) | 1.0058(0.0074) | -0.003(0.005) |
| Duration of other exercises | 0 | -0.0205(0.1166) | -0,176 | 8,60E-01 | 9,87E-01 | 0.019(0.0031) | 1.0095(0.0071) | -0.0069(0.0041) |
| Forced expiratory volume in 1-second (FEV1)_ predicted | 0 | -0.0126(0.0748) | -0,1688 | 8,66E-01 | 9,87E-01 | 0.1721(0.0124) | 1.0391(0.0117) | -0.0011(0.0048) |
| Diagnoses - main ICD10: J22 Unspecified acute lower respiratory infection | 0 | -0.0392(0.2358) | -0,1662 | 8,68E-01 | 9,87E-01 | 0.0026(0.0014) | 0.9996(0.0067) | 0.0006(0.0041) |
| Age at Menopause | 26414677 | -0.0153(0.0922) | -0,166 | 8,68E-01 | 9,87E-01 | 0.1331(0.0159) | 0.9906(0.0139) | -0.0013(0.0052) |
| Free cholesterol in medium HDL | 27005778 | -0.0332(0.201) | -0,1654 | 8,69E-01 | 9,87E-01 | 0.0636(0.0234) | 0.9983(0.0084) | 0.0046(0.0045) |
| Illnesses of siblings: High blood pressure | 0 | -0.0135(0.082) | -0,1644 | 8,69E-01 | 9,87E-01 | 0.0363(0.0027) | 1.0067(0.0079) | -0.0008(0.0049) |
| Lumbar Spine bone mineral density | 26367794 | -0.0172(0.1054) | -0,1633 | 8,70E-01 | 9,87E-01 | 0.1299(0.0148) | 0.9766(0.0079) | 0.0004(0.0046) |
| Chronotype | 27494321 | 0.0126(0.0778) | 0,1616 | 8,72E-01 | 9,87E-01 | 0.102(0.0057) | 1.0125(0.0075) | 0.001(0.0054) |
| Total cholesterol in IDL | 27005778 | 0.0366(0.2282) | 0,1606 | 8,72E-01 | 9,87E-01 | 0.071(0.0468) | 1.0179(0.02) | -0.0016(0.0046) |
| Mothers age at death | 27015805 | -0.0235(0.152) | -0,1545 | 8,77E-01 | 9,87E-01 | 0.0361(0.0072) | 1.0119(0.0069) | 0.0008(0.0045) |
| Concentration of medium VLDL particles | 27005778 | -0.0223(0.1448) | -0,1536 | 8,78E-01 | 9,87E-01 | 0.1506(0.0357) | 0.9819(0.0067) | 0.0008(0.0047) |
| Apolipoprotein B | 27005778 | 0.0305(0.2049) | 0,1488 | 8,82E-01 | 9,87E-01 | 0.0825(0.0405) | 1.0088(0.0149) | 0.0008(0.0047) |
| Insomnia | 28604731 | 0.0161(0.1085) | 0,1487 | 8,82E-01 | 9,87E-01 | 0.0471(0.0047) | 1.0033(0.0068) | 0.0024(0.0043) |
| Diagnoses - main ICD10: M70 Soft tissue disorders related to use_ overuse and pressure | 0 | 0.0287(0.1931) | 0,1487 | 8,82E-01 | 9,87E-01 | 0.004(0.0013) | 0.9822(0.0062) | 0.000094498(0.0049) |
| Bring up phlegm/sputum/mucus on most days | 0 | 0.0246(0.166) | 0,1485 | 8,82E-01 | 9,87E-01 | 0.026(0.0063) | 1.008(0.0074) | -0.0009(0.0049) |
| Illnesses of siblings: Diabetes | 0 | -0.0151(0.1026) | -0,1472 | 8,83E-01 | 9,87E-01 | 0.0202(0.002) | 1.0096(0.0066) | -0.0052(0.0049) |
| Diagnoses - main ICD10: K60 Fissure and fistula of anal and rectal regions | 0 | 0.0253(0.1729) | 0,1461 | 8,84E-01 | 9,87E-01 | 0.0055(0.0014) | 0.9821(0.0064) | 0.0094(0.0045) |
| Total cholesterol in medium HDL | 27005778 | -0.0294(0.204) | -0,144 | 8,86E-01 | 9,87E-01 | 0.0544(0.025) | 0.9992(0.0089) | 0.0044(0.0042) |
| Femoral Neck bone mineral density | 26367794 | 0.0137(0.0968) | 0,1413 | 8,88E-01 | 9,87E-01 | 0.12(0.0145) | 0.9773(0.0079) | 0.0026(0.0045) |
| Non-cancer illness code_ self-reported: joint disorder | 0 | -0.0356(0.2559) | -0,1391 | 8,89E-01 | 9,87E-01 | 0.0023(0.0013) | 0.9905(0.0063) | 0.0029(0.0043) |
| Lung cancer (squamous cell) | 24880342 | 0.0259(0.1901) | 0,1362 | 8,92E-01 | 9,87E-01 | 0.0506(0.0196) | 1.0053(0.0066) | -0.0018(0.0048) |
| Sitting height | 0 | -0.0068(0.0502) | -0,1347 | 8,93E-01 | 9,87E-01 | 0.32(0.0185) | 1.1378(0.0424) | 0.0021(0.0066) |
| 2hr glucose adjusted for BMI | 20081857 | -0.0273(0.2031) | -0,1343 | 8,93E-01 | 9,87E-01 | 0.0869(0.0344) | 0.9953(0.0069) | 0.0019(0.0046) |
| Illness_ injury_ bereavement_ stress in last 2 years: None of the above | 0 | 0.0127(0.0947) | 0,1344 | 8,93E-01 | 9,87E-01 | 0.0225(0.0017) | 1.0132(0.0062) | 0.0063(0.0048) |
| Diagnoses - main ICD10: K57 Diverticular disease of intestine | 0 | -0.0147(0.1101) | -0,1334 | 8,94E-01 | 9,87E-01 | 0.0132(0.0015) | 1.0032(0.0066) | 0.0034(0.0046) |
| Diagnoses - main ICD10: N32 Other disorders of bladder | 0 | -0.0266(0.2006) | -0,1324 | 8,95E-01 | 9,87E-01 | 0.0041(0.0012) | 0.9895(0.0062) | 0.0055(0.0046) |
| Medication for pain relief_ constipation_ heartburn: Paracetamol | 0 | 0.0103(0.0796) | 0,1287 | 8,98E-01 | 9,87E-01 | 0.0426(0.0024) | 1.0276(0.0085) | -0.0012(0.0054) |
| Pain type(s) experienced in last month: Neck or shoulder pain | 0 | -0.011(0.0859) | -0,1282 | 8,98E-01 | 9,87E-01 | 0.0325(0.002) | 1.0253(0.0076) | 0.0017(0.0051) |
| Pulse rate_ automated reading | 0 | -0.0072(0.0567) | -0,1275 | 8,99E-01 | 9,87E-01 | 0.1532(0.0108) | 1.0487(0.0282) | 0.0016(0.0062) |
| Triglycerides in very small VLDL | 27005778 | 0.0199(0.1566) | 0,127 | 8,99E-01 | 9,87E-01 | 0.139(0.0372) | 0.9989(0.0097) | 0.0003(0.0044) |
| Mono-unsaturated fatty acids | 27005778 | 0.0233(0.1837) | 0,1269 | 8,99E-01 | 9,87E-01 | 0.1044(0.0422) | 0.9884(0.0078) | -0.0013(0.0044) |
| Hearing difficulty/problems with background noise | 0 | 0.0104(0.0826) | 0,1264 | 9,00E-01 | 9,87E-01 | 0.0511(0.0026) | 1.0213(0.0084) | -0.0037(0.0059) |
| Obesity class 1 | 23563607 | 0.0087(0.0704) | 0,1242 | 9,01E-01 | 9,87E-01 | 0.2153(0.0115) | 1.019(0.0112) | -0.0058(0.006) |
| Free cholesterol | 27005778 | 0.0318(0.2564) | 0,1239 | 9,01E-01 | 9,87E-01 | 0.0611(0.0433) | 1.0235(0.0108) | -0.0027(0.0045) |
| Phospholipids in very large VLDL | 27005778 | 0.0203(0.164) | 0,1238 | 9,02E-01 | 9,87E-01 | 0.1089(0.0284) | 0.986(0.0066) | -0.004(0.005) |
| Sensitivity / hurt feelings | 0 | 0.0077(0.0661) | 0,1169 | 9,07E-01 | 9,87E-01 | 0.0615(0.0031) | 1.0229(0.009) | -0.0022(0.0056) |
| Pack years adult smoking as proportion of life span exposed to smoking PREVIEW ONLY | 0 | -0.009(0.0771) | -0,1169 | 9,07E-01 | 9,87E-01 | 0.1256(0.0113) | 1.0157(0.0099) | -0.0007(0.0051) |
| Insomnia | 27992416 | 0.0107(0.0919) | 0,1165 | 9,07E-01 | 9,87E-01 | 0.1312(0.0112) | 1.0038(0.0078) | 0.0033(0.0045) |
| Cigarettes smoked per day | 20418890 | 0.0181(0.1572) | 0,115 | 9,09E-01 | 9,87E-01 | 0.0582(0.0155) | 1.0058(0.0061) | -0.0018(0.0044) |
| Cholesterol esters in large HDL | 27005778 | -0.0172(0.1515) | -0,1134 | 9,10E-01 | 9,87E-01 | 0.1232(0.031) | 1.0011(0.014) | -0.003(0.0048) |
| Non-cancer illness code_ self-reported: polio / poliomyelitis | 0 | 0.0258(0.2289) | 0,1127 | 9,10E-01 | 9,87E-01 | 0.0026(0.0013) | 0.9932(0.0065) | 0.0029(0.0043) |
| Non-cancer illness code_ self-reported: hiatus hernia | 0 | 0.0178(0.1606) | 0,1109 | 9,12E-01 | 9,87E-01 | 0.008(0.0014) | 1.034(0.0067) | 0.0026(0.0047) |
| Free cholesterol in large HDL | 27005778 | -0.017(0.1539) | -0,1108 | 9,12E-01 | 9,87E-01 | 0.1051(0.029) | 1.0091(0.0139) | -0.0034(0.0046) |
| Primary biliary cirrhosis | 26394269 | -0.0128(0.1177) | -0,109 | 9,13E-01 | 9,87E-01 | 0.3602(0.0618) | 1.0053(0.0097) | -0.0031(0.0052) |
| Number of cigarettes previously smoked daily | 0 | -0.0089(0.0845) | -0,1048 | 9,17E-01 | 9,87E-01 | 0.0984(0.014) | 1.0033(0.0089) | 0.0012(0.0047) |
| Mothers age at death | 0 | -0.0121(0.1167) | -0,104 | 9,17E-01 | 9,87E-01 | 0.0242(0.0031) | 1.0283(0.0077) | -0.0019(0.005) |
| Diagnoses - main ICD10: K30 Dyspepsia | 0 | 0.0226(0.2176) | 0,1038 | 9,17E-01 | 9,87E-01 | 0.0031(0.0014) | 1.0035(0.0065) | 0.0013(0.0043) |
| Eye problems/disorders: Glaucoma | 0 | 0.0121(0.1173) | 0,1035 | 9,18E-01 | 9,87E-01 | 0.0382(0.0056) | 1.0188(0.0079) | -0.0004(0.005) |
| Mean Caudate | 25607358 | -0.0139(0.1367) | -0,1017 | 9,19E-01 | 9,87E-01 | 0.2542(0.0396) | 0.9675(0.0061) | 0.0008(0.0047) |
| Frequency of unenthusiasm / disinterest in last 2 weeks | 0 | 0.008(0.0793) | 0,1009 | 9,20E-01 | 9,87E-01 | 0.0386(0.0026) | 1.0177(0.0085) | -0.0018(0.0055) |
| Fluid intelligence score | 0 | 0.006(0.0591) | 0,1008 | 9,20E-01 | 9,87E-01 | 0.2372(0.0109) | 1.0428(0.0104) | -0.0037(0.0056) |
| Extreme waist-to-hip ratio | 23563607 | 0.015(0.149) | 0,1007 | 9,20E-01 | 9,87E-01 | 0.3646(0.0612) | 0.9758(0.0084) | 0.0112(0.006) |
| Phospholipids in very small VLDL | 27005778 | -0.0195(0.1957) | -0,0999 | 9,21E-01 | 9,87E-01 | 0.0864(0.0384) | 1.0136(0.0153) | 0.000087857(0.0044) |

| | | | | | | | | |
|---|---|---|---|---|---|---|---|---|
| Triglycerides in chylomicrons and largest VLDL particles | 27005778 | -0.0168(0.1753) | -0,0959 | 9,24E-01 | 9,87E-01 | 0.0913(0.0272) | 0.9913(0.0065) | -0.0021(0.0048) |
| Reason for glasses/contact lenses: For astigmatism | 0 | -0.0133(0.1394) | -0,0956 | 9,24E-01 | 9,87E-01 | 0.0089(0.0015) | 1.0264(0.0067) | -0.0022(0.005) |
| No-wear time bias adjusted acceleration standard deviation | 0 | -0.0092(0.0968) | -0,0955 | 9,24E-01 | 9,87E-01 | 0.0867(0.0079) | 1.003(0.0074) | 0.0018(0.0046) |
| Mean Putamen | 25607358 | 0.0114(0.1254) | 0,0911 | 9,27E-01 | 9,87E-01 | 0.2912(0.0474) | 0.9501(0.0072) | -0.0021(0.0049) |
| Maximum workload during fitness test | 0 | 0.0159(0.1759) | 0,0902 | 9,28E-01 | 9,87E-01 | 0.042(0.0104) | 1.0082(0.0072) | 0.0025(0.0052) |
| Mouth/teeth dental problems: Loose teeth | 0 | -0.0102(0.1138) | -0,09 | 9,28E-01 | 9,87E-01 | 0.0146(0.0017) | 1.0099(0.007) | -0.0032(0.0051) |
| Non-cancer illness code_ self-reported: osteoarthritis | 0 | -0.0087(0.0987) | -0,0877 | 9,30E-01 | 9,87E-01 | 0.0189(0.0017) | 1.0269(0.0073) | -0.0038(0.0044) |
| Total cholesterol in large HDL | 27005778 | -0.0135(0.1567) | -0,0864 | 9,31E-01 | 9,87E-01 | 0.1046(0.029) | 1.007(0.0142) | -0.0031(0.0047) |
| Mean Accumbens | 25607358 | 0.018(0.2082) | 0,0863 | 9,31E-01 | 9,87E-01 | 0.0891(0.0371) | 0.9784(0.0068) | -0.0025(0.0045) |
| Length of working week for main job | 0 | -0.0115(0.133) | -0,0861 | 9,31E-01 | 9,87E-01 | 0.0193(0.0029) | 1.009(0.0065) | 0.0067(0.0052) |
| Type 2 Diabetes | 22885922 | -0.0081(0.0944) | -0,0854 | 9,32E-01 | 9,87E-01 | 0.0863(0.0091) | 1.0075(0.0069) | 0.000031261(0.0043) |
| 18:2 linoleic acid (LA) | 27005778 | 0.015(0.1797) | 0,0837 | 9,33E-01 | 9,87E-01 | 0.1341(0.0498) | 1.0015(0.0101) | -0.0019(0.0045) |
| Ratio of bisallylic groups to double bonds | 27005778 | -0.009(0.1164) | -0,077 | 9,39E-01 | 9,91E-01 | 0.2995(0.1094) | 0.9759(0.0078) | 0.0002(0.0048) |
| Non-cancer illness code_ self-reported: vaginal prolapse/uterine prolapse | 0 | -0.0136(0.1795) | -0,0758 | 9,40E-01 | 9,91E-01 | 0.0045(0.0014) | 1.0066(0.0066) | -0.0014(0.0045) |
| Started insulin within one year diagnosis of diabetes | 0 | -0.0111(0.1499) | -0,0739 | 9,41E-01 | 9,91E-01 | 0.1487(0.045) | 1.0086(0.008) | 0.0063(0.005) |
| Most recent bowel cancer screening | 0 | 0.0154(0.2339) | 0,0657 | 9,48E-01 | 9,93E-01 | 0.0126(0.006) | 1.0108(0.0062) | -0.0052(0.0043) |
| Vitamin and mineral supplements: Vitamin A | 0 | 0.0146(0.222) | 0,0657 | 9,48E-01 | 9,93E-01 | 0.0036(0.0015) | 1.0031(0.0062) | 0.0092(0.0047) |
| College completion | 23722424 | 0.0057(0.0879) | 0,0654 | 9,48E-01 | 9,93E-01 | 0.0786(0.0059) | 1.0213(0.0087) | 0.0071(0.0053) |
| Handedness (chirality/laterality): Left-handed | 0 | 0.0086(0.1323) | 0,0648 | 9,48E-01 | 9,93E-01 | 0.0097(0.0016) | 1.0046(0.007) | 0.0021(0.0046) |
| Excessive daytime sleepiness | 27992416 | -0.006(0.0977) | -0,0609 | 9,51E-01 | 9,93E-01 | 0.0547(0.005) | 1.002(0.0066) | -0.0034(0.0046) |
| Total cholesterol in small VLDL | 27005778 | -0.0104(0.1744) | -0,0598 | 9,52E-01 | 9,93E-01 | 0.092(0.0322) | 1.0055(0.0098) | 0.0005(0.0045) |
| Eye problems/disorders: Injury or trauma resulting in loss of vision | 0 | -0.0322(0.5487) | -0,0588 | 9,53E-01 | 9,93E-01 | 0.0015(0.004) | 1.0096(0.0062) | 0.0033(0.0041) |
| Shortness of breath walking on level ground | 0 | 0.0069(0.1192) | 0,0575 | 9,54E-01 | 9,93E-01 | 0.0488(0.0052) | 1.0071(0.007) | -0.007(0.0052) |
| Neo-conscientiousness | 21173776 | 0.0122(0.2152) | 0,0568 | 9,55E-01 | 9,93E-01 | 0.0704(0.0309) | 1.0011(0.007) | -0.0032(0.0047) |
| Acetate | 27005778 | 0.0111(0.1979) | 0,0558 | 9,56E-01 | 9,93E-01 | 0.0507(0.0195) | 1.0019(0.0069) | 0.0044(0.004) |
| Concentration of small VLDL particles | 27005778 | 0.0066(0.1485) | 0,0446 | 9,64E-01 | 9,97E-01 | 0.1512(0.0375) | 0.9912(0.0079) | 0.0003(0.0046) |
| Diagnoses - main ICD10: Z80 Family history of malignant neoplasm | 0 | -0.0149(0.3374) | -0,044 | 9,65E-01 | 9,97E-01 | 0.0011(0.0014) | 0.9905(0.0059) | 0.0029(0.0046) |
| Mean diameter for VLDL particles | 27005778 | -0.0066(0.151) | -0,0438 | 9,65E-01 | 9,97E-01 | 0.1362(0.0343) | 0.9907(0.0081) | -0.0002(0.0048) |
| Free cholesterol in IDL | 27005778 | -0.0087(0.2127) | -0,0411 | 9,67E-01 | 9,97E-01 | 0.0693(0.0419) | 1.0195(0.0199) | -0.0012(0.0044) |
| Autism spectrum disorder | 0 | -0.0042(0.1102) | -0,0384 | 9,69E-01 | 9,97E-01 | 0.4662(0.0525) | 0.9639(0.0068) | -0.0012(0.0046) |
| Pack years of smoking PREVIEW ONLY | 0 | -0.0031(0.0822) | -0,0383 | 9,70E-01 | 9,97E-01 | 0.1082(0.01) | 1.0164(0.0094) | -0.0008(0.005) |
| Cancer code_ self-reported: squamous cell carcinoma | 0 | 0.0084(0.2491) | 0,0336 | 9,73E-01 | 9,97E-01 | 0.0023(0.0013) | 0.9994(0.0068) | -0.0025(0.0047) |
| Concentration of very small VLDL particles | 27005778 | 0.0053(0.1608) | 0,0328 | 9,74E-01 | 9,97E-01 | 0.1182(0.0364) | 1.0048(0.0121) | 0.0003(0.004) |
| Concentration of chylomicrons and largest VLDL particles | 27005778 | -0.0053(0.1643) | -0,0325 | 9,74E-01 | 9,97E-01 | 0.1157(0.0262) | 0.9888(0.0064) | -0.0011(0.0047) |
| Drive faster than motorway speed limit | 0 | 0.0023(0.0746) | 0,0303 | 9,76E-01 | 9,97E-01 | 0.0569(0.0028) | 1.0282(0.0088) | -0.0007(0.0053) |
| Diagnoses - main ICD10: G47 Sleep disorders | 0 | 0.0053(0.1752) | 0,0302 | 9,76E-01 | 9,97E-01 | 0.006(0.0015) | 0.9954(0.0073) | -0.0046(0.0053) |
| HOMA-IR | 20081858 | -0.0036(0.1399) | -0,0261 | 9,79E-01 | 9,97E-01 | 0.066(0.0123) | 1.0033(0.0064) | -0.0025(0.0048) |
| Omega-9 and saturated fatty acids | 27005778 | -0.0048(0.2112) | -0,0228 | 9,82E-01 | 9,97E-01 | 0.0865(0.0424) | 0.9962(0.0078) | -0.0013(0.0046) |
| Number of depression episodes | 0 | 0.005(0.2195) | 0,0226 | 9,82E-01 | 9,97E-01 | 0.0305(0.0119) | 1.0055(0.0063) | -0.002(0.005) |
| Fed-up feelings | 0 | 0.0014(0.069) | 0,0197 | 9,84E-01 | 9,97E-01 | 0.0713(0.0033) | 1.025(0.0101) | -0.0047(0.006) |
| Phospholipids in IDL | 27005778 | 0.0042(0.2292) | 0,0185 | 9,85E-01 | 9,97E-01 | 0.0598(0.0421) | 1.0204(0.0194) | -0.0004(0.0044) |
| Free cholesterol in small VLDL | 27005778 | 0.0028(0.1608) | 0,0171 | 9,86E-01 | 9,97E-01 | 0.1146(0.0343) | 1.0001(0.0087) | 0.0008(0.0045) |
| Qualifications: Other professional qualifications eg: nursing_ teaching | 0 | -0.0012(0.071) | -0,0172 | 9,86E-01 | 9,97E-01 | 0.0469(0.0025) | 1.0271(0.0091) | 0.0004(0.0049) |
| Total lipids in very small VLDL | 27005778 | 0.0027(0.1697) | 0,0162 | 9,87E-01 | 9,97E-01 | 0.1094(0.0369) | 1.0089(0.0127) | -0.0001(0.0042) |
| Triglycerides in small VLDL | 27005778 | -0.0023(0.1588) | -0,0144 | 9,89E-01 | 9,97E-01 | 0.1215(0.0355) | 0.9948(0.0073) | 0.002(0.0048) |
| Body mass index | 20935630 | -0.0009(0.0701) | -0,0134 | 9,89E-01 | 9,97E-01 | 0.1877(0.0096) | 1.0115(0.0107) | -0.0064(0.006) |
| Vitamin and mineral supplements: Vitamin E | 0 | 0.0019(0.1595) | 0,0121 | 9,90E-01 | 9,97E-01 | 0.0063(0.0014) | 0.9965(0.0064) | 0.0046(0.0047) |
| Fractured bone site(s): Arm | 0 | -0.0019(0.1683) | -0,0115 | 9,91E-01 | 9,97E-01 | 0.0053(0.0015) | 0.9965(0.0067) | -0.0072(0.0046) |
| Non-cancer illness code_ self-reported: hypothyroidism/myxoedema | 0 | 0.0007(0.0671) | 0,0102 | 9,92E-01 | 9,97E-01 | 0.0492(0.0058) | 1.0333(0.0151) | -0.0064(0.0051) |
| Child birth weight | 23202124 | 0.0012(0.1343) | 0,009 | 9,93E-01 | 9,97E-01 | 0.1091(0.0183) | 1.0051(0.0057) | -0.0039(0.0045) |
| Total lipids in small VLDL | 27005778 | 0.0011(0.1502) | 0,0072 | 9,94E-01 | 9,97E-01 | 0.1458(0.0374) | 0.9926(0.0081) | 0.0004(0.0045) |
| Nap during day | 0 | -0.0002(0.0634) | -0,0032 | 9,98E-01 | 9,99E-01 | 0.0793(0.0032) | 1.0254(0.0096) | -0.0062(0.0055) |
| Illnesses of father: Diabetes | 0 | -0.0002(0.1151) | -0,0014 | 9,99E-01 | 9,99E-01 | 0.0173(0.0022) | 1.0287(0.0072) | -0.0008(0.0049) |

Source: number denotes PMID; UKBB, UK Biobank; rg, genetic correlation; se, standard error; FDR, false discovery rate adjusted p-value; h² obs, narrow-sense heritability observed; h² int, narrow-sense heritability intercept, gcov_int, cross-trait intercept

**Supplementary Table 6. Runs of homozygosity within PD known genes/loci identified in the Spanish population**

| Chr | Bp1 | Bp2 | Kb | NSNP | PD Gene | Case | Control | Freq cases | Freq controls | P |
|---|---|---|---|---|---|---|---|---|---|---|
| 6 | 163235289 | 163452712 | 217,424 | 16 | *PARK2* | 68 | 18 | 0,0142 | 0,0059 | 0,0008 |
| 6 | 162802886 | 162847889 | 45,004 | 4 | *PARK2* | 65 | 18 | 0,0136 | 0,0059 | 0,0016 |
| 6 | 162683669 | 162697914 | 14,246 | 5 | *PARK2* | 63 | 19 | 0,0132 | 0,0062 | 0,0044 |
| 15 | 89680126 | 89921477 | 241,352 | 84 | *POLG1* | 33 | 7 | 0,0069 | 0,0023 | 0,0083 |
| 11 | 133897018 | 134123514 | 226,497 | 65 | *MIR4697* | 35 | 8 | 0,0073 | 0,0026 | 0,0093 |
| 16 | 20262776 | 20326407 | 63,632 | 4 | *SYT17* | 105 | 43 | 0,0220 | 0,0140 | 0,0149 |
| 6 | 162283824 | 162291767 | 7,944 | 2 | *PARK2* | 50 | 16 | 0,0105 | 0,0052 | 0,0187 |
| 6 | 162182126 | 162236682 | 54,557 | 8 | *PARK2* | 47 | 15 | 0,0098 | 0,0049 | 0,0227 |
| 17 | 77416940 | 77434466 | 17,527 | 5 | *DNAH17* | 35 | 10 | 0,0073 | 0,0033 | 0,0301 |
| 13 | 98832895 | 98902577 | 69,683 | 9 | *MBNL2* | 30 | 8 | 0,0063 | 0,0026 | 0,0345 |
| 6 | 162048933 | 162141896 | 92,964 | 12 | *PARK2* | 48 | 17 | 0,0100 | 0,0055 | 0,0440 |
| 17 | 75407883 | 75488782 | 80,9 | 4 | *DNAH17* | 31 | 9 | 0,0065 | 0,0029 | 0,0466 |
| 6 | 161811421 | 161830541 | 19,121 | 2 | *PARK2* | 54 | 21 | 0,0113 | 0,0068 | 0,0637 |
| 1 | 162439692 | 162513969 | 74,278 | 5 | *FCGR2A* | 30 | 10 | 0,0063 | 0,0033 | 0,0959 |
| 3 | 150063219 | 150159767 | 96,549 | 10 | *MED12L* | 38 | 14 | 0,0079 | 0,0046 | 0,0974 |
| 2 | 134632112 | 134633660 | 1,549 | 2 | *TMEM163* | 59 | 25 | 0,0123 | 0,0082 | 0,1002 |
| 12 | 122316185 | 122359542 | 43,358 | 5 | *HIP1R* | 350 | 196 | 0,0732 | 0,0639 | 0,1270 |
| 8 | 15684006 | 15734010 | 50,005 | 6 | *FGF20* | 46 | 20 | 0,0096 | 0,0065 | 0,1809 |
| 21 | 39828318 | 39861730 | 33,413 | 5 | *DYRK1A* | 28 | 11 | 0,0059 | 0,0036 | 0,2192 |
| 15 | 61966513 | 61994134 | 27,622 | 10 | *VPS13C* | 41 | 18 | 0,0086 | 0,0059 | 0,2233 |
| 1 | 204729758 | 204739515 | 9,758 | 2 | *RAB29* | 64 | 31 | 0,0134 | 0,0101 | 0,2353 |
| 8 | 22575108 | 22631572 | 56,465 | 5 | *SORBS3* | 33 | 14 | 0,0069 | 0,0046 | 0,2472 |
| 16 | 51557928 | 51772569 | 214,642 | 21 | *NOD2* | 38 | 17 | 0,0079 | 0,0055 | 0,2691 |
| 4 | 113570885 | 113609023 | 38,139 | 2 | *CAMK2D* | 62 | 31 | 0,0130 | 0,0101 | 0,3020 |
| 3 | 19190206 | 19401472 | 211,267 | 7 | *SATB1* | 345 | 202 | 0,0721 | 0,0659 | 0,3101 |
| 15 | 60992255 | 61016235 | 23,981 | 5 | *VPS13C* | 31 | 14 | 0,0065 | 0,0046 | 0,3456 |
| 3 | 18436286 | 18461878 | 25,593 | 2 | *SATB1* | 277 | 162 | 0,0579 | 0,0528 | 0,3658 |
| 12 | 40671282 | 40676801 | 5,52 | 13 | *LRRK2* | 215 | 124 | 0,0450 | 0,0404 | 0,3673 |
| 4 | 113429572 | 113454932 | 25,361 | 2 | *CAMK2D* | 60 | 31 | 0,0125 | 0,0101 | 0,3818 |
| 9 | 17736118 | 17760260 | 24,143 | 3 | *SH3GL2* | 55 | 29 | 0,0115 | 0,0095 | 0,4564 |
| 9 | 17772909 | 17792845 | 19,937 | 2 | *SH3GL2* | 53 | 28 | 0,0111 | 0,0091 | 0,4722 |
| 6 | 161655357 | 161672443 | 17,087 | 2 | *PARK2* | 63 | 34 | 0,0132 | 0,0111 | 0,4777 |
| 4 | 113281951 | 113303632 | 21,682 | 2 | *ANK2* | 74 | 41 | 0,0155 | 0,0134 | 0,5100 |
| 1 | 156169610 | 156237315 | 67,706 | 3 | *GBAP1* | 509 | 313 | 0,1064 | 0,1021 | 0,5662 |
| 6 | 111221374 | 111277745 | 56,372 | 2 | *FYN* | 257 | 155 | 0,0537 | 0,0506 | 0,5728 |
| 14 | 76179407 | 76489451 | 310,045 | 15 | *RPS6KL1* | 33 | 18 | 0,0069 | 0,0059 | 0,6823 |
| 14 | 74371655 | 74427993 | 56,339 | 4 | *RPS6KL1* | 144 | 89 | 0,0301 | 0,0290 | 0,8364 |
| 2 | 165081440 | 165098966 | 17,527 | 2 | *SCN3A* | 166 | 103 | 0,0347 | 0,0336 | 0,8410 |

| | | | | | | | | | | |
|---|---|---|---|---|---|---|---|---|---|---|
| 14 | 54837026 | 54857286 | 20,261 | 3 | *GCH1* | 47 | 28 | 0,0098 | 0,0091 | 0,8497 |
| 2 | 165134402 | 165159764 | 25,363 | 2 | *SCN3A* | 161 | 100 | 0,0337 | 0,0326 | 0,8513 |
| 8 | 23521248 | 23526463 | 5,216 | 2 | *BIN3* | 58 | 35 | 0,0121 | 0,0114 | 0,8595 |
| 10 | 120222457 | 120492868 | 270,412 | 17 | *BAG3* | 27 | 17 | 0,0056 | 0,0055 | 1 |

Chr; chromosome, Bp1; start base-pair position; Bp2; end base-pair position, KB; kilobases, NSNP; number of SNPs

**Supplementary Table 7. Significant functional associations for PD known/nominated genes via two-sample Mendelian randomization**

| Gene | Probe | beta | se | P-adjusted | Data source | Analyte | Top QTL SNP | Chr | Bp |
|---|---|---|---|---|---|---|---|---|---|
| *NSF* | ILMN_1680353 | -0,877869 | 0,19141 | 3,29E-04 | Expression (Westra et al., 2013) | Blood | rs183211 | 17 | 44788310 |
| *DGKQ* | ILMN_1793017 | -0,0965771 | 0,0358463 | 2,03E-02 | Expression (Westra et al., 2013) | Blood | rs11724804 | 4 | 965779 |
| *BST1* | ILMN_1770161 | -0,171468 | 0,0586181 | 2,03E-02 | Expression (Westra et al., 2013) | Blood | rs4698412 | 4 | 15737348 |
| *KANSL1* | ENSG00000120071 | -0,364123 | 0,064984 | 4,56E-06 | Expression (Qi et al., 2018) | Brain | rs2532233 | 17 | 44273218 |
| *KANSL1* | ENSG00000120071 | -0,364123 | 0,064984 | 4,56E-06 | Expression (Qi et al., 2018) | Brain | rs2532233 | 17 | 44273218 |
| *TMEM163* | ENSG00000152128 | 0,198515 | 0,093362 | 3,47E-03 | Expression (Qi et al., 2018) | Brain | rs512499 | 2 | 135221034 |
| *WNT3* | ENSG00000108379 | -0,0895389 | 0,0577794 | 3,47E-03 | Expression (Qi et al., 2018) | Brain | rs12452064 | 17 | 44868187 |
| *GAK* | ENSG00000178950 | 0,64831 | 0,175293 | 3,47E-03 | Expression (Qi et al., 2018) | Brain | rs11248057 | 4 | 906131 |
| *KAT8* | ENSG00000103510 | -0,162825 | 0,068547 | 7,27E-03 | Expression (Qi et al., 2018) | Brain | rs12597511 | 16 | 31145219 |
| *CD38* | ENSG00000004468 | -0,313842 | 0,116388 | 1,78E-02 | Expression (Qi et al., 2018) | Brain | rs11724635 | 4 | 15737101 |
| *DGKQ* | ENSG00000145214 | 0,0149581 | 0,0342918 | 3,40E-02 | Expression (Qi et al., 2018) | Brain | rs7441544 | 4 | 961987 |
| *HLA-DRB6* | ENSG00000229391 | -0,0652757 | 0,045313 | 3,62E-02 | Expression (Qi et al., 2018) | Brain | rs9271406 | 6 | 32587588 |
| *TMEM175* | ENSG00000127419 | -0,331655 | 0,11129 | 3,85E-02 | Expression (Qi et al., 2018) | Brain | rs60492656 | 4 | 875008 |
| *HLA-DQA1* | ENSG00000196735 | 0,0295384 | 0,0358717 | 4,22E-02 | Expression (Qi et al., 2018) | Brain | rs3828789 | 6 | 32635754 |
| *TMEM175* | ENSG00000127419.12 | -0,352456 | 0,125892 | 3,01E-02 | Expression (GTEx Consortium 2015) | Brain | rs60492656 | 4 | 875008 |
| *KAT8* | ENSG00000103510.15 | -0,109742 | 0,0470337 | 3,01E-02 | Expression (GTEx Consortium 2015) | Brain | rs12597511 | 16 | 31145219 |
| *HLA-DRB6* | ENSG00000229391.3 | -0,0652757 | 0,045313 | 4,13E-02 | Expression (GTEx Consortium 2015) | Brain | rs9271406 | 6 | 32587588 |
| *CTSB* | ENSG00000164733.16 | -0,305612 | 0,101981 | 4,13E-02 | Expression (GTEx Consortium 2015) | Brain | rs1692821 | 8 | 11699988 |
| *CTSB* | ENSG00000164733.16 | -0,305612 | 0,101981 | 4,13E-02 | Expression (GTEx Consortium 2015) | Brain | rs1692821 | 8 | 11699988 |
| *KANSL1* | cg20120463 | 0,258633 | 0,0453715 | 9,36E-07 | Methylation (Qi et al., 2018) | Brain | rs571792649 | 17 | 44283022 |
| *KANSL1* | cg20120463 | 0,258633 | 0,0453715 | 9,36E-07 | Methylation (Qi et al., 2018) | Brain | rs571792649 | 17 | 44283022 |
| *KANSL1* | cg13704117 | 0,267191 | 0,0461202 | 1,11E-06 | Methylation (Qi et al., 2018) | Brain | rs17661015 | 17 | 44192957 |
| *KANSL1* | cg13704117 | 0,267191 | 0,0461202 | 1,11E-06 | Methylation (Qi et al., 2018) | Brain | rs17661015 | 17 | 44192957 |
| *KANSL1* | cg01570182 | -0,273096 | 0,0493174 | 1,11E-06 | Methylation (Qi et al., 2018) | Brain | rs561504284 | 17 | 44340253 |
| *KANSL1* | cg01570182 | -0,273096 | 0,0493174 | 1,11E-06 | Methylation (Qi et al., 2018) | Brain | rs561504284 | 17 | 44340253 |
| *KANSL1* | cg11204664 | -0,294739 | 0,0543752 | 1,16E-06 | Methylation (Qi et al., 2018) | Brain | rs62073157 | 17 | 44334052 |
| *KANSL1* | cg11204664 | -0,294739 | 0,0543752 | 1,16E-06 | Methylation (Qi et al., 2018) | Brain | rs62073157 | 17 | 44334052 |
| *ARHGAP27* | cg27060340 | 0,162083 | 0,0271755 | 1,23E-06 | Methylation (Qi et al., 2018) | Brain | rs62065376 | 17 | 43521161 |
| *ARHGAP27* | cg18124852 | 0,253036 | 0,0463789 | 4,77E-06 | Methylation (Qi et al., 2018) | Brain | rs113575082 | 17 | 43524526 |
| *ARHGAP27* | cg21931864 | 0,309902 | 0,0608942 | 5,11E-06 | Methylation (Qi et al., 2018) | Brain | rs62065379 | 17 | 43525365 |
| *KANSL1* | cg14517863 | 0,398133 | 0,0821297 | 1,50E-05 | Methylation (Qi et al., 2018) | Brain | rs578109777 | 17 | 44319369 |
| *KANSL1* | cg14517863 | 0,398133 | 0,0821297 | 1,50E-05 | Methylation (Qi et al., 2018) | Brain | rs578109777 | 17 | 44319369 |
| *CRHR1* | cg15413793 | 0,416744 | 0,0862881 | 1,53E-05 | Methylation (Qi et al., 2018) | Brain | rs3108317 | 17 | 43719870 |
| *ARHGAP27* | cg22458832 | 0,260951 | 0,048371 | 2,39E-05 | Methylation (Qi et al., 2018) | Brain | rs62065379 | 17 | 43525365 |
| *CRHR1* | cg23762722 | 0,385354 | 0,0764167 | 2,39E-05 | Methylation (Qi et al., 2018) | Brain | rs62054387 | 17 | 43805525 |
| *KANSL1* | ch.17.41650897R | 0,457858 | 0,0994899 | 3,62E-05 | Methylation (Qi et al., 2018) | Brain | rs2668665 | 17 | 44290047 |
| *KANSL1* | ch.17.41650897R | 0,457858 | 0,0994899 | 3,62E-05 | Methylation (Qi et al., 2018) | Brain | rs2668665 | 17 | 44290047 |
| *ARHGAP27* | cg12346284 | 0,300318 | 0,0583272 | 4,11E-05 | Methylation (Qi et al., 2018) | Brain | rs62065379 | 17 | 43525365 |
| *KANSL1* | cg22192454 | 0,484445 | 0,106622 | 4,11E-05 | Methylation (Qi et al., 2018) | Brain | rs41374248 | 17 | 44120442 |
| *KANSL1* | cg22192454 | 0,484445 | 0,106622 | 4,11E-05 | Methylation (Qi et al., 2018) | Brain | rs41374248 | 17 | 44120442 |
| *KANSL1* | cg23647744 | 0,446944 | 0,100381 | 5,76E-05 | Methylation (Qi et al., 2018) | Brain | rs2732705 | 17 | 44351929 |
| *KANSL1* | cg23647744 | 0,446944 | 0,100381 | 5,76E-05 | Methylation (Qi et al., 2018) | Brain | rs2732705 | 17 | 44351929 |
| *KANSL1* | cg15633388 | 0,258667 | 0,0452142 | 7,11E-05 | Methylation (Qi et al., 2018) | Brain | rs545648705 | 17 | 44276330 |

| | | | | | | | | | |
|---|---|---|---|---|---|---|---|---|---|
| *KANSL1* | cg15633388 | 0,258667 | 0,0452142 | 7,11E-05 | Methylation (Qi et al., 2018) | Brain | rs545648705 | 17 | 44276330 |
| *KANSL1* | cg18391203 | -0,49959 | 0,115554 | 8,90E-05 | Methylation (Qi et al., 2018) | Brain | rs578109777 | 17 | 44319369 |
| *KANSL1* | cg18391203 | -0,49959 | 0,115554 | 8,90E-05 | Methylation (Qi et al., 2018) | Brain | rs578109777 | 17 | 44319369 |
| *ARHGAP27* | cg12548348 | 0,506038 | 0,118868 | 1,15E-04 | Methylation (Qi et al., 2018) | Brain | rs56212100 | 17 | 43501940 |
| *KANSL1* | cg22131547 | 0,188652 | 0,0718072 | 4,76E-04 | Methylation (Qi et al., 2018) | Brain | rs551913443 | 17 | 44274474 |
| *KANSL1* | cg22131547 | 0,188652 | 0,0718072 | 4,76E-04 | Methylation (Qi et al., 2018) | Brain | rs551913443 | 17 | 44274474 |
| *HLA-DRB5* | cg19575208 | -0,167448 | 0,0449188 | 9,71E-04 | Methylation (Qi et al., 2018) | Brain | rs477515 | 6 | 32569691 |
| *HLA-DRB5* | cg08845336 | -0,177885 | 0,0479992 | 1,03E-03 | Methylation (Qi et al., 2018) | Brain | rs477515 | 6 | 32569691 |
| *HLA-DRB5* | cg00211215 | -0,197642 | 0,0539661 | 1,18E-03 | Methylation (Qi et al., 2018) | Brain | rs477515 | 6 | 32569691 |
| *HLA-DRB5* | cg09139047 | -0,262544 | 0,0749921 | 2,13E-03 | Methylation (Qi et al., 2018) | Brain | rs477515 | 6 | 32569691 |
| *HLA-DRB5* | cg06032479 | -0,271721 | 0,0781605 | 2,26E-03 | Methylation (Qi et al., 2018) | Brain | rs477515 | 6 | 32569691 |
| *ARHGAP27* | cg14154330 | 0,313657 | 0,0904413 | 2,27E-03 | Methylation (Qi et al., 2018) | Brain | rs1230085 | 17 | 43501599 |
| *HLA-DRB5* | cg10765922 | -0,294047 | 0,086089 | 2,68E-03 | Methylation (Qi et al., 2018) | Brain | rs477515 | 6 | 32569691 |
| *HLA-DRB5* | cg26036029 | -0,274152 | 0,0834837 | 4,19E-03 | Methylation (Qi et al., 2018) | Brain | rs556025 | 6 | 32570880 |
| *DGKQ* | cg19408572 | -0,101842 | 0,0529988 | 6,05E-03 | Methylation (Qi et al., 2018) | Brain | rs4690163 | 4 | 975137 |
| *HLA-DRB5* | cg10632894 | -0,327684 | 0,103386 | 6,05E-03 | Methylation (Qi et al., 2018) | Brain | rs556025 | 6 | 32570880 |
| *HLA-DQA1* | cg21223191 | -0,0726812 | 0,0704359 | 6,16E-03 | Methylation (Qi et al., 2018) | Brain | rs3129762 | 6 | 32585755 |
| *DGKQ* | cg00741675 | -0,108155 | 0,0564098 | 6,24E-03 | Methylation (Qi et al., 2018) | Brain | rs4690163 | 4 | 975137 |
| *HLA-DRB5* | cg14075454 | -0,344767 | 0,110095 | 6,31E-03 | Methylation (Qi et al., 2018) | Brain | rs556025 | 6 | 32570880 |
| *GAK* | cg14517359 | 0,229644 | 0,0750298 | 7,45E-03 | Methylation (Qi et al., 2018) | Brain | rs3775127 | 4 | 886061 |
| *HLA-DRB5* | cg24242384 | -0,380517 | 0,124736 | 7,73E-03 | Methylation (Qi et al., 2018) | Brain | rs556025 | 6 | 32570880 |
| *HLA-DRB5* | cg17480035 | 0,0913986 | 0,0612456 | 7,73E-03 | Methylation (Qi et al., 2018) | Brain | rs9270931 | 6 | 32572547 |
| *TMEM175* | cg06157924 | 0,100912 | 0,0443705 | 7,80E-03 | Methylation (Qi et al., 2018) | Brain | rs2290403 | 4 | 942006 |
| *HLA-DQA1* | cg24143305 | -0,117974 | 0,0490467 | 9,56E-03 | Methylation (Qi et al., 2018) | Brain | rs9272219 | 6 | 32602269 |
| *TMEM163* | cg10711022 | -0,279012 | 0,0828581 | 1,12E-02 | Methylation (Qi et al., 2018) | Brain | rs10170773 | 2 | 135362505 |
| *IGSF9B* | cg12362517 | -0,166516 | 0,052661 | 1,22E-02 | Methylation (Qi et al., 2018) | Brain | rs11223627 | 11 | 133800769 |
| *IGSF9B* | cg25790212 | -0,194568 | 0,0628878 | 1,41E-02 | Methylation (Qi et al., 2018) | Brain | rs11223627 | 11 | 133800769 |
| *HLA-DQA1* | cg14353184 | 0,149566 | 0,060264 | 1,45E-02 | Methylation (Qi et al., 2018) | Brain | rs2858861 | 6 | 32580331 |
| *HLA-DQA1* | cg03787837 | 0,126192 | 0,0881042 | 1,45E-02 | Methylation (Qi et al., 2018) | Brain | rs9271406 | 6 | 32587588 |
| *HLA-DRB5* | cg00598125 | -0,126283 | 0,0535458 | 1,86E-02 | Methylation (Qi et al., 2018) | Brain | rs616187 | 6 | 32574305 |
| *HLA-DRB5* | cg01812434 | -0,0766439 | 0,0399939 | 3,40E-02 | Methylation (Qi et al., 2018) | Brain | rs9271492 | 6 | 32589122 |
| *HLA-DQA1* | cg04054303 | 0,183726 | 0,0960181 | 3,79E-02 | Methylation (Qi et al., 2018) | Brain | rs6928482 | 6 | 32626249 |

Chr; chromosome; Bp, base pair; se, standard error; P-adjusted, adjusted false discovery rate p value after performing Mendelian randomization between QTL resources (exposures) versus Spanish summary statistics (outcome)

**Supplementary Table 8. Genetic variants in PD and Parkinsonism related genes overrepresented in Spanish cases versus controls**

| Chr | Position | rs number | GENE | Variant | Codon | Zygosity | Ncases | Ncontrols | A1 | A2 | Freq cases | Freq controls | CHISQ | P | OR | SE | SNP probe NeuroChip ID |
|---|---|---|---|---|---|---|---|---|---|---|---|---|---|---|---|---|---|
| 16 | 46694455 | rs192419029 | *VPS35* | p.Leu774Met | CTG-ATG | Het | 1 | 0 | A | C | 0,0001045 | 0 | 0,6378 | 0,4245 | NA | NA | rs192419029 |
| 12 | 40645075 | rs78501232 | *LRRK2* | p.Glu334Lys | GAG-AAG | Het | 2 | 1 | A | G | 0,0003133 | 0 | 1,916 | 0,1663 | NA | NA | rs78501232 |
| 12 | 40657677 | rs79996249 | *LRRK2* | p.Lys544Glu | AAA-GAA | Het | 3 | 1 | G | A | 0,0003138 | 0,000164 | 0,3272 | 0,5673 | 1,914 | 1,13 | rs79996249 |
| 12 | 40677735 | rs200861727 | *LRRK2* | p.Arg767His | CGT-CAT | Het | 1 | 0 | A | G | 0,0001046 | 0 | 0,6378 | 0,4245 | NA | NA | seq-prion1987 |
| 12 | 40692290 | rs35808389 | *LRRK2* | p.Leu1114= | N/A | Het | 1 | 0 | G | A | 0,0001044 | 0 | 0,6383 | 0,4243 | NA | NA | newrs35808389 |
| 12 | 40696668 | rs281865047 | *LRRK2* | p.Ile1192Val | ATT-GTT | Het | 9 | 4 | G | A | 0,000941 | 0,0006555 | 0,3662 | 0,5451 | 1,436 | 0,59 | seq-prion2116 |
| 12 | 40702420 | rs17466213 | *LRRK2* | p.Ile1371Val | ATA-GTA | Het | 3 | 1 | G | A | 0,0003141 | 0,0001638 | 0,3293 | 0,5661 | 1,918 | 1,13 | rs17466213 |
| 12 | 40702426 | No | *LRRK2* | p.Val1373Met | GTG-ATG | Het | 2 | 0 | A | G | 0,0002088 | 0 | 1,277 | 0,2585 | NA | NA | seq-prion2169 |
| 12 | 40702947 | rs72546327 | *LRRK2* | p.Thr1410Met | ACG-ATG | Het | 3 | 0 | A | G | 0,0003135 | 0,0001638 | 0,3271 | 0,5673 | 1,914 | 1,13 | rs72546327 |
| 12 | 40707778 | rs35507033 | *LRRK2* | p.Arg1514Gln | CGA-CAA | Het | 109 | 67 | A | G | 0,01141 | 0,01108 | 0,03738 | 0,8467 | 1,031 | 0.15 | rs35507033 |
| 12 | 40713899 | rs35303786 | *LRRK2* | p.Met1646Thr | ATG-ACG | Hom | 59 | 24 | G | A | 0,03822 | 0,02355 | 23,86 | 1,04E-06 | 1,648 | 0,1 | LRRK2:NM_198578.3:c.4937T>C:p.(Met1646Thr) |
| | | | | | | Het | 227 | 86 | | | | | | | | | |
| 12 | 40715947 | No | *LRRK2* | p.Ser1761Arg | AGT-CGT | Het | 3 | 0 | C | A | 0,0003132 | 0 | 1,916 | 0,1663 | NA | NA | 12:40715947 |
| 12 | 40716270 | rs72547980 | *LRRK2* | p.Gln1823Lys | CAA-AAA | Het | 1 | 0 | A | C | 0,0001062 | 0 | 0,6321 | 0,4266 | NA | NA | rs72547980 |
| 12 | 40717058 | rs35602796 | *LRRK2* | p.Met1869Thr | ATG-ACG | Het | 5 | 1 | G | A | 0,000522 | 0,0001635 | 1,255 | 0,2625 | 3,194 | 1,03 | LRRK2:NM_198578.3:c.5606T>C:p.(Met1869Thr) |
| 12 | 40728833 | rs77428810 | *LRRK2* | p.Arg1941His | CGT-CAT | Het | 3 | 0 | A | G | 0,0003138 | 0 | 1,916 | 0,1662 | NA | NA | rs77428810 |
| 12 | 40734202 | rs34637584 | *LRRK2* | p.Gly2019Ser | GGC-AGC | Hom | 1 | 0 | A | G | 0,0143 | 0,001799 | 62,45 | 2,73E-15 | 8,05 | 0,26 | LRRK2:NM_198578.3:c.6055G>A:p.(Gly2019Ser) |
| | | | | | | Het | 134 | 12 | A | G | | | | | | | |
| 12 | 40740686 | rs33995883 | *LRRK2* | p.Asn2081Asp | AAT-GAT | Hom | 2 | 5 | G | A | 0,0186 | 0,01757 | 0,2194 | 0,6395 | 1,06 | 0,12 | rs33995883 |
| | | | | | | Het | 172 | 99 | G | A | | | | | | | |
| 12 | 40757328 | rs34778348 | *LRRK2* | p.Gly2385Arg | GGA-AGA | Het | 1 | 0 | A | G | 0,0001045 | 0 | 0,6365 | 0,425 | NA | NA | rs34778348 |
| 12 | 40757343 | rs79546190 | *LRRK2* | p.Val2390Met | GTG-ATG | Het | 1 | 0 | A | G | 0,0001048 | 0 | 0,6316 | 0,4268 | NA | NA | newrs79546190 |
| 12 | 40704236 | rs33939927 | *LRRK2* | p.Arg1441Gly | CGC-GGC | Het | 1 | 0 | C | G | 0,0001045 | 0 | 0,6365 | 0,425 | NA | NA | 12_40704236 |
| 1 | 155205022 | rs76071730 | *GBA* | p.His490Arg | CAT-CGT | Het | 1 | 0 | G | A | 0,0001044 | 0 | 0,6385 | 0,4243 | NA | NA | 1:155205022 |
| 1 | 155205551 | No | *GBA* | p.Val437Ile | GTC-ATC | Het | 1 | 0 | A | G | 0,0001044 | 0 | 0,6381 | 0,4244 | NA | NA | GBA_V437I |
| 1 | 155206172 | No | *GBA* | p.Leu363Pro | CTA-CCA | Het | 1 | 0 | G | A | 0,0001044 | 0 | 0,6386 | 0,4242 | NA | NA | GBA_L363P |
| 1 | 155207985 | rs74462743 | *GBA* | p.Gly234Glu | GGG-GAG | Het | 1 | 0 | A | G | 0,0001044 | 0 | 0,6386 | 0,4242 | NA | NA | rs74462743 |
| 1 | 155208006 | rs364897 | *GBA* | p.Asn227Ser | AAT-AGT | Het | 3 | 0 | G | A | 0,0003132 | 0 | 1,916 | 0,1663 | NA | NA | rs364897 |
| 1 | 155209824 | rs121908302 | *GBA* | p.Val54Leu | GTG-CTG | Het | 1 | 0 | A | C | 0,0001044 | 0 | 0,6385 | 0,4243 | NA | NA | rs121908302 |
| 1 | 155210451 | rs387906315 | *GBA* | p.Leu29Alafs*18 | CTT-GTT | Het | 2 | 0 | I | D | 0,0002088 | 0 | 1,277 | 0,2585 | NA | NA | Neuro-1:155210451 |
| 1 | 155205518 | rs1064651 | *GBA* | p.Asp448His | GAC-CAC | Het | 14 | 3 | G | C | 0,001464 | 0,0004892 | 3,279 | 0,07018 | 2,994 | 0,6 | Neuro-1:155205518 |
| 1 | 155206167 | rs2230288 | *GBA* | p.Glu365Lys | GAG-AAG | Het | 97 | 31 | A | G | 0,01043 | 0,005229 | 11,75 | 0,000609 | 2,005 | 0,2 | Only imputed |
| 1 | 155205634 | rs76763715 | *GBA* | p.Asn409Ser | AAC-AGC | Het | 53 | 12 | G | A | 0,005753 | 0,001958 | 12,65 | 0,0003763 | 2,949 | 0,3 | seq-rs76763715.2-B3 |
| 1 | 155210498 | rs150466109 | *GBA* | p.Lys13Arg | AAG-AGG | Het | 1 | 1 | G | A | 0,0002088 | 0 | 1,276 | 0,2586 | NA | NA | rs150466109 |
| 17 | 44087737 | rs63751392 | *MAPT* | p.Asn613del | AAT to --- | Het | 1 | 0 | D | I | 0,0001044 | 0 | 0,6383 | 0,4243 | NA | NA | rs63751392 |
| 6 | 161771195 | No | *PARK2* | p.Trp445Term | TGG-TAG | Hom | 3 | 0 | A | G | 0,001046 | 0 | 6,386 | 0,0115 | NA | NA | 6:161771195 |
| | | | | | | Het | 4 | 0 | A | G | | | | | | | |
| 6 | 161781161 | No | *PARK2* | p.Thr415Asn | ACC-AAC | Hom | 2 | 0 | A | C | 0,0007307 | 0 | 4,471 | 0,03448 | NA | NA | PARK2:NM_004562.2:c.1244C>A:p.(Thr415Asn) |
| | | | | | | Het | 3 | 0 | A | C | | | | | | | |
| 6 | 161990427 | No | *PARK2* | p.Ile298Ser | ATT-AGT | Het | 1 | 0 | C | A | 0,0001044 | 0 | 0,6383 | 0,4243 | NA | NA | PARK2:NM_004562.2:c.893T>G:p.(Ile298Ser) |
| 6 | 162206852 | rs34424986 | *PARK2* | p.Arg275Trp | CGG-TGG | Hom | 4 | 0 | A | G | 0,006054 | 0,003435 | 5,113 | 0,02374 | 1,767 | 0,25 | rs34424986 |
| | | | | | | Het | 49 | 22 | A | G | | | | | | | |
| 6 | 162206909 | rs150562946 | *PARK2* | p.Arg256Cys | CGC-TGC | Het | 5 | 3 | A | G | 0,0005218 | 0,000491 | 0,006952 | 0,9336 | 1,063 | 0,63 | rs150562946 |
| 6 | 162206917 | No | *PARK2* | p.Cys253Phe | TGC-TAC | Het | 1 | 0 | A | C | 0,0001044 | 0 | 0,6383 | 0,4243 | NA | NA | PARK2:NM_004562.2:c.758G>T:p.(Cys253Phe) |
| 6 | 162394367 | rs144032774 | *PARK2* | p.Arg234Gln | TGC-TTC | Het | 7 | 2 | A | G | 0,0007307 | 0,0003271 | 1,061 | 0,3031 | 2,235 | 0,78 | newrs144032774 |
| 6 | 162394433 | rs137853058 | *PARK2* | p.Cys212Tyr | TGT-TAT | Hom | 4 | 0 | A | G | 0,001252 | 0,0001636 | 5,347 | 0,02076 | 7,665 | 0,88 | PARK2:NM_004562.2:c.635G>A:p.(Cys212Tyr) |
| | | | | | | Het | 4 | 1 | A | G | | | | | | | |
| 6 | 162394434 | No | *PARK2* | p.Cys212Gly | TGT-GGT | Het | 4 | 0 | C | A | 0,0004176 | 0 | 2,554 | 0,11 | NA | NA | PARK2:NM_004562.2:c.634T>G:p.(Cys212Gly) |
| 6 | 162622179 | No | *PARK2* | p.Thr173Met | ACG-ATG | Hom | 3 | 0 | A | G | 0,001007 | 0 | 5,792 | 0,0161 | NA | NA | PARK2:NM_004562.2:c.518C>T:p.(Thr173Met) |
| | | | | | | Het | 2 | 0 | A | G | | | | | | | |
| 6 | 162683724 | rs55774500 | *PARK2* | p.Ala82Glu | GCA-GAA | Het | 8 | 4 | A | C | 0,001358 | 0,0009814 | 0,4378 | 0,5082 | 1,384 | 0,49 | rs55774500 |
| 6 | 162683747 | No | *PARK2* | p.Trp74Cysfs*8 | C-CCA | Hom | 1 | 0 | I | D | 0,001358 | 0,000654 | 1,708 | 0,1912 | 2,078 | 0,55 | indel.94051 |
| | | | | | C-CCA | Het | 11 | 4 | I | D | | | | | | | |

| | | | | | | | | | | | | | | | | | |
|---|---|---|---|---|---|---|---|---|---|---|---|---|---|---|---|---|---|
| 6 | 162864346 | rs137853059 | *PARK2* | p.Val56Glu | GTG-GAG | Het | 1 | 0 | A | T | 0,0001044 | 0 | 0,6387 | 0,4242 | NA | NA | PARK2:NM_004562.2:c.167T>A:p.(Val56Glu) |
| 6 | 162864377 | rs75860381 | *PARK2* | p.Ala46Thr | GCA-ACA | Het | 1 | 0 | A | G | 0,0001044 | 0 | 0,6389 | 0,4241 | NA | NA | rs75860381 |
| 6 | 162864388 | rs368134308 | *PARK2* | p.Arg42Pro | CGT-CCT | Het | 1 | 0 | A | G | 0,0002089 | 0,0001635 | 0,04016 | 0,8412 | 1,277 | 1,22 | rs368134308 |
| 6 | 162864388 | rs368134308 | *PARK2* | p.Arg42His | CGT-CAT | Het | 3 | 0 | A | G | | | | | | | |
| 6 | 162864411 | rs55777503 | *PARK2* | p.Gln34Argfs | CAG-CGG | Hom | 1 | 0 | D | I | 0,000731 | 0 | 4,473 | 0,03444 | NA | NA | rs55777503 |
| | | | | | | Het | 5 | 0 | D | I | | | | | | | |
| 6 | 162864412 | rs148851677 | *PARK2* | p.Gln34Arg | CAG-CGG | Het | 1 | 0 | G | A | 0,0001044 | 0 | 0,6389 | 0,4241 | NA | NA | rs148851677 |
| 6 | 162864461 | rs146288080 | *PARK2* | p.Asp18Asn | GAT-AAT | Het | 3 | 0 | A | G | 0,0003132 | 0 | 1,916 | 0,1663 | NA | NA | PARK2:NM_004562.2:c.52G>A:p.(Asp18Asn) |
| 1 | 20960135 | No | *PINK1* | p.Gly32Arg | GGG-AGG | Het | 6 | 0 | A | G | 0,0006687 | 0 | 3,534 | 0,06013 | NA | NA | PINK1:NM_032409.2:c.94G>A:p.(Gly32Arg) |
| 1 | 20960418 | No | *PINK1* | p.Gln126Pro | CAG-CCG | Het | 2 | 1 | C | A | 0,0002115 | 0,000164 | 0,04341 | 0,835 | 1,29 | 1,22 | PINK1:NM_032409.2:c.377A>C:p.(Gln126Pro) |
| 1 | 20964505 | rs143204084 | *PINK1* | p.Lys186Asn | AAG-AAC | Het | 1 | 0 | C | G | 0,0001045 | 0 | 0,6393 | 0,424 | NA | NA | rs143204084 |
| 1 | 20966446 | rs375601968 | *PINK1* | p.Arg246Gln | CGA-CAA | Het | 5 | 1 | A | G | 0,000522 | 0,0001636 | 1,255 | 0,2627 | 3,193 | 1,03 | PINK1:NM_032409.2:c.737G>A:p.(Arg246Gln) |
| 1 | 20971008 | rs372280083 | *PINK1* | p.Leu268Val | CTA-GTA | Het | 1 | 0 | G | C | 0,0001044 | 0 | 0,6383 | 0,4243 | NA | NA | PINK1:NM_032409.2:c.802C>G:p.(Leu268Val) |
| 1 | 20971132 | rs74315355 | *PINK1* | p.Gly309Asp | GGC-GAC | Hom | 1 | 0 | A | G | 0,0002092 | 0 | 1,28 | 0,258 | NA | NA | PINK1:NM_032409.2:c.926G>A:p.(Gly309Asp) |
| 1 | 20972199 | No | *PINK1* | p.Leu369Pro | CTT-CCT | Het | 1 | 0 | G | A | 0,0001044 | 0 | 0,6383 | 0,4243 | NA | NA | PINK1:NM_032409.2:c.1106T>C:p.(Leu369Pro) |
| 1 | 20977063 | No | *PINK1* | p.Asn542Ser | AAC-AGC | Het | 1 | 0 | G | A | 0,0001044 | 0 | 0,6377 | 0,4246 | NA | NA | PINK1:NM_032409.2:c.1625A>G:p.(Asn542Ser) |
| 15 | 89865073 | rs41549716 | *POLG1* | p.Ty831-Cys | TAT-TGT | Hom | 1 | 0 | G | A | 0,003966 | 0,00278 | 1,504 | 0,22 | 1,428 | 0,29 | rs41549716 |
| | | | | | | Het | 36 | 17 | G | A | | | | | | | |
| 15 | 89868870 | rs113994096 | *POLG1* | p.Pro587Leu | CCG-CTG | Hom | 1 | 0 | A | G | 0,002823 | 0,002619 | 0,05695 | 0,8114 | 1,078 | 0,31 | rs113994096 |
| | | | | | | Het | 25 | 16 | A | G | | | | | | | |
| 15 | 89868895 | rs556925652 | *POLG1* | p.Arg579Trp | CGG-TGG | Het | 1 | 0 | A | G | 0,0001044 | 0 | 0,6383 | 0,4243 | NA | NA | hrm_15_89868895_G_A |
| 15 | 89870429 | rs145843073 | *POLG1* | p.Asn468Asp | AAT-GAT | Het | 6 | 1 | G | A | 0,0006262 | 0,0001635 | 1,793 | 0,1806 | 3,831 | 1 | rs145843073 |
| 15 | 89870432 | rs113994095 | *POLG1* | p.Ala467Thr | GCC-ACC | Het | 2 | 1 | A | G | 0,0002088 | 0,0001635 | 0,04002 | 0,8414 | 1,277 | 1,22 | rs113994095 |
| 15 | 89873472 | rs113994093 | *POLG1* | p.Arg232His | CGT-CAT | Het | 1 | 0 | A | G | 0,0001046 | 0 | 0,6383 | 0,4243 | NA | NA | rs113994093 |
| 1 | 17313685 | rs150748722 | *ATP13A2* | p.Arg980His | CGC-CAC | Het | 1 | 0 | A | G | 0,0001063 | 0 | 0,6333 | 0,4262 | NA | NA | rs150748722 |
| 1 | 17314656 | rs55708915 | *ATP13A2* | p.Ile946Phe | ATC-TTC | Het | 10 | 5 | A | T | 0,001044 | 0,0008178 | 0,1997 | 0,655 | 1,277 | 0,54 | rs55708915 |
| 22 | 38508219 | rs121908680 | *FBXO7* | p.Tyr790Term | TAT-TAG | Het | 1 | 0 | C | A | 0,0001045 | 0 | 0,6375 | 0,4246 | NA | NA | rs121908680 |
| 22 | 38511664 | rs387906863 | *FBXO7* | p.Arg635Gln | CGA-CAA | Het | 1 | 0 | A | G | 0,0001044 | 0 | 0,6383 | 0,4243 | NA | NA | qs387906863 |
| 22 | 38528888 | rs11570680 | *FBXO7* | p.Ala343Thr | GCG-ACG | Het | 119 | 65 | A | G | 0,01243 | 0,01063 | 1,051 | 0,3053 | 1,172 | 0,15 | rs11570680 |
| 2 | 74588717 | rs72466496 | *DCTN1* | p.Thr1249Ile | ACC-ATC | Het | 8 | 2 | A | G | 0,0008349 | 0,000327 | 1,513 | 0,2187 | 2,554 | 0,76 | rs72466496 |
| 2 | 74593101 | rs145130328 | *DCTN1* | p.Ile935Met | ATC-ATG | Het | 1 | 0 | G | C | 0,0001044 | 0 | 0,6383 | 0,4243 | NA | NA | rs145130328 |
| 2 | 74594023 | rs121909344 | *DCTN1* | p.Arg785Trp | CGG-TGG | Het | 15 | 4 | A | G | 0,001565 | 0,000654 | 2,565 | 0,1092 | 2,396 | 0,54 | rs121909344 |

**IPDGC consortium members and affiliations:**

**United Kingdom:**
Alastair J Noyce (Preventive Neurology Unit, Wolfson Institute of Preventive Medicine, QMUL, London, UK and Department of Molecular Neuroscience, UCL, London, UK),
Rauan Kaiyrzhanov (Department of Molecular Neuroscience, UCL Institute of Neurology, London, UK),
Ben Middlehurst (Institute of Translational Medicine, University of Liverpool, Liverpool, UK),
Demis A Kia (UCL Genetics Institute; and Department of Molecular Neuroscience, UCL Institute of Neurology, London, UK),
Manuela Tan (Department of Clinical Neuroscience, University College London, London, UK),
Henry Houlden (Department of Molecular Neuroscience, UCL Institute of Neurology, London, UK),
Huw R Morris (Department of Clinical Neuroscience, University College London, London, UK),
Helene Plun-Favreau (Department of Molecular Neuroscience, UCL Institute of Neurology, London, UK),
Peter Holmans (Biostatistics & Bioinformatics Unit, Institute of Psychological Medicine and Clinical Neuroscience, MRC Centre for Neuropsychiatric Genetics & Genomics, Cardiff, UK),
John Hardy (Department of Molecular Neuroscience, UCL Institute of Neurology, London, UK),
Daniah Trabzuni (Department of Molecular Neuroscience, UCL Institute of Neurology, London, UK; Department of Genetics, King Faisal Specialist Hospital and Research Centre, Riyadh, 11211 Saudi Arabia),
Jose Bras (UK Dementia Research Institute at UCL and Department of Molecular Neuroscience, UCL Institute of Neurology, London, UK),
John Quinn PhD (Institute of Translational Medicine, University of Liverpool, Liverpool, UK),
Kin Y Mok (Department of Molecular Neuroscience, UCL Institute of Neurology, London, UK),
Kerri J. Kinghorn (Institute of Healthy Ageing, University College London, London, UK),
Kimberley Billingsley (Institute of Translational Medicine, University of Liverpool, Liverpool, UK),
Nicholas W Wood (UCL Genetics Institute; and Department of Molecular Neuroscience, UCL Institute of Neurology, London, UK),
Patrick Lewis (University of Reading, Reading, UK),
Sebastian Schreglmann (Department of Molecular Neuroscience, UCL Institute of Neurology, London, UK),
Rita Guerreiro (UK Dementia Research Institute at UCL and Department of Molecular Neuroscience, UCL Institute of Neurology, London, UK),
Ruth Lovering (University College London, London, UK),
Lea R'Bibo (Department of Molecular Neuroscience, UCL Institute of Neurology, London, UK),
Claudia Manzoni (University of Reading, Reading, UK),
Mie Rizig (Department of Molecular Neuroscience, UCL Institute of Neurology, London, UK),
Mina Ryten (Department of Molecular Neuroscience, UCL Institute of Neurology, London, UK),
Sebastian Guelfi (Department of Molecular Neuroscience, UCL Institute of Neurology, London, UK),
Valentina Escott-Price (MRC Centre for Neuropsychiatric Genetics and Genomics, Cardiff University School of Medicine, Cardiff, UK),
Viorica Chelban (Department of Molecular Neuroscience, UCL Institute of Neurology, London, UK),
Thomas Foltynie (UCL Institute of Neurology, London, UK),
Nigel Williams (MRC Centre for Neuropsychiatric Genetics and Genomics, Cardiff, UK), Karen E. Morrison (Faculty of Medicine, University of Southampton, UK), Carl Clarke (University of Birmingham, Birmingham, UK and Sandwell and West Birmingham Hospitals NHS Trust, Birmingham, UK).

**France:**
Alexis Brice (Institut du Cerveau et de la Moelle épinière, ICM, Inserm U 1127, CNRS, UMR 7225, Sorbonne Universités, UPMC University Paris 06, UMR S 1127, AP-HP, Pitié-Salpêtrière Hospital, Paris, France),
Fabrice Danjou (Institut du Cerveau et de la Moelle épinière, ICM, Inserm U 1127, CNRS, UMR 7225, Sorbonne Universités, UPMC University Paris 06, UMR S 1127, AP-HP, Pitié-Salpêtrière Hospital, Paris, France),
Suzanne Lesage (Institut du Cerveau et de la Moelle épinière, ICM, Inserm U 1127, CNRS, UMR 7225, Sorbonne Universités, UPMC University Paris 06, UMR S 1127, AP-HP, Pitié-Salpêtrière Hospital, Paris, France),

Jean-Christophe Corvol (Institut du Cerveau et de la Moelle épinière, ICM, Inserm U 1127, CNRS, UMR 7225, Sorbonne Universités, UPMC University Paris 06, UMR S 1127, Centre d'Investigation Clinique Pitié Neurosciences CIC-1422, AP-HP, Pitié-Salpêtrière Hospital, Paris, France),
Maria Martinez (INSERM UMR 1220; and Paul Sabatier University, Toulouse, France),

**Germany:**
Claudia Schulte (Department for Neurodegenerative Diseases, Hertie Institute for Clinical Brain Research, University of Tübingen, and DZNE, German Center for Neurodegenerative Diseases, Tübingen, Germany),
Kathrin Brockmann (Department for Neurodegenerative Diseases, Hertie Institute for Clinical Brain Research, University of Tübingen, and DZNE, German Center for Neurodegenerative Diseases, Tübingen, Germany),
Javier Simón-Sánchez (Department for Neurodegenerative Diseases, Hertie Institute for Clinical Brain Research, University of Tübingen, and DZNE, German Center for Neurodegenerative Diseases, Tübingen, Germany),
Peter Heutink (DZNE, German Center for Neurodegenerative Diseases and Department for Neurodegenerative Diseases, Hertie Institute for Clinical Brain Research, University of Tübingen, Tübingen, Germany),
Patrizia Rizzu (DZNE, German Center for Neurodegenerative Diseases),
Manu Sharma (Centre for Genetic Epidemiology, Institute for Clinical Epidemiology and Applied Biometry, University of Tubingen, Germany),
Thomas Gasser (Department for Neurodegenerative Diseases, Hertie Institute for Clinical Brain Research, and DZNE, German Center for Neurodegenerative Diseases, Tübingen, Germany),

**United States of America:**
Aude Nicolas (Laboratory of Neurogenetics, National Institute on Aging, Bethesda, MD, USA),
Mark R Cookson (Laboratory of Neurogenetics, National Institute on Aging, Bethesda, USA),
Sara Bandres-Ciga (Laboratory of Neurogenetics, National Institute on Aging, Bethesda, MD, USA),
Cornelis Blauwendraat (National Institute on Aging and National Institute of Neurological Disorders and Stroke, USA),
David W. Craig (Department of Translational Genomics, Keck School of Medicine, University of Southern California, Los Angeles, USA),
Faraz Faghri (Laboratory of Neurogenetics, National Institute on Aging, Bethesda, USA; Department of Computer Science, University of Illinois at Urbana-Champaign, Urbana, IL, USA),
J Raphael Gibbs (Laboratory of Neurogenetics, National Institute on Aging, National Institutes of Health, Bethesda, MD, USA),
Dena G. Hernandez (Laboratory of Neurogenetics, National Institute on Aging, Bethesda, MD, USA),
Kendall Van Keuren-Jensen (Neurogenomics Division, TGen, Phoenix, AZ USA),
Joshua M. Shulman (Departments of Neurology, Neuroscience, and Molecular & Human Genetics, Baylor College of Medicine, Houston, Texas, USA; Jan and Dan Duncan Neurological Research Institute, Texas Children's Hospital, Houston, Texas, USA),
Hirotaka Iwaki (Laboratory of Neurogenetics, National Institute on Aging, Bethesda, MD, USA),
Hampton L. Leonard (Laboratory of Neurogenetics, National Institute on Aging, Bethesda, MD, USA),
Mike A. Nalls (Laboratory of Neurogenetics, National Institute on Aging, Bethesda, USA; CEO/Consultant Data Tecnica International, Glen Echo, MD, USA),
Laurie Robak (Baylor College of Medicine, Houston, Texas, USA),
Steven Lubbe (Ken and Ruth Davee Department of Neurology, Northwestern University Feinberg School of Medicine, Chicago, IL, USA),
Steven Finkbeiner (Departments of Neurology and Physiology, University of California, San Francisco; Gladstone Institute of Neurological Disease; Taube/Koret Center for Neurodegenerative Disease Research, San Francisco, CA, USA),
Niccolo E. Mencacci (Northwestern University Feinberg School of Medicine, Chicago, IL, USA),
Codrin Lungu (National Institutes of Health Division of Clinical Research, NINDS, National Institutes of Health, Bethesda, MD, USA),
Andrew B Singleton (Laboratory of Neurogenetics, National Institute on Aging, Bethesda, MD, USA),

Sonja W. Scholz (Neurodegenerative Diseases Research Unit, National Institute of Neurological Disorders and Stroke, Bethesda, MD, USA),
Xylena Reed (Laboratory of Neurogenetics, National Institute on Aging, Bethesda, MD, USA).
Roy N. Alcalay (Department of Neurology, College of Physicians and Surgeons, Columbia University Medical Center, New York, NY, USA, Taub Institute for Research on Alzheimer's Disease and the Aging Brain, College of Physicians and Surgeons, Columbia University Medical Center, New York, NY, USA)

**Canada:**
Ziv Gan-Or (Montreal Neurological Institute and Hospital, Department of Neurology & Neurosurgery, Department of Human Genetics, McGill University, Montréal, QC, H3A 0G4, Canada),
Guy A. Rouleau (Montreal Neurological Institute and Hospital, Department of Neurology & Neurosurgery, Department of Human Genetics, McGill University, Montréal, QC, H3A 0G4, Canada), Lynne Krohn (Montreal Neurological Institute and Hospital, Department of Neurology & Neurosurgery, Department of Human Genetics, McGill University, Montréal, QC, H3A 0G4, Canada)

**The Netherlands:**
Jacobus J van Hilten (Department of Neurology, Leiden University Medical Center, Leiden, Netherlands),
Johan Marinus (Department of Neurology, Leiden University Medical Center, Leiden, Netherlands)

**Spain:**
Astrid D. Adarmes-Gómez (Instituto de Biomedicina de Sevilla (IBiS), Hospital Universitario Virgen del Rocío/CSIC/Universidad de Sevilla, Seville),
Miquel Aguilar (Fundació Docència i Recerca Mútua de Terrassa and Movement Disorders Unit, Department of Neurology, University Hospital Mutua de Terrassa, Terrassa, Barcelona.),
Ignacio Alvarez (Fundació Docència i Recerca Mútua de Terrassa and Movement Disorders Unit, Department of Neurology, University Hospital Mutua de Terrassa, Terrassa, Barcelona.),
Victoria Alvarez (Hospital Universitario Central de Asturias, Oviedo),
Francisco Javier Barrero (Hospital Universitario San Cecilio de Granada, Universidad de Granada),
Jesús Alberto Bergareche Yarza (Instituto de Investigación Sanitaria Biodonostia, San Sebastián),
Inmaculada Bernal-Bernal (Instituto de Biomedicina de Sevilla (IBiS), Hospital Universitario Virgen del Rocío/CSIC/Universidad de Sevilla, Seville),
Marta Blazquez (Hospital Universitario Central de Asturias, Oviedo),
Marta Bonilla-Toribio (Instituto de Biomedicina de Sevilla (IBiS), Hospital Universitario Virgen del Rocío/CSIC/Universidad de Sevilla, Seville),
Juan A. Botía (Universidad de Murcia, Murcia),
María Teresa Boungiorno (Fundació Docència i Recerca Mútua de Terrassa and Movement Disorders Unit, Department of Neurology, University Hospital Mutua de Terrassa, Terrassa, Barcelona.)
Dolores Buiza-Rueda (Instituto de Biomedicina de Sevilla (IBiS), Hospital Universitario Virgen del Rocío/CSIC/Universidad de Sevilla, Seville),
Ana Cámara (Hospital Clinic de Barcelona),
Fátima Carrillo (Instituto de Biomedicina de Sevilla (IBiS), Hospital Universitario Virgen del Rocío/CSIC/Universidad de Sevilla, Seville),
Mario Carrión-Claro (Instituto de Biomedicina de Sevilla (IBiS), Hospital Universitario Virgen del Rocío/CSIC/Universidad de Sevilla, Seville),
Debora Cerdan (Hospital General de Segovia, Segovia),
Jordi Clarimón (Memory Unit, Department of Neurology, IIB Sant Pau, Hospital de la Santa Creu i Sant Pau, Universitat Autònoma de Barcelona and Centro de Investigación Biomédica en Red en Enfermedades Neurodegenerativas (CIBERNED), Madrid)
Yaroslau Compta (Hospital Clinic de Barcelona),
Beatríz de la Casa (Hospital Gregorio Marañon, Madrid)
Monica Diez-Fairen (Fundació Docència i Recerca Mútua de Terrassa and Movement Disorders Unit, Department of Neurology, University Hospital Mutua de Terrassa, Terrassa, Barcelona.),
Oriol Dols-Icardo (Memory Unit, Department of Neurology, IIB Sant Pau, Hospital de la Santa Creu i Sant Pau, Universitat Autònoma de Barcelona, Barcelona, and Centro de Investigación Biomédica en Red en Enfermedades Neurodegenerativas (CIBERNED), Madrid),

Jacinto Duarte (Hospital General de Segovia, Segovia),
Raquel Duran (Centro de Investigacion Biomedica, Universidad de Granada, Granada),
Francisco Escamilla-Sevilla (Hospital Universitario Virgen de las Nieves, Instituto de Investigación Biosanitaria de Granada, Granada),
Mario Ezquerra (Hospital Clinic de Barcelona)
Cici Feliz (Departmento de Neurologia, Instituto de Investigación Sanitaria Fundación Jiménez Díaz, Madrid, Spain),
Manel Fernández (Hospital Clinic de Barcelona),
Rubén Fernández-Santiago (Hospital Clinic de Barcelona),
Ciara Garcia (Hospital Universitario Central de Asturias, Oviedo),
Pedro García-Ruiz (Instituto de Investigación Sanitaria Fundación Jiménez Díaz, Madrid),
Pilar Gómez-Garre (Instituto de Biomedicina de Sevilla (IBiS), Hospital Universitario Virgen del Rocío/CSIC/Universidad de Sevilla, Seville),
Maria Jose Gomez Heredia (Hospital Universitario Virgen de la Victoria, Malaga),
Isabel Gonzalez-Aramburu (Hospital Universitario Marqués de Valdecilla-IDIVAL, Santander) ,
Ana Gorostidi Pagola (Instituto de Investigación Sanitaria Biodonostia, San Sebastián),
Janet Hoenicka (Institut de Recerca Sant Joan de Déu, Barcelona),
Jon Infante (Hospital Universitario Marqués de Valdecilla-IDIVAL and University of Cantabria, Santander, and Centro de Investigación Biomédica en Red en Enfermedades Neurodegenerativas (CIBERNED),
Silvia Jesús (Instituto de Biomedicina de Sevilla (IBiS), Hospital Universitario Virgen del Rocío/CSIC/Universidad de Sevilla, Seville),
Adriano Jimenez-Escrig (Hospital Universitario Ramón y Cajal, Madrid),
Jaime Kulisevsky (Movement Disorders Unit, Department of Neurology, IIB Sant Pau, Hospital de la Santa Creu i Sant Pau, Universitat Autònoma de Barcelona, Barcelona, and Centro de Investigación Biomédica en Red en Enfermedades Neurodegenerativas (CIBERNED)),
Miguel A. Labrador-Espinosa (Instituto de Biomedicina de Sevilla (IBiS), Hospital Universitario Virgen del Rocío/CSIC/Universidad de Sevilla, Seville),
Jose Luis Lopez-Sendon (Hospital Universitario Ramón y Cajal, Madrid),
Adolfo López de Munain Arregui (Instituto de Investigación Sanitaria Biodonostia, San Sebastián),
Daniel Macias (Instituto de Biomedicina de Sevilla (IBiS), Hospital Universitario Virgen del Rocío/CSIC/Universidad de Sevilla, Seville),
Irene Martínez Torres (Department of Neurology, Instituto de Investigación Sanitaria La Fe, Hospital Universitario y Politécnico La Fe, Valencia),
Juan Marín (Movement Disorders Unit, Department of Neurology, IIB Sant Pau, Hospital de la Santa Creu i Sant Pau, Universitat Autònoma de Barcelona, Barcelona, and Centro de Investigación Biomédica en Red en Enfermedades Neurodegenerativas (CIBERNED)),
Maria Jose Marti (Hospital Clinic Barcelona),
Juan Carlos Martínez-Castrillo (Instituto Ramón y Cajal de Investigación Sanitaria, Hospital Universitario Ramón y Cajal, Madrid),
Carlota Méndez-del-Barrio (Instituto de Biomedicina de Sevilla (IBiS), Hospital Universitario Virgen del Rocío/CSIC/Universidad de Sevilla, Seville),
Manuel Menéndez González (Hospital Universitario Central de Asturias, Oviedo),
Marina Mata (Department of Neurology, Hospital Universitario Infanta Sofía, Madrid, Spain)
Adolfo Mínguez (Hospital Universitario Virgen de las Nieves, Granada, Instituto de Investigación Biosanitaria de Granada),
Pablo Mir (Instituto de Biomedicina de Sevilla (IBiS), Hospital Universitario Virgen del Rocío/CSIC/Universidad de Sevilla, Seville),
Elisabet Mondragon Rezola (Instituto de Investigación Sanitaria Biodonostia, San Sebastián),
Esteban Muñoz (Hospital Clinic Barcelona),
Javier Pagonabarraga (Movement Disorders Unit, Department of Neurology, IIB Sant Pau, Hospital de la Santa Creu i Sant Pau, Universitat Autònoma de Barcelona, Barcelona, and Centro de Investigación Biomédica en Red en Enfermedades Neurodegenerativas (CIBERNED)),
Berta Pascual-Sedano (Unidad de Trastornos del Movimiento, Hospital Sant Pau, Barcelona)
Pau Pastor (Fundació Docència i Recerca Mútua de Terrassa and Movement Disorders Unit, Department of Neurology, University Hospital Mutua de Terrassa, Terrassa, Barcelona.),
Francisco Perez Errazquin (Hospital Universitario Virgen de la Victoria, Malaga),

Teresa Periñán-Tocino (Instituto de Biomedicina de Sevilla (IBiS), Hospital Universitario Virgen del Rocío/CSIC/Universidad de Sevilla, Seville),
Javier Ruiz-Martínez (Hospital Universitario Donostia, Instituto de Investigación Sanitaria Biodonostia, San Sebastián),
Clara Ruz (Centro de Investigacion Biomedica, Universidad de Granada, Granada),
Antonio Sanchez Rodriguez (Hospital Universitario Marqués de Valdecilla-IDIVAL, Santander),
María Sierra (Hospital Universitario Marqués de Valdecilla-IDIVAL, Santander),
Esther Suarez-Sanmartin (Hospital Universitario Central de Asturias, Oviedo),
Cesar Tabernero (Hospital General de Segovia, Segovia),
Juan Pablo Tartari (Fundació Docència i Recerca Mútua de Terrassa and Movement Disorders Unit, Department of Neurology, University Hospital Mutua de Terrassa, Terrassa, Barcelona),
Cristina Tejera-Parrado (Instituto de Biomedicina de Sevilla (IBiS), Hospital Universitario Virgen del Rocío/CSIC/Universidad de Sevilla, Seville),
Eduard Tolosa (Hospital Clinic Barcelona),
Francesc Valldeoriola (Hospital Clinic Barcelona),
Laura Vargas-González (Instituto de Biomedicina de Sevilla (IBiS), Hospital Universitario Virgen del Rocío/CSIC/Universidad de Sevilla, Seville),
Lydia Vela (Department of Neurology, Hospital Universitario Fundación Alcorcón, Madrid),
Francisco Vives (Centro de Investigacion Biomedica, Universidad de Granada, Granada).

**Austria:**
Alexander Zimprich (Department of Neurology, Medical University of Vienna, Austria)

**Norway:**
Lasse Pihlstrom (Department of Neurology, Oslo University Hospital, Oslo, Norway),
Mathias Toft (Department of Neurology and Institute of Clinical Medicine, Oslo University Hospital, Oslo, Norway)

**Estonia**:
Sulev Koks (Department of Pathophysiology, University of Tartu, Tartu, Estonia; Department of Reproductive Biology, Estonian University of Life Sciences, Tartu, Estonia),
Pille Taba (Department of Neurology and Neurosurgery, University of Tartu, Tartu, Estonia)

**Israel:**
Sharon Hassin-Baer (The Movement Disorders Institute, Department of Neurology and Sagol Neuroscience Center, Chaim Sheba Medical Center, Tel-Hashomer, 5262101, Ramat Gan, Israel, Sackler Faculty of Medicine, Tel Aviv University, Tel Aviv, Israel)
